# Ecophysiological niche expansion driven by biofilm lifestyle in an archaeal soil nitrifier

**DOI:** 10.64898/2026.05.09.724019

**Authors:** Thomas Pribasnig, Maximilian Dreer, Zhen-Hao Luo, Andrea Malits, Logan H. Hodgskiss, Christa Schleper

**Affiliations:** Department of Functional and Evolutionary Ecology, Archaea Biology and Ecogenomics Unit, University of Vienna, Djerassiplatz 1, 1030 Vienna, Austria; Thomas Pribasnig, Center for Microbial Communities, Department of Chemistry and Bioscience, Aalborg University, Aalborg, Denmark; Maximilian Dreer, Architecture and Dynamics of Biological Macromolecules, Institut Pasteur, Université Paris Cité, Paris, France

**Keywords:** ammonia oxidizing archaea, Nitrososphaera viennensis, nitrification inhibition, acidic soils

## Abstract

As key drivers of nitrification, ammonia-oxidizing archaea (AOA) play a central role in the global nitrogen cycle and contribute significantly to the emissions of the potent greenhouse gas nitrous oxide (N_2_O). However, the ecological implications of AOA growth as biofilms, remain poorly understood. Since nitrite production can be used to follow cellular activities directly we were able to compare biofilms with planktonic cells of the terrestrial model AOA *Nitrososphaera viennensis* at ecologically and agriculturally relevant conditions. Biofilms were more resistant across nearly all tested conditions and remained active at lower temperatures, acidic pH, and high ammonium concentrations. Collectively, activities in biofilm help reconcile discrepancies between earlier laboratory and environmental observations of soil AOA. Additionally, biofilms showed a high general resilience and lowered sensitivities to nitrification inhibitors. Although *in situ* biofilms grown in microrespiratory chambers exhibited activity and ammonia affinity similar to planktonic cells, biofilm cultures produced only half as much N₂O. The enhanced fitness of biofilms across all tested conditions vastly expands the potential ecophysiological niche of AOA and supports the hypothesis that biofilm growth represents the *in situ* phenotype of AOA in soil environments.

## Introduction

Ammonia-oxidizing archaea (AOA) are key players in the global nitrogen cycle by catalyzing the first step of nitrification, the oxidation of ammonia to nitrite^1–4^. Their activity affects nitrogen availability in natural and agricultural systems and contributes to the emission of nitrous oxide (N_2_O)^5–9^, a potent greenhouse gas. Based on molecular studies, AOA have a remarkably broad environmental distribution, being detected in virtually all oxic environments, including marine^10,11^, soil^12,13^, sediment^10^, hot spring^14,15^, acidic^16^, and arctic ecosystems^17–19^. Due to their high numbers in marine and terrestrial ecosystems, they range among the most abundant prokaryotes on Earth^20^.

Despite their ubiquity, the physiological traits that allow AOA to thrive across such diverse environments remain poorly understood. For example, although many AOA clades are found across a broad soil pH range (3.5–9.0)^21^, the basis for this broad tolerance (compared to their bacterial counterparts) is not reflected by the cultured organisms. Suggested possibilities include a coupling of AOA evolution to pH adaptation^22,23^, the emergence of few distinct low pH ecotypes^24^, and a coupling of nitrite removal with nitrite-oxidizing bacteria *in situ* ^25^, but these do not account for the existence of seemingly neutrophilic clades in acidic environments^16,21,24^. Similarly, many habitats in which AOA are found experience temperatures well below the growth ranges of cultured AOA. For example, while the average annual soil temperature at 5 cm depth in central Europe was ∼12 °C between 2010-2020, reaching ∼22 °C in summer^26^, no growth below 28°C and 30°C has been observed in the two terrestrial clades of AOA represented by *Nitrososphaera viennensis*^27^ and *Nitrosocosmicus franklandianus*^28^. The high temperature optima of AOA have been attributed to a secondary adaptation to mesophily^29^, but how they remain active under seemingly unfavorable low temperatures in the environment remains to be explored. While AOA are generally considered the dominant ammonia oxidizers in oligotrophic and acidic environments^11–13,16,30^, ammonia-oxidizing bacteria (AOB) are often thought to take over under high ammonia conditions, such as those after fertilizer application^31–33^. This aligns with the observation that AOA, with the exception of *Nitrosocosmicus* spp., display ammonia affinities one to two orders of magnitude greater than those of AOB^34–37^. However, AOA are nevertheless ubiquitously found at high numbers in agricultural environments under high ammonia levels even outnumbering AOB ^12,13,38,39^. Seemingly counterintuitive, many cultivated AOA strains display susceptibility to higher concentrations of ammonia when compared to AOB^27,28,40,41^. Additionally, nitrification inhibitors (NI) are used in agricultural soils to improve the low nitrogen use efficiency (NUE) by plants^42–44^, yet the reported effectiveness of these compounds varies between soils and pure-culture based studies^45–48^ which impairs the search of novel, less biohazardous inhibitors. Taken together, these apparent contradictions underscore a central knowledge gap. We lack a mechanistic understanding of how AOA persist and remain active in the environments under conditions that appear unfavorable based on pure-culture physiology.

Biofilm formation offers a compelling explanation. Biofilms are hypothesized to be the predominant microbial lifestyle in soil environments^49^, while offering protection from environmental stress^50,51^, buffering against pH^52,53^ and temperature stress^54,55^, and enabling resilience to fluctuating conditions^56^. If AOA form biofilms with similar traits, this could resolve many of the discrepancies outlined above: mitigating ammonia toxicity, reducing sensitivity to nitrification inhibitors, and sustaining activity under suboptimal pH or temperature. Until recently, this hypothesis could not be systematically tested, but we have now demonstrated reproducible and measurable biofilm formation and activity in six ammonia-oxidizing archaeal (AOA) strains from terrestrial and marine clades^57^. In this comparison, the soil genera *Nitrosocosmicus* and *Nitrososphaera* formed the strongest biofilms and showed distinct colonization strategies^57^. Having established this model system for AOA biofilm growth^57^, we are now able to directly assess how this lifestyle could contribute to the ecophysiology of AOA.

By comparing and quantifying the nitrification activity of biofilms and planktonic cultures of equivalent cell number, we show here that biofilms of the model AOA *Nitrososphaera viennensis*^27,58^ confer remarkable resistance and resilience across a variety of different ecologically and agriculturally relevant conditions, including acidic pH, low temperatures, starvation, increased ammonia concentration and exposure to nitrification inhibitors. These results highlight that biofilm lifestyle could contribute to the remarkable ecological success of AOA by explaining how they maintain activity in a broad range of environments and stress conditions.

## Results and Discussion

*Nitrososphaera viennensis* biofilms were grown on soda-lime glass microscope slides (MS) under optimal growth conditions until maximum nitrite production was achieved, as previously described^57^. After reaching peak activity, biofilms were exposed to various stress conditions, including changes in temperature, pH, and ammonia concentration, as well as the presence of biological or synthetic nitrification inhibitors, desiccation, or starvation. Concentrated planktonic controls containing an equal number of cells as biofilms were evaluated under the same conditions. All experiments other than controls were conducted in triplicates unless specified otherwise and nitrite production was used as a proxy for growth. Controls for biofilm (n=9) and concentrated planktonic cultures (n=6) performed at different times with respective experiments were combined for all plotting and analysis.

### Biofilms exhibit increased nitrification at ecologically relevant low temperatures

Planktonic cultures of *N. viennensis* (isolated from garden soil in Vienna, Austria) have been reported to grow between 28–47 °C with an optimum at 42 °C^58^, which does not reflect typical Austrian soil temperatures (∼1–19 °C annually)^59^. In this study, both biofilms and highly concentrated planktonic cultures remained active well below the previously reported temperature range (Fig. 1).

**Figure 1.**
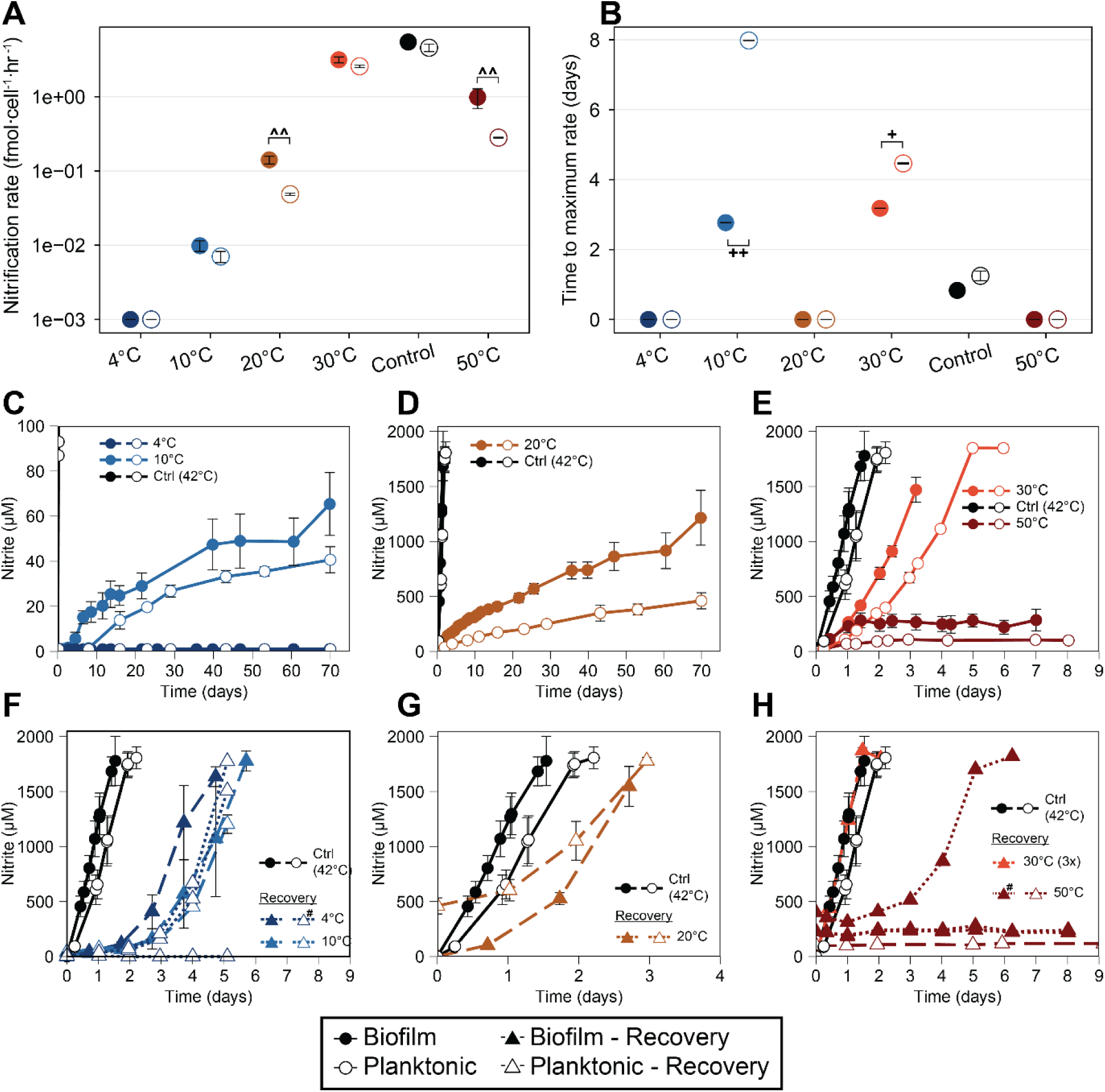
Biofilm resistance and resilience towards different temperatures. Known temperature ranges for *N. viennensis* : 28-47°C. Filled circles indicate biofilms, open circles planktonic cells. **(A)** Maximum ammonia oxidation rates (n_max_ ) per cell (fmol cell^−1^ h^−1^) plotted on a log scale at low (4°C, 10°C), medium (20°C), and high temperatures (30°C, 42°C, 50°C). Rates were calculated per replicate. Carrot indicates significant biofilm to planktonic ratios: (^ 1.5-2.5, ^^ 2.5-5.0) **(B)** Time to reach maximum rates (*t*_max_) at the same temperature ranges as in (A), calculated per replicate; shown are means ± standard deviation. Plus signs indicated significant differences of means: (+1.0-3.0, ++3.0-7.0, +++ >7) **(C-E)** Nitrite production of *N. viennensis* as biofilms or planktonic controls. Planktonic cultures were inoculated with cell numbers matching those in the biofilms. **(F-H**) Nitrite production of recoveries after one (4°C, 10°C, 20°C, 50°C) or three consecutive transfers (30°C) at the respective temperatures. Shown are means ± standard deviation of nitrite production (µM) over time, grouped by temperature range. Filled and open symbols indicate biofilms and planktonic cells respectively. Growth curves where replicates greatly diverged are plotted individually as dotted lines and marked with (#) in the legend. In all panels, if not plotted individually, data points represent averages ± standard deviation as error bars (biofilm controls, n=9; planktonic controls n=6; all others, n=3). All maximum rates and times are provided in Dataset S1.

Neither growth mode produced nitrite at 4 °C, but production occurred at 10 °C, where biofilms outperformed planktonic cultures in response time (t_max_; Fig. 1ABC). Nitrite production did not start immediately, likely due to cold shock following the abrupt temperature shift. Both growth modes recovered after returning to optimal conditions, with slightly faster recovery in biofilms (Fig. 1F). Although no nitrite production was detected in biofilms at 4 °C, these cultures recovered faster than both planktonic 4 °C cultures and all 10 °C cultures.

The largest differences between growth modes occurred at 20 °C and 50 °C. While activity was reduced at 20 °C compared to 42 °C controls (Fig. 1A), biofilms showed almost threefold higher maximum nitrification rate (n_max_) than planktonic cultures and both maintained stable activity throughout the 70-day incubation (Fig. 1D). After transfer back to 42 °C, activity resumed after a short delay and was nearly fully restored within 24 h (Fig. 1G). These findings suggest that *N. viennensis* remains active at low temperatures but that activity in low-density planktonic laboratory cultures may fall below detection limits. Given that high planktonic AOA concentrations are unlikely in soils, biofilm growth may explain nitrification attributed to AOA at environmentally relevant temperatures. Even when accounting for cell density, biofilms still outperformed planktonic cultures at 10 °C, 20 °C, and during recovery from 4 °C.

At higher temperatures relevant to soil environments (30 °C, 42 °C, and 50 °C), biofilms also showed greater activity or adaptability. In Central Europe, topsoil temperatures can reach ∼34 °C in natural soils and ∼37 °C in urban soils^60^, with extremes of 47 °C beneath pavement^60^, and are expected to increase with climate change^61–63^. At 30 °C both growth modes remained active with similar n_max_ (∼50% of optimal; Fig. 1A), but biofilms reached maximum activity faster (lower t_max_; Fig. 1B), suggesting higher adaptability under fluctuating temperatures. Repeated transfers of biofilms at 30 °C further increased growth rates (Fig. S1), and returning them to 42 °C immediately restored initial n_max_ (Fig. 1H). At 50 °C neither growth mode oxidized all supplied ammonia, but biofilms oxidized more ammonia at higher rates before growth ceased (Fig. 1E). After seven days at 50 °C, one of three biofilm replicates recovered at 42 °C, whereas none of the planktonic cultures recovered (Fig. 1H).

Overall, *N. viennensis* showed strong resilience across a wide temperature range, particularly in biofilms (Fig. 1FGH). Cultures rapidly resumed activity after prolonged periods of minimal (10 °C) or undetectable (4 °C) activity, suggesting an ability to persist through seasonal low temperatures. Biofilm growth enabled *N. viennensis* to maintain substantially higher activity at low temperatures than previously recognized, providing a plausible explanation for its success in soils well below its apparent temperature optimum.

### Biofilms confer advantage in anthropologically influenced environments

Modern agriculture depends heavily on large-scale nitrogen-based fertilizer application. The impact of fertilizer application on microbial communities has been widely studied^64,65^, but general conclusions are difficult to draw due to the high diversity and complexity of soil habitats. In the context of nitrifiers, AOB rather than AOA have been shown to be the dominant nitrifiers at high ammonium levels, even in environments dominated by AOA^31,32^. Once fertilizer is depleted and nitrogen levels have lowered, AOA would be more active^32,33^. This ecological pattern would suggest that while AOB outcompete AOA at high nitrogen levels, AOA are resilient to these large nitrogen fluxes.

The described optimal NH_4_Cl concentration for *N. viennensis* is 2.6 mM, but concentrations of up to 15 mM are tolerated^58^. However, both biofilm and planktonic cultures displayed activity at 20, 30, and 40 mM of total NH_4_ ^+^/NH_3_ (hereafter referred to as ammonium) (Fig. 2A), which is likely partially due to the increased cell density, as also seen for temperature. However, in some cases biofilms exposed to higher ammonium showed even higher nitrification compared to controls suggesting an additional advantage outside of purely cell concentration effects. At 20 mM ammonium, biofilms not only achieved higher n_max_ than planktonic cells, but also performed better than the control conditions (2mM ammonium), implying that biofilm growth offers a competitive advantage at elevated ammonium concentrations. Although the n_max_ rates of planktonic and control cultures were similar at 20 mM ammonium (Fig. 2A), planktonic cultures had to first adapt to elevated ammonium concentrations as reflected by increased t_max_ values compared to biofilms and controls respectively (Fig. 2C).

**Figure 2.**
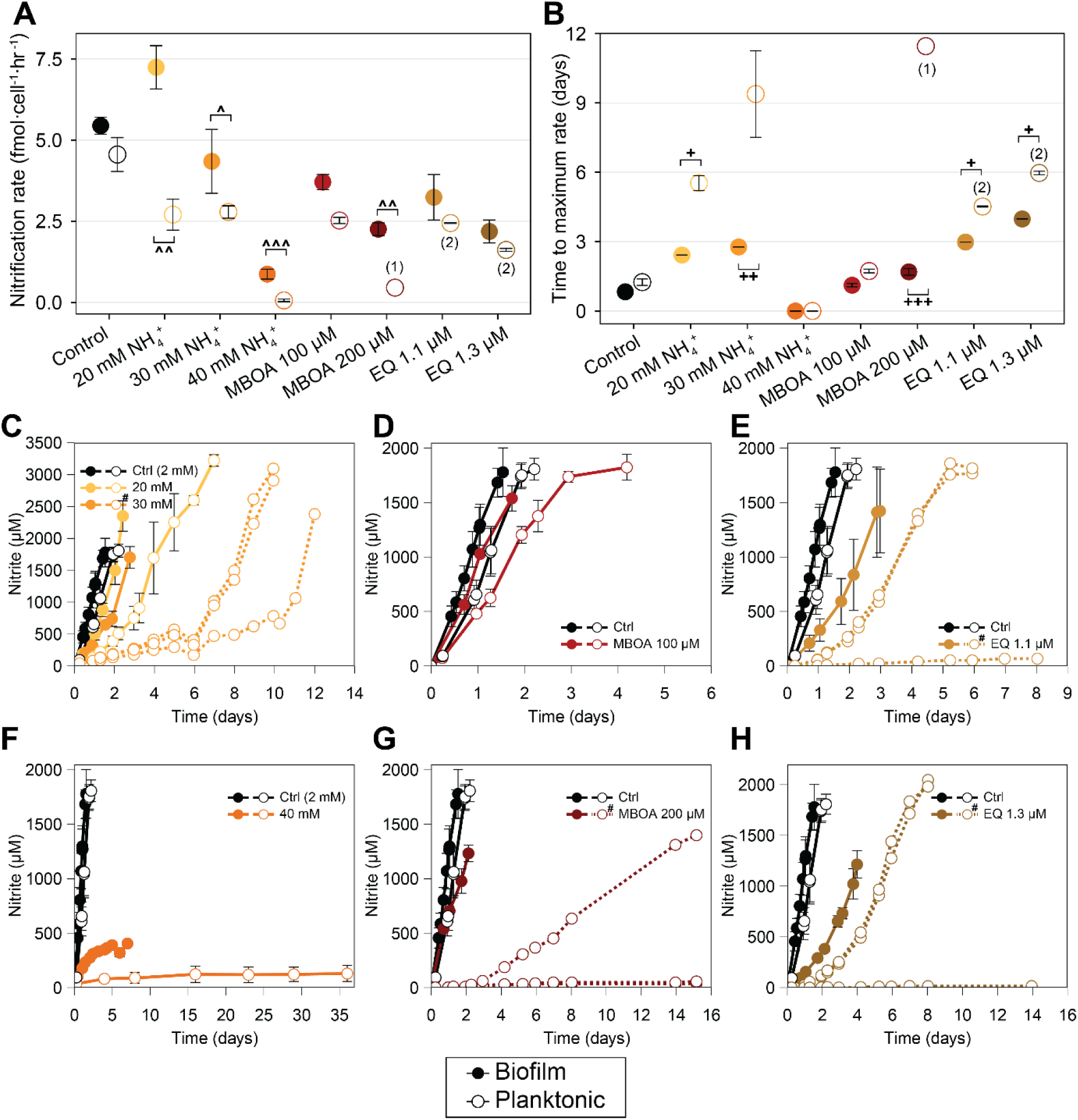
Biofilm resistance towards different ammonium concentrations and nitrification inhibitors (NIs). Known maximum ammonium ranges for *N. viennensis* : 15 mM. Filled circles indicate biofilms, open circles planktonic cells. **(A)** Maximum ammonia oxidation rates (n_max_) per cell (fmol cell^−1^ h^−1^) plotted on a linear scale at 20, 30, and 40 mM total ammonium, 100 and 200 µM MBOA, and 1.1 and 1.3 µM ethoxyquin (EQ). Rates were calculated per replicate. Parentheses represent conditions where less than three replicates were used. Carrot indicates significant biofilm to planktonic ratios: (^ 1.5-2.5, ^^ 2.5-5.0) **(B)** Time to reach maximum rates (*t*_max_) under the same conditions as in (A), calculated per replicate. Parentheses represent conditions where less than three replicates were used. Plus signs indicated significant differences of means: (+1.0-3.0, ++3.0-7.0, +++ >7). Effects of different ammonium **(C, F),** MBOA **(D, G)**, and EQ concentrations **(E,H)** on nitrite production of *N. viennensis* biofilms or planktonic controls. Planktonic cultures were inoculated with cell numbers matching those in the biofilms. Growth curves where replicates greatly diverged are plotted individually as dotted lines and marked with (#) in the legend. In all panels, if not plotted individually, data points represent averages ± standard deviation as error bars (n≥3), average ± range (n=2), or single data points (n=1) (biofilm controls, n=9; planktonic controls n=6; all others, n=3 unless otherwise indicated). All maximum rates and times are provided in Dataset S1.

Activities at 30 mM of ammonium again highlighted the faster adaptability of biofilms. Biofilms again displayed a higher n_max_, and were faster in reaching them (Fig. 2AB). At the highest concentration tested (40 mM ammonium), both growth modes showed activity before being completely inhibited. However, biofilms oxidized four times as much ammonium and reached a higher n_max_, before being inhibited (Fig. 2AF) similar to the trend observed at 50°C (Fig. 1E).

The advantage of biofilms in both n_max_ (20 mM, 30 mM, and 40 mM) and t_max_ (20 mM and 30 mM) suggests that biofilms may exist in a physiological state that allows the cells to respond more readily to sudden changes in ammonium/ammonia availability, whereas planktonic cells would require time to adapt. One possible explanation for this difference is the higher physiological and transcriptional heterogeneity observed in biofilms^66^, exemplified by the upregulation of genes for urea utilization, even in its absence^57^. Another likely explanation is that biofilms provide a protective barrier alleviating the effects of excessive ammonium concentrations.

The continuous transfer of biofilm cultures at 20mM and 30 mM of ammonium showed either no impairment or a steady decline of nitrification rates respectively (Fig. S2). Likewise, the recovery of biofilms to optimal conditions after repeated transfers was immediate at 20 mM, and slower at 30 mM (Fig. S2). Biofilms recovered from a seven day long exposure to 40 mM with an extended lag phase (4 days) before exhibiting a quick recovery. This suggests that biofilms remain active up to 20 mM of ammonium and are resilient to prolonged exposure of ammonium concentrations of up to at least 40 mM.

Growth of AOA as biofilms therefore offers a plausible explanation for the observed ecological patterns of nitrifiers in fertilized soils: although activity is reduced, they are resilient to transient spikes in ammonium and resume normal activity once the ammonium stress has been alleviated by AOB and plant activities.

Another common application to agricultural fields along with ammonium are nitrification inhibitors (NIs) with the intended purpose of reducing the activity of microbial nitrifiers. However, the efficacy of NIs in laboratory cultures and field trials remains inconsistent^45–48^, indicating the role of yet unknown factors that affect it. Therefore, the impacts of biofilm growth on the sensitivity to the biologic nitrification inhibitor 6-methoxy-2(*3H*)-benzoxalone (MBOA)^67^ and the synthetic nitrification inhibitor ethoxyquin (EQ)^68^ was tested using *N. viennensis*. Biofilms retained activity in the presence of both NIs, and recovered rapidly after exposure, while planktonic cultures were more susceptible at every concentration tested (Fig. 2DEGH). While biofilms were not significantly different in n_max_ and t_max_ than planktonic cells at 100 µM MBOA, they conferred a significant advantage at 200 µM of MBOA, with all biofilms continuously growing. Conversely, only one planktonic triplicate was able to maintain growth (Fig. 2G). Biofilms exhibited similar n_max_ for both tested concentrations of EQ of 1.1 and 1.3 µM (Fig. 2EH). However, only two planktonic cultures of a triplicate measurement of both concentrations were consistently active, giving biofilms an advantage in growth. Additionally, the benefit of biofilm growth when exposed to ethoxyquin was observed in t_max_ as biofilms reached a maximum rate faster in both conditions tested. Overall, biofilms were resilient to all tested NI concentrations, displaying none or minimal lag phases upon recovery to optimal medium (Fig. S3).

Biofilms are well known to enable resistance to antimicrobial compounds^51^ (i.e., antibiotics^69^) presumably through a variety of mechanisms^70^, which could similarly happen with nitrification inhibitors. While lab grown planktonic cultures are often strongly inhibited by NIs, equivalent concentrations frequently fail to suppress nitrification in soil studies^45–48^. NI efficacy is dependent on several factors, including physicochemical properties of soils like soil pH^71,72^, but also community composition^72^. The results shown here suggest that growth of ammonium oxidizers as biofilms might additionally account for the reduced efficacy of NIs *in situ*.

### Biofilms confer a growth advantage at low pH

A natural consequence of nitrification in soils and other environments is a decrease in pH. *N. viennensis* is described to grow between pH 6.0 - 8.5 with an optimum at a neutral pH of 7.5 when grown planktonically^27^. However, *N*. *viennensis* is detected in soils of a much wider pH range (Fig. 3A) while maintaining comparable abundances to neutrophilic conditions (Fig. S4). Therefore, the possibility of biofilms conferring resistance to acidic pH, as proposed previously for AOB^73,74^, was tested. Both biofilms and planktonic cultures actively oxidized ammonia in all tested acidic pH conditions (5.0, 5.5, 5.7, 6.0), with biofilms reaching higher n_max_ rates for all conditions except pH 5.0 at which ammonia oxidation of both growth conditions was slowed the most, only producing ∼65 µM NO_2_ ^−^ before inhibition (Fig. 3F). The n_max_ was consistently reached during the initial phase of nitrite production for both biofilm and planktonic cultures leading to a negligible t_max_ in all cases (Fig. S5).

**Figure 3.**
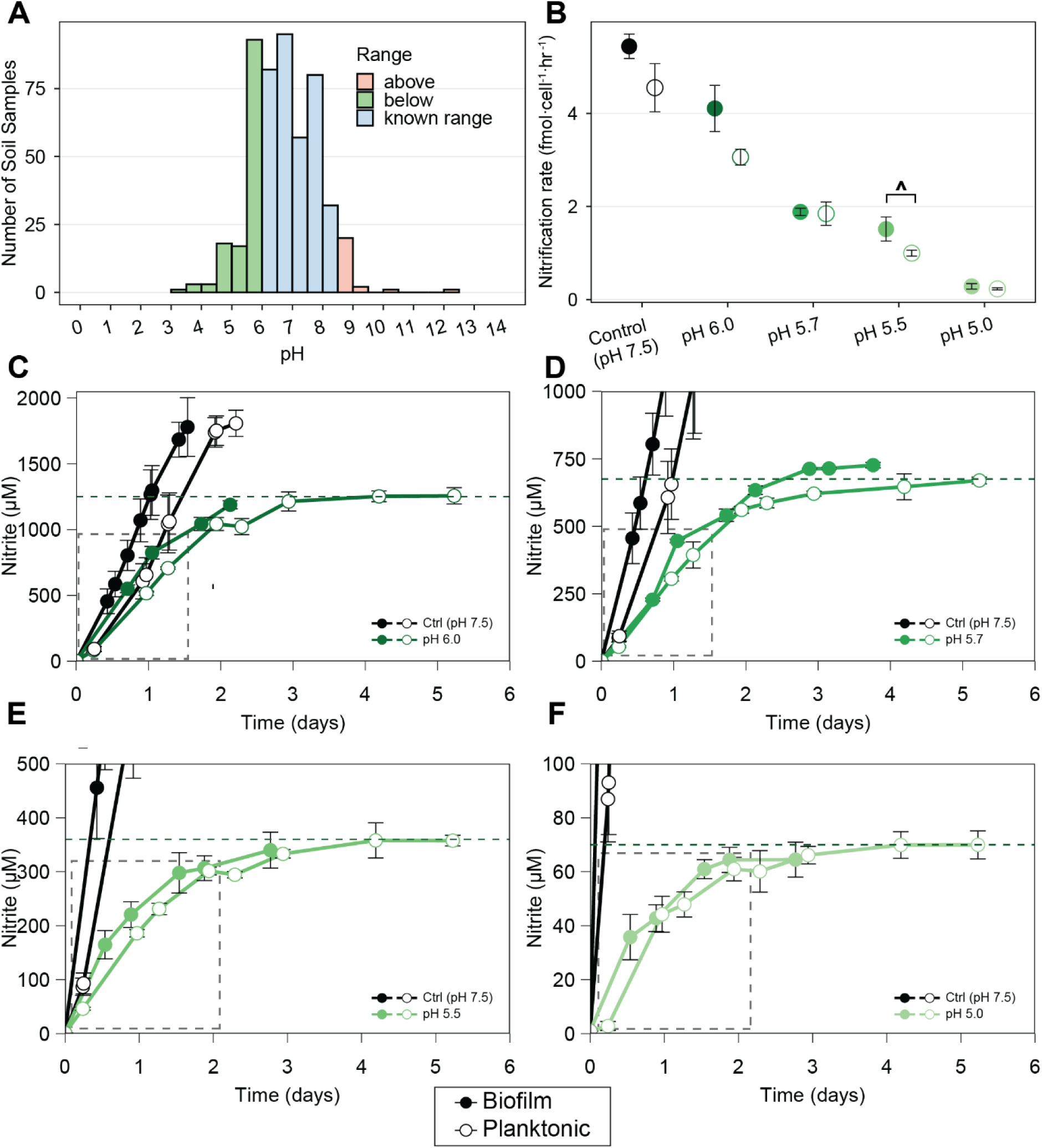
Biofilm resistance to pH. Known pH range for *N. viennensis* : 6.0-8.5. Filled circles indicate biofilms, open circles planktonic cells. **(A)** Detection of *N. viennensis* EN76 in soil samples with varying pH values. **(B)** Maximum ammonia oxidation rates (*n*_max_ ) per cell (fmol cell^−1^ h^−1^) plotted on a linear scale at pH 6.0, 5.7, 5.5 and 5.0. Rates were calculated per replicate. Filled circles indicate biofilms, open circles planktonic cells. Carrot indicates significant biofilm to planktonic ratios: (^ 1.5-2.5, ^^ 2.5-5.0) **(C-F).** Growth of *N. viennensis* as biofilms or planktonic controls. Planktonic cultures were inoculated with cell numbers matching those in the biofilms. Averages of nitrite production (µM) over time (days), at pH 6.0 B **(C)**, 5.7 **(D)**, 5.5 **(E)**, and 5.0 **(F)** are shown. In all panels, data points represent averages ± standard deviation as error bars (biofilm controls, n=9; planktonic controls n=6; all others, n=3). Colored dotted lines mark the pH-dependent maximum tolerable nitrite concentration. Grey dashed line boxes indicate the faster response of biofilms to acidic pH based on nitrite production. All maximum rates and times are provided in Dataset S1.

Ammonia oxidation was inhibited at different maximal nitrite concentrations in a pH dependent manner - the lower the pH, the lower the observed maximal nitrite concentration. This is in line with a previous study showing that the continuous removal of nitrite by coculturing *Nitrosotalea* spp. with the NOB *Nitrobacter laanbroekii* NHB1 lowered the pH ranges at which *Nitrosotalea* spp. grew by ∼0.5 pH units^25^. The inhibition of AOA by accumulating nitrite concentrations at acidic pH is hypothesized to be due to formation of nitrite derived nitrous acid (HNO_2_, pKa = ∼3.3). This would explain both inhibitory nitrite concentrations directly correlated with decreasing pH values and biofilms reaching maximal nitrite concentrations faster than planktonic cells, as the biofilm and its matrix could act as protective barrier against slowly rising concentrations of HNO_2_. At the higher pHs tested here, it is also possible that the alternative by-product peroxynitrite (pKa = ∼6.8)^75^, which is produced by the interaction of nitric oxide, a metabolite implicated in AOA metabolism^76^, and reactive oxygen species, is responsible for this inhibition.

Biofilm cultures initially exposed to low pH were continually transferred to determine the effect of longer exposure times to low pH. In all cases, n_max_ rates steadily decreased during successive transfers in acidic pH (Fig. 4). After the first two transfers, biofilms were challenged by being exposed to an approximate of the previously observed maximal nitrite concentration respective for each condition from time point zero (third transfer-challenge). As expected, little to no additional nitrite was produced during this transfer (Fig.4, Fig. S6). Upon the fourth transfer (a return to low pH without nitrite) rates either remained minimal (pH 5.0), decreased further (pH 5.5 and 5.7), or exhibited a small amount of increase (pH 6.0). After this prolonged exposure to acidic pH (4 transfers, ∼10-18 days), all biofilms were able to fully oxidize supplied ammonia upon a return to optimal growth medium (recovery), albeit at impaired n_max_ (Fig. 4). The lower the pH, the longer it took all replicates of respective pHs to fully oxidize ammonia, with the standard deviation increasing considerably with decreasing pH.

**Figure 4.**
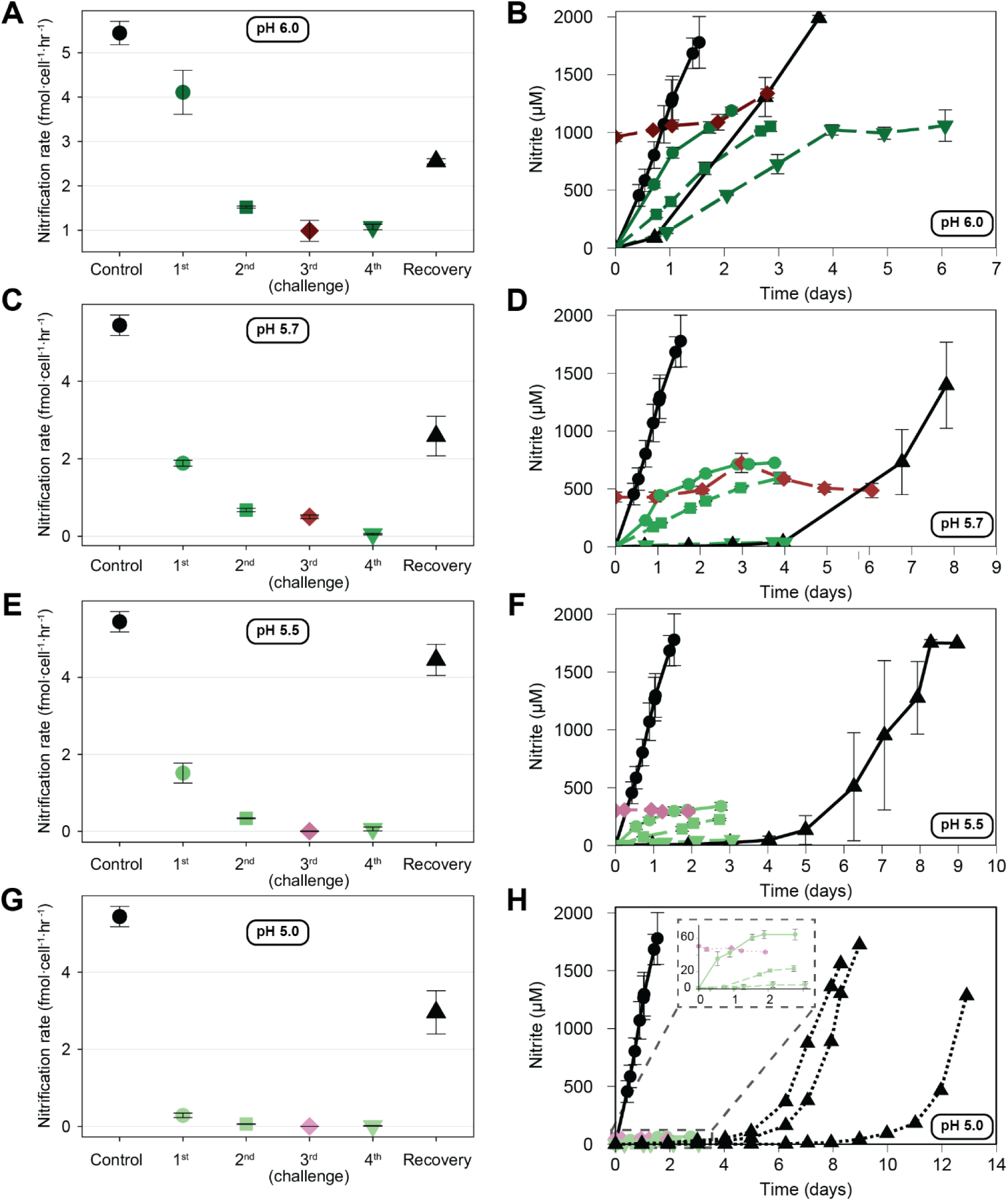
Biofilm resilience to pH. **(A,C,E,G)** Maximum ammonia oxidation rates (*n*_max_ ) of biofilms per cell (fmol cell^−1^ h^−1^) plotted on a linear scale at pH 6.0, 5.7, 5.5 and 5.0 including all consecutive transfers at those pH and recovery at 7.5. Biofilms were transferred twice at the respective pH (circle, rectangle), followed by a third transfer into medium containing the maximum nitrite concentration tolerable at that pH (diamond, challenge), and a subsequent transfer without nitrite (inverted triangle). Challenges are indicated in shades of red. Biofilm recovery by being transferred to medium with pH 7.5 (triangle). Rates were calculated for individual replicates. **(B,D,F,H)** Growth of *N. viennensis* biofilms at pH 6.0, 5.7, 5.5, and 5.0 following the same transfer sequence, and recovery at pH 7.5 as described in (A,C,E,G). Challenges are indicated in shades of red. Grey dashed box in **(H)** indicates a zoomed in portion of growth curves. Enlarged figures for resistance at pH 5.5 and 5.0 can be found in Figure S6. Growth curves where replicates greatly diverged are plotted individually as dotted lines. In all panels, if not plotted individually, data points represent averages ± standard deviation as error bars (biofilm controls, n=9; all others, n=3). All maximum rates and times are provided in Dataset S1.

Biofilm-mediated resistance to acidic pH, including tolerance to prolonged exposure, provides a potential mechanism for the survival and prevalence (Fig. 3A) of neutrophilic AOA in low pH soils. The clear dependence on nitrite and pH levels shown here for a neutrophilic strain, and previously observed for an acidic AOA strain^25^, is also a critical factor to consider as most soils contain nitrite-oxidizing bacteria (NOB) that readily convert nitrite to nitrate^77^. Therefore, it is highly probable that a biofilm growth mode of AOA, along with the activity of naturally present NOB, significantly expands the ecophysiological niches in which non-acidophilic AOA can survive or thrive.

### Biofilms are resilient to starvation or desiccation phases

Other likely environmental stressors for soil organisms include starvation and desiccation. AOA enrichment cultures can recover after starvation for up to 50 days^78^, but their tolerance to desiccation remains poorly understood. After starvation for 7 or 30 days both biofilms and planktonic cells recovered similarly, though with extended lag phases of 2 and 7 days, respectively, as indicated by elevated t_max_ values (Fig. S7). Tolerance to starvation might offer a competitive advantage in natural environments where nutrient availability is highly variable. Desiccation for one or three days similarly allowed complete recovery of both biofilms and planktonic cells, albeit with prolonged lag phases. After 1 day of desiccation, planktonic cells recovered more rapidly than biofilms, whereas recovery was comparable after 3 days, although with high variability between replicates (Fig. S7). These differences may be due to methodological limitations in achieving uniform desiccation across all cultures. Given that increased resistance to desiccation is considered a hallmark feature of biofilms^50^, the resistance of planktonic cells might point towards a more general resistance of AOA which warrants further research.

### Biofilms display similar activity with less nitrous oxide emissions than planktonic cultures

In earlier studies the affinity of different AOA for ammonia has been determined with concentrated planktonic cells ^34,35,37^. To better understand affinities relevant in natural environments, biofilms were grown directly in microrespiratory glass chambers later used for rate measurements and compared to planktonic controls containing the same amount of cells. When comparing the Michaelis-Menten kinetics between biofilms and planktonic cells, remarkably similar values were observed (Fig. 5A) with K_m(app)_ of 27.60 µM NH_4_ ^+^/NH_3_ and 34.11 µM NH_4_ ^+^/NH_3_ as well as V_max_ of 203.70 µM NH_3_/hr and 200.46 µM NH_3_/hr for planktonic and biofilm cells respectively. It remains unclear whether this similarity reflects a uniform activity level across all cells in the biofilm or results from a heterogeneous structure in which the outer layers harbor highly active cells, while deeper layers remain metabolically less active. Nevertheless, this study confirms that the measured affinities of *N. viennensis*, and perhaps all AOA, remain valid when biofilm-associated growth is taken into account.

**Figure 5.**
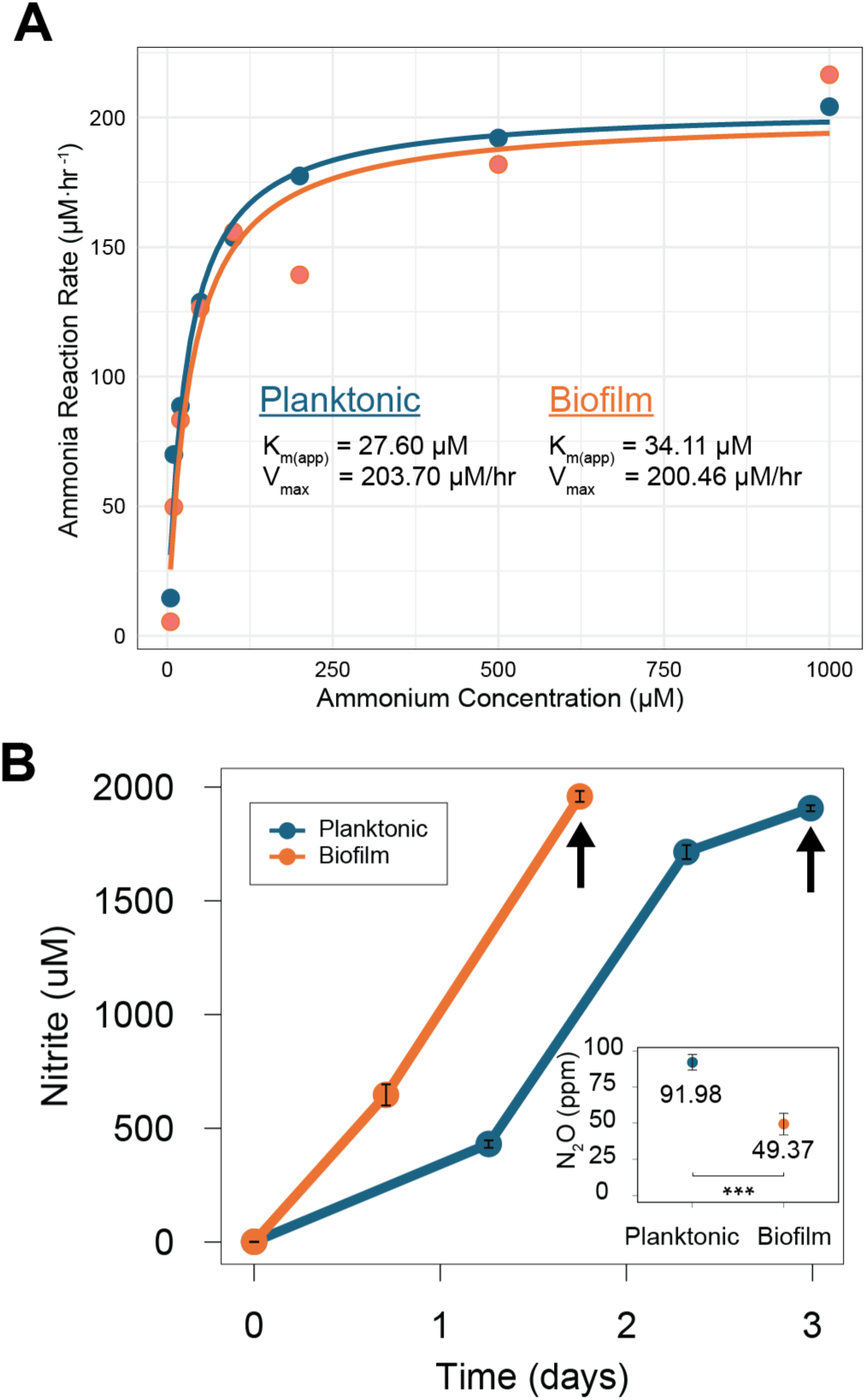
Michaelis-Menten kinetics and N_2_O measurements: **(A)** Ammonia oxidation rate (µM NH_3_/hr) as a function of substrate (NH_4_^+^/NH_3_) concentration. Data points represent individual measurements; the fitted Michaelis–Menten curve is shown with the apparent Michaelis constant (Km_(app)_) and maximum rate (V_max_) indicated. Experiments were done with biofilms and an equal number of concentrated planktonic cells. Microrespiratory graphs and data can be found in Figure S# and Dataste S1 respectively. **(B)** Growth curves for planktonic and biofilm cultures in sealed culture vessels shown as average ± standard deviation (n=5). Inset: N_2_O measured for biofilms and the same amount of planktonic cells in 5 replicates, displayed are produced ppm N_2_O measured in the gas phase of cultivation vessels. Asterisks indicated level of significance: (*** p-value ≤ 0.001).

While activity remains unchanged, the same cannot be said for the metabolic by-product nitrous oxide (N_2_O). Nitrous oxide is a potent greenhouse gas that is known to be produced by the activity of ammonia-oxidizing microorganisms^5–9^. The production of N_2_O was assessed by measuring the headspace in Schott bottles of quintuplicate cultures containing either biofilm or planktonic cultures containing the same number of cells.

Surprisingly, planktonic cells emitted approximately double the amount of N_2_O as *N. viennensis* biofilms (Fig. 5B). Production of N_2_O in pure cultures of AOA has been previously attributed to the abiotic reaction of reactive nitrogen species^76^, but recent evidence also indicates a biological route^9^. Additionally, a biochemical basis for the production of N_2_O in AOA has been revealed by the recent characterization of a multicopper oxidase enzyme^79^. This enzyme oxidized hydroxylamine during nitrification and generated nitroxyl (HNO) as an intermediate that can react with itself to form nitrous oxide. This enzyme was characterized from a marine AOA, but is particularly widespread in terrestrial strains^57^ and could serve as an electron sink when an excess of hydroxylamine is produced. However, this gene was shown to be up-regulated in biofilms of *N. viennensis*^57^, which would not intuitively match a decrease in the N_2_O production shown here. Alternatively, N_2_O could be produced by another mechanism via dark oxygen production^80,81^, which has also recently been proposed as an electron-overflow pathway in AOA, active under oxic growth conditions ^82^. Both scenarios frame the production of N_2_O as a by-product of balancing electrons in the ammonia oxidation process. Therefore, biofilms might exhibit enhanced capacity for managing electron flow and maintaining redox homeostasis, potentially reducing the N_2_O produced via these pathways. Considering that biofilms are known to sequester CO_2_ and different nitrogen species^83^, another possibility is that N_2_O could be trapped in the biofilm’s matrix, reducing the measurable amount of N_2_O compared to planktonic cells.

### Biofilms and aggregates of ammonia-oxidizing archaea

Many of the advantages conferred to cells by growing as biofilms, including those described here, are attributed to the matrix of EPS in which they are embedded. This matrix acts as a protective coating by physically separating cells from the immediate environment. Although the exact three-dimensional structure and chemical composition of AOA biofilms and their EPS remain to be elucidated, the autofluorescence of AOA based on the expression of cofactor F_420_ enabled us to perform live imaging of cells relative to their position in the biofilm matrix without using destructive staining techniques (Fig. 6). While it was apparent from phase-contrast light microscopy that biofilms contain a massive number of single cells in a three-dimensional structure (Fig. 6ADG), fluorescence microscopy confirmed the distribution of single cells and clusters thereof encased within the EPS structure of the biofilm.

**Figure 6.**
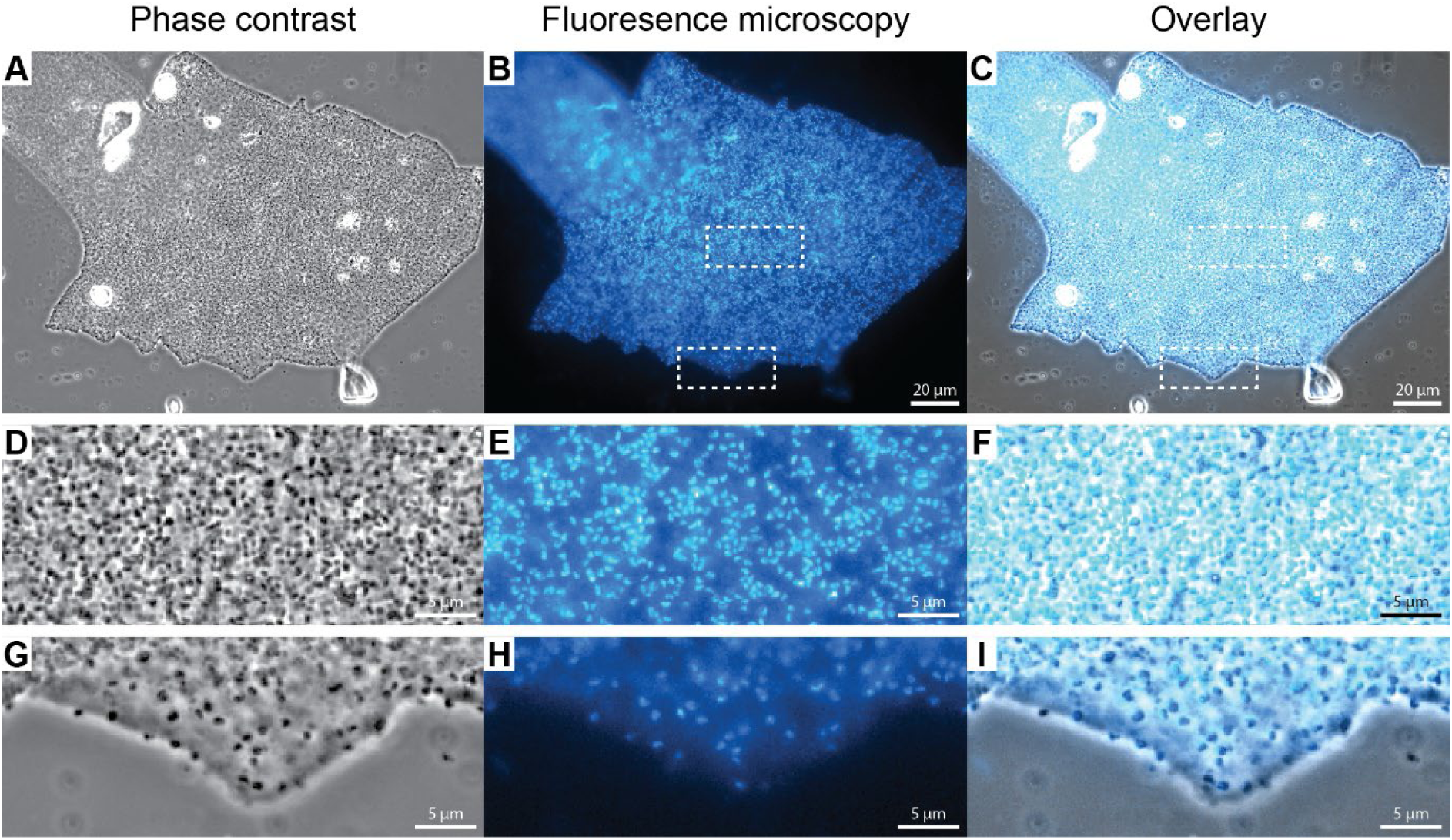
Cofactor F_420_ autofluorescence of an *N. viennensis* biofilm. **(A, D, G)** Phase contrast light microscopy **(B, E, H)** Fluorescence microscopy images based on autofluorescence due to cofactor F_420_. Images were taken with a standard Nikon fluorescent filter cuber for DAPI and are false colored in Cyan. **(C, F, I)** Overlays of corresponding phase contrast and fluorescent images. **(D-F)** and **(G-I)** are magnifications of the central and peripheral areas marked in **(A-C)** respectively. Images are from a biofilm of *N. viennensis* grown on microscopy slides.

The encasement of AOA cells within EPS matrices is not restricted to *N. viennensis* and was also observed using F_420_ autofluorescence in multiple other species (Fig. S8), i.e. in biofilms of *Nitrosocosmicus franklandianus* (Fig. S8 A-C), aggregates of *Nitrosocosmicus arcticus* (Fig. S8 D-F), and in biofilms of a recently isolated strain from human skin, ‘*Candidatus* Nitrosocosmicus epidermidis’ (Fig. S8 G-I). Equally, biofilms of the marine strains *Nitrosopumilus maritimus* (Fig. S8 J-L) and *Nitrosopumilus adriaticus* (Fig. S8 M-O) could be visualized. The presence of multiple AOA species embedded within a protective coat of EPS during biofilm growth is suggestive that many of their prior established ecophysiological ranges can also be expanded and the benefits of biofilm growth shown here are not exclusive to *N. viennensis*.

## Conclusion

Ammonia oxidizing microorganisms are ideally suited to measure cellular activity in biofilms because their metabolic product (nitrite) can be easily monitored in the supernatant. This gave us the unique opportunity to directly compare activities in biofilm versus planktonic cells.

Biofilms of *N. viennensis* were more resistant to temperature, acidic pH, elevated ammonium concentrations, and application of both biological and synthetic nitrification inhibitors compared to planktonic controls. Moreover, *N. viennensis* biofilms displayed a high universal resilience to stress conditions with the exception of a high temperature extreme of 50°C. Although the activity of biofilms and concentrated planktonic cultures of *N. viennensis* were virtually identical under optimal conditions, biofilms released almost half the amount of the greenhouse gas N_2_O.

The underlying mechanisms of how biofilms confer increased resistance and resilience remain to be elucidated, but the presence of EPS remains an extremely likely candidate. The protective effects of biofilms on small molecules like elevated ammonia concentrations, addition of NI compounds, or the accumulation of reactive nitrite-based compounds at acidic pH could be similar to the proposed mechanisms for antibiotic resistance in bacterial biofilms. While the results presented here indicate that reduced growth rate is not a factor for *N. viennensis,* the physical separation of molecules from cells by the EPS matrix, an increase in adaptive responses, or the presence of microenvironments within the biofilm could all play a role^84^. While biofilms are often described as reservoirs of microbial cells that enable survival, seeding, or nutrient capture under reduced per cell activity^85,86^, the findings here suggest that AOA biofilms can maintain high cell specific activities comparable to planktonic cultures. This active biofilm lifestyle may contribute, together with the observed protective benefits and resilience of biofilms, to the ecological success of AOA.

Based on this and our previous work it is highly likely that many of the observed benefits conferred by biofilm formation are conserved in other AOA species, both terrestrial and marine strains.

We anticipate that our findings are likely conservative, as they are based on pure cultures grown on inert glass surfaces. In natural soils, microorganisms are embedded in complex multispecies biofilms surrounded by a structured matrix, where enhanced EPS production and community interactions are expected to further amplify the protective advantages of the biofilm lifestyle^87^. Regardless, this study represents first insights into the ecophysiological relevance of biofilm formation by AOA. The increased fitness exhibited by biofilms for all tested conditions helps to resolve many discrepancies between lab observations and the natural environment and thus supports the hypothesis that biofilm growth is the preferred *in situ* phenotype of AOA in soil environments.

## Materials and Methods

### Cultivation

*N. viennensis* was grown in freshwater medium (FWM) supplemented with 2 mM NH_4C_l, 2 mM NaHCO_3,_ 1 mM pyruvate, 50 µg/ml kanamycin, 7.5 µM FeNaEDTA, non-chelated trace elements (0.1% volume/volume (v/v)), and buffered with 10 mM HEPES to pH 7.5. A detailed description of the growth medium has recently been published (Dreer et al., 2025 Table S1^57^). Cultures were grown at 42°C, 80 rpm shaking in the dark. All solutions were sterilized by filtration (0.2 µm mixed cellulose ester (MCE) membrane filters), except FWM (autoclaved). Nitrite production was followed by colorimetrically measuring nitrite concentrations via the Griess reaction as described previously^88^.

#### Biofilm growth on microscopy slides

Biofilms were grown as described recently^57^. Briefly, silanized soda-lime glass microscopy slides (MS) (Carl Roth, Histobond, CEX0.1, 76×26×1 mm) were individually added to 250 ml schott bottles containing 125 ml growth medium, using polymethylpentene tweezers (Vitlab, 145 mm). MS were transferred five times between 1.3 - 1.8 mM nitrite, carefully dipping MS three times in prewarmed growth medium to remove all remaining liquid from the previous transfer using tweezers. The sixth transfer of MS was used for experiments based on reaching previously described maximal nitrite production^57^. MS and tweezers were sterilized by autoclaving.

#### Determination of AOA abundances by fluorescence-activated cell sorting (FACS)

Cell abundances for planktonic AOA and AOA growing in biofilms were quantified by fluorescence-activated cell sorting (FACS) as in Malits et al., 2023^89^. Briefly, cells were fixed in 0.2 µm-filtered glutaraldehyde (0.5% final concentration) for 10-20 min at 4°C and deep-frozen. After thawing, samples were diluted in 0.2 µm-filtered TE buffer (pH 8, 10 mM TRIS-1 mM EDTA) and stained with SYBR Green I (1:20000, diluted in ultrapure water) in the dark for 10 min. Samples were analyzed using a FACS Aria II (BD Biosciences) flow cytometer. The sample flow rate was accurately calibrated daily and used to calculate the abundances of the AOA population either derived from biofilms or from the planktonic phase.

#### Planktonic culture controls

To determine appropriate cell numbers for planktonic controls, three replicate biofilm cultures were grown as described above. After 5 complete transfers (the time to reach maximum nitrite production), biofilm triplicates were washed and resuspended thoroughly with full medium using a syringe and needle to dislocate biofilm cells from microscopy slides and break up aggregates. Microscopy slides were checked via phase contrast light microscope to ensure the complete removal of cells. Resuspended cells were counted using FACS (see above). The cell numbers obtained here were used as the benchmark for all performed planktonic controls.

To reliably determine exact cell numbers of subsequently grown planktonic cultures, fluorescence-activated cell sorting (FACS) was performed over a complete growth curve of *N. viennensis*, linking nitrite values with cell numbers. From this data, a script based on linear interpolation to calculate cell numbers between 1-1809 µM of nitrite was created and is available on github (https://github.com/pribasnig/N_viennensis_NO2_to_cellnumber_tool). The script was validated using an independent FACS cell counting experiment.

Planktonic controls were then grown separately from biofilm cultures. For all planktonic culture controls, *N. viennensis* was grown in 500mL of medium in 1 L culture vessels and cell numbers were determined by measuring nitrite and calculating the corresponding cells/ml with the aforementioned script. Cells were centrifuged at 21,000xg for 30min at 40°C ( using a Thermo Scientific Fiberlite F12-6×500 LEX rotor and 400 ml centrifugation vessels), resuspended in a known volume of fresh growth medium, and appropriate cell numbers were aliquoted into the respective experimental conditions of 250 mL of medium in 500 mL culture bottles, identical to the medium and culture vessels used for biofilm growth.

#### Biofilm growth in microrespiratory chambers

A microrespiration chamber (Unisense, 2 ml) made of glass was added to a 30 ml polystyrene container (Greiner) containing 20 ml growth medium. Using tweezers, the chamber was transferred to fresh growth medium between 1.3 mM - 1.8 mM nitrite, carefully submerging the chamber in prewarmed growth medium three times, slowly pouring the medium out of the chamber after each submersion to remove all remaining liquid from the previous transfer. As described previously for biofilms on glass slides^57^, nitrite production accelerated with each subsequent transfer until reaching a maximum (Fig. S9). The resulting biofilm forming directly on the chamber was used to measure oxygen consumption after 30 transfers as described below. The chamber and tweezers were sterilized by autoclaving. After microrespiratory measurements, biofilms were resuspended thoroughly with full medium using a syringe and needle, as described for microscopy slides, and cell abundances determined via FACS. The acquired cell number (2.91×10^8^ cells) was used to set up planktonic cultures for comparison.

### In situ *pH Range of* N. viennensis

Metagenomic abundance profiles for *Nitrososphaera viennensis* were retrieved from the MicrobeAtlas repository (https://microbeatlas.org/)^90^. Specifically, to ensure taxonomic accuracy, the “representative” dataset of *Nitrososphaera viennensis* was used (downloaded at the end of 2025). Abundance was quantified in parts per million (ppm), a normalized metric representing the number of reads mapped to the target genome per one million total sequenced reads. To correlate abundance with environmental conditions, sample metadata was parsed from the NCBI Sequence Read Archive (SRA) using unique Run IDs to extract pH values. We visualized the distribution of *N. viennensis* across the pH gradient by plotting using the ‘ggplot2’ package^91^ in R version 4.5.3^92^. To address the wide dynamic range of the abundance data, we applied a log_10 t_ransformation to the y-axis, while the x-axis was constrained to a biological pH range of 2 to 14. The number of soil samples where *N. viennensis* was detected was then binned and counted into pH ranges in increments of 0.5.

### Resistance and resilience tests

Due to the necessity to develop biofilm samples over multiple transfers and the need to concentrate actively growing cells for planktonic controls, biofilm and planktonic cultures could not be performed simultaneously in an accurate manner. However, biofilm controls and planktonic controls (42°C, 2 mM NH_4C_l, pH 7.5) were always performed with their respective stressed counterparts. Biofilm experiments were performed in three different groups (Group 1: controls, 30°C, 50°C, 20mM NH_4_Cl, 40mM NH_4_Cl, 1 week starvation 4 weeks starvation; Group 2: controls, pH 6.0, pH 5.7, EQ 1.1 µM, EQ 1.3 µM, MBOA 100 µM, MBOA 200 µM, 1 day desiccation, 3 days desiccation; Group 3: controls, 30mM NH_4_Cl, pH 5.0, pH 5.5, 4°C, 10°C, 20°C). Planktonic cultures were performed in two different groups (Group 1: controls, 20 mM NH_4_Cl, 30 mM NH_4_Cl, 40 mM NH_4_Cl, 4°C, 10°C, 20°C, 30°C, 1 day desiccation , 3 days desiccation, 1 week starvation, 4 weeks starvation; Group 2: controls, pH 6.0, pH 5.7, pH 5.5, pH 5.0, EQ 1.1 µM, EQ 1.3 µM, MBOA 100 µM, MBOA 200 µM, 50°C). For plotting and statistical analysis, all performed controls were averaged together (biofilm n=9, planktonic n=6) (Fig. S10). Points for control groups were only used if data could be averaged from multiple groups (i.e., final points representing only one group were removed).

#### Temperature Experiments

For temperature treatments, optimal media was preincubated at the respective temperature for at least 24 hours before the start of the experiment. To assess the effect of temperature on the activity of biofilms, biofilms grown under optimal conditions at 42°C and then transferred to 4°C, 10°C, 20°C, 30°C, 42°C (control), or 50°C. Subsequently, biofilm and planktonic cultures were transferred back to 42°C to test their resilience. When all ammonia was oxidized within a week (30°C), biofilms were continuously transferred three times at this temperature before being returned to 42°C to assess their resilience. At lower temperatures (10°C, 20°C), where ammonia oxidation was slower, or at temperatures (4°C, 50°C), where activity was minimal or absent (4°C, 50°C), biofilms were transferred only once — either after one week (50°C) or after 70 days of low (10°C, 20°C) or no activity (4°C). Recovery of planktonic cells at 30°C was not performed as the test condition consumed all ammonium before it could be transferred to 42°C and would have necessitated additional centrifugation.

#### Ammonia and Nitrification Inhibitors

The effects of ammonia toxicity and nitrification inhibitors were tested on biofilm and concentrated planktonic cultures. Biofilms were transferred from optimal conditions (2 mM ammonium) to either 20 mM, 30 mM, or 40 mM of ammonium. If all ammonia was oxidized within a week (20 mM and 30 mM), biofilms were continuously transferred three times at the respective concentrations before being returned to 2 mM to assess their resilience. When activity ceased (40mM), biofilms were returned to optimal conditions after 7 days. The biological nitrification inhibitor 6-methoxy-2(*3H*)-benzoxolone (MBOA)^67^ and the synthetic nitrification inhibitor ethoxyquin (EQ)^68^ were tested at two concentrations. MBOA and ethoxyquin stocks were prepared freshly in dimethyl sulfoxide (DMSO), sterilized with 0.22 µM polytetrafluoroethylene (PTFE) filters and added to the medium shortly before the start of the experiment. No significant pH changes were observed. MBOA was tested at 100 µM and 200 µM while EQ was tested at 1.1 µM and 1.3 µM. After oxidizing all ammonia, or when ceasing activity after 7 days, biofilms were returned to optimal medium to test resilience.

#### pH Experiments

To assess resistance to low pH, biofilms were transferred from optimal conditions (pH 7.5) to medium with a lower pH once maximum nitrification rates were reached (see above). The acidic pHs of 6.0, 5.7, 5.5, and 5.0 were each tested individually. The pH of media was adjusted by addition of either HEPES (7.5) or MES (5.0, 5.5, 5.7, 6.0) and stable throughout growth. For acidic pH experiments 30 mM of 2-(N-morpholino) ethanesulfonic acid (MES) buffer titrated to pHs 6.0, 5.7, 5.5, and 5.0 were added, respectively. Pre-tests confirmed that 30 mM MES was sufficient to maintain stable pH throughout the incubation.

After the initial growth curve (first transfer), biofilms were transferred again to the same pH to determine the maximum tolerable nitrite concentration at each pH (second transfer). A third transfer was conducted into medium already containing this maximum nitrite concentration to challenge the growth with nitrite at a low pH (third transfer/challenge), followed by a fourth transfer into respective pH fresh medium without added nitrite. Following the fourth transfer, biofilms were returned to medium at a neutral pH of 7.5 to assess recovery and resilience.

#### Starvation and Desiccation

For starvation, NH_4_Cl was omitted from growth medium and cultures incubated at 42°C for 1 or 4 weeks, after which 2 mM NH_4_CL were added. For desiccation, microscopy slides were air dried sterile in a laminar flow hood and incubated in empty 250mL Schott bottles for 1 or 3 days. Planktonic control cultures were concentrated to a volume of approximately 500 µL and then transferred to empty Schott bottles, air-dried, and incubated under the same conditions as the biofilms. After all conditions, biofilms were incubated at optimal conditions again to test their resilience.

#### Cell specific nitrification rates and time to maximum rate

As the various stressors initiated a variety of growth responses that were not sigmoidal, each growth type (planktonic or biofilm) and each condition were modeled using the linear regression function (lm) in the R statistical package. One of three types of models was chosen: linear (represented by y=mx+b; lm(NO_2_ ^−^ ∼ time, data = data)), quadratic (represented by y=ax^2^+bx+c; lm(NO_2_ ^−^ ∼ poly(time, 2, raw = TRUE), data = data)), or cubic (represented by y=ax^3^+bx^2^+cx+d; lm(NO_2_ ^−^ ∼ poly(time, 3, raw = TRUE), data = df)). Points in each condition that represented lag or stationary extensions were removed before model fitting. Maximum nitrification rates (n_max_) and time to reach the maximum nitrification rate (t_max_) were determined by calculating the maximum of the first derivative (i.e. maximum slope) with respect to the chosen model. In the case of linear models where the slope is constant across all time points, t_max_ was set as the time when the nitrite level reached 1000 µM. In the case of replicate 3 of planktonic growth following 4 weeks of starvation (linear fit), the t_max_ for reaching 1000 µM nitrite was extrapolated based on the model parameters. A full list of all growth types and conditions, along with the selected model and the point of the maximum rates can be found in Supplementary Material S2-Models.

The values for n_max_ and t_max_ were used for the comparison of rates between biofilm and planktonic conditions. In cases where the amount of oxidized ammonia differed greatly between growth types due to the early transfer of biofilm cultures (30°C, 20 mM NH_4_ ^+^, 30 mM NH ^+^, 1.1 µM ethoxyquin, 1.3 µM ethoxyquin, and starvation 1 week) or differing stationary phase end points (dessication 1 day), n_max_ and t_max_ were calculated at comparable concentrations of produce nitrite (see Supplementary Material S3). A visualization of all compared conditions and associated points used for maximum rates can be found in Supplementary Material S3-Rate Comparisons.

#### Statistical Analysis of Maximum Rates

Biofilm replicates were compared to their planktonic counterparts for each stress condition. With both cases having n=3, tests for normality difficult to rely on. Additionally, non-parametric tests are too limited in power to yield informative *P* values. Therefore, statistical significance was determined based on a complete separation of values (i.e., all of one condition greater than all of the other condition) and for n_max_, a ratio of the average biofilm values to the average of planktonic values that is greater than 1.5, and for t_max_ a difference in means greater than 1.0 days. Carrots are used to identify significant biofilm to planktonic ratios for n_max_ at the following levels: ^ 1.5-2.5 and ^^ 2.5-5.0. Plus signs are used to indicate a significant differences of means for t_max_ at the following levels: +1.0-3.0 days, ++3.0-7.0 days, +++ >7. A full list of whether values showed complete separation, biofilm to planktonic ratios (n_max_), and differences of means (t_max_) can be found in supplementary material (Dataset S1). Data was processed using the tidyverse statistical package^93^ in R.

### Substrate-dependent oxygen consumption measurements

Oxygen consumption rates of an *N. viennensis* biofilm formed in a MicroRespiration glass chamber (described above) were directly measured using a MicroRespiration system (Unisense), consisting of an O_2 M_icroOptode, a temperature sensor, a microsensor amplifier (fx-6 UniAmp), a rack (MR-2), a stirrer controller (MR2-Co), and a glass coated magnet. The sensor was calibrated at 42°C a day before measurements.

The chamber was transferred to prewarmed ammonia-free growth medium and let equilibrate for 3 mins to remove any remaining ammonia. The chamber was then removed from the growth medium and residual liquids carefully poured out. Before the start of oxygen measurements, fresh, prewarmed ammonia-free growth medium was added, the chamber closed with the lid and put in a rack, which was transferred to a water bath at 42°C, ensuring that the chamber was fully submerged. The sensor was left to equilibrate until the signal was stable (1 - 1.5 hours), before a range of final total NH_4_ ^+^/NH_3 c_oncentrations, hereafter referred to as ammonium, were tested (5, 10, 20, 100, 200, 500, 1000 µM) in the same chamber using a 10 µl Hamilton syringe (Hamilton). Sequential ammonium addition was used to reach each concentration (from lowest to highest) and ammonia oxidation rates were calculated based on a thermodynamically calculated ammonia to oxygen consumption ratio of 1:1.4^94^. Higher ammonium was added once rates of the previous measurements were stable. Once 1000 µM of total ammonium was added the measurement was continued until all oxygen was consumed to ensure that decreasing O_2_ concentrations did not affect the measurement. After finishing the measurements, the lid was carefully taken off and all liquid transferred to sterile eppendorf tubes. To remove the biofilm, the internal chamber surface was rigorously washed with 1 ml growth medium using a 2 ml syringe and needle, and transferred to sterile Eppendorf tubes. Both the liquid and biofilm fraction were fixed as described above and cell numbers counted using FACS.

To compare the activity of biofilms and planktonic cells of *N. viennensis*, planktonic controls with the same corresponding cell numbers were treated as described above. Measurements were done in fully filled (∼2 ml), headspace-free chambers containing a stirrer set to 300 rpm, submerged in a recirculating water bath. Oxygen consumption graphs for planktonic and biofilm cultures can be found in Fig. S11. Data points used for the calculation of rates can be found in Fig S12 (planktonic) and Fig. S13 (biofilm).

### Michaelis-Menten kinetics

K_m(app) a_nd V_max w_ere calculated from ammonia consumption rates derived from oxygen consumption measurements assuming a thermodynamically calculated ammonia to oxygen consumption ratio of 1:1.4^94^ to, which were fitted to a Michaelis-Menten equation:

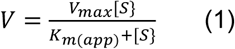

V is the reaction rate (µM NH_3/_hr), V_max i_s the maximal reaction rate (µM NH_3/_hr), S the ammonium concentration (µM NH_4+_/NH_3)_ and K_m(app) t_he half saturation concentration (µM NH_4+_/NH_3)_. K_m(app) a_nd V_max w_ere calculated using a nonlinear least squares regression analysis in R.

### Nitrous oxide

Biofilm and planktonic cultures containing an equal number of cells grown in quintuplicates were grown in sealed schott bottles at 42°C and 80 rpm as described above. After producing ∼1700 - 1900 µM nitrite, 15 ml headspace gas was sampled and transferred to sealed glass vials using a syringe. The sample headspace N_2O_ concentrations were determined with a Trace Gas Chromatograph (TRACE Ultra Gas Chromatograph, Thermo Scientific, Germany) equipped with a pulsed discharge ionization detector. Results were tested for normality and homogeneity of variance before being analyzed with a t-test to determine if differences were significant.

### Imaging F_420_ in Biofilms and Aggregates

Biofilms of *Nitrososphaera viennensis*, *Nitroscosmicus franklandianus*, *Nitrosopumilus maritimus*, and *Nitrosopumilus adriaticus*, were grown as biofilms as previously described^57^. *Nitrosocosmicus arcticus* was grown as previously described^17^ and aggregates were collected for imaging. The strain ‘*Candidatus* Nitrosocosmicus epidermidis’ was grown as a biofilm with methods described here and in Dreer et al.^57^ in medium previously described for this strain^95^. Biofilms or aggregates were imaged using phase contrast light microscopy and F_420_ autofluorescent images were taken with a standard Nikon fluorescent filter cube for 4’,6-diamidino-2-phenyldole (DAPI).

## Data Availability

All raw data for this manuscript can be found at the following GitHub page: https://github.com/hodgskiss/Nviennensis_Biofilm_vs_Planktonic_Cells.

## Acknowledgements

We thank Evangelia S. Papadopoulou and Dimitrios Dalkidis (Department of Environmental Sciences, University of Thessaly) for providing guidance on nitrification inhibitor concentrations and supplying ethoxyquin. We also acknowledge and thank Margarete Watzka and Wolfgang Wanek (Division of Terrestrial Ecosystem Research, Center of Microbiology and Environmental Systems Science, University of Vienna) for headspace N_2_O analysis.

