## Supplementary Material for "Ecophysiological niche expansion driven by biofilm lifestyle in an archaeal soil nitrifier"

**Included in supplementary material:**

Figures S1-S12

Supplementary Material S2-Models

Supplementary Material S3-Rate Comparisons

Dataset S1-max rate and time data

### Supplementary Figures

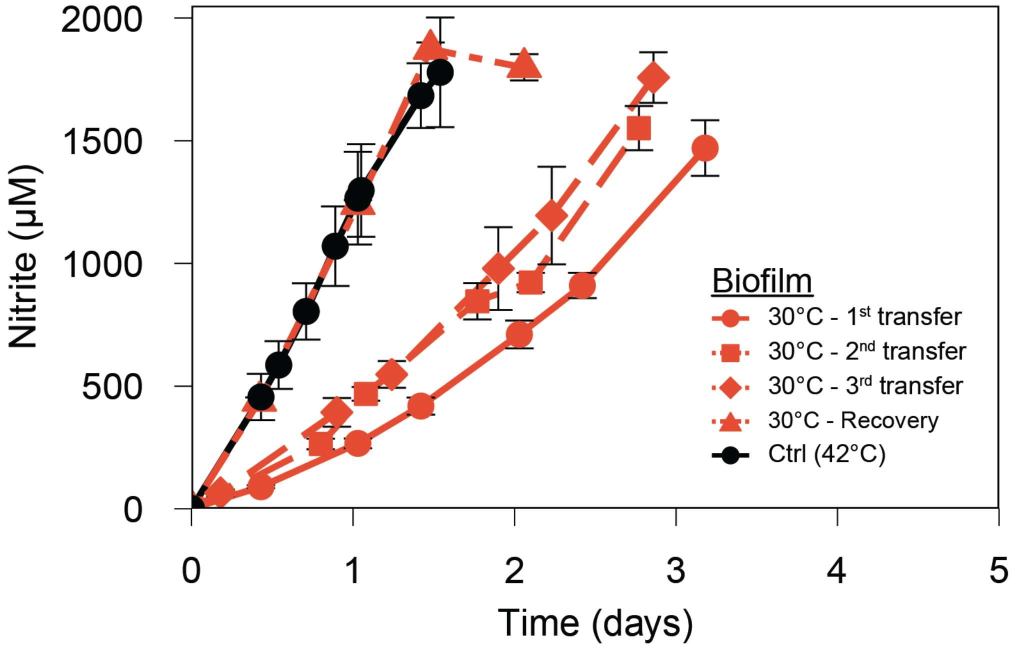


**Figure S1. Biofilm resilience to 30°C**. Growth of *N. viennensis* biofilms were transferred three times before they recovered from 30°C. Biofilms were recovered (triangle) by transferring microscopy slides (MS) back to optimal growth temperature. Data points represent averages ± standard deviation as error bars (biofilm controls, n=9; all others, n=3). All maximum rates and times are provided in Dataset S1.

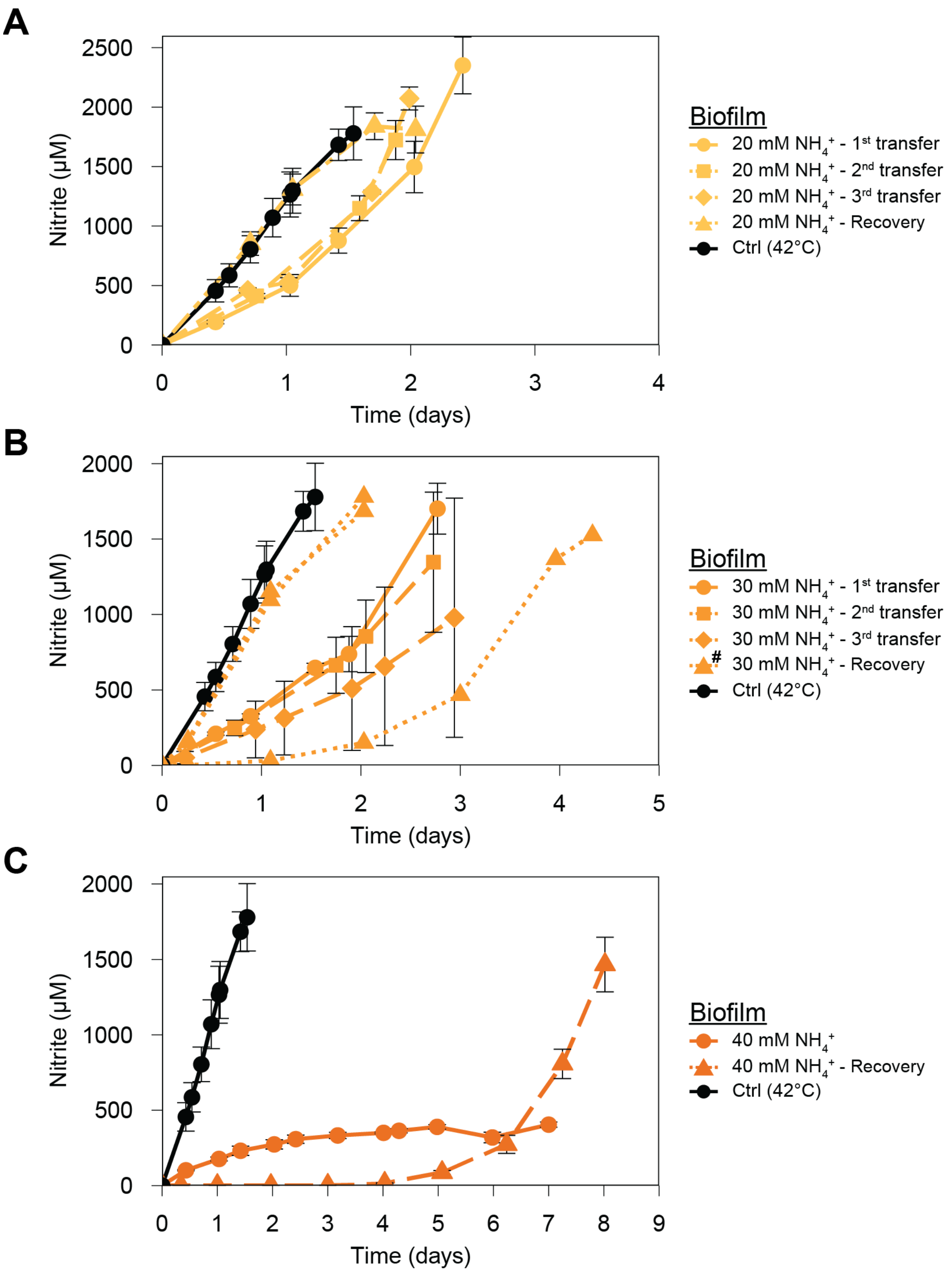


**Figure S2. Biofilm resilience (20 mM and 30 mM $NH_4^+$) and recovery from high ammonium concentrations (20mM, 30 mM, and 40 mM $NH_4^+$).** Growth of *N. viennensis* biofilms recovered from different ammonia concentrations: **(A)** 20 mM $NH_4^+$, **(B)** 30 mM $NH_4^+$, and **(C)** 40 mM $NH_4^+$. Biofilms were recovered (triangle) by transferring microscopy slides (MS) back to optimal growth medium with 2 mM $NH_4^+$ Biofilms grown in the presence of 20 and 30 mM (circle) included two additional transfers (rectangle, diamond) at the respective ammonia concentrations before being recovered to optimal medium (triangle). Growth curves where replicates greatly diverged are plotted individually as dotted lines and marked with (#) in the legend. In all panels, if not plotted individually, data points represent averages ± standard deviation as error bars (biofilm controls, n=9; all others, n=3). All maximum rates and times are provided in Dataset S1.

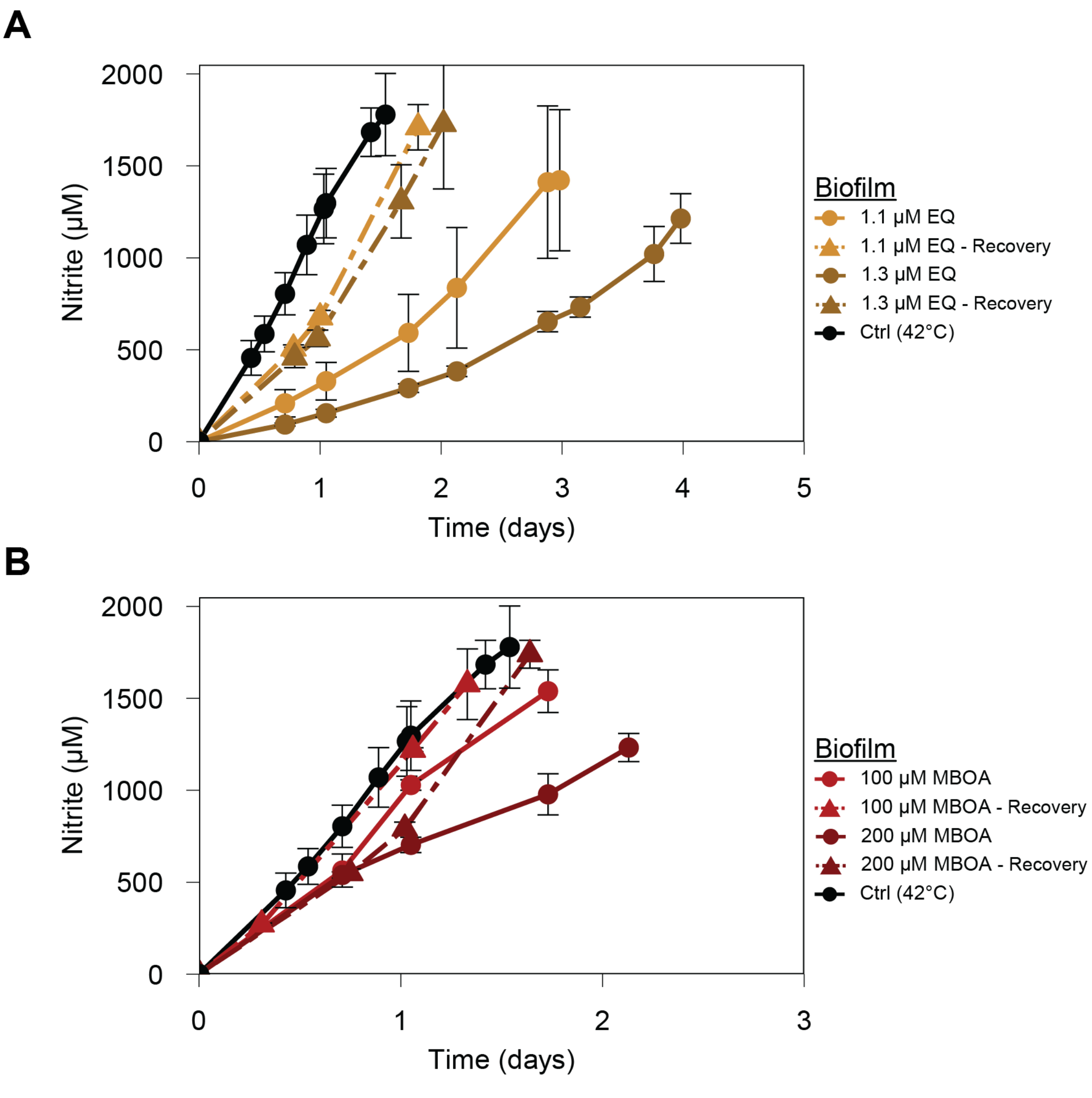


**Figure S3. Biofilm recovery from nitrification inhibitors (NI).** Growth of *N. viennensis* biofilms recovered from: **(A)** 1.1 and 1.3 µM EQ and **(B)** 100 and 200 µM MBOA. Biofilms were recovered (triangle) by transferring microscopy slides (MS) back to optimal growth medium. In all panels, data points represent averages ± standard deviation as error bars (biofilm controls, n=9; all others, n=3). All maximum rates and times are provided in Dataset S1.

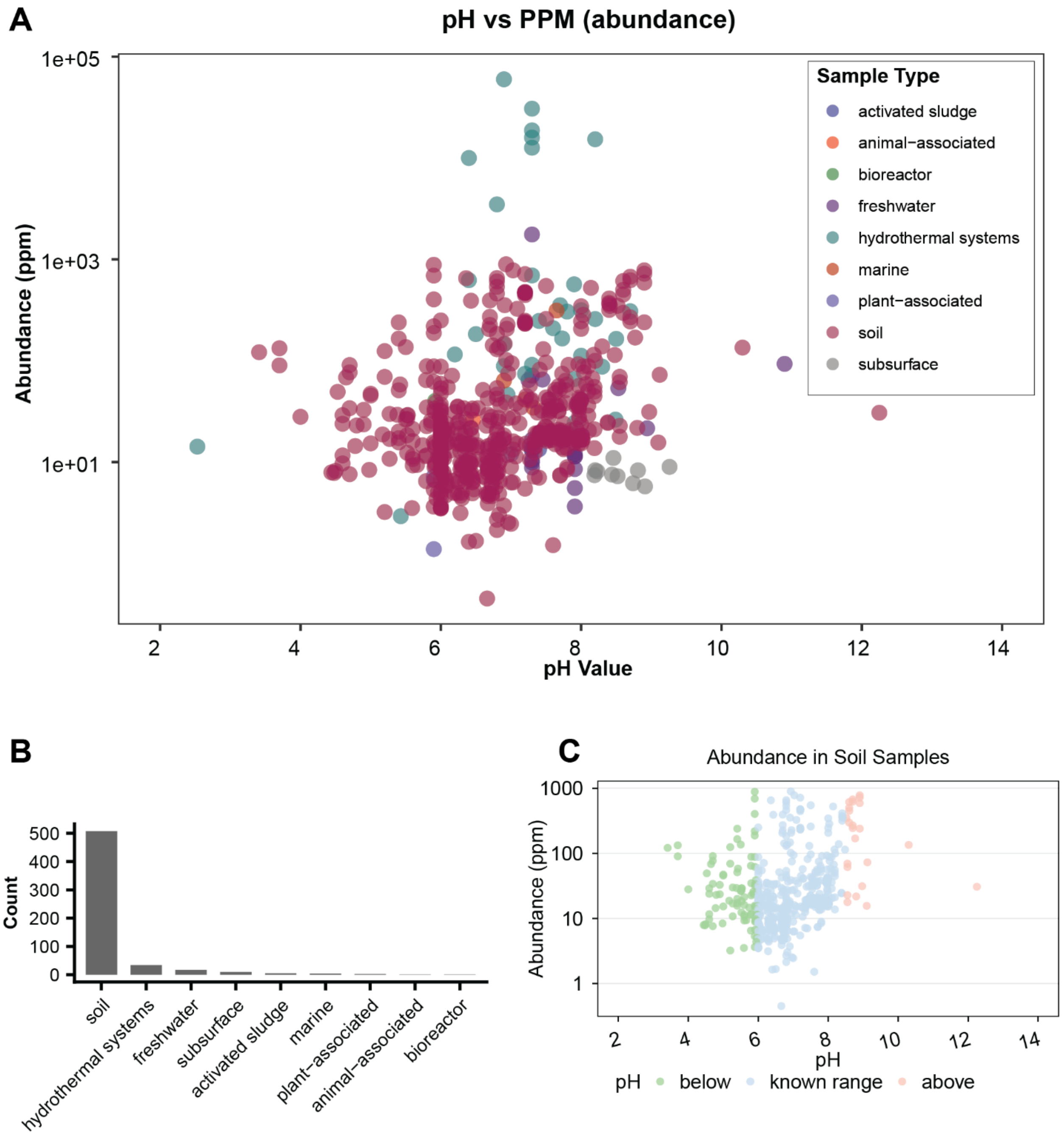


**Figure S4. Abundance of *N. viennensis* in samples of varying pH.** **(A)** Abundance of *N. viennensis* at different pH values across different sample types. **(B)** A histogram representing the number of sample types that *N. viennensis* was detected in. **(C)** Abundance of *N. viennensis* specifically in soil samples at varying pH values. Data downloaded from https://microbeatlas.org/ .

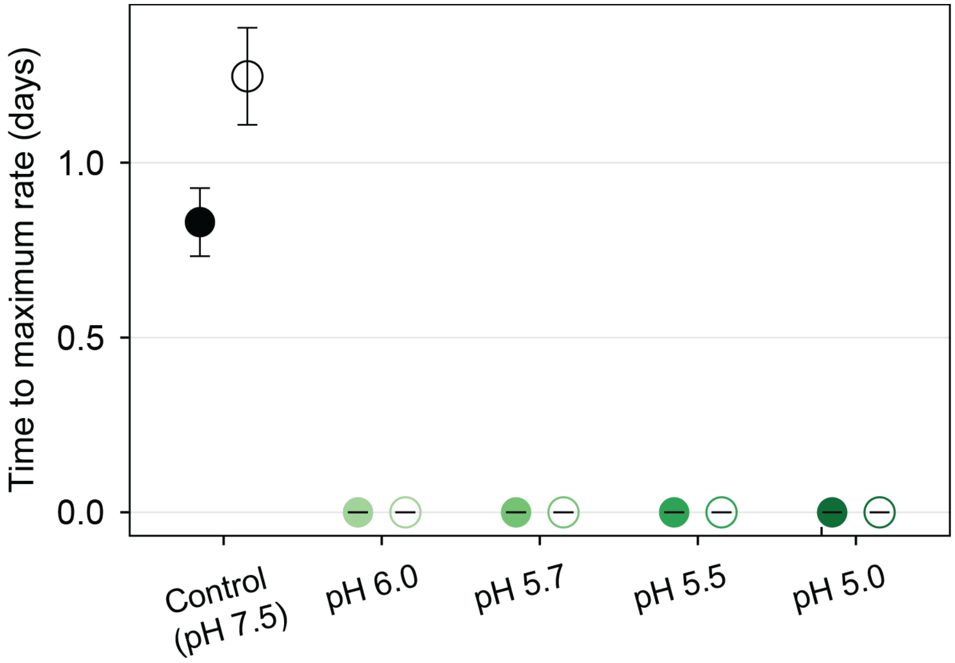


**Figure S5. Time to maximum nitrification rate ($t_{max}$) for controls and biofilms at pH of 6.0, 5.7, 5.5, and 5.0.** Filled and open symbols indicate biofilms and planktonic cells respectively. Data points represent averages ± standard deviation as error bars (biofilm controls, n=9; planktonic controls n=6; all others, n=3). In all cases of pH tests cubic models were used which placed the $n_{max}$ at a time of zero.

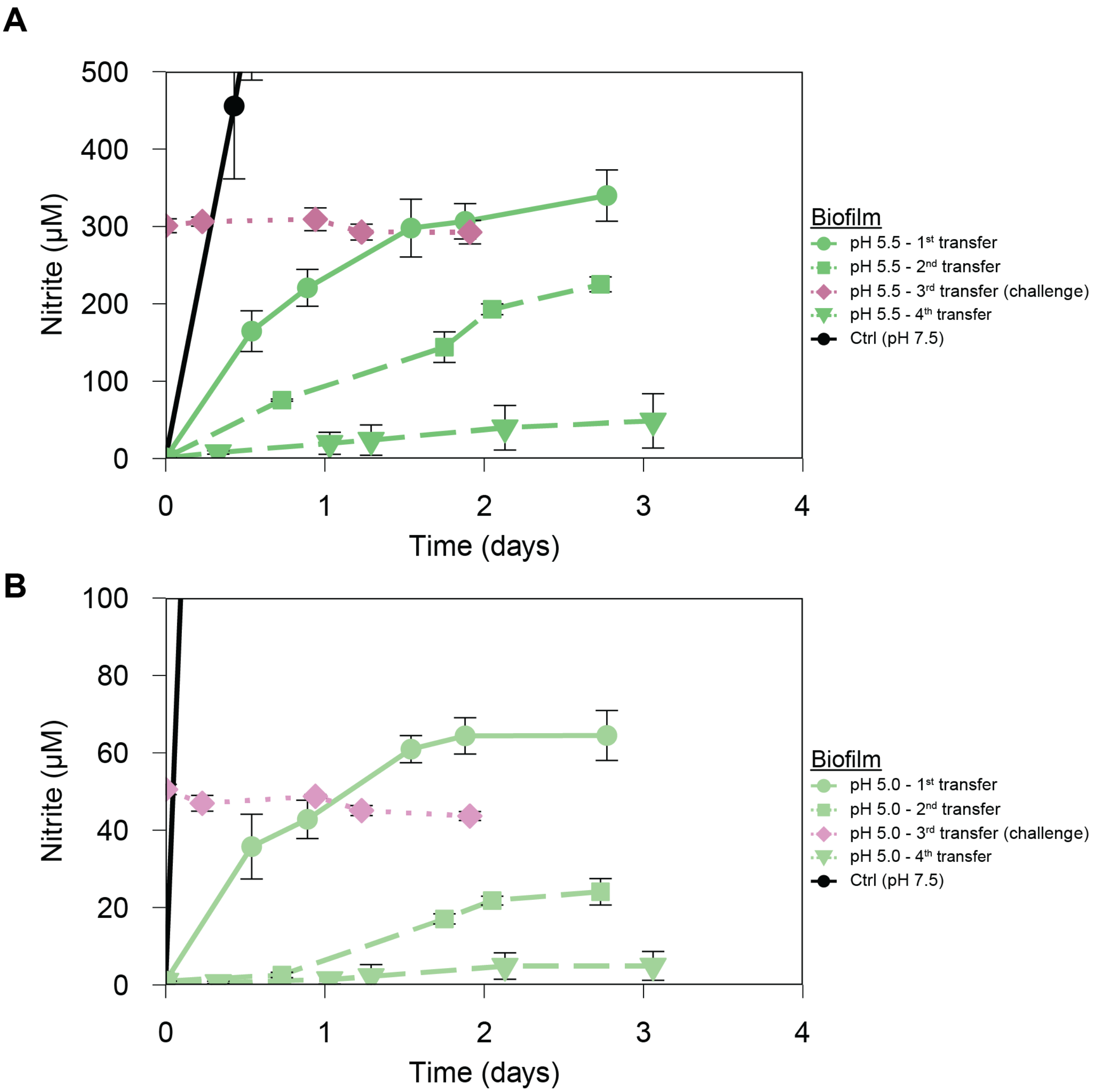


**Figure S6. Resistance of biofilms at pH 5.5 and 5.0.** Growth of *N. viennensis* biofilms at **(A)** pH 5.5 and **(B)** pH 5.0 following the same transfer sequence. Biofilms were transferred twice at the respective pH (circle, rectangle), followed by a third transfer into medium containing the maximum nitrite concentration tolerable at that pH (diamond, challenge), and a subsequent transfer without nitrite (inverted triangle). Challenges are indicated in shades of red. In all panels, data points represent averages ± standard deviation as error bars (biofilm controls, n=9; all others, n=3). All maximum rates and times are provided in Dataset S1.

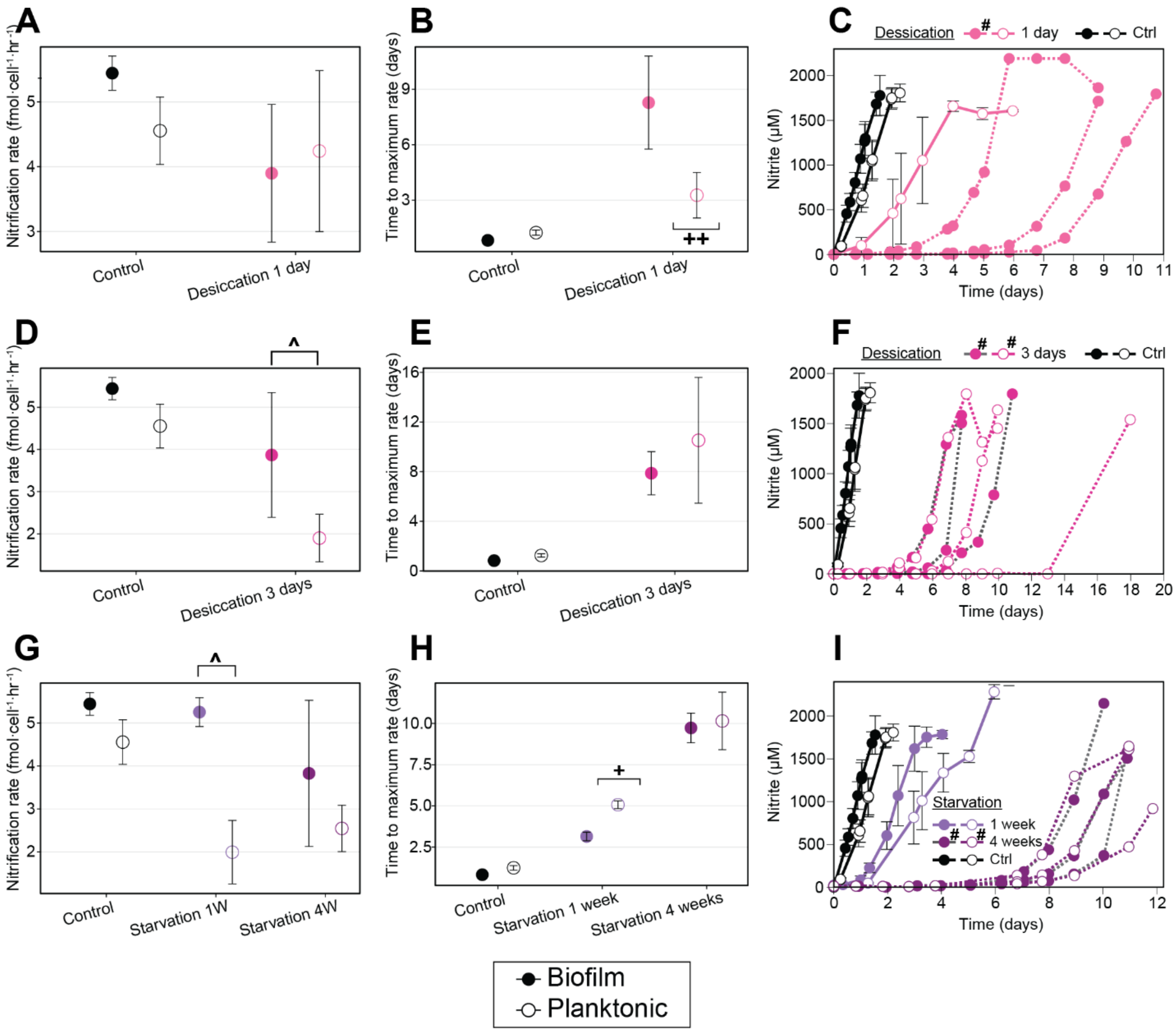


**Figure S7. Biofilm resilience to desiccation and ammonia starvation. (A,D,H)** Maximum ammonia oxidation rates ($n_{max}$) per cell (fmol cell$^{-1}$ h$^{-1}$) after one day of desiccation, three days of desiccation, and one or four weeks of ammonia starvation, upon recovery to optimal growth medium. Rates were calculated per replicate. Carrot indicates significant biofilm to planktonic ratios: (^ 1.5-2.5, ^^ 2.5-5.0). **(B, E, H)** Time to reach maximum rates ($t_{max}$) after the same periods of desiccation, or starvation as in (A,D,H respectively), calculated per replicate; shown are averages ± standard deviation. Plus signs indicated significant differences of means: (+1.0-3.0, ++3.0-10.0, +++ >10). **(C,F,I)** Nitrite production of *N. viennensis* biofilms or planktonic controls. Planktonic cultures were inoculated with cell numbers matching those in the biofilms. Filled and open symbols indicate biofilms and planktonic cells respectively. Growth curves where replicates greatly diverged are plotted individually as dotted lines and marked with (#) in the legend. . In all panels, if not plotted individually, data points represent averages ± standard deviation as error bars (biofilm controls, n=9; planktonic controls n=6; all others, n=3). All maximum rates and times are provided in Dataset S1.

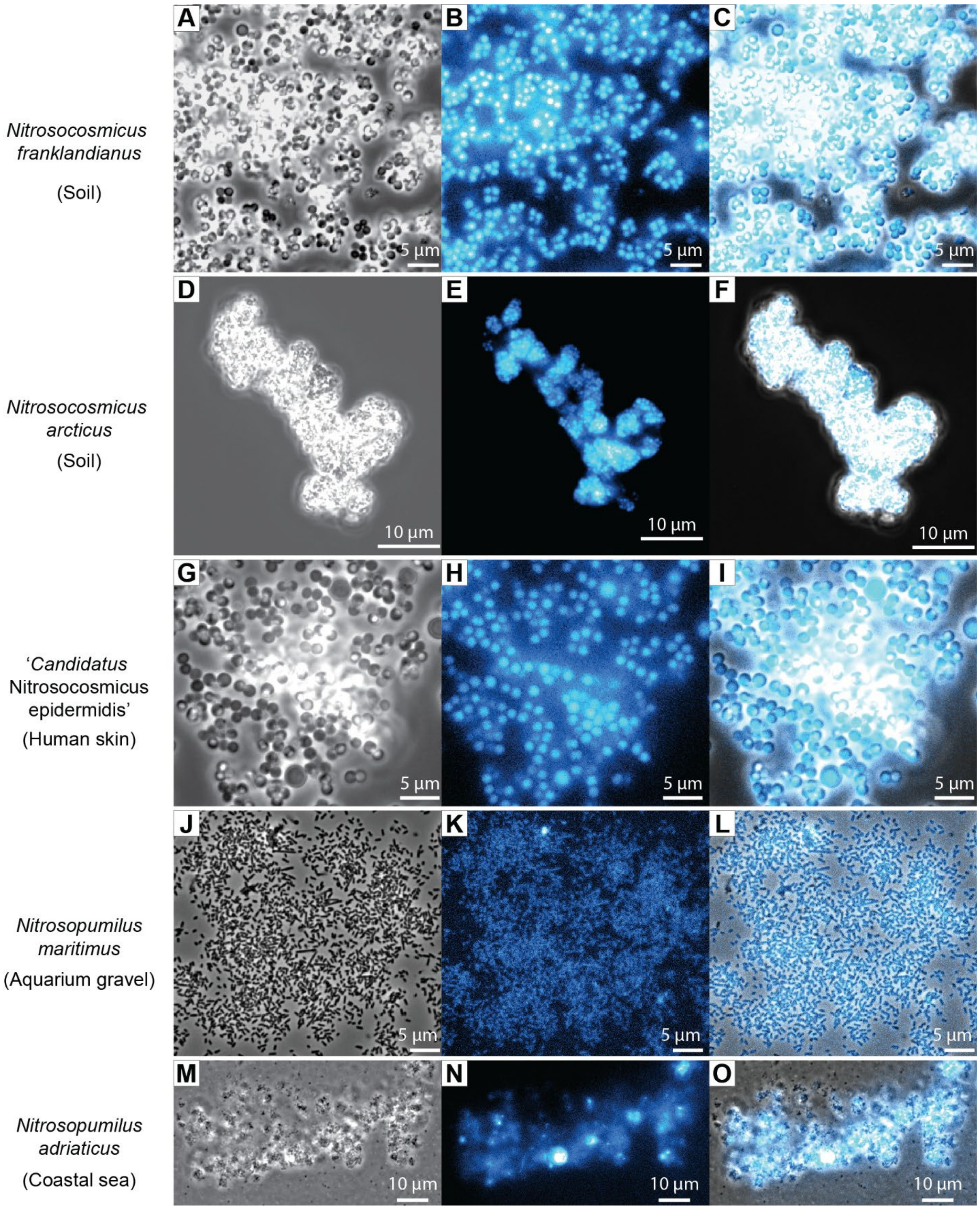

A
B
C
Nitrosocosmicus franklandianus
(Soil)
D
E
F
Nitrosocosmicus arcticus
(Soil)
G
H
I
'Candidatus Nitrosocosmicus epidermidis'
(Human skin)
J
K
L
Nitrosopumilus maritimus
(Aquarium gravel)
M
N
O
Nitrosopumilus adriaticus
(Coastal sea)
5 µm
10 µm

**Figure S8. Cofactor $F_{420}$ autofluorescence of AOA biofilm and aggregates. (A,D,G,J,M)** Phase contrast light microscopy **(B,E,H,K,N)** Fluorescence microscopy images based on autofluorescence due to cofactor $F_{420}$. Images were taken with a standard Nikon fluorescent filter cuber for DAPI and are false colored in Cyan. **(C,F,I,L,O)** Overlays of corresponding phase contrast and fluorescent images. Represented strains are *N. franklandianus* **(A-C)**, *N. arcticus* **(D-F)**, '*Ca.* N. epidermidis' **(G-I)**, *N. maritimus* **(J-L)**, and *N. adriaticus* **(M-O)**.

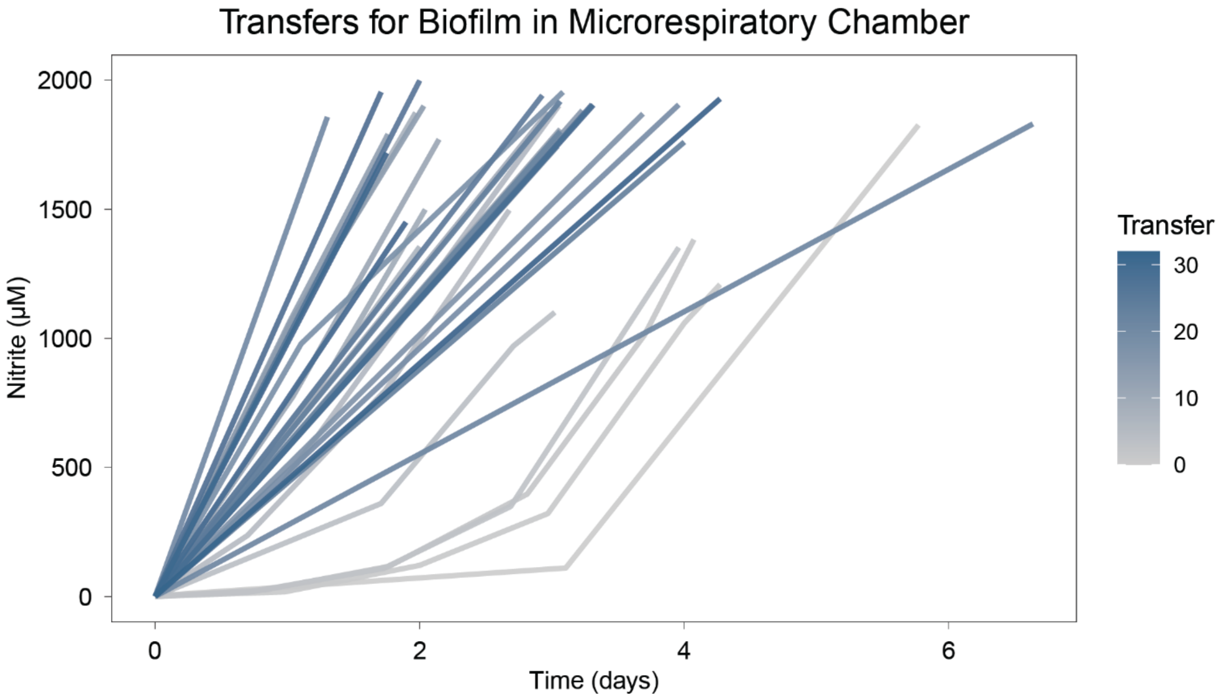


**Figure S9. Biofilm formation on microrespiratory chamber.** Nitrite production of the transfers of *N. viennensis* biofilms grown on microrespiratory chambers.

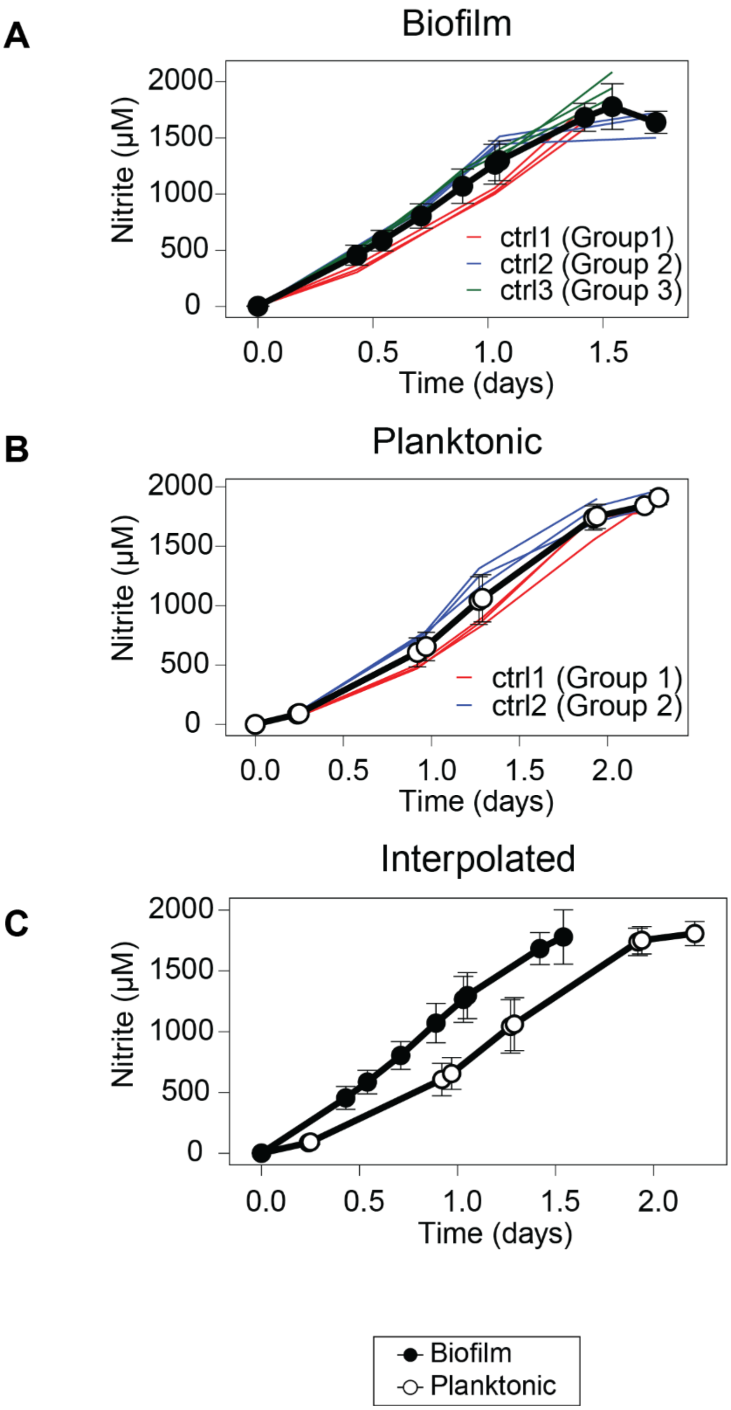


**Figure S10. Interpolated controls of biofilm and planktonic growth types. (A)** Control growth curves from the three separate biofilm groups. Interpolated growth curve is shown in black. **(B)** Control growth curves from the two separate planktonic groups. Interpolated growth curve is shown in black. Biofilm and planktonic controls were grown at different times (see Materials and Methods. **(C)** Interpolated controls used for all growth curves. Data points represent averages ± standard deviation as error bars (biofilm controls, n=9; planktonic controls n=6). Final points were removed as they only represent data from one group (see Materials and Methods).

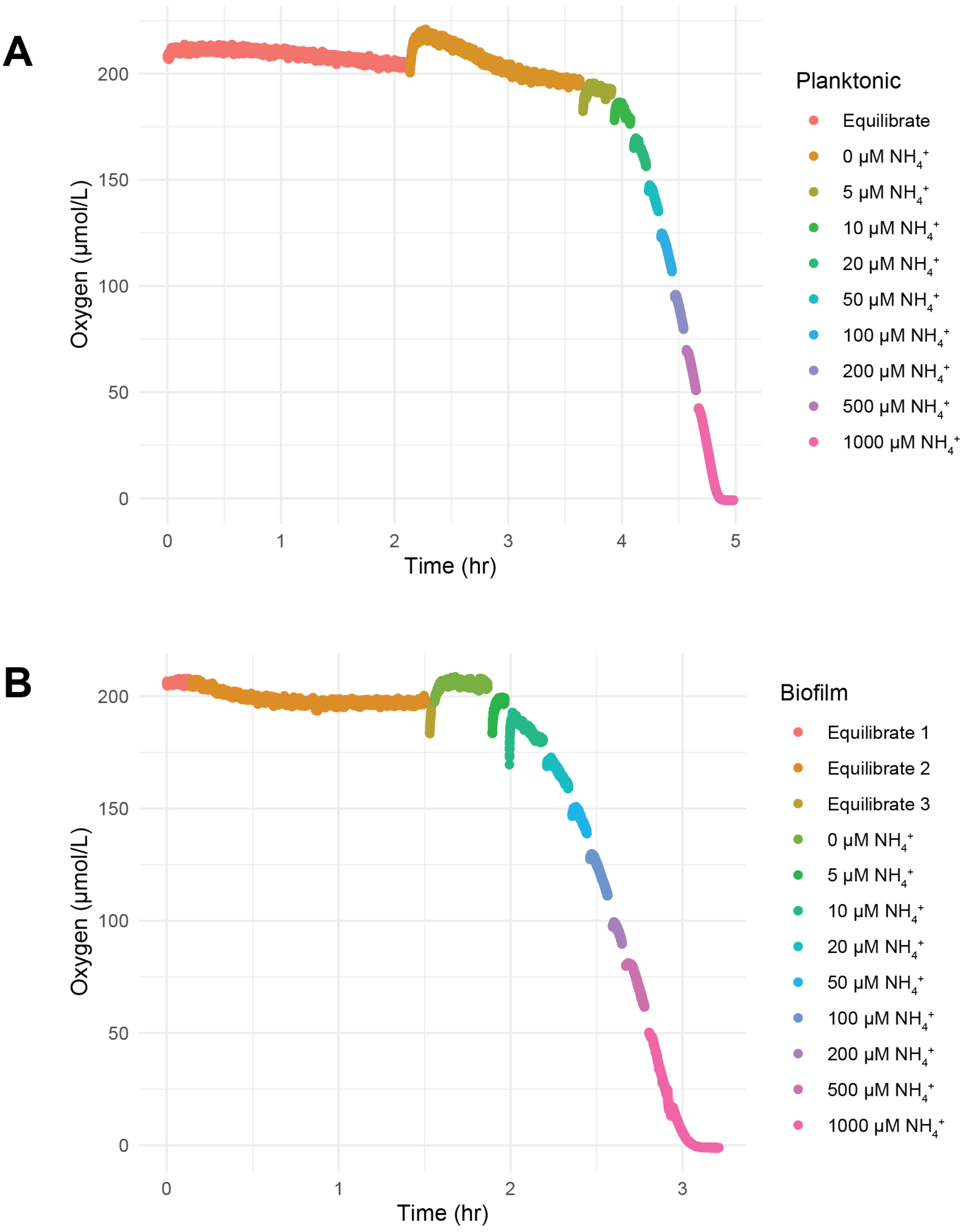


**Figure S11. Oxygen consumption of (A) planktonic and (B) biofilm cells in microrespiratory chambers.**

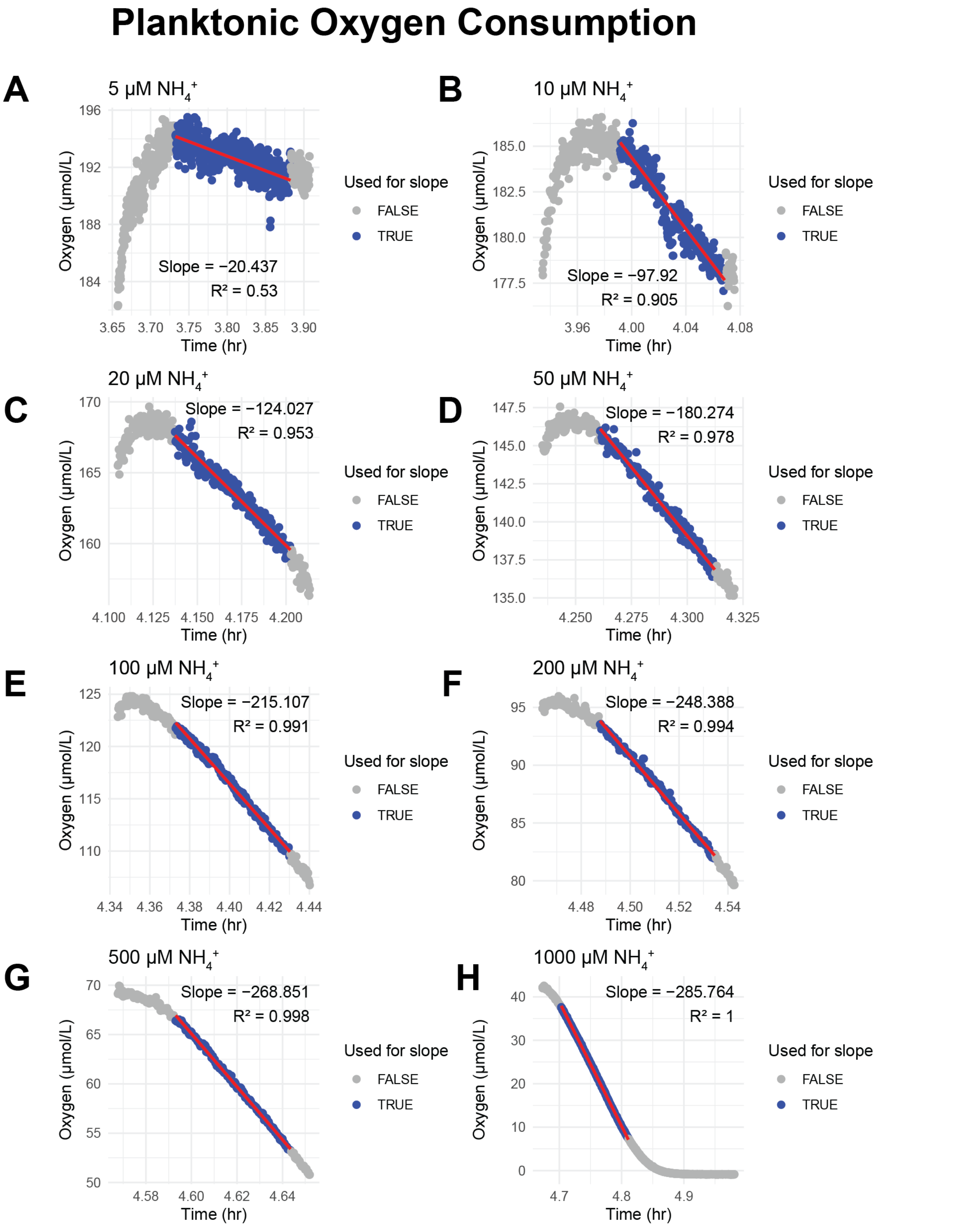


**Figure S12. Oxygen consumption rates for planktonic cells at varying ammonium concentrations.**

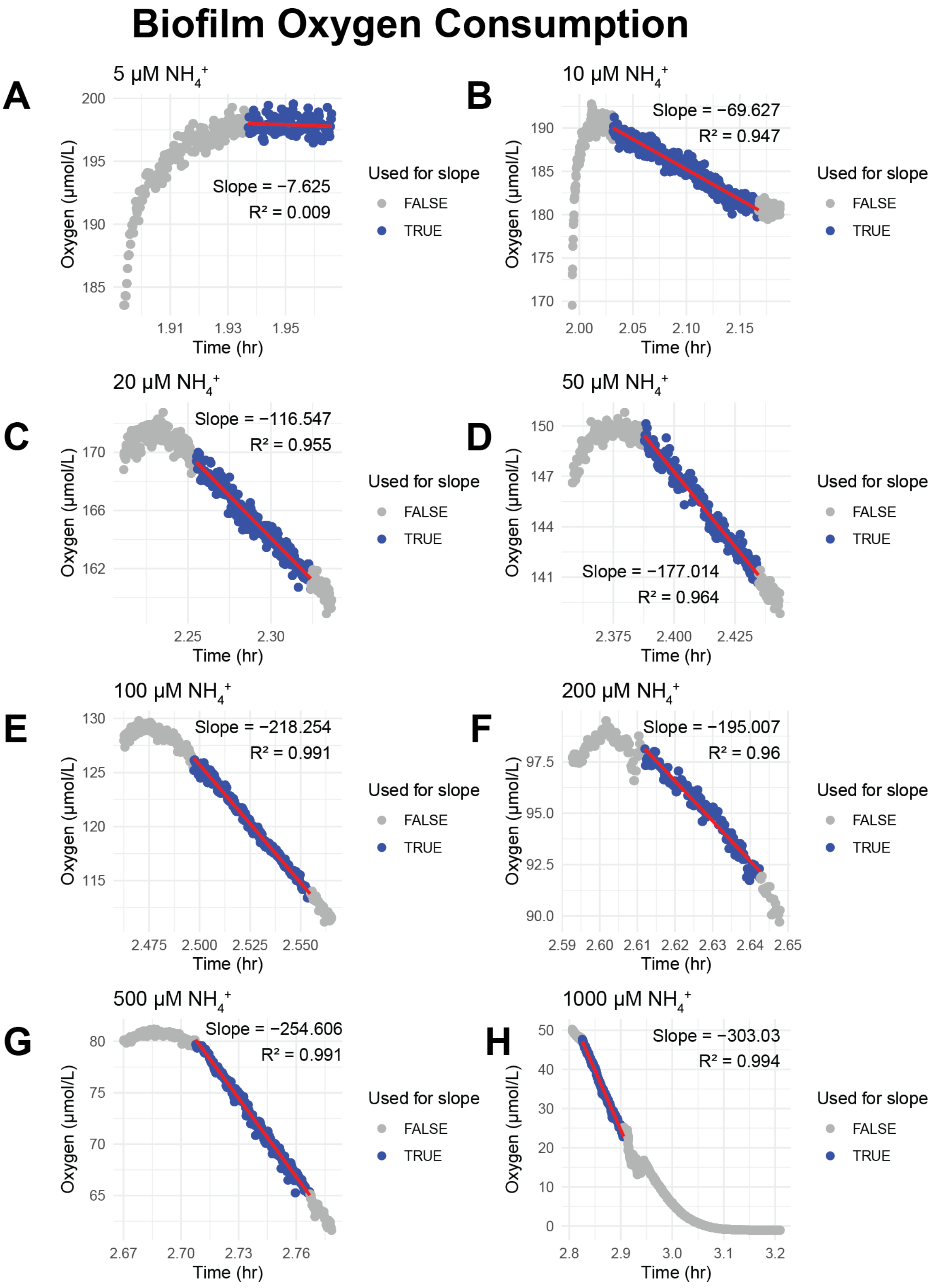


**Figure S13. Oxygen consumption rates for biofilm cells at varying ammonium concentrations.**
