## Supplementary Material S2 for "Ecophysiological niche expansion driven by biofilm lifestyle in an archaeal soil nitrifier"

**Models of Individual Growth Curves**

Each graph represents one condition of one growth type (biofilm or planktonic). Growth type, condition, and the chosen model (linear, quadratic, or cubic) are indicated in the figure titles. Dotted lines represent growth curves. Solid lines represent fitted models. Models only cover regions of the growth curve for which points were used. Dashed grey lines indicate points not used in the model for any of the three replicates. Large colored points represent the point in the model of the maximum rate ($n_{max}$).

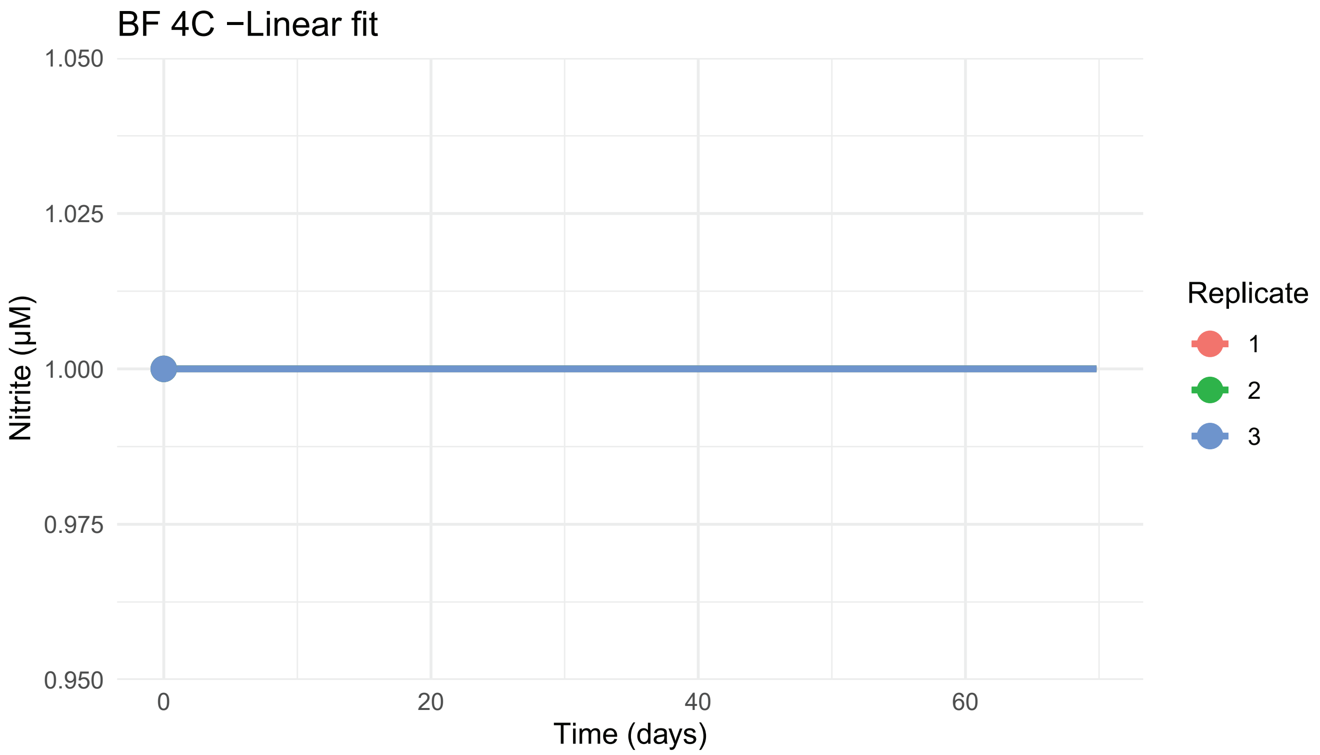
BF 4C −Linear fit
1.050
1.025
1.000
0.975
0.950
Nitrite (µM)
0
20
40
60
Time (days)
Replicate
1
2
3

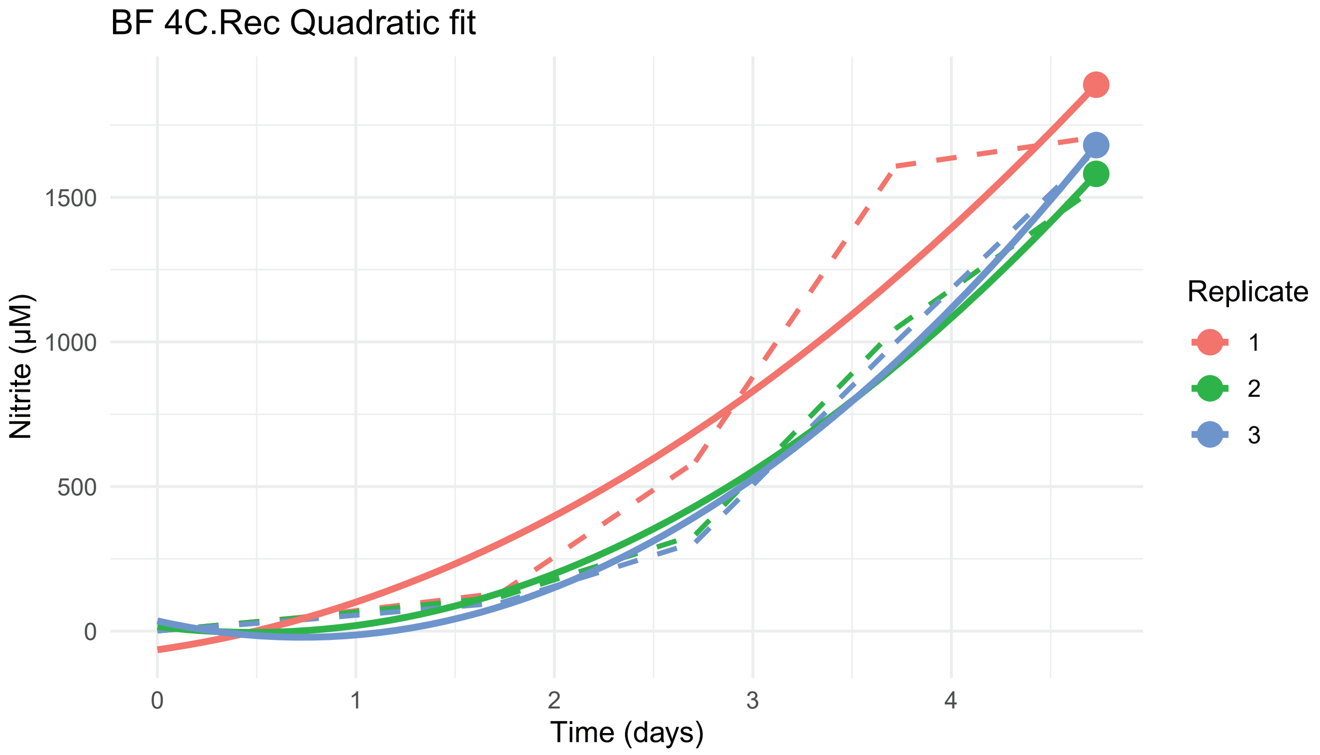
BF 4C.Rec Quadratic fit
1500
1000
500
0
Nitrite (µM)
0
1
2
3
4
Time (days)
Replicate
1
2
3

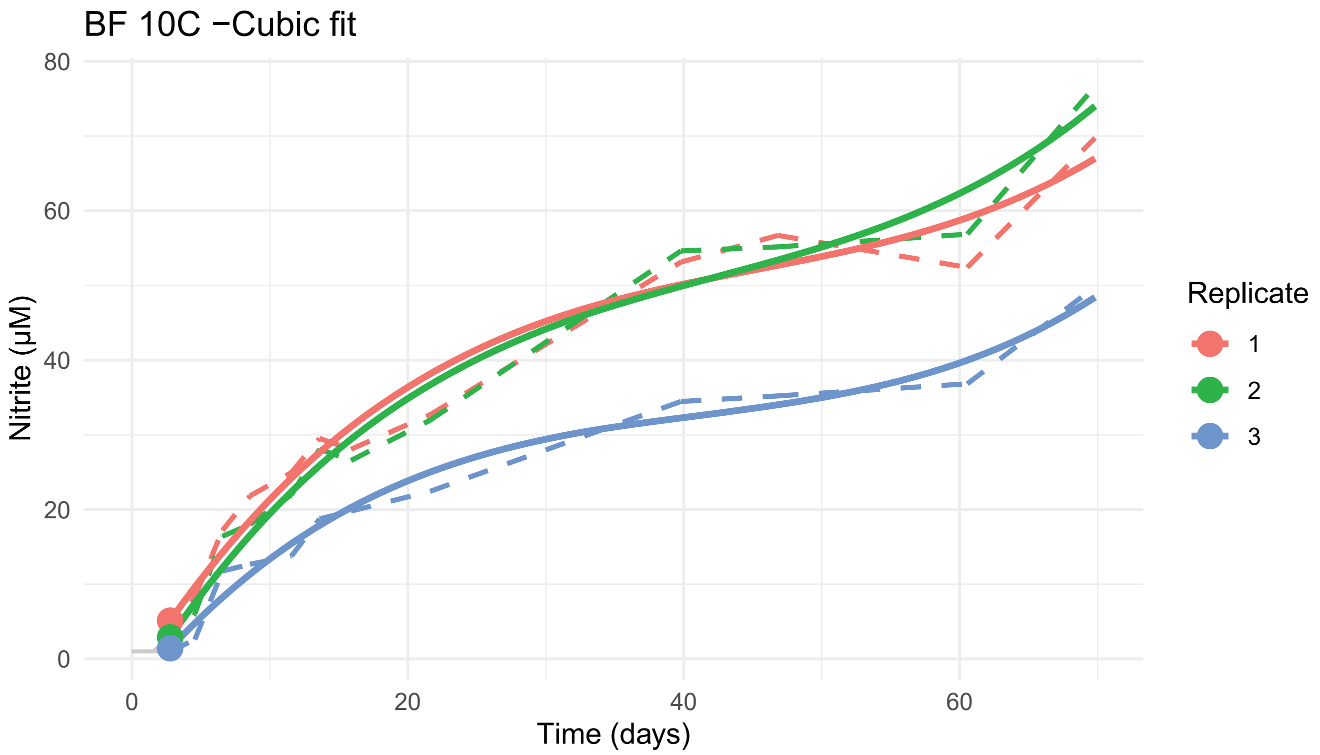
BF 10C −Cubic fit
80
60
40
20
0
Nitrite (µM)
0
20
40
60
Time (days)
Replicate
1
2
3

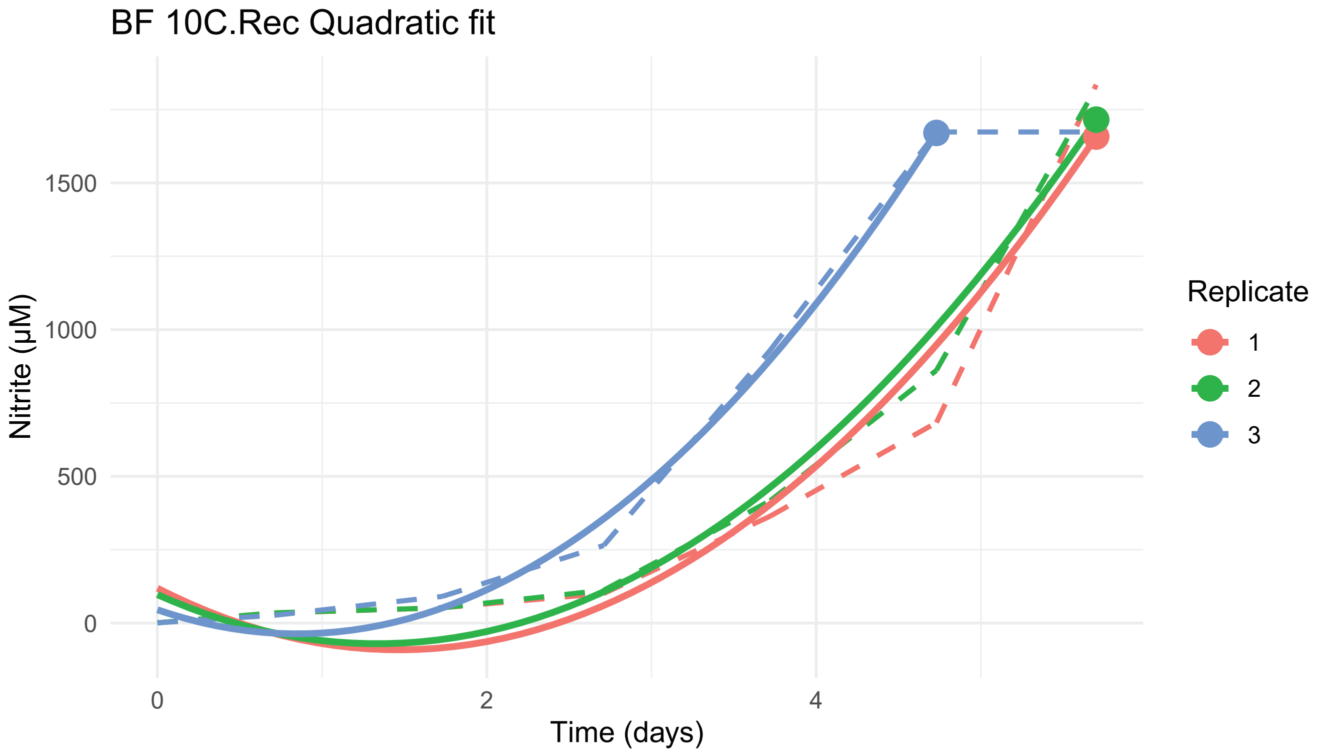
BF 10C.Rec Quadratic fit
1500
1000
500
0
Nitrite (µM)
0
2
4
Time (days)
Replicate
1
2
3

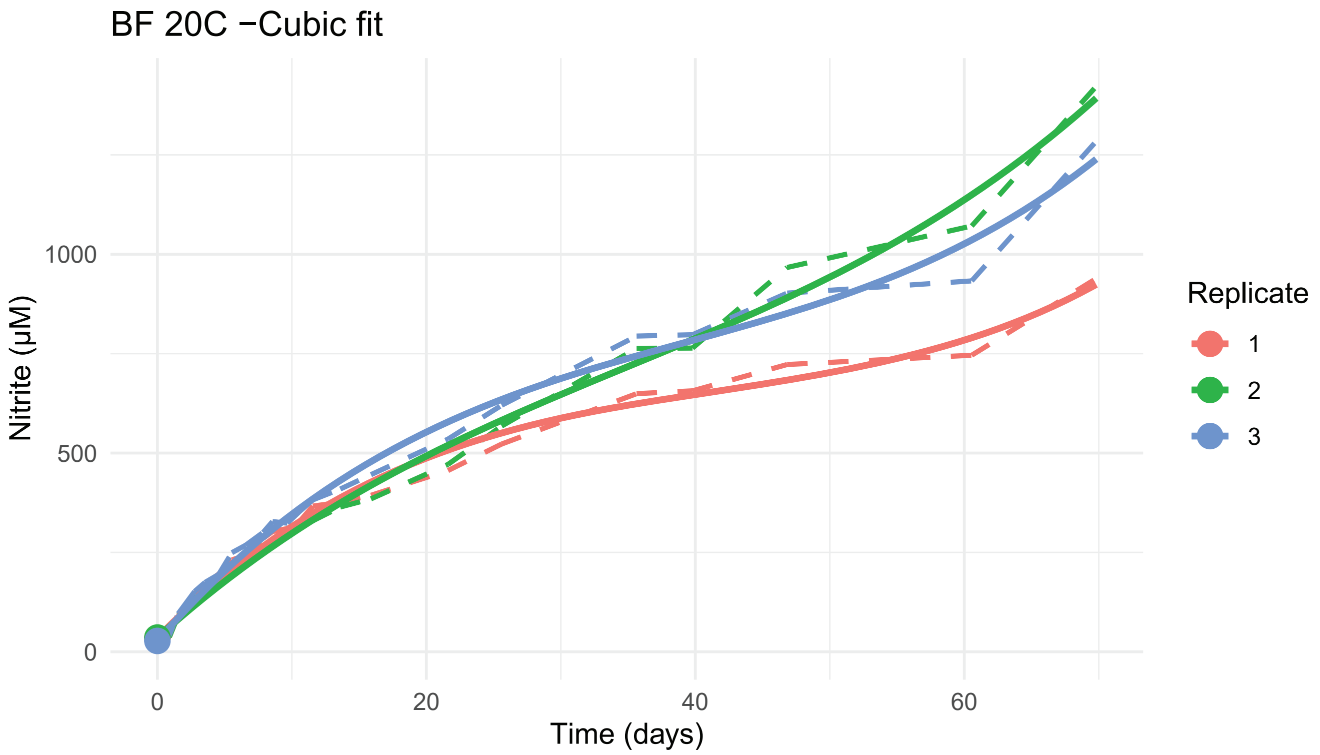
BF 20C −Cubic fit
Nitrite (μM)
0
500
1000
0
20
40
60
Time (days)
Replicate
1
2
3

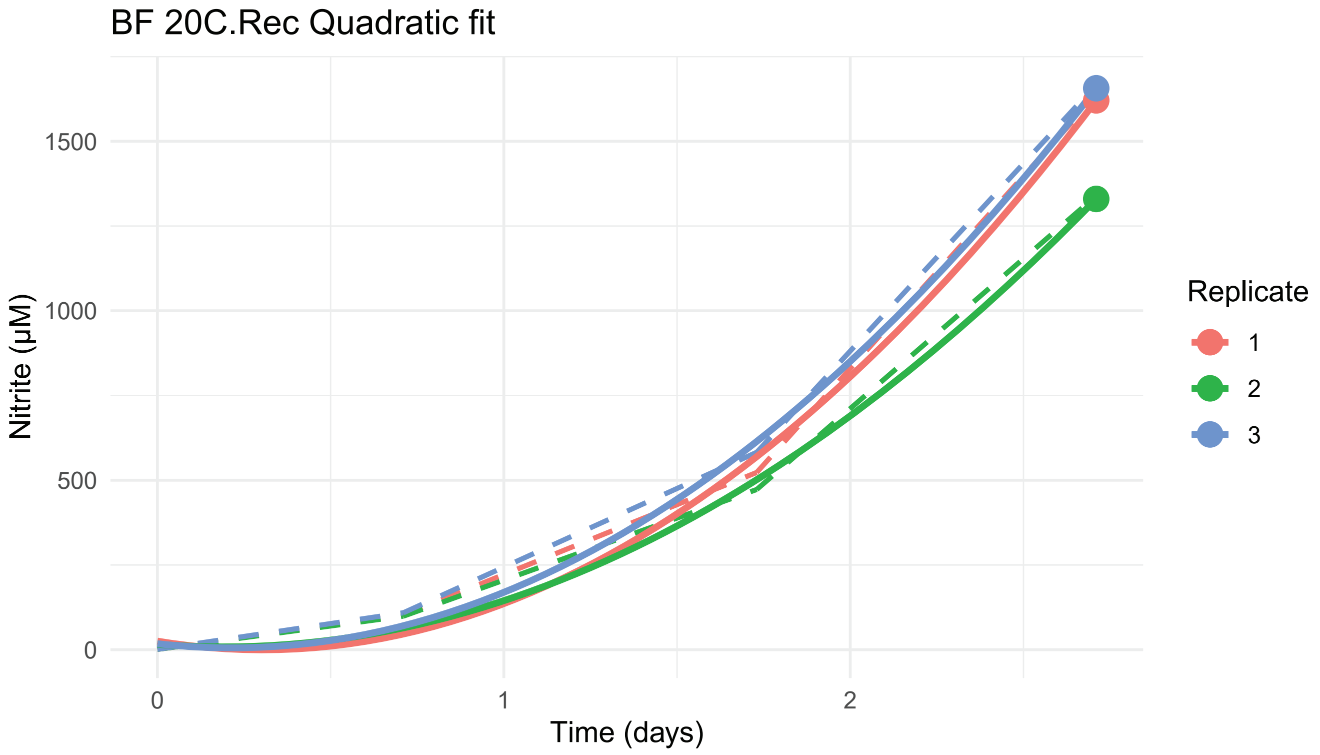
BF 20C.Rec Quadratic fit
Nitrite (μM)
0
500
1000
1500
0
1
2
Time (days)
Replicate
1
2
3

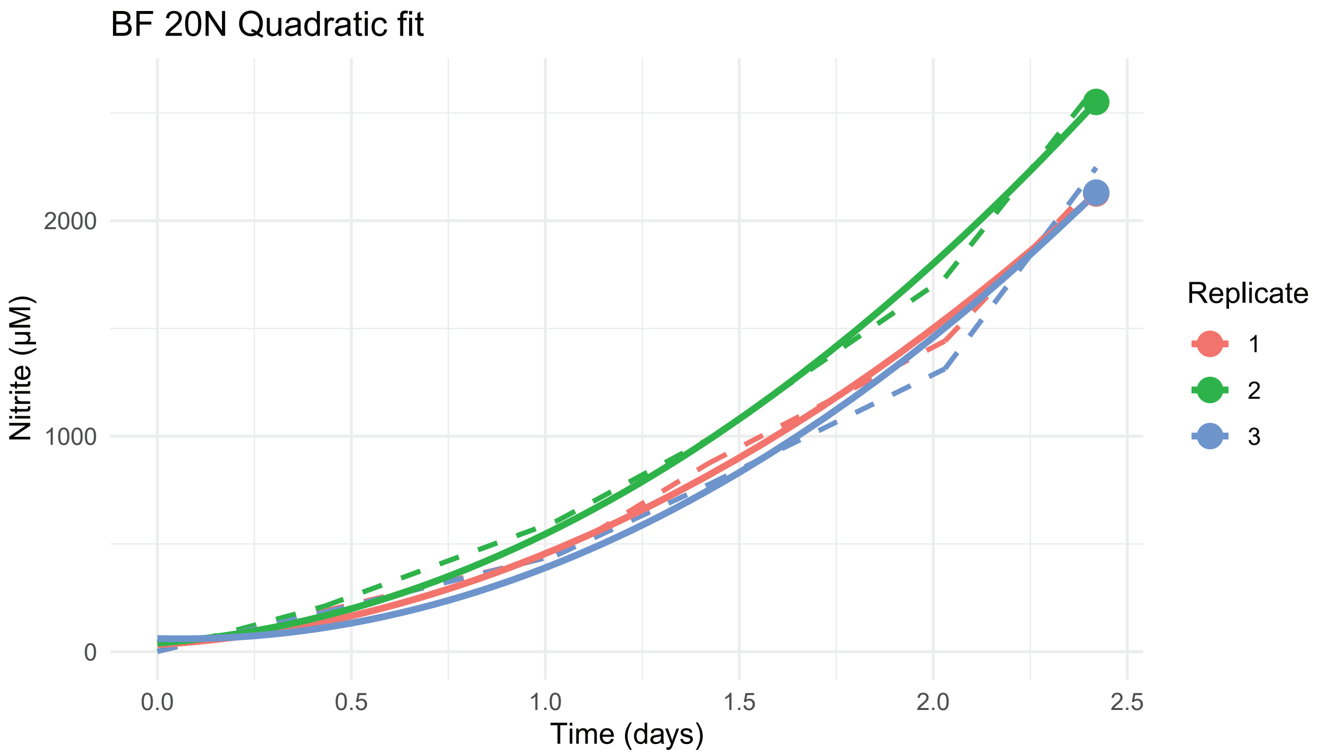
BF 20N Quadratic fit
Nitrite (µM)
2000
1000
0
0.0
0.5
1.0
1.5
2.0
2.5
Time (days)
Replicate
1
2
3

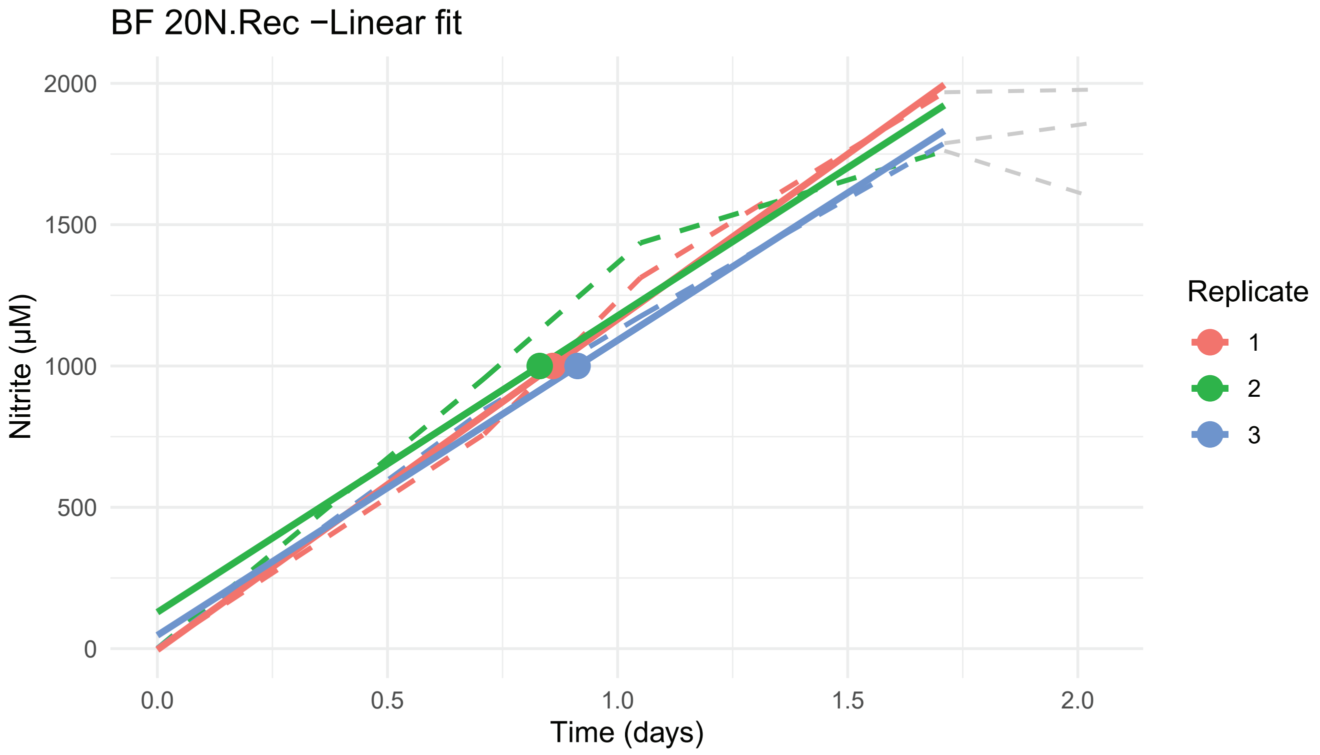
BF 20N.Rec −Linear fit
Nitrite (µM)
2000
1500
1000
500
0
0.0
0.5
1.0
1.5
2.0
Time (days)
Replicate
1
2
3

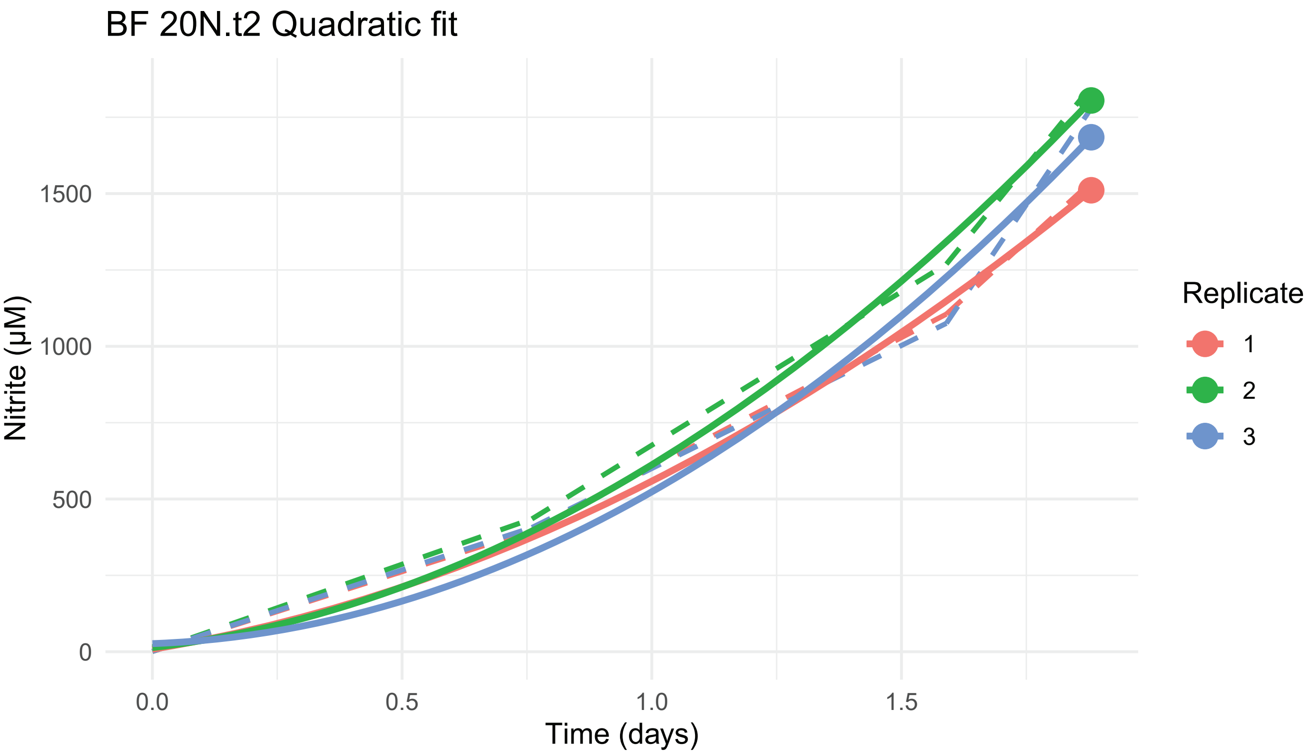
BF 20N.t2 Quadratic fit
Nitrite (µM)
0
500
1000
1500
0.0
0.5
1.0
1.5
Time (days)
Replicate
1
2
3

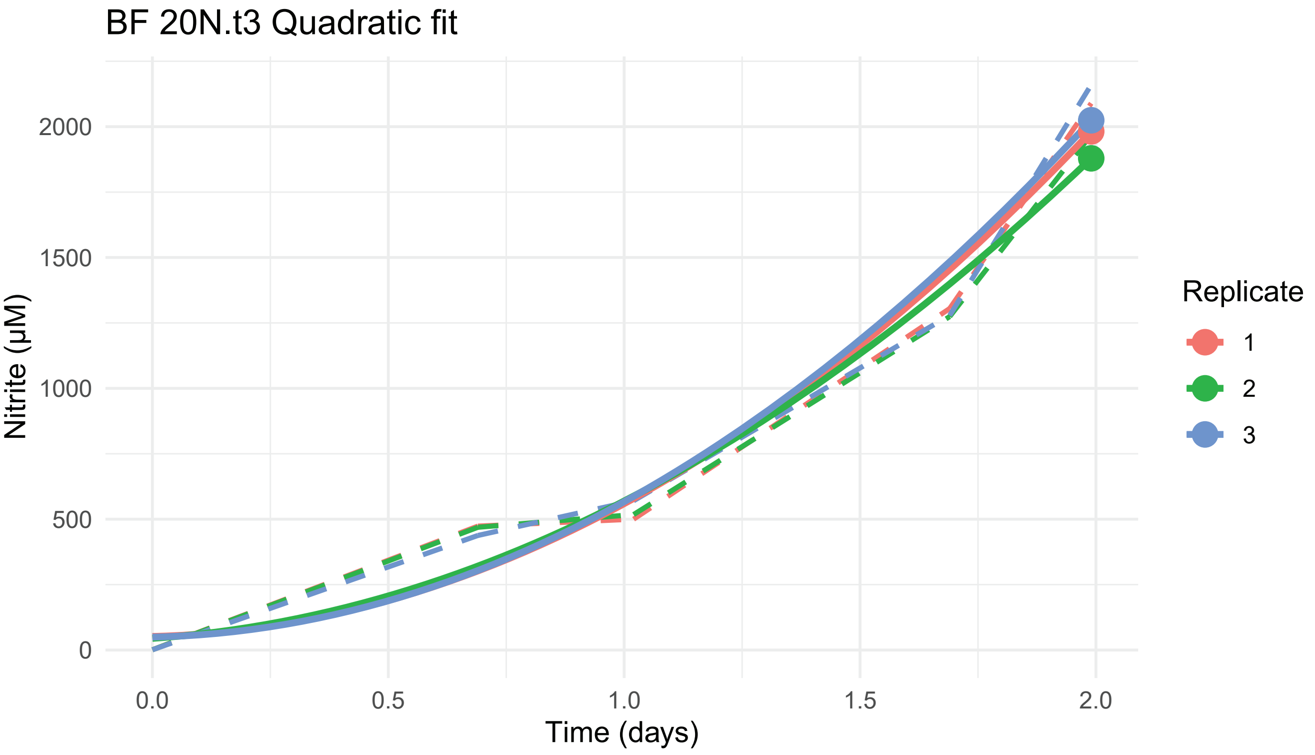
BF 20N.t3 Quadratic fit
Nitrite (µM)
0
500
1000
1500
2000
0.0
0.5
1.0
1.5
2.0
Time (days)
Replicate
1
2
3

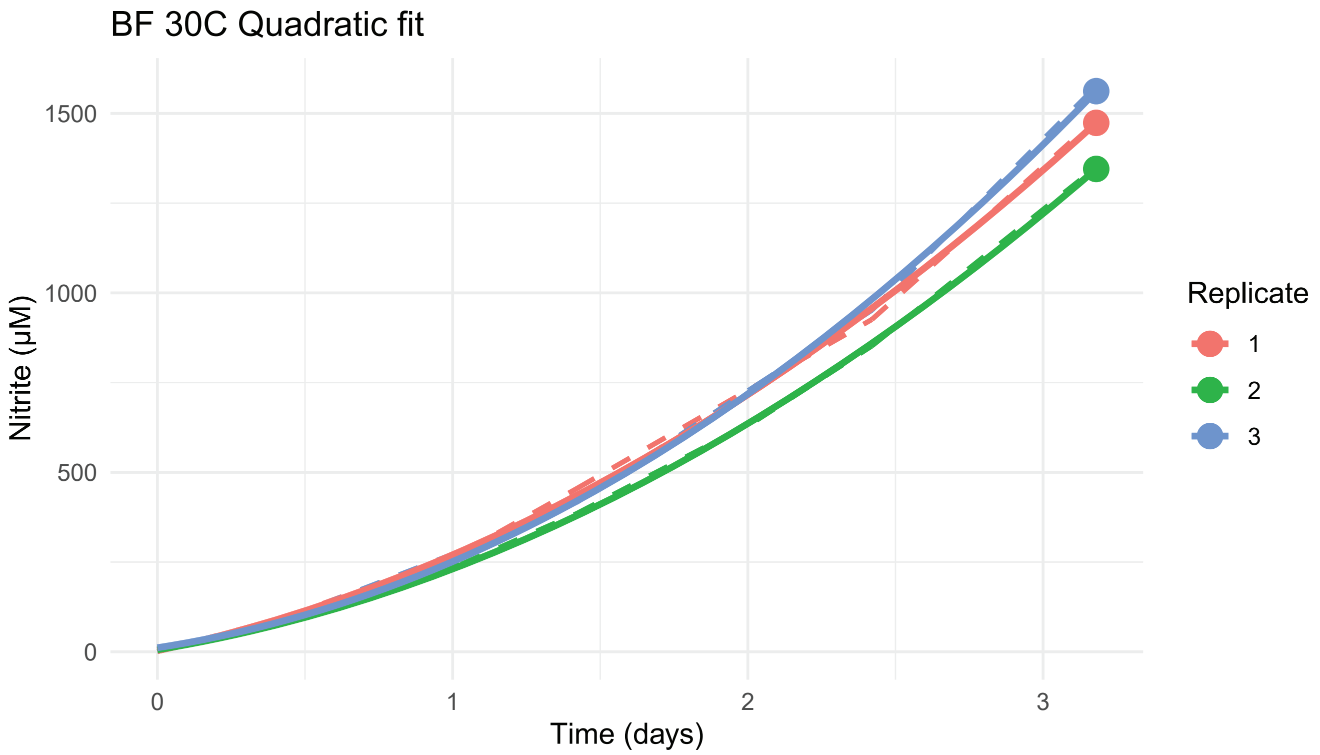

BF 30C Quadratic fit
Nitrite (µM)
1500
1000
500
0
0
1
2
3
Time (days)
Replicate
1
2
3


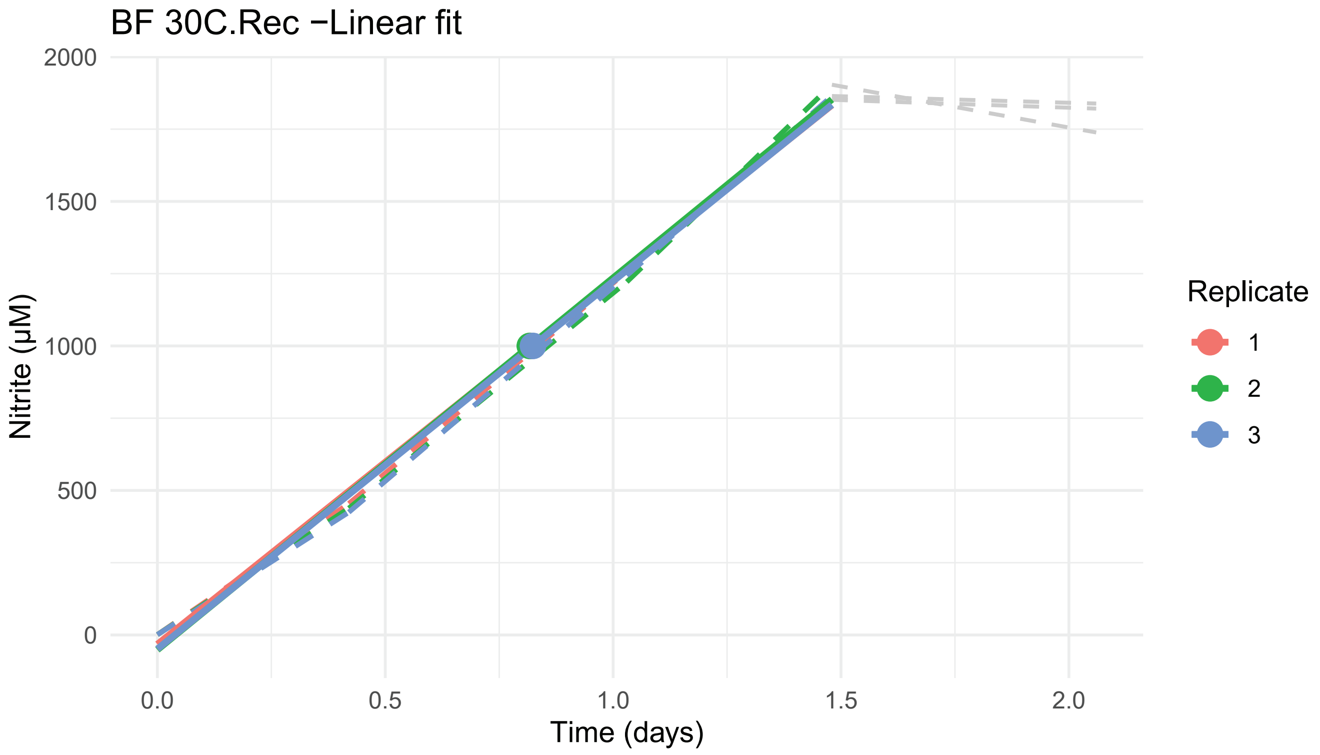

BF 30C.Rec −Linear fit
Nitrite (µM)
2000
1500
1000
500
0
0.0
0.5
1.0
1.5
2.0
Time (days)
Replicate
1
2
3

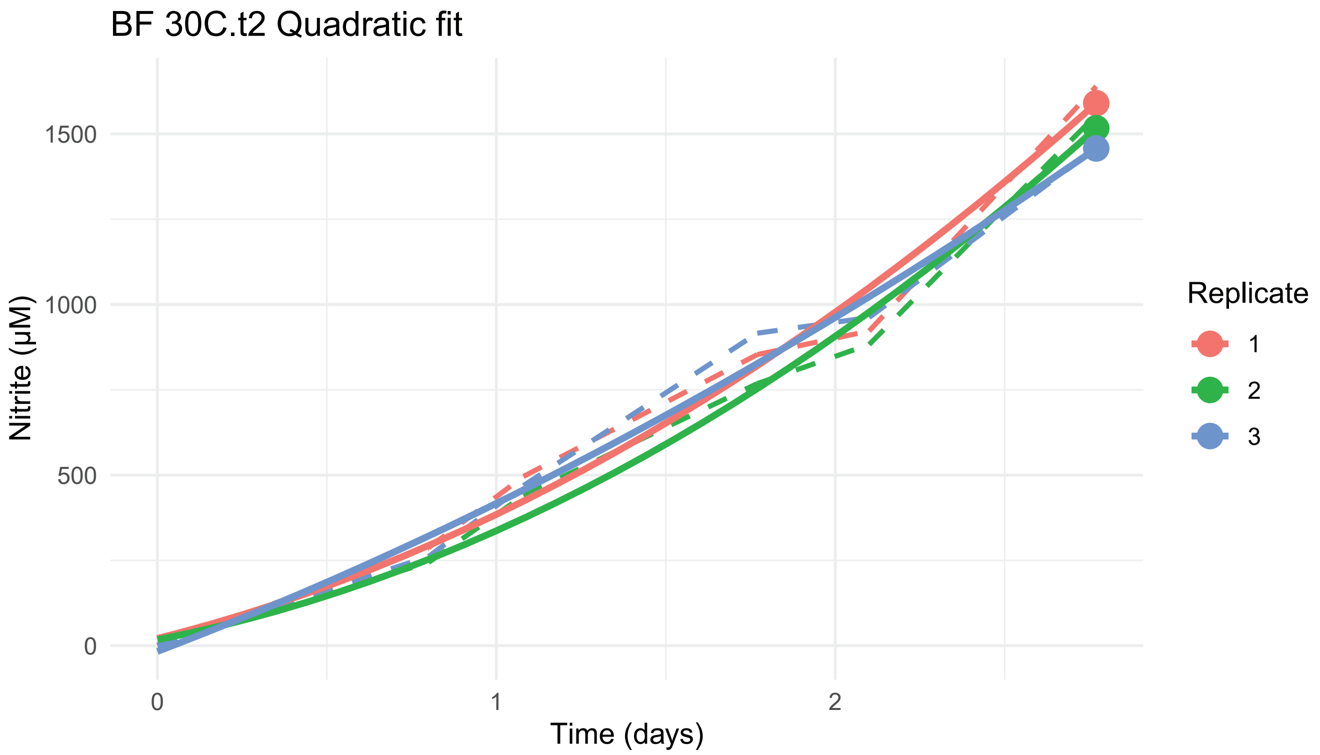

BF 30C.t2 Quadratic fit
Nitrite (µM)
0
500
1000
1500
Time (days)
0
1
2
Replicate
1
2
3


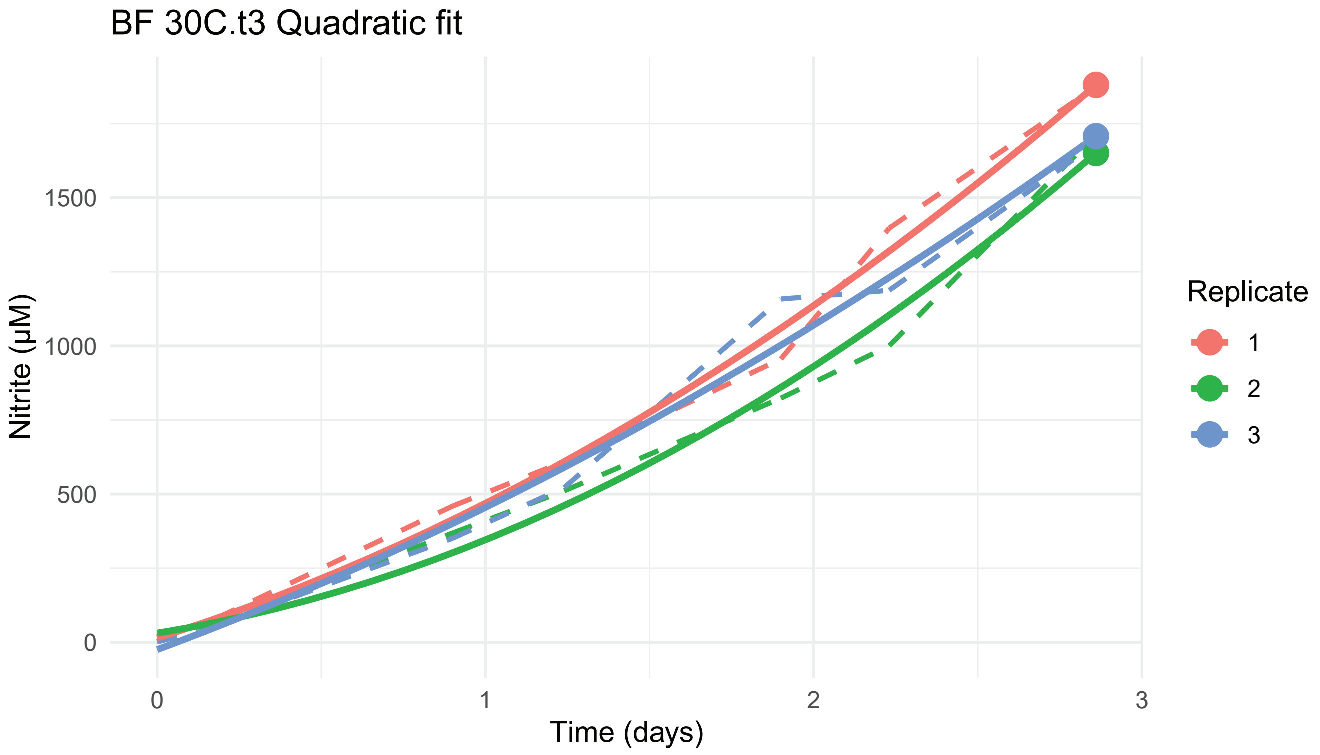

BF 30C.t3 Quadratic fit
Nitrite (µM)
0
500
1000
1500
Time (days)
0
1
2
3
Replicate
1
2
3

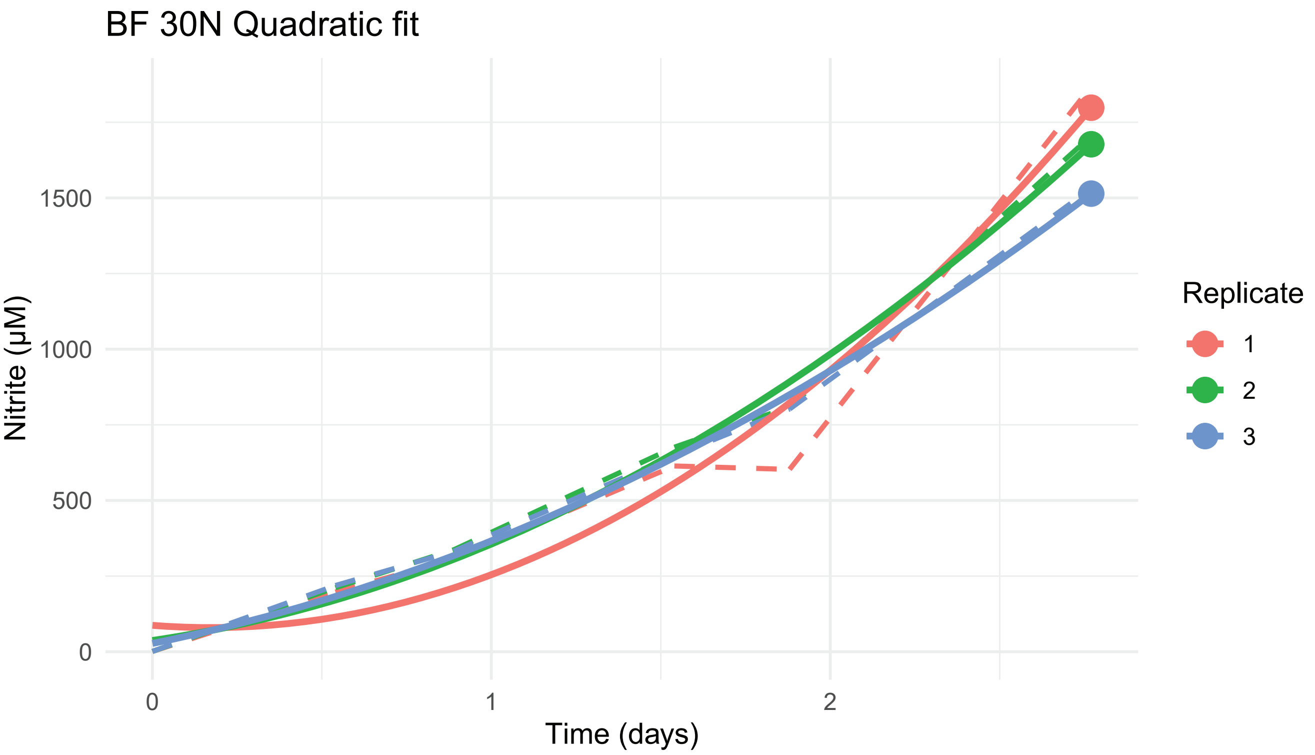
BF 30N Quadratic fit
Nitrite (µM)
0
500
1000
1500
0
1
2
Time (days)
Replicate
1
2
3

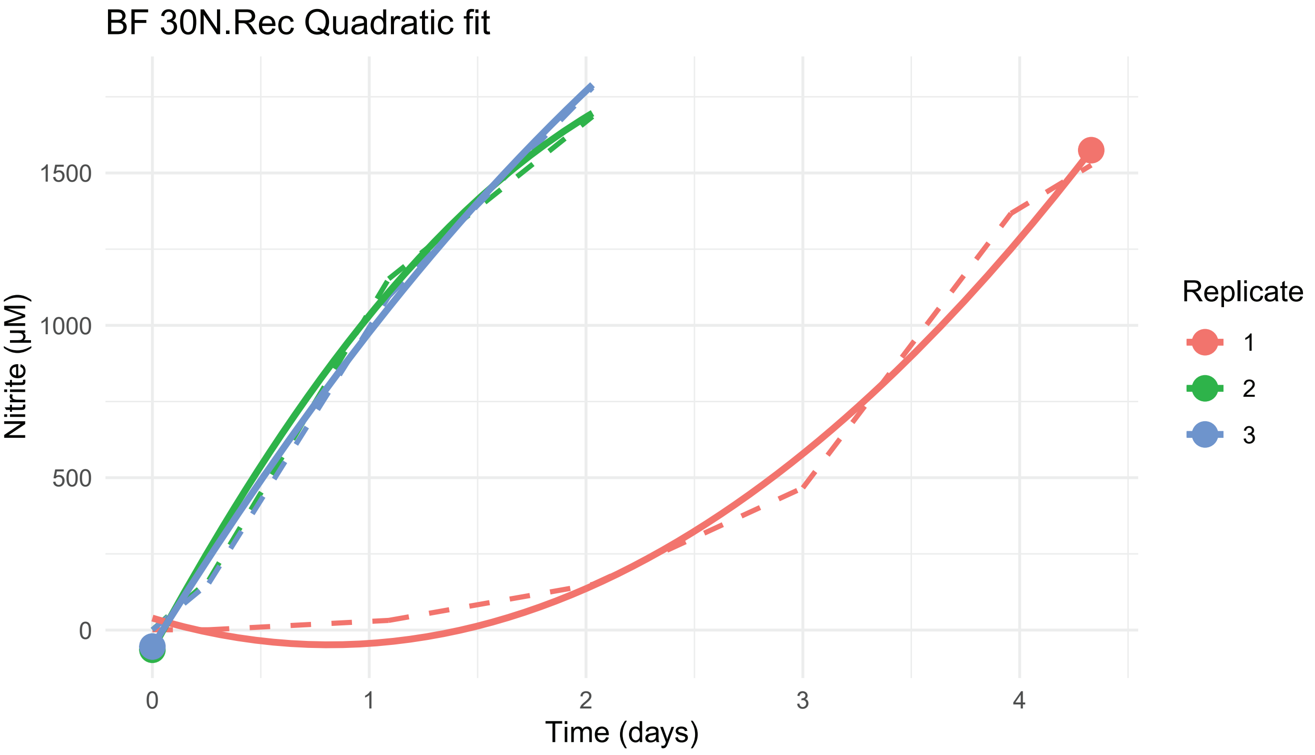
BF 30N.Rec Quadratic fit
Nitrite (µM)
0
500
1000
1500
0
1
2
3
4
Time (days)
Replicate
1
2
3

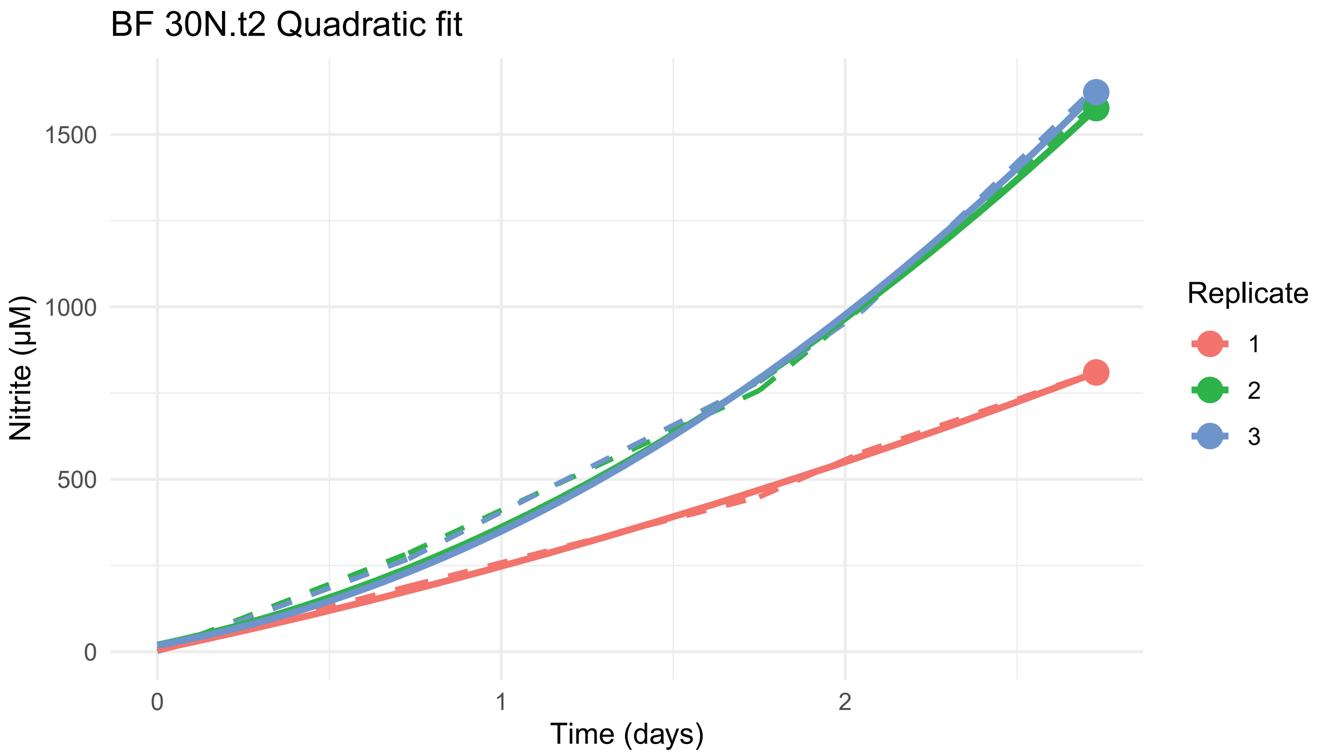
BF 30N.t2 Quadratic fit
Nitrite (µM)
0
500
1000
1500
0
1
2
Time (days)
Replicate
1
2
3

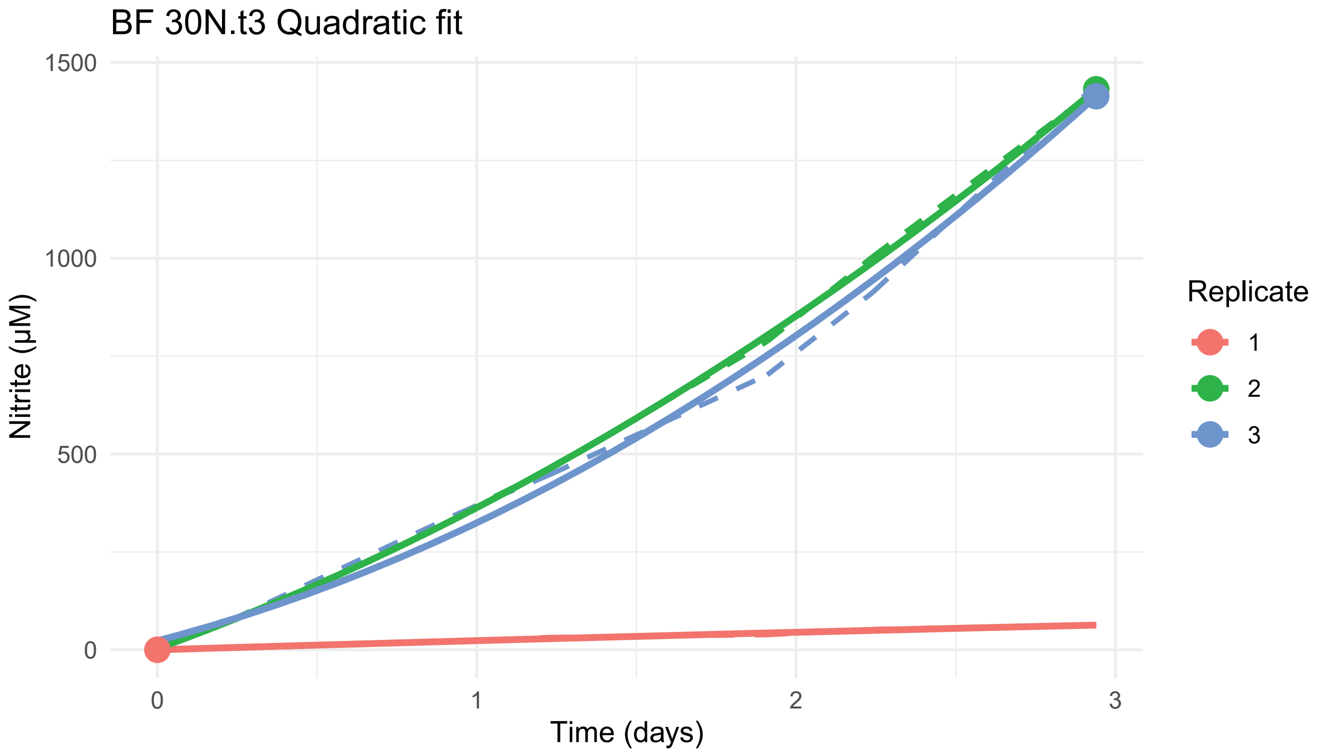
BF 30N.t3 Quadratic fit
Nitrite (µM)
0
500
1000
1500
0
1
2
3
Time (days)
Replicate
1
2
3

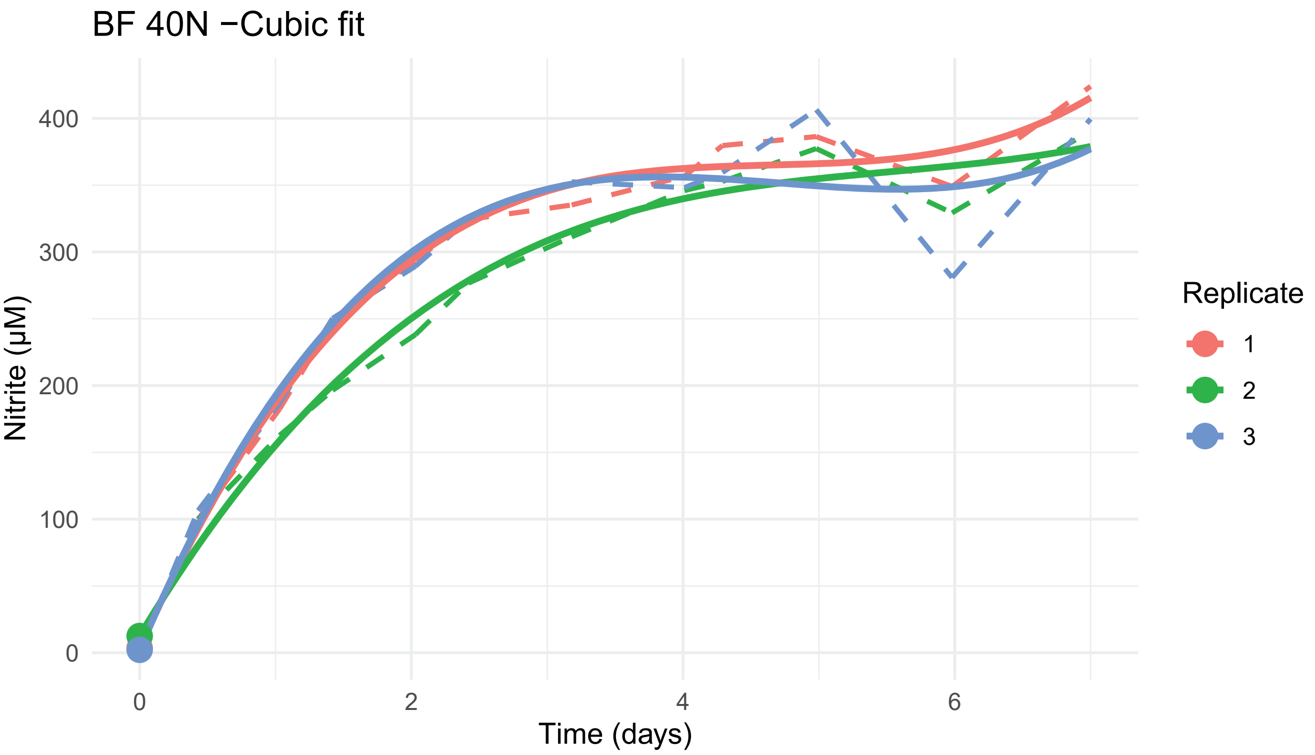
BF 40N −Cubic fit
Nitrite (μM)
0
100
200
300
400
0
2
4
6
Time (days)
Replicate
1
2
3

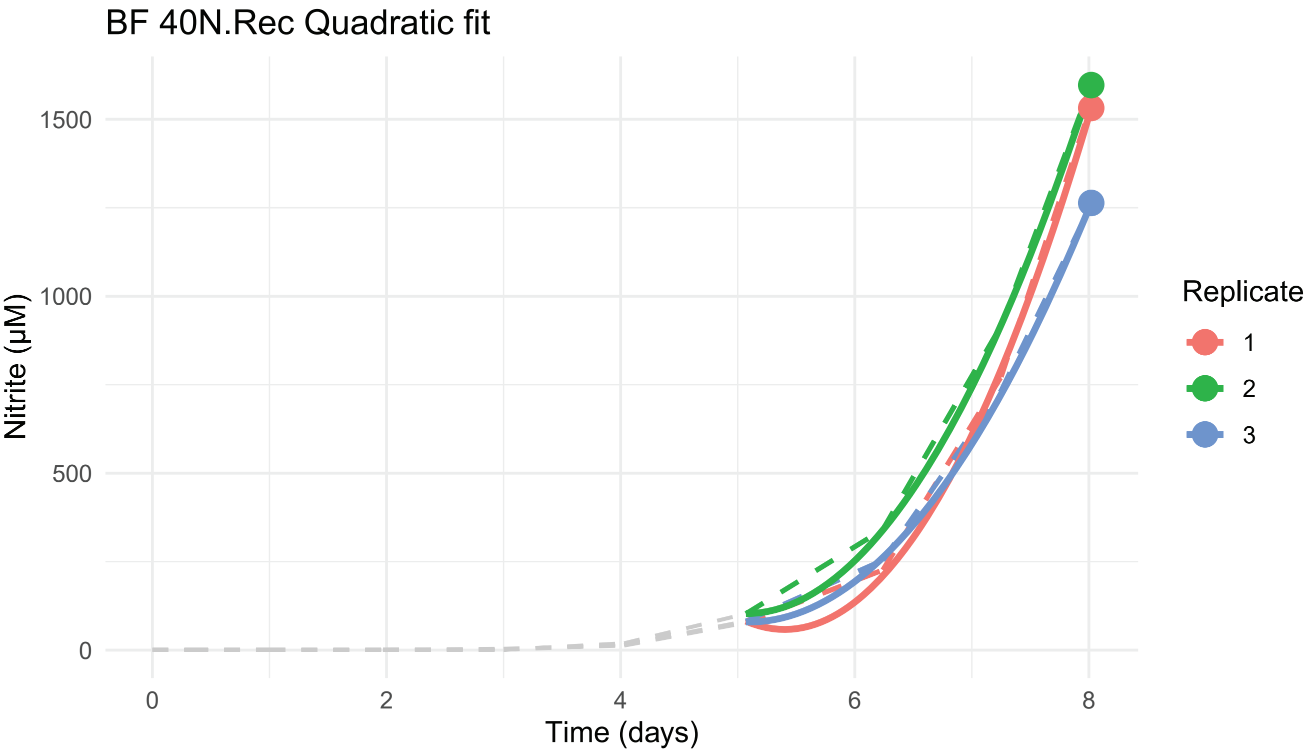
BF 40N.Rec Quadratic fit
Nitrite (μM)
0
500
1000
1500
0
2
4
6
8
Time (days)
Replicate
1
2
3

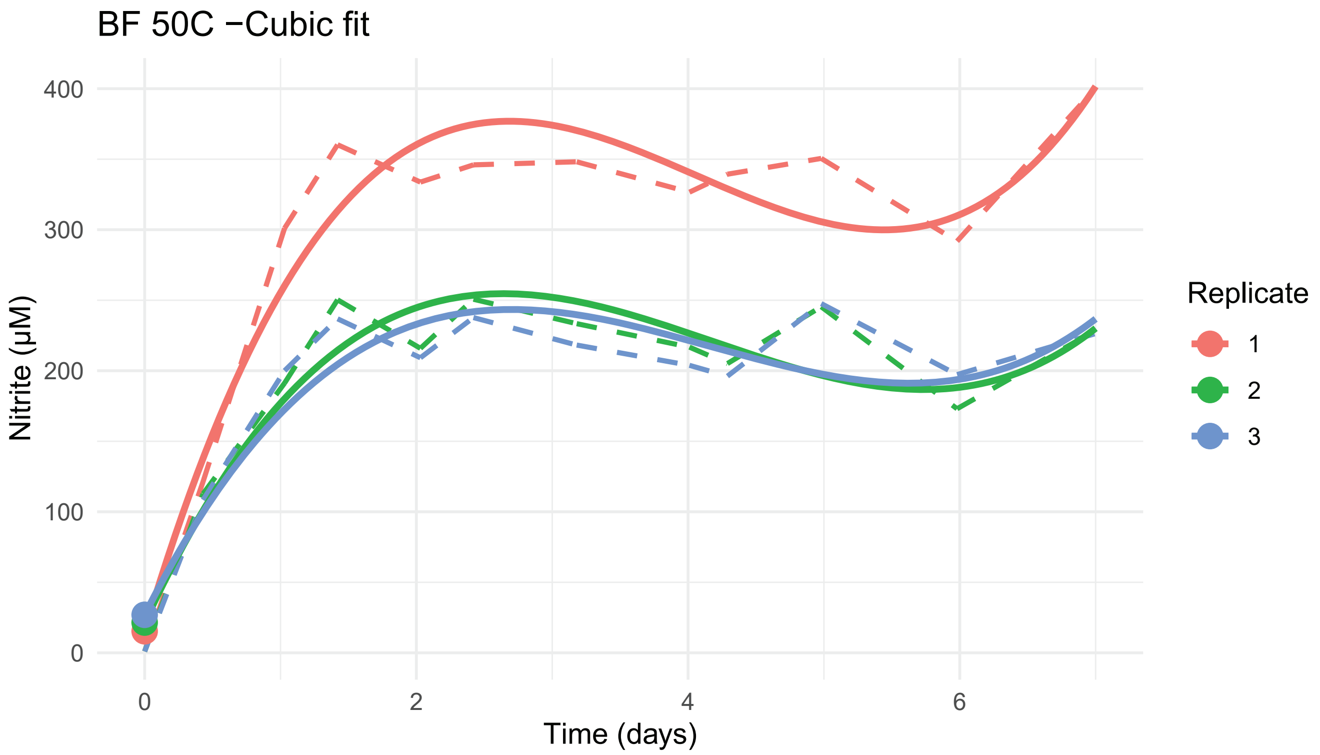
BF 50C −Cubic fit
Nitrite (µM)
400
300
200
100
0
0
2
4
6
Time (days)
Replicate
1
2
3

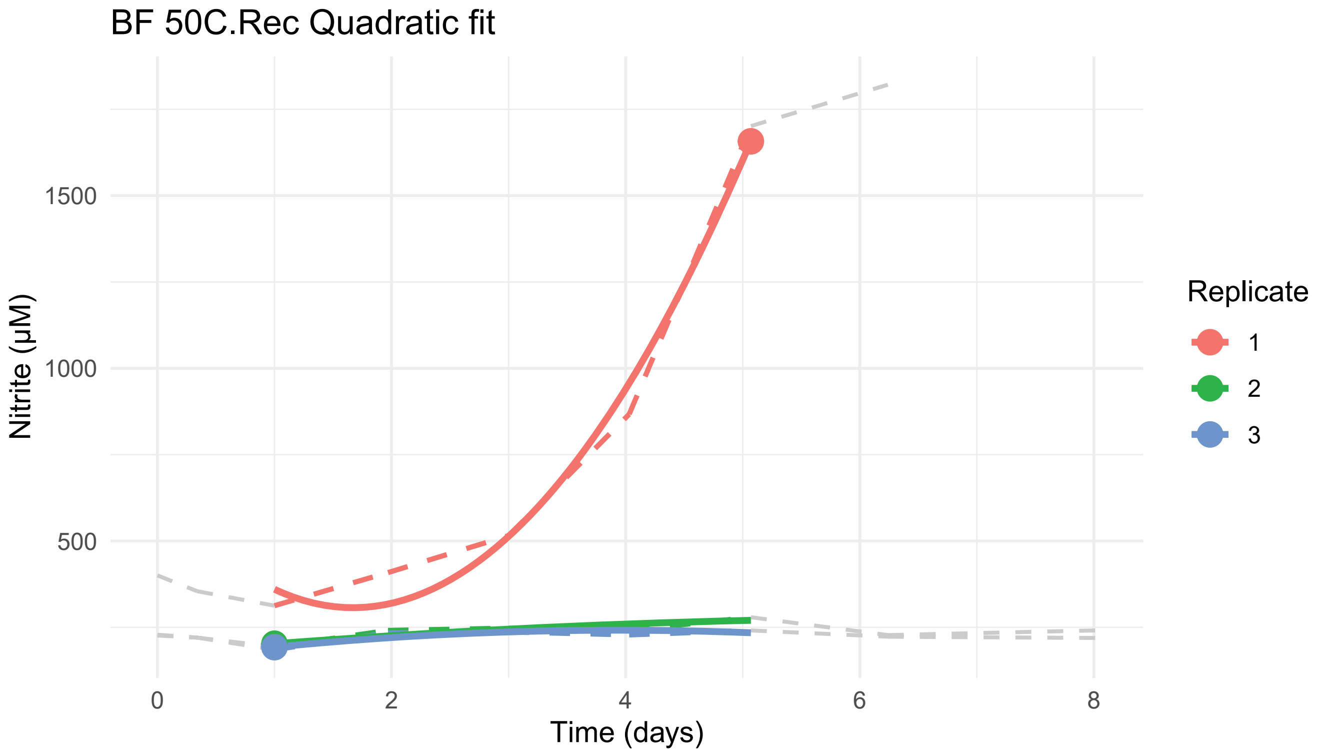
BF 50C.Rec Quadratic fit
Nitrite (µM)
1500
1000
500
0
2
4
6
8
Time (days)
Replicate
1
2
3

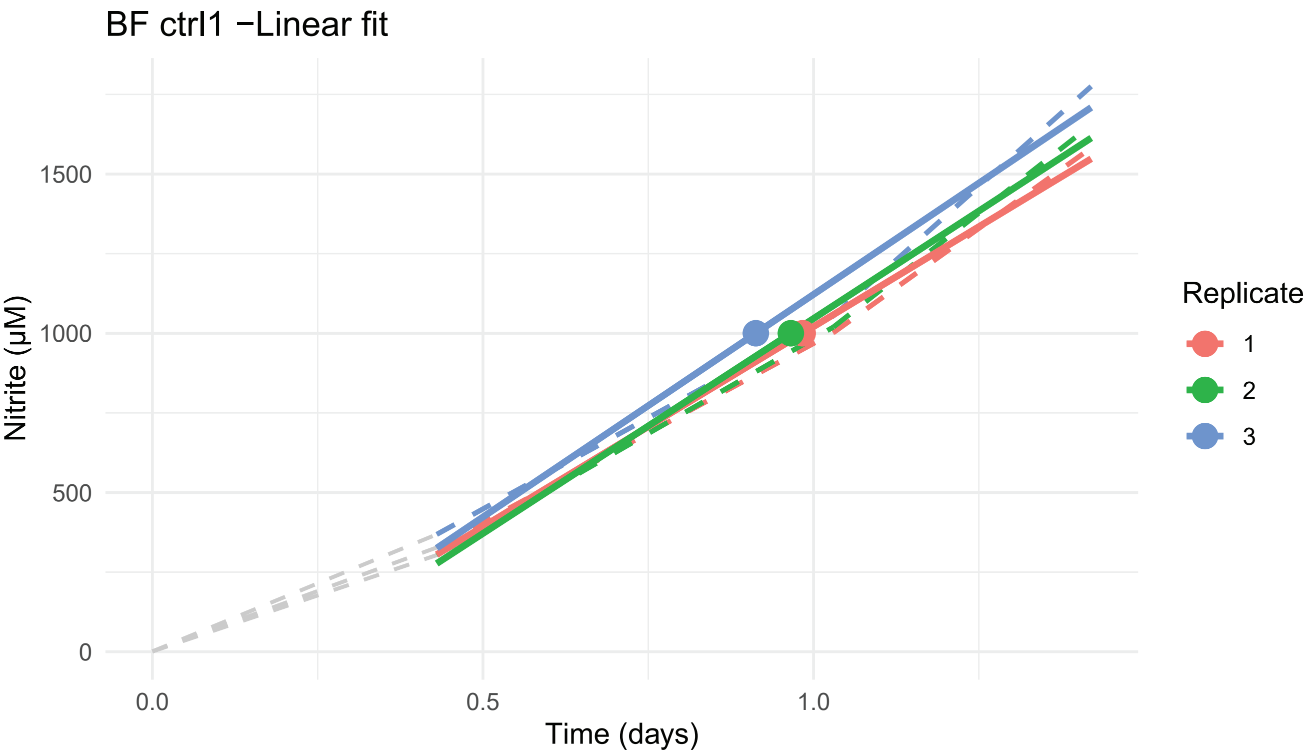
BF ctrl1 −Linear fit
Nitrite (µM)
0
500
1000
1500
Time (days)
0.0
0.5
1.0
Replicate
1
2
3

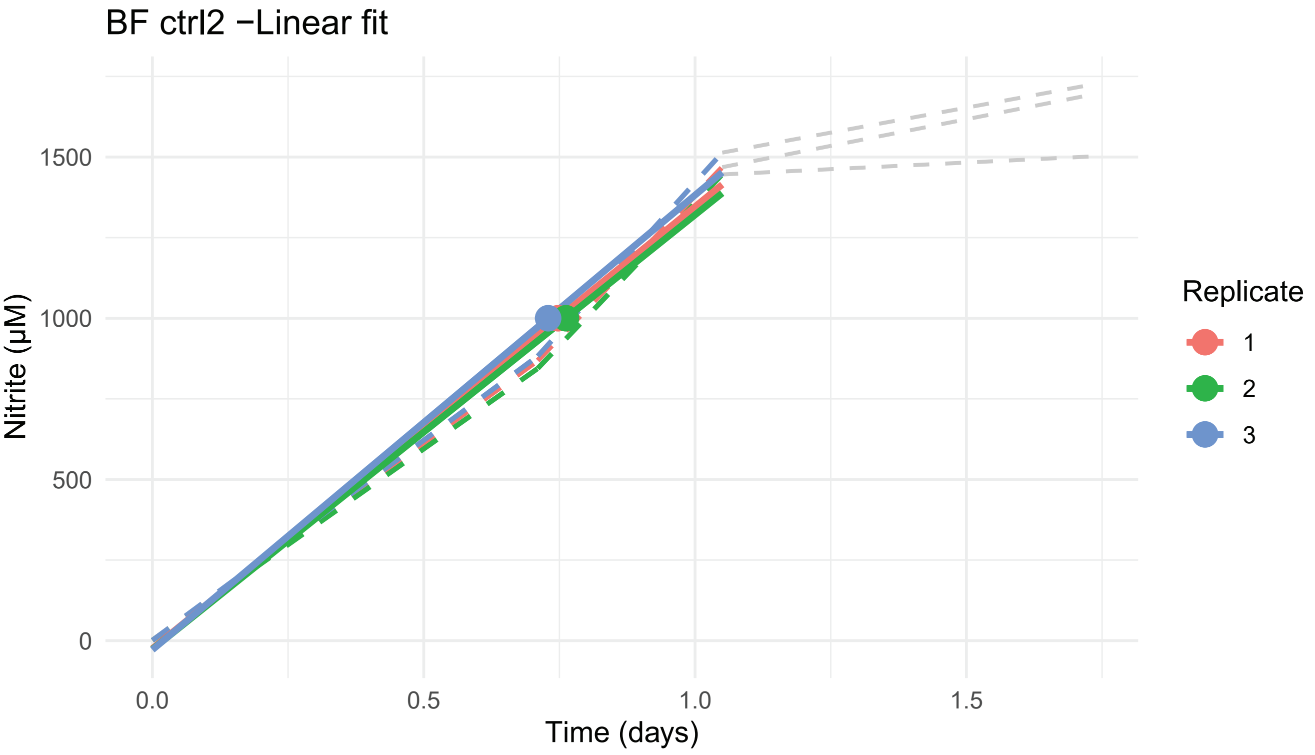
BF ctrl2 −Linear fit
Nitrite (µM)
0
500
1000
1500
Time (days)
0.0
0.5
1.0
1.5
Replicate
1
2
3

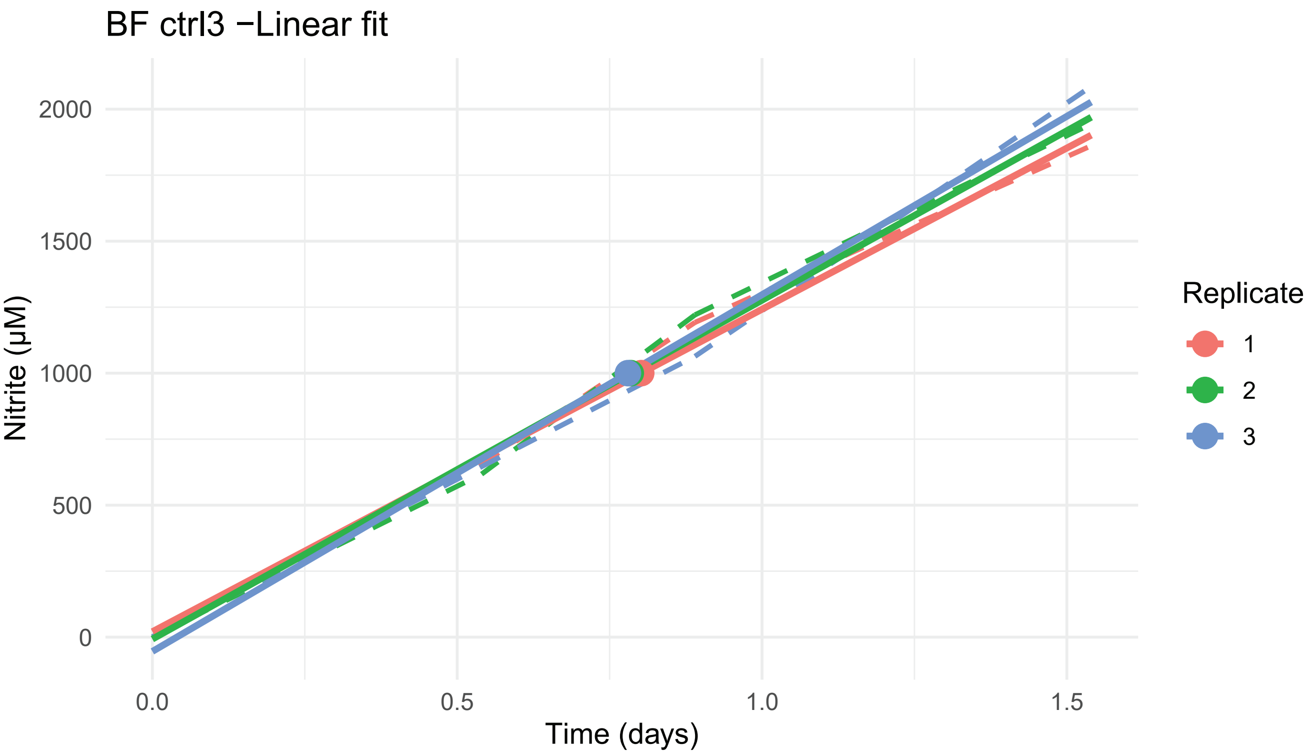
BF ctrl3 −Linear fit
Nitrite (µM)
2000
1500
1000
500
0
0.0
0.5
1.0
1.5
Time (days)
Replicate
1
2
3

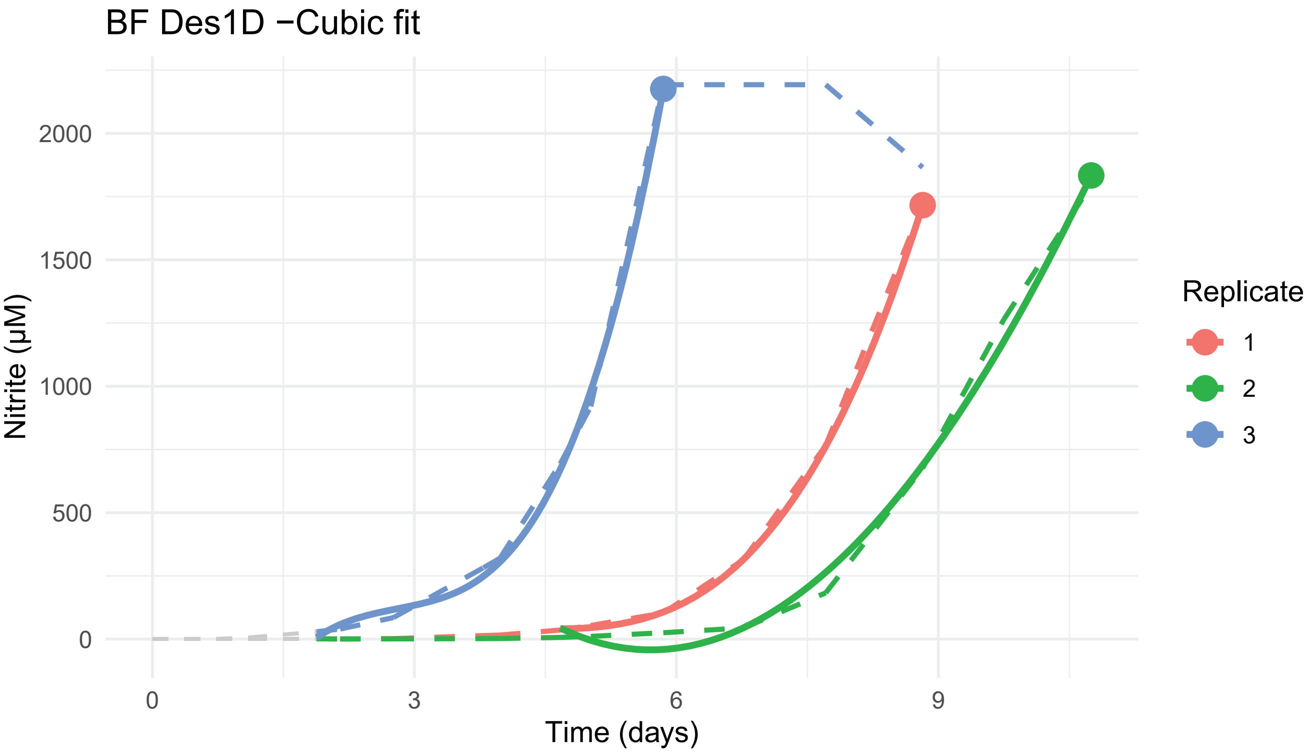
BF Des1D −Cubic fit
Nitrite (µM)
2000
1500
1000
500
0
0
3
6
9
Time (days)
Replicate
1
2
3

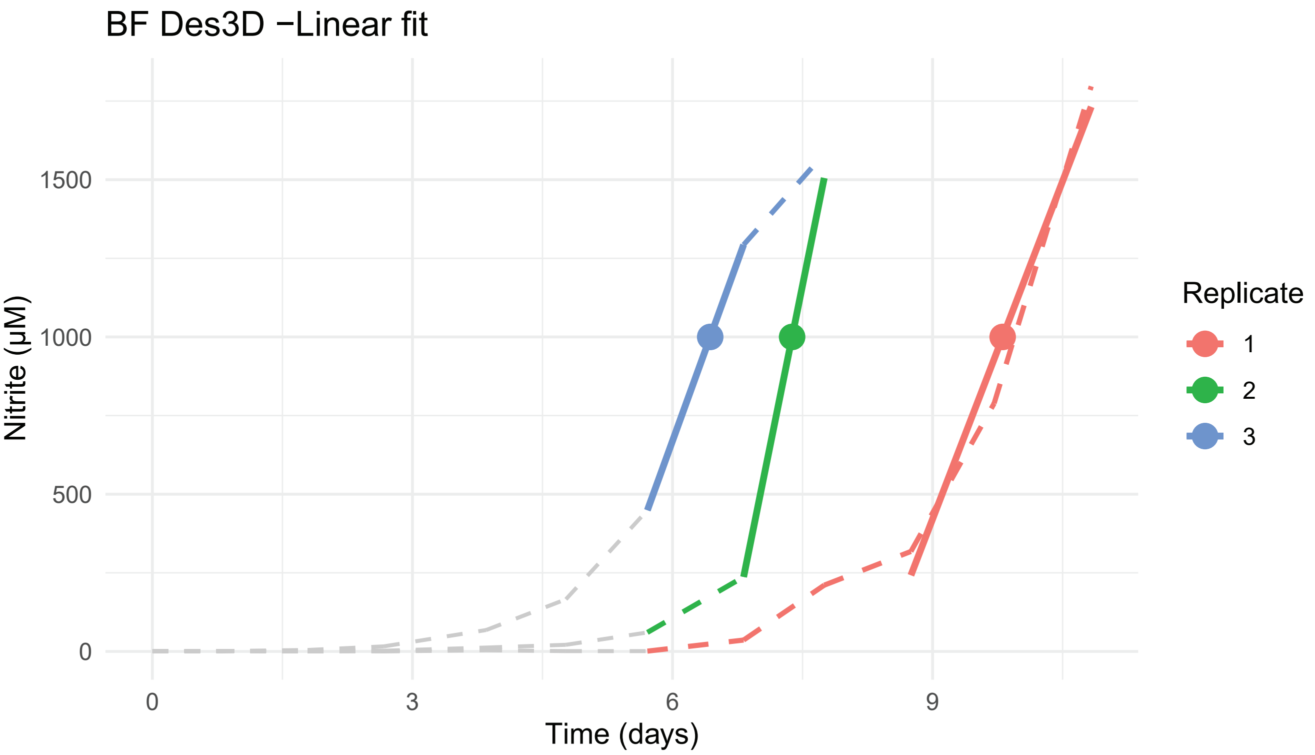

BF Des3D −Linear fit
Nitrite (µM)
0
500
1000
1500
0
3
6
9
Time (days)
Replicate
1
2
3


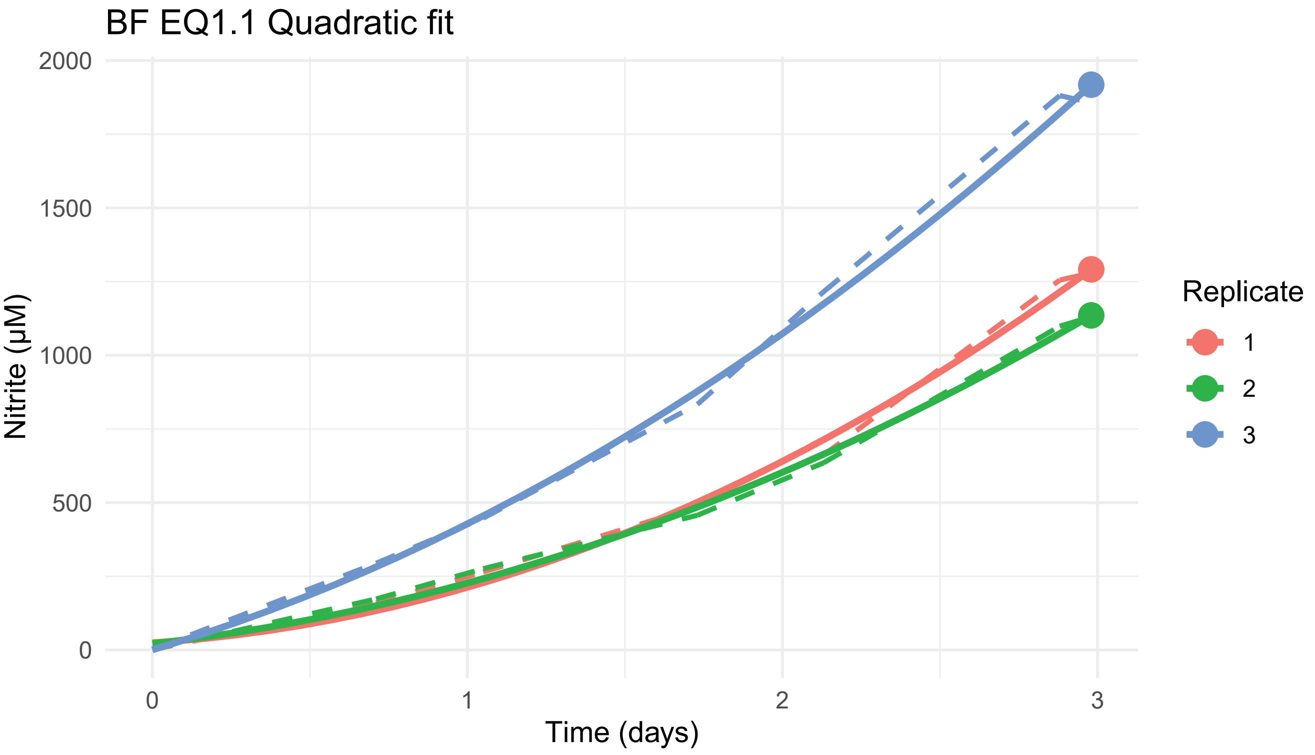

BF EQ1.1 Quadratic fit
Nitrite (µM)
0
500
1000
1500
2000
0
1
2
3
Time (days)
Replicate
1
2
3

BF EQ1.1.Rec Quadratic fit

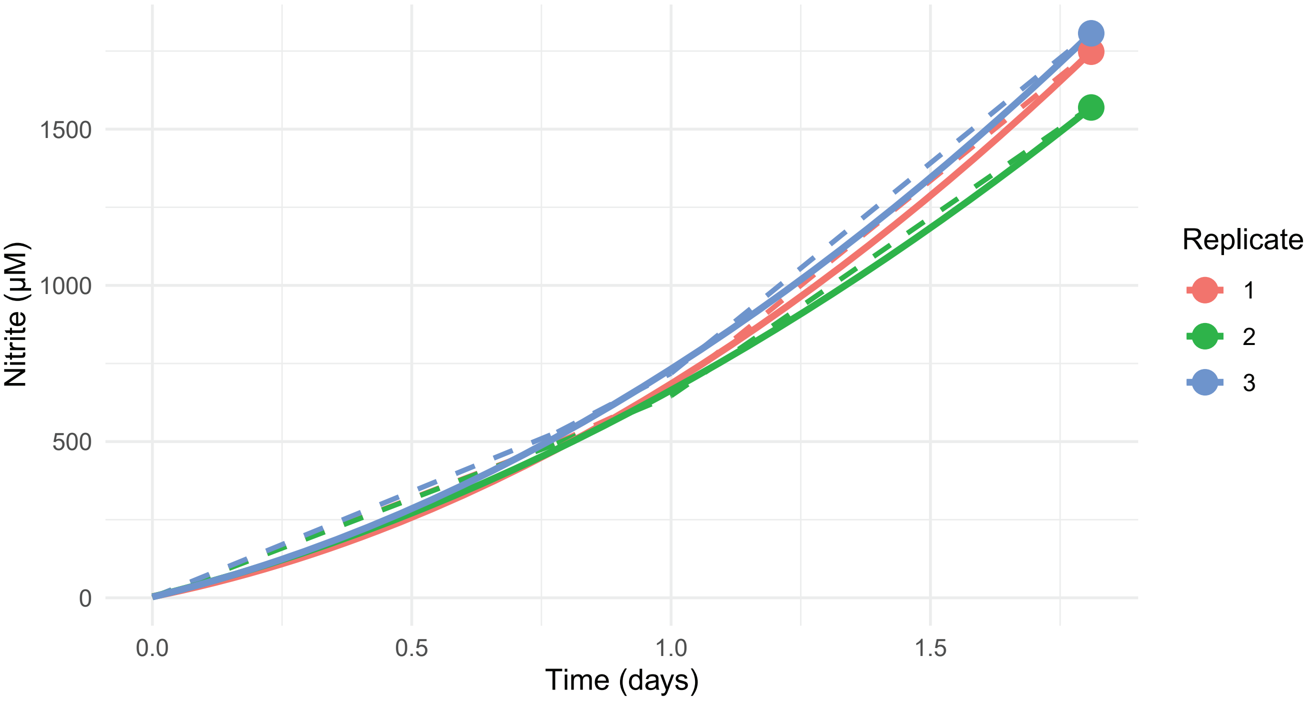


BF EQ1.3 Quadratic fit

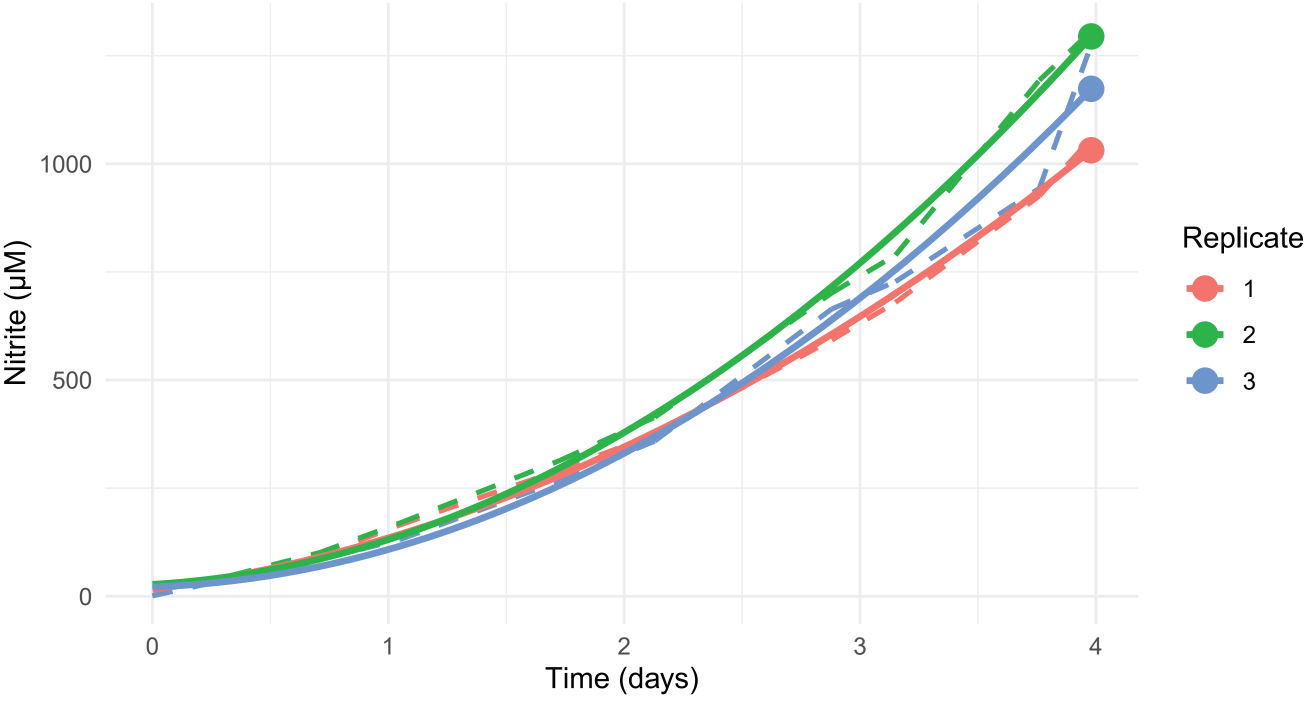

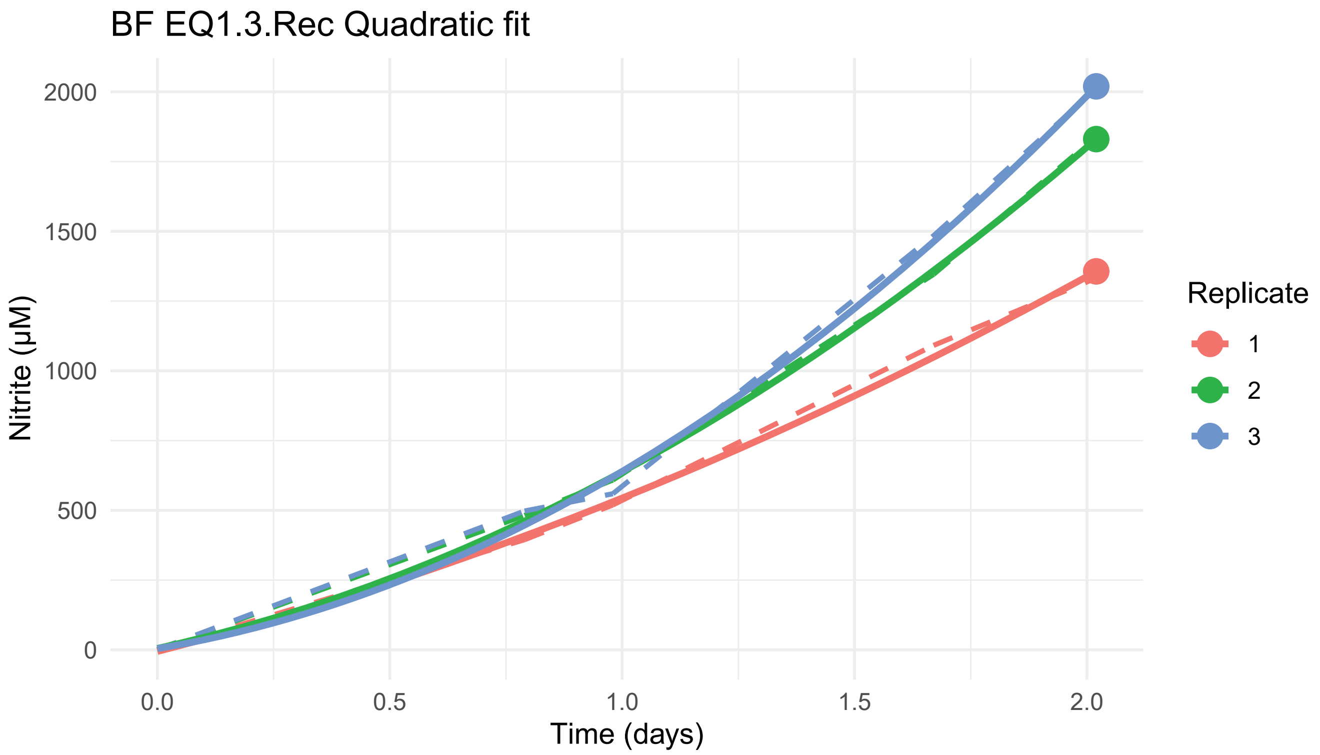
BF EQ1.3.Rec Quadratic fit
Nitrite (μM)
0
500
1000
1500
2000
0.0
0.5
1.0
1.5
2.0
Time (days)
Replicate
1
2
3

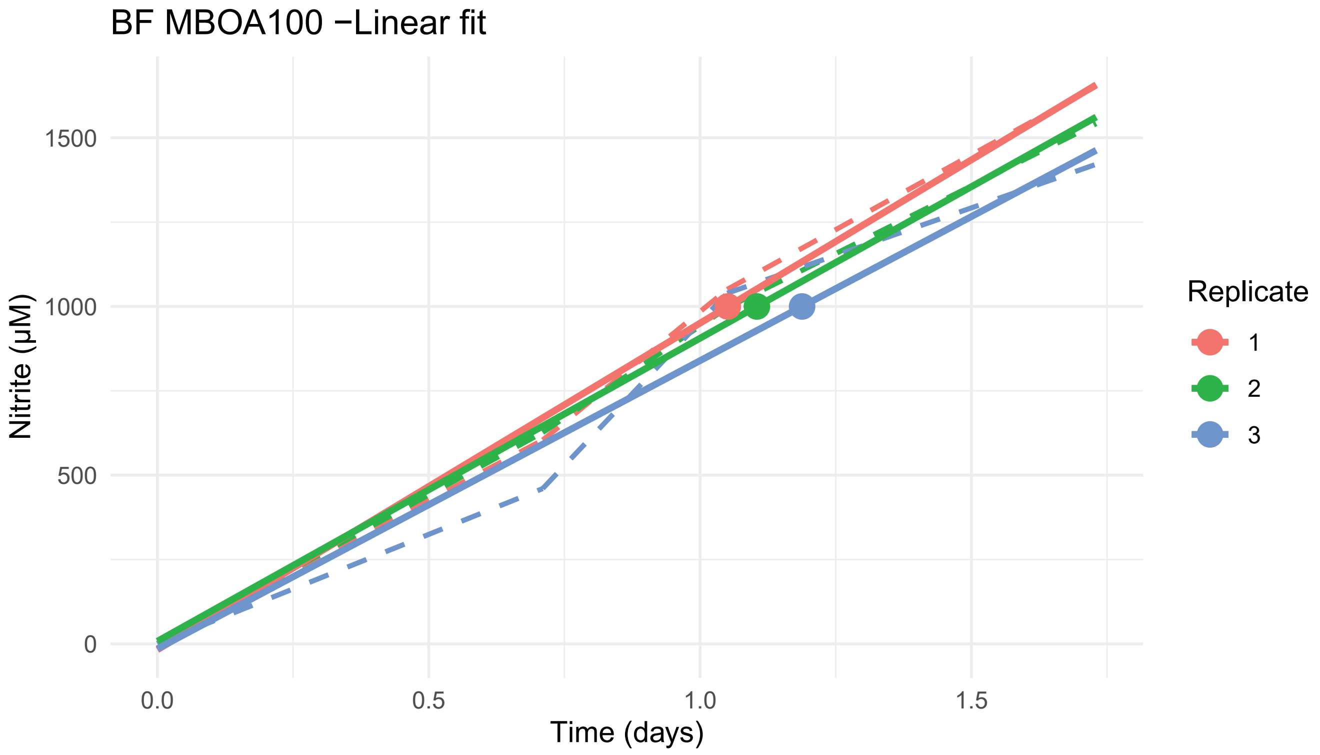
BF MBOA100 −Linear fit
Nitrite (μM)
0
500
1000
1500
0.0
0.5
1.0
1.5
Time (days)
Replicate
1
2
3

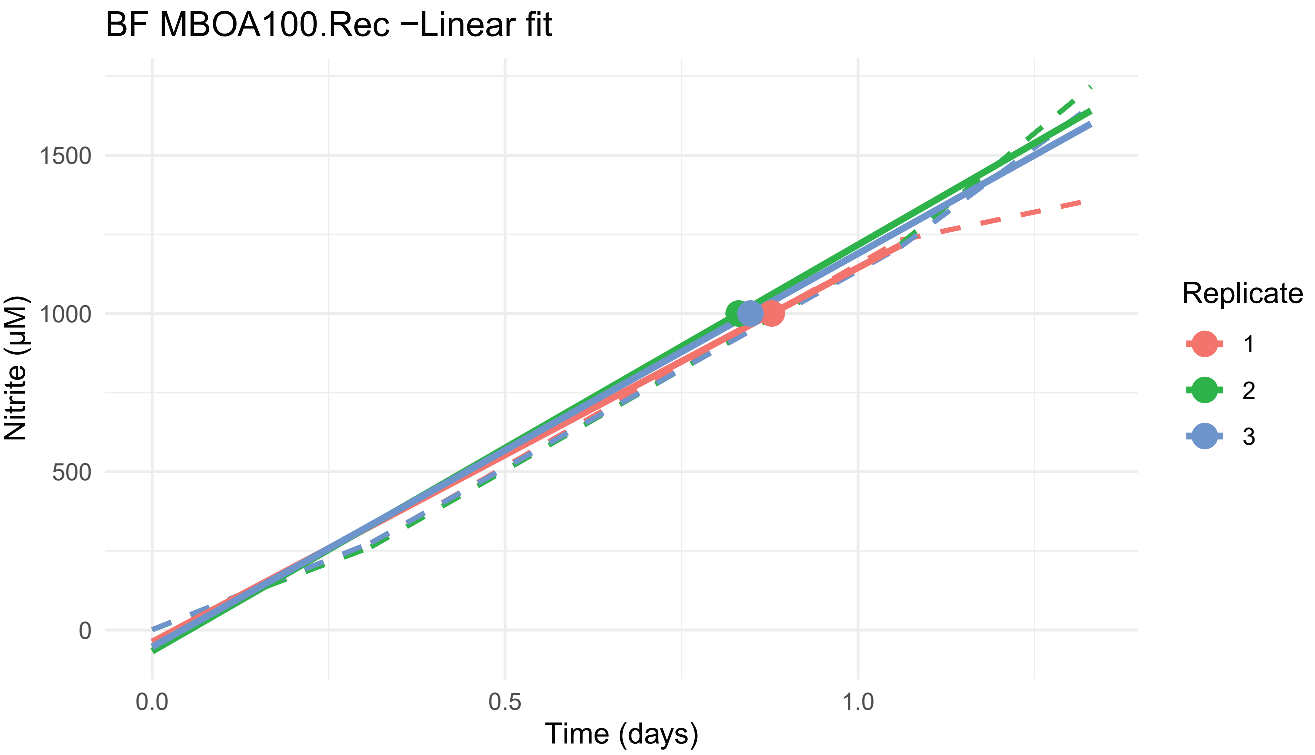
BF MBOA100.Rec −Linear fit
Nitrite (µM)
Time (days)
0
500
1000
1500
0.0
0.5
1.0
Replicate
1
2
3

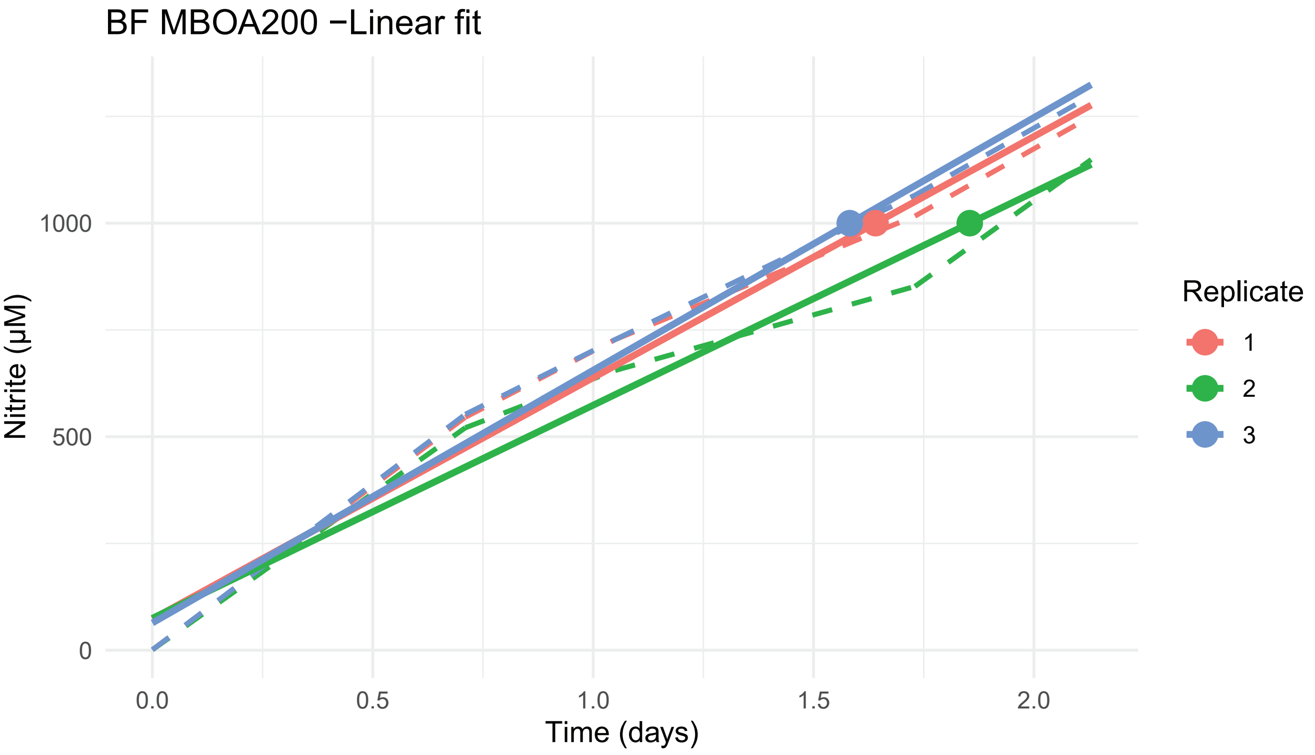
BF MBOA200 −Linear fit
Nitrite (µM)
Time (days)
0
500
1000
0.0
0.5
1.0
1.5
2.0
Replicate
1
2
3

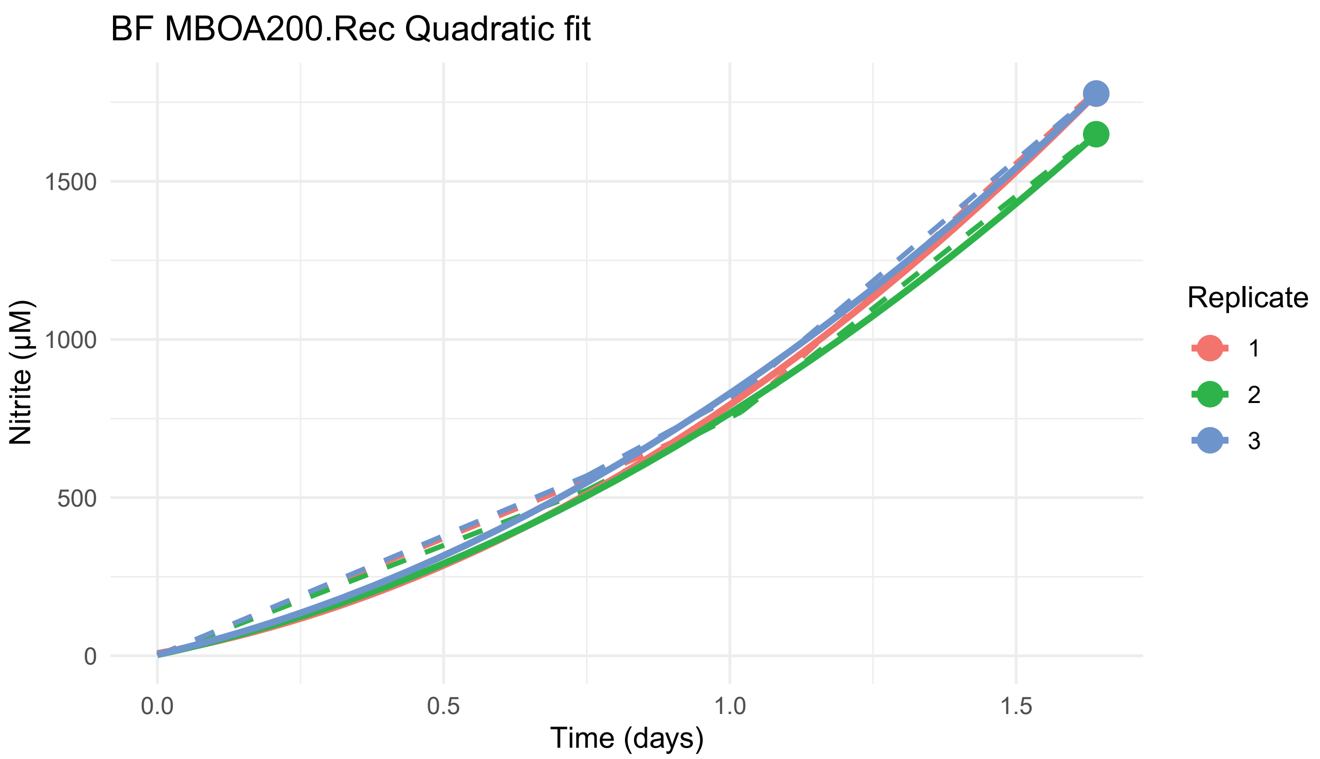
BF MBOA200.Rec Quadratic fit
Nitrite (µM)
1500
1000
500
0
0.0
0.5
1.0
1.5
Time (days)
Replicate
1
2
3

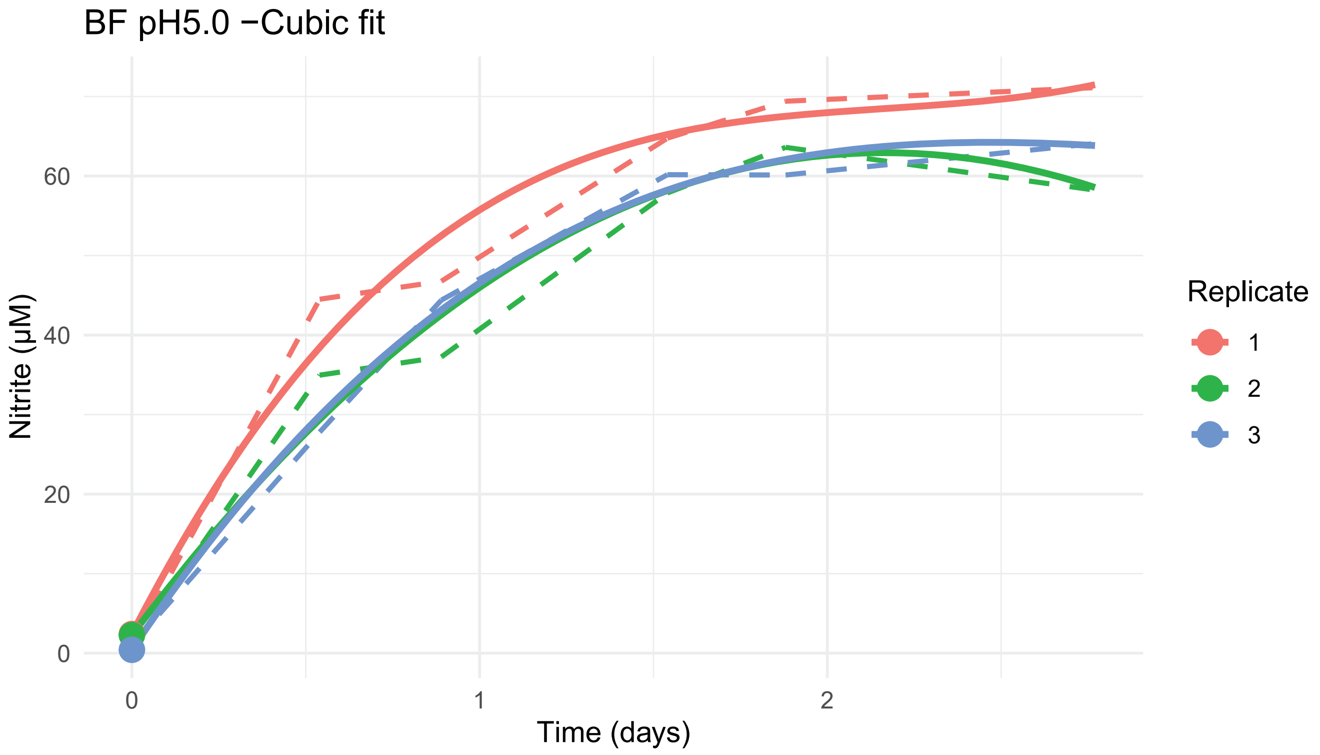
BF pH5.0 −Cubic fit
Nitrite (µM)
60
40
20
0
0
1
2
Time (days)
Replicate
1
2
3

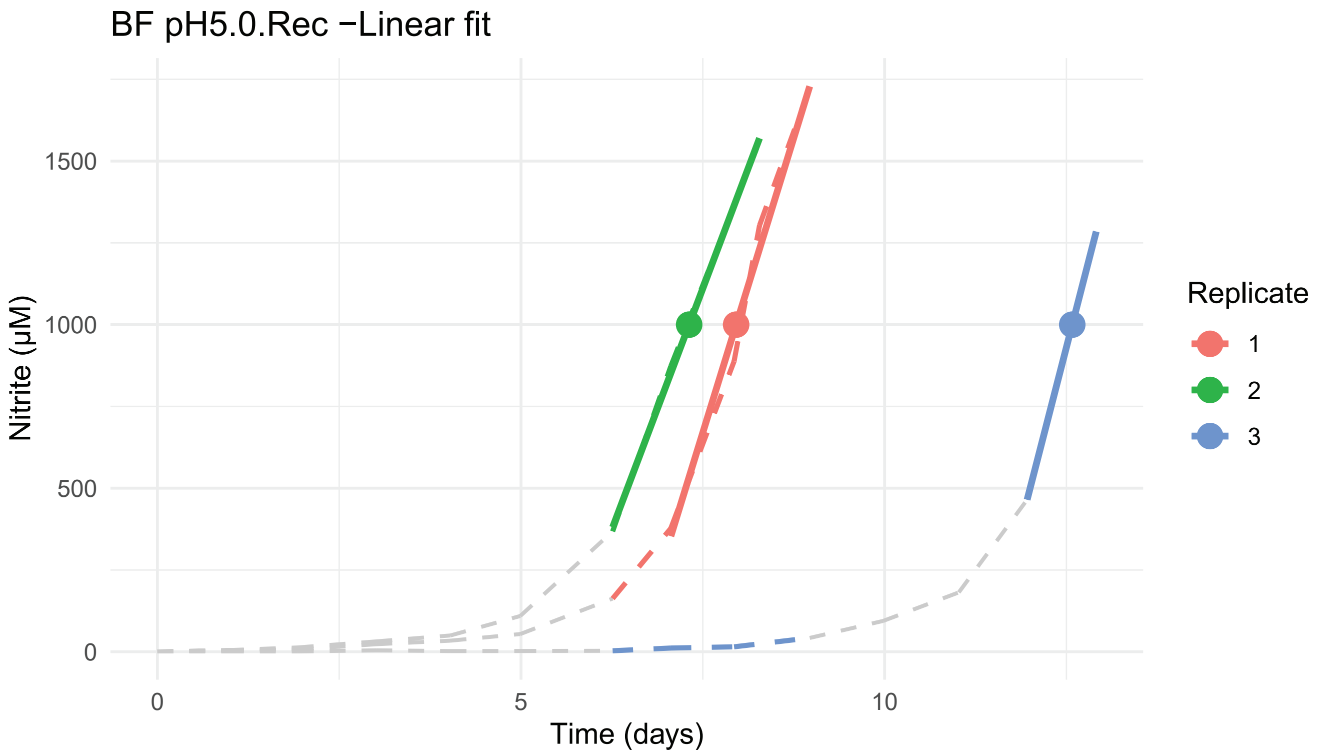
BF pH5.0.Rec −Linear fit
Nitrite (µM)
0
500
1000
1500
0
5
10
Time (days)
Replicate
1
2
3

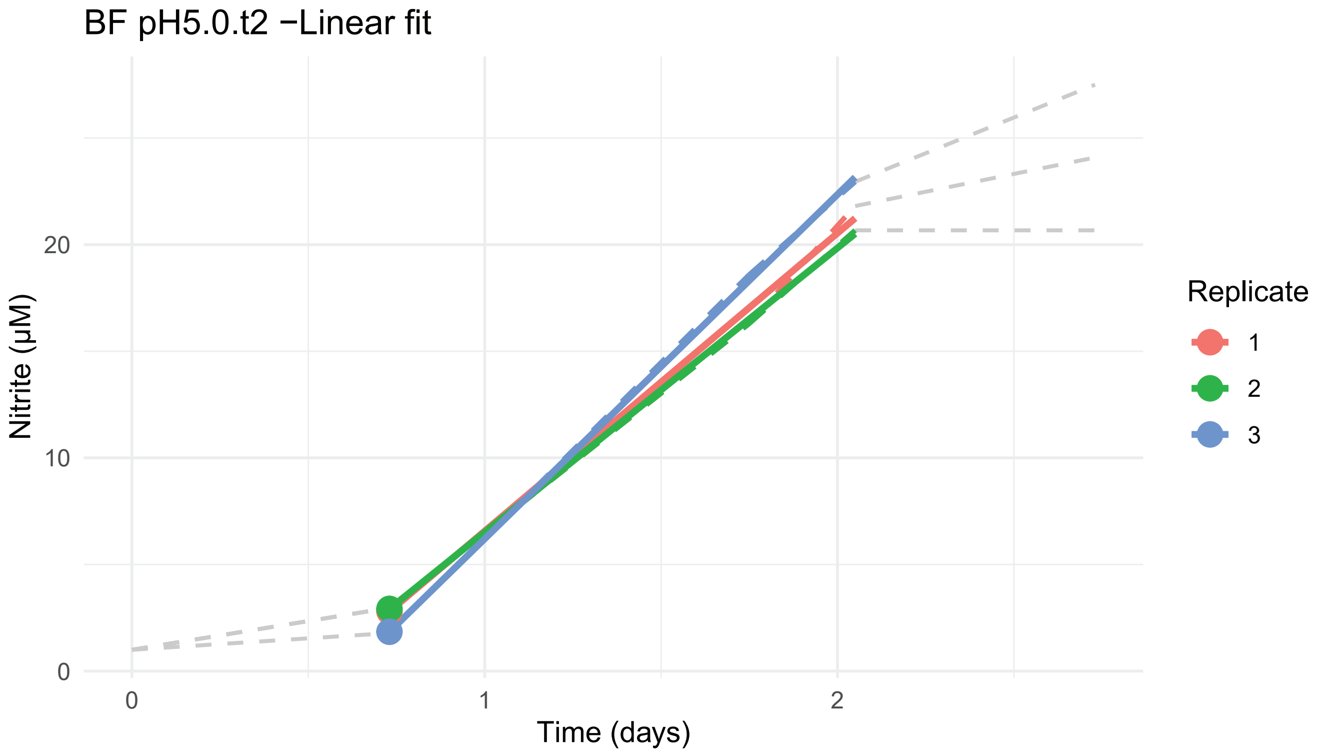
BF pH5.0.t2 −Linear fit
Nitrite (µM)
0
10
20
0
1
2
Time (days)
Replicate
1
2
3

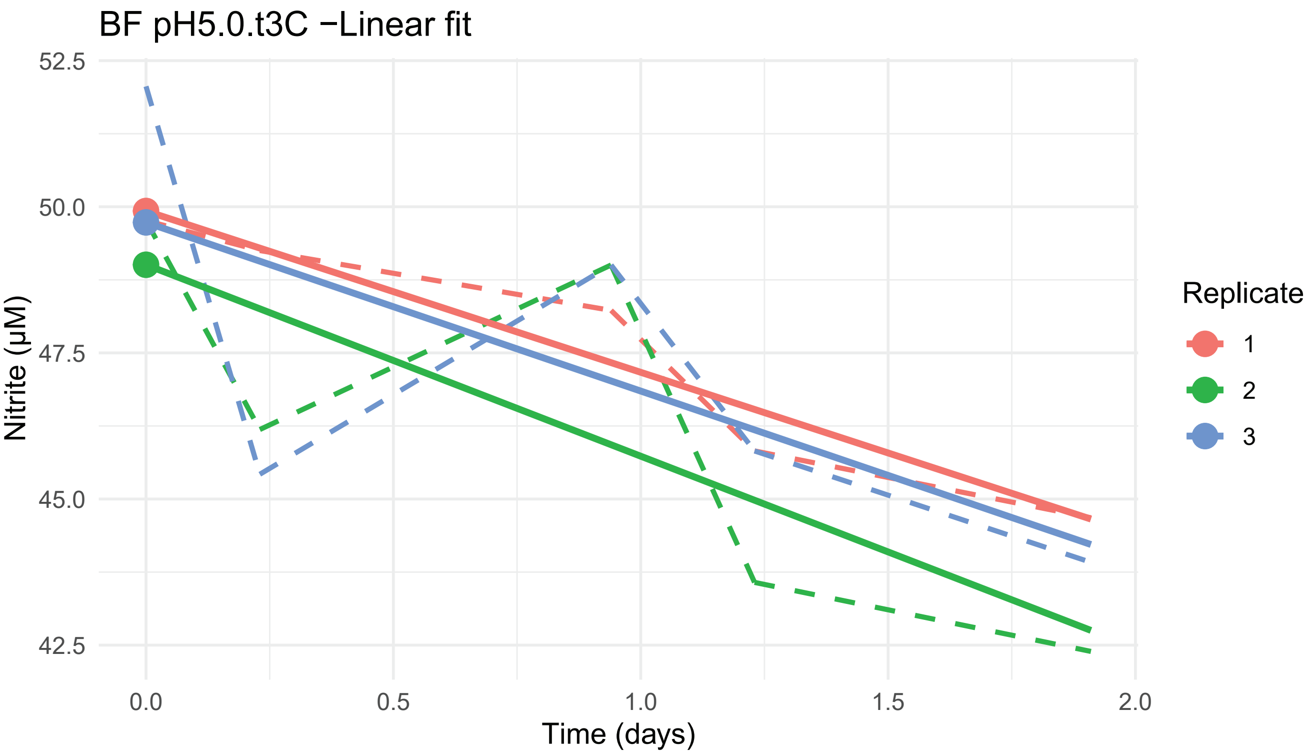
BF pH5.0.t3C −Linear fit
52.5
50.0
47.5
45.0
42.5
Nitrite (µM)
0.0
0.5
1.0
1.5
2.0
Time (days)
Replicate
1
2
3

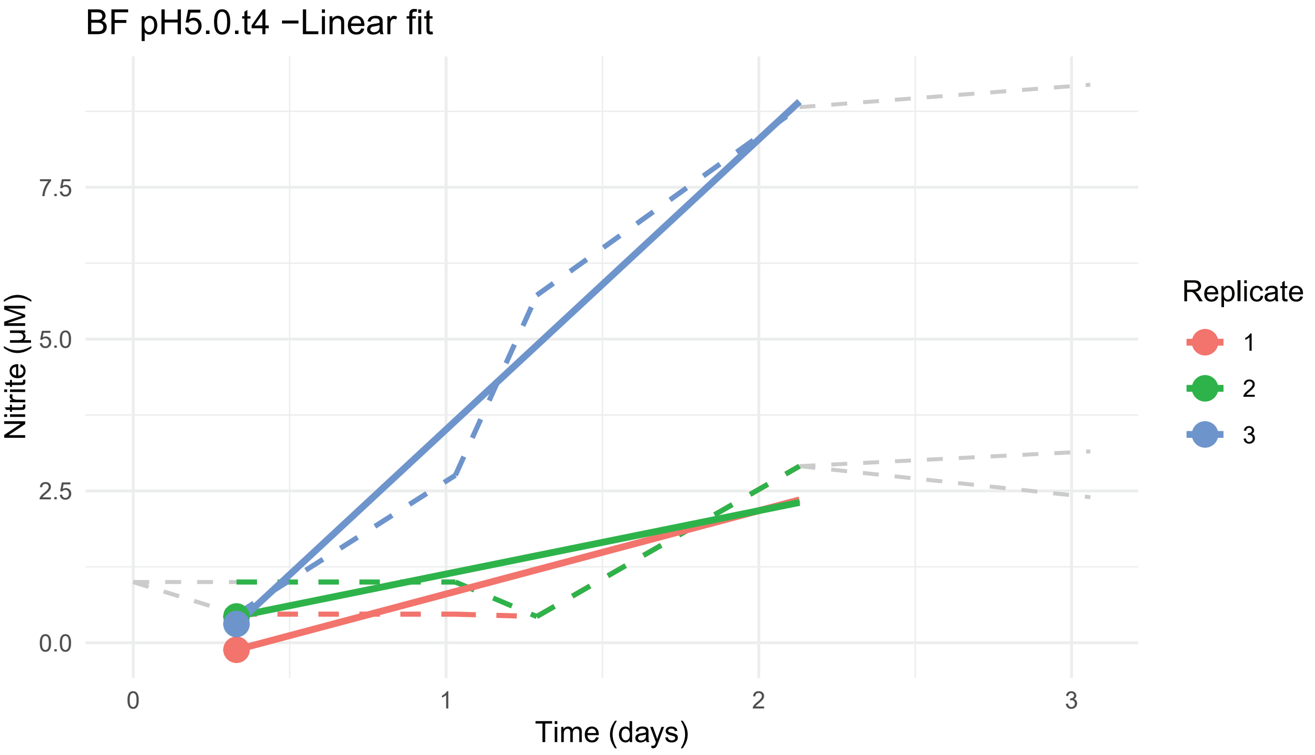
BF pH5.0.t4 −Linear fit
7.5
5.0
2.5
0.0
Nitrite (µM)
0
1
2
3
Time (days)
Replicate
1
2
3

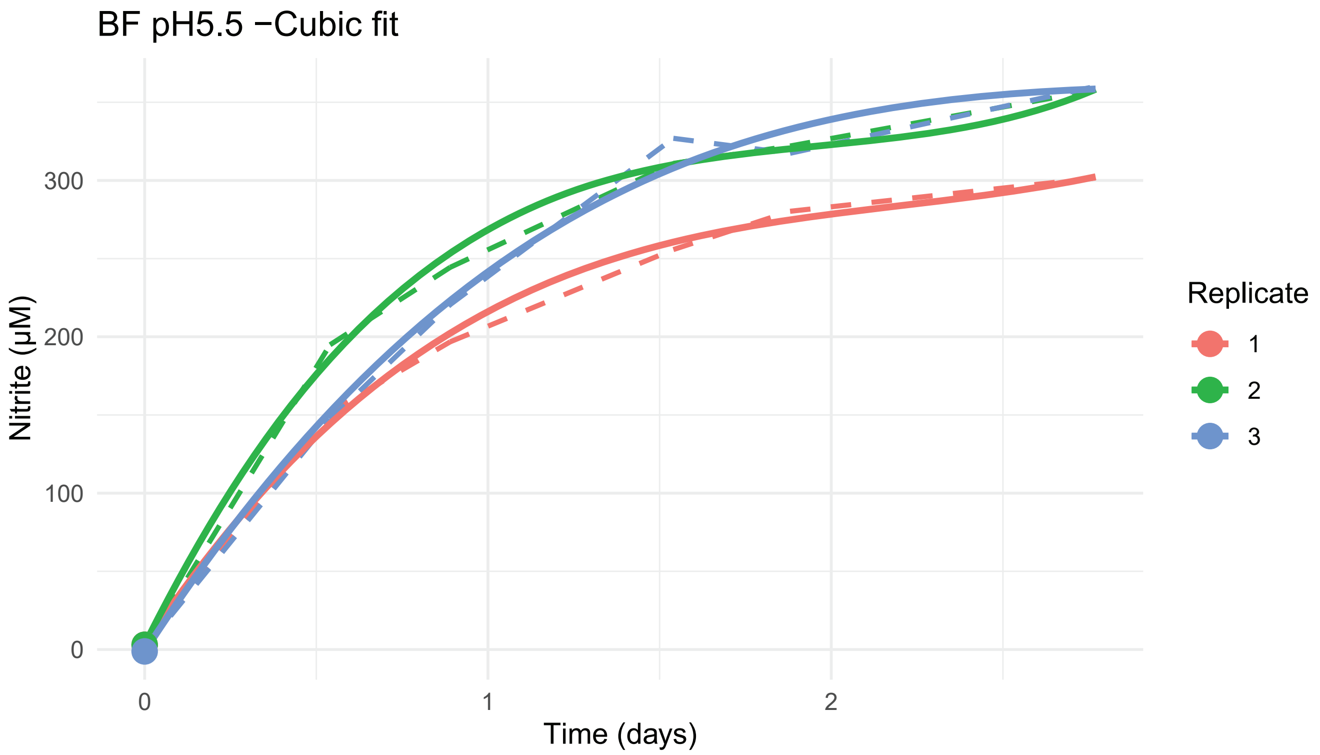

BF pH5.5 −Cubic fit
Nitrite (µM)
300
200
100
0
0
1
2
Time (days)
Replicate
1
2
3


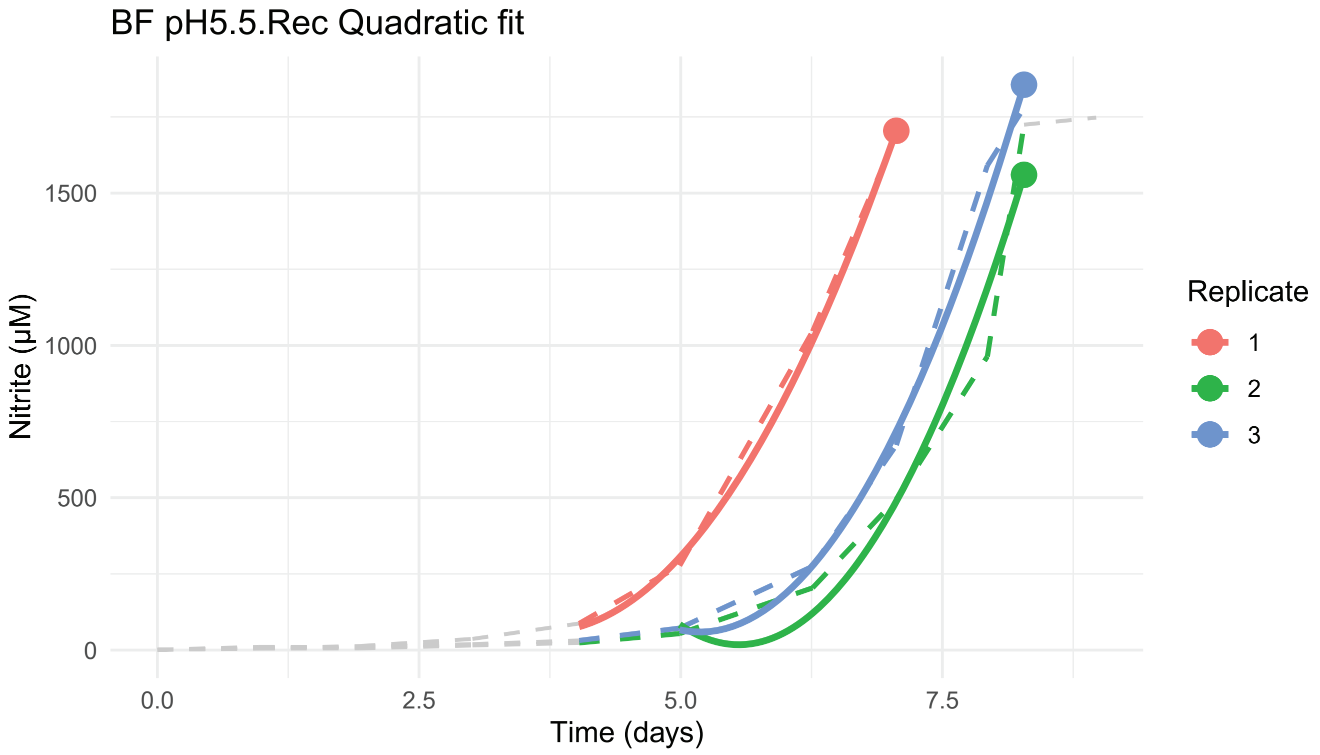

BF pH5.5.Rec Quadratic fit
Nitrite (µM)
1500
1000
500
0
0.0
2.5
5.0
7.5
Time (days)
Replicate
1
2
3

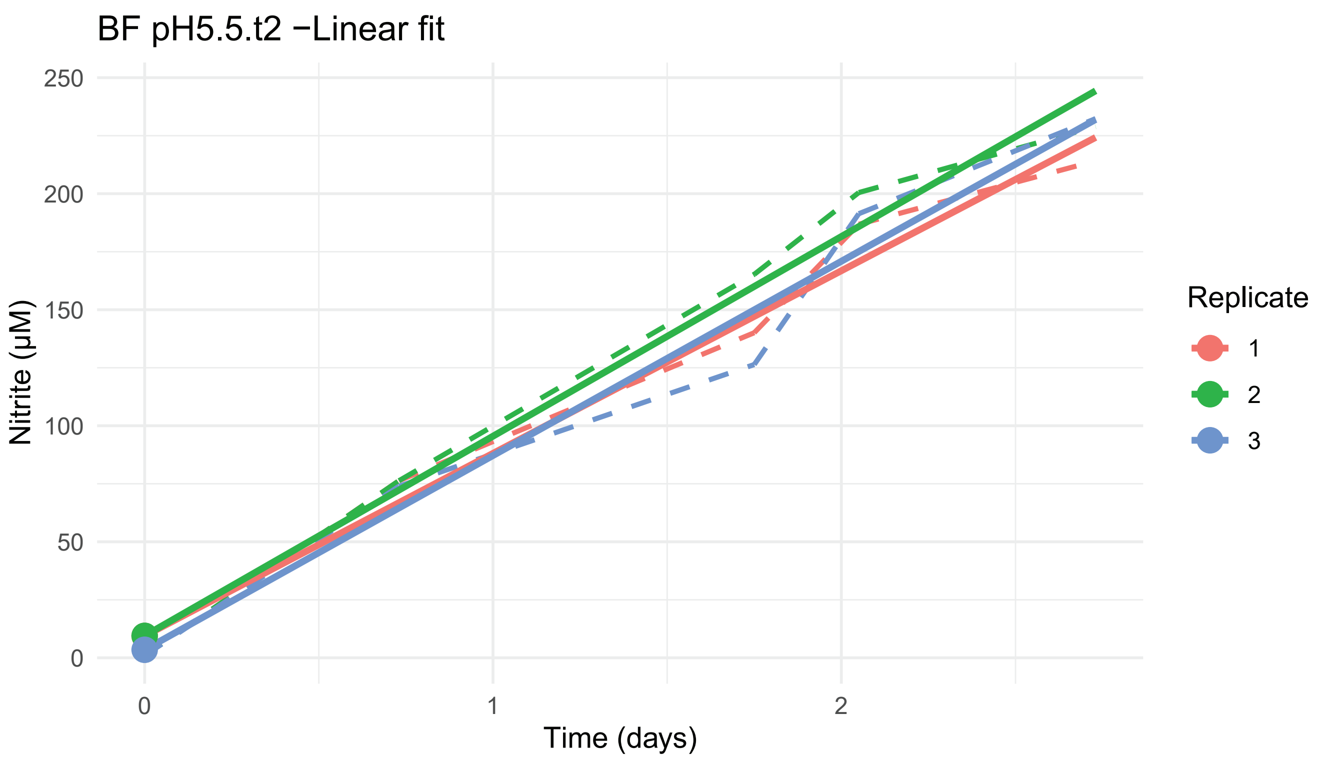
BF pH5.5.t2 −Linear fit
Nitrite (µM)
250
200
150
100
50
0
0
1
2
Time (days)
Replicate
1
2
3

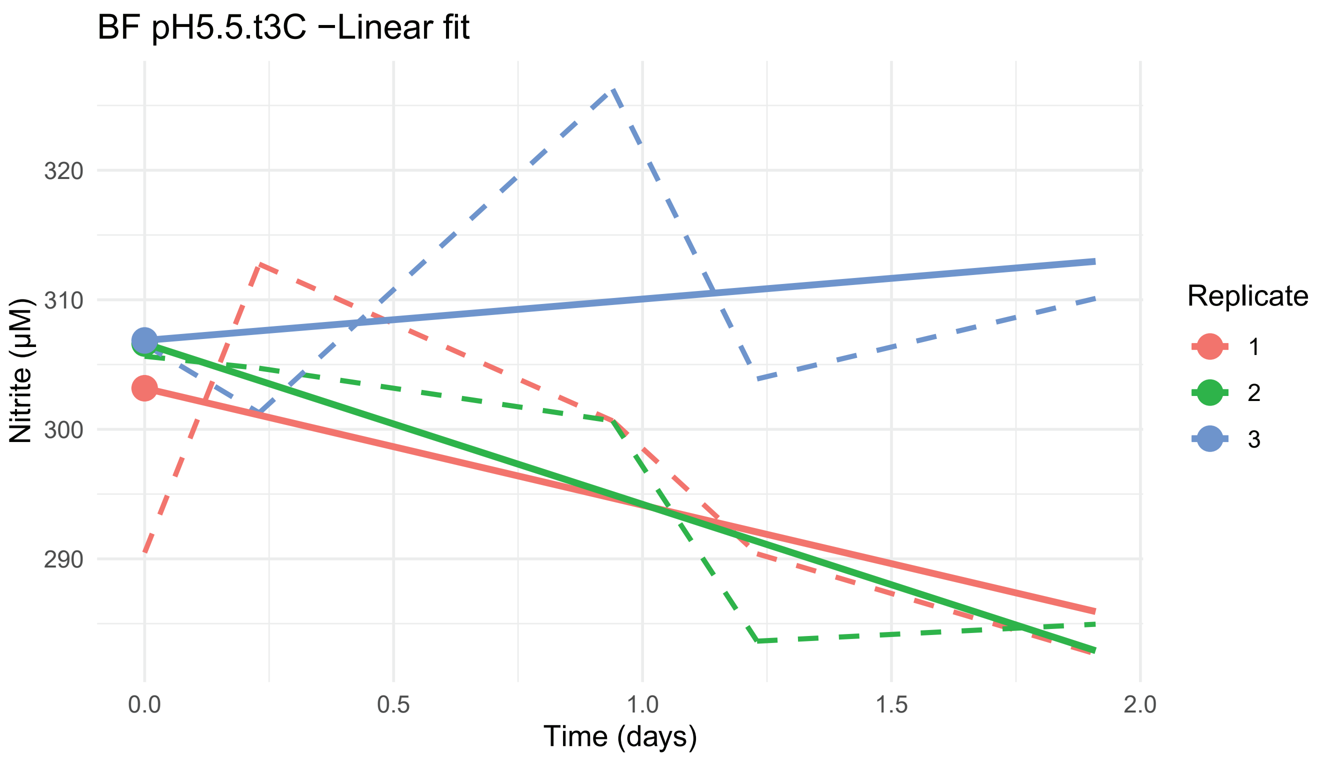
BF pH5.5.t3C −Linear fit
Nitrite (µM)
320
310
300
290
0.0
0.5
1.0
1.5
2.0
Time (days)
Replicate
1
2
3

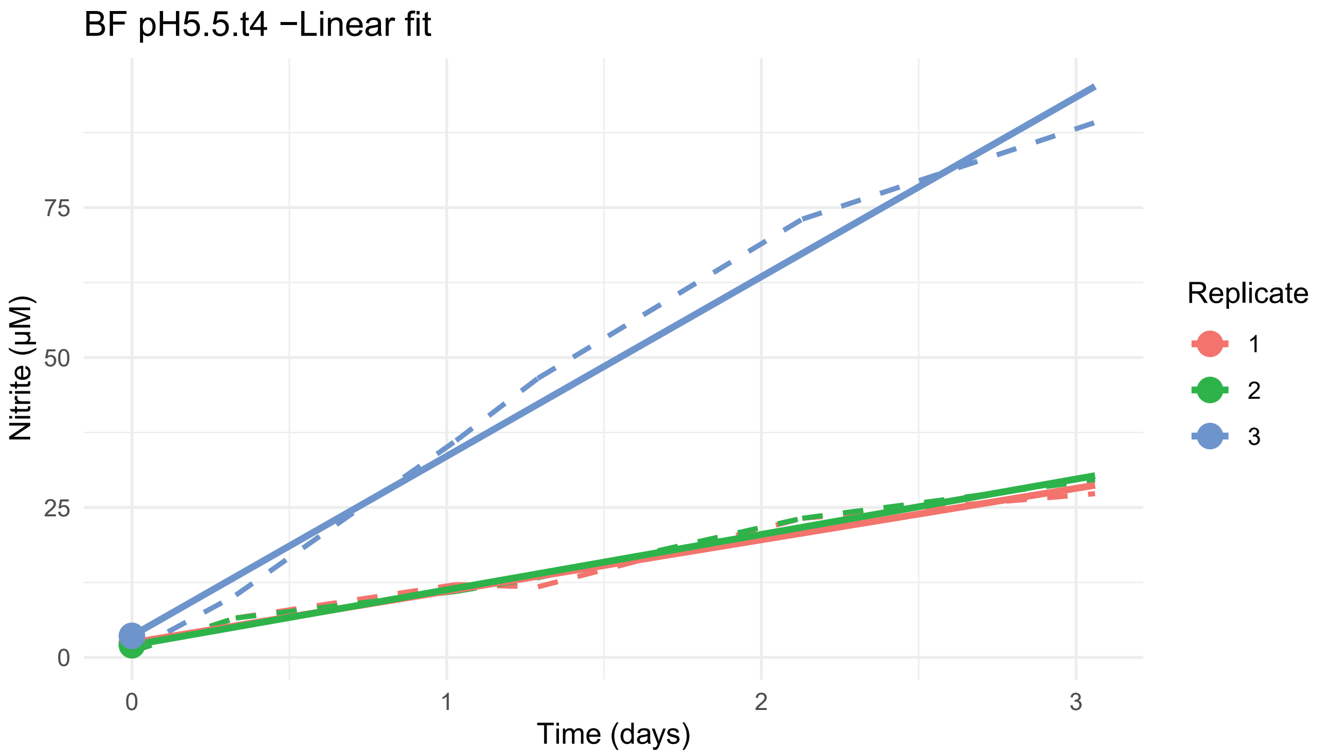
BF pH5.5.t4 −Linear fit
Nitrite (μM)
75
50
25
0
0
1
2
3
Time (days)
Replicate
1
2
3

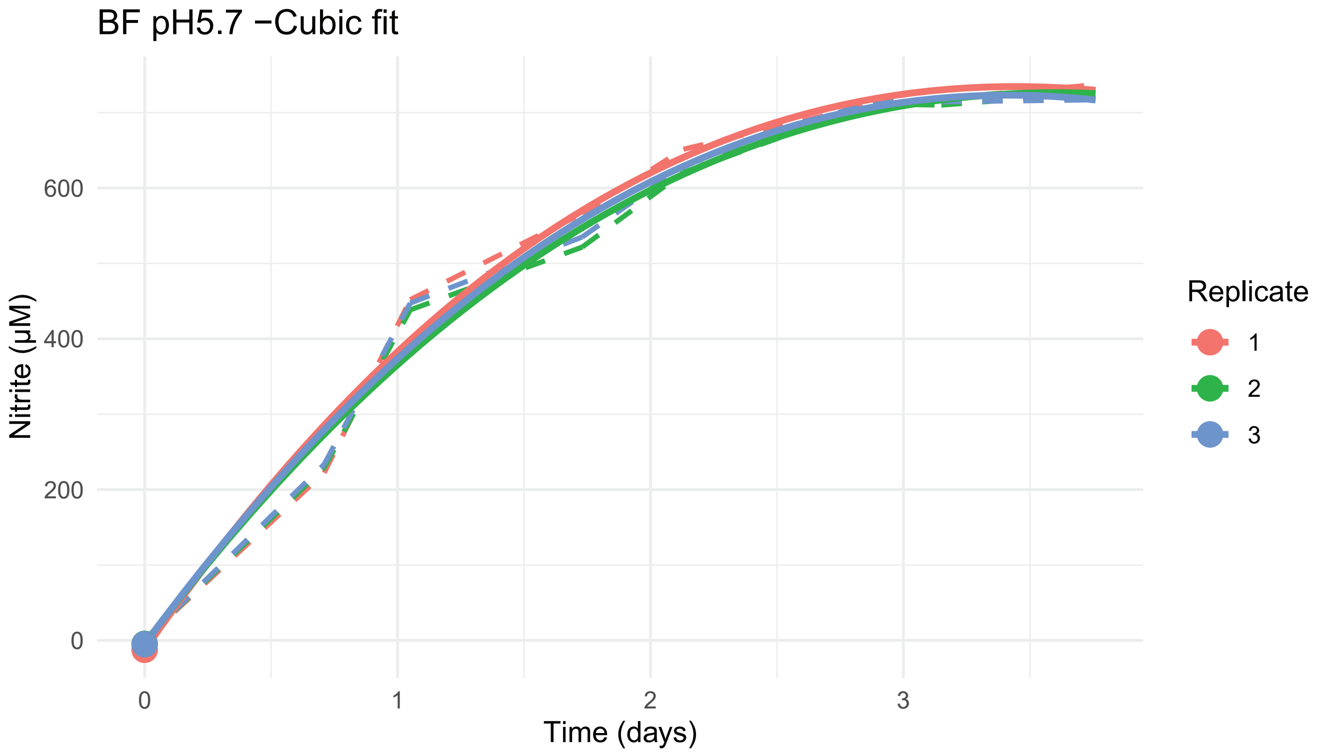
BF pH5.7 −Cubic fit
Nitrite (μM)
600
400
200
0
0
1
2
3
Time (days)
Replicate
1
2
3

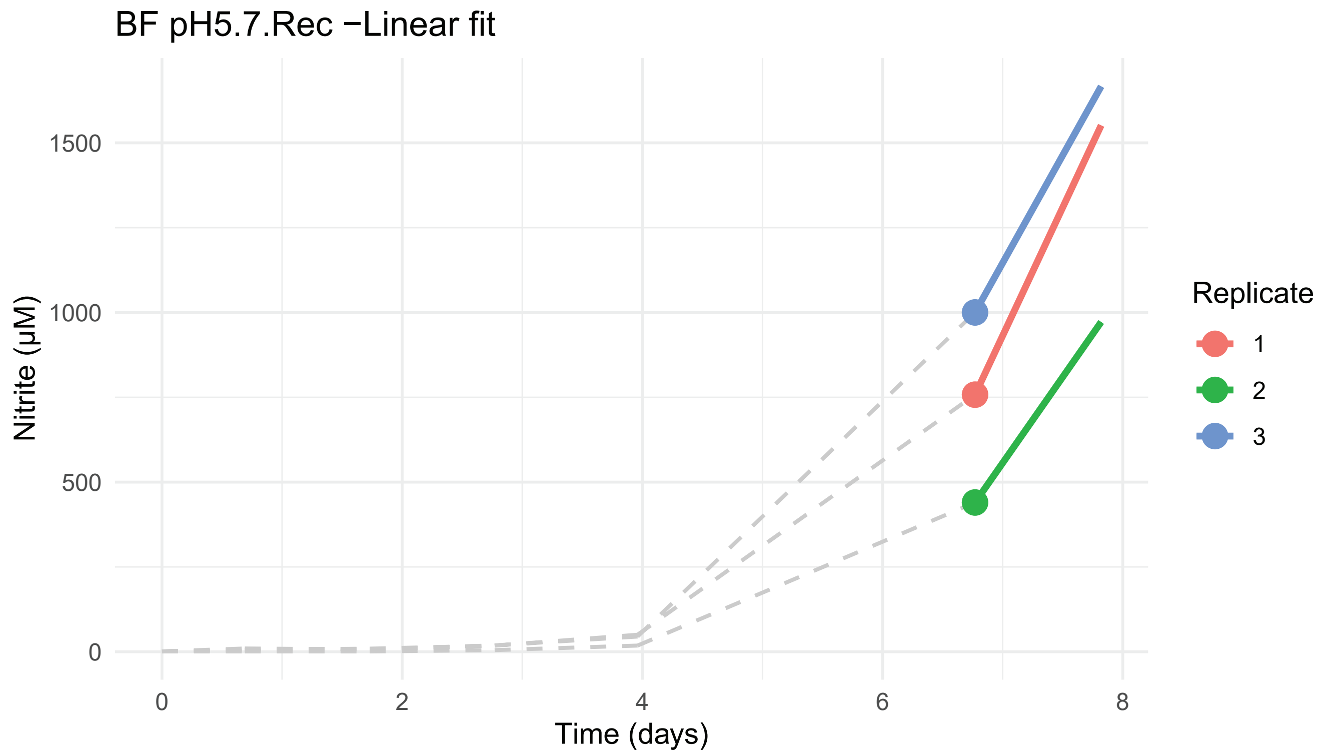
BF pH5.7.Rec −Linear fit
Nitrite (µM)
0
500
1000
1500
0
2
4
6
8
Time (days)
Replicate
1
2
3

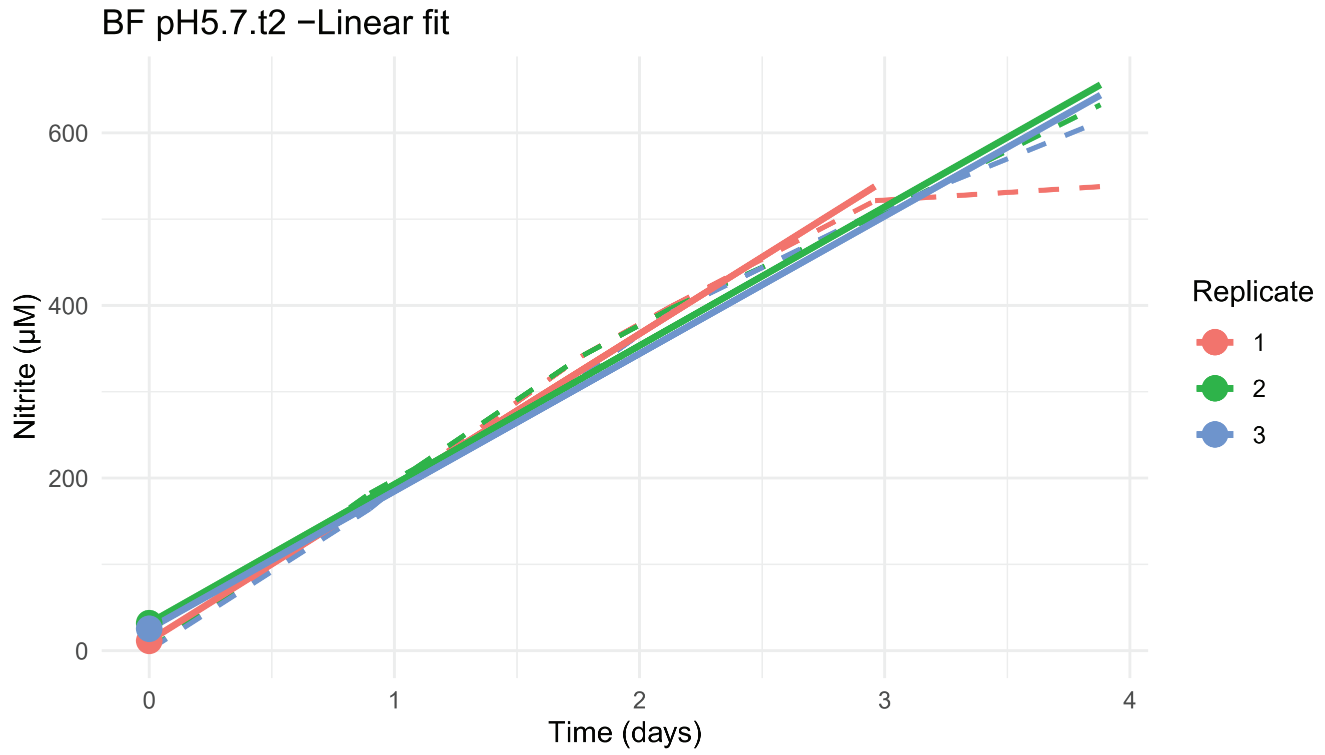
BF pH5.7.t2 −Linear fit
Nitrite (µM)
0
200
400
600
0
1
2
3
4
Time (days)
Replicate
1
2
3

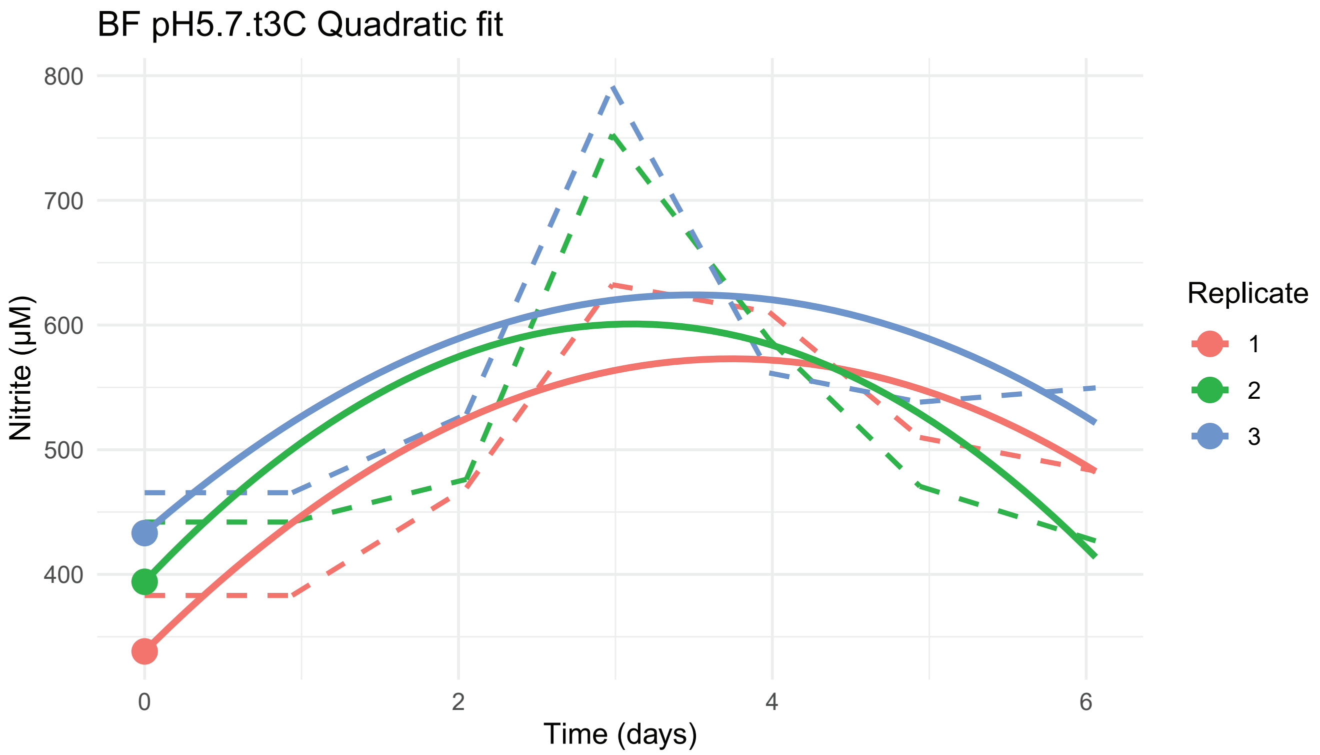
BF pH5.7.t3C Quadratic fit
800
700
600
500
400
Nitrite (µM)
0
2
4
6
Time (days)
Replicate
1
2
3

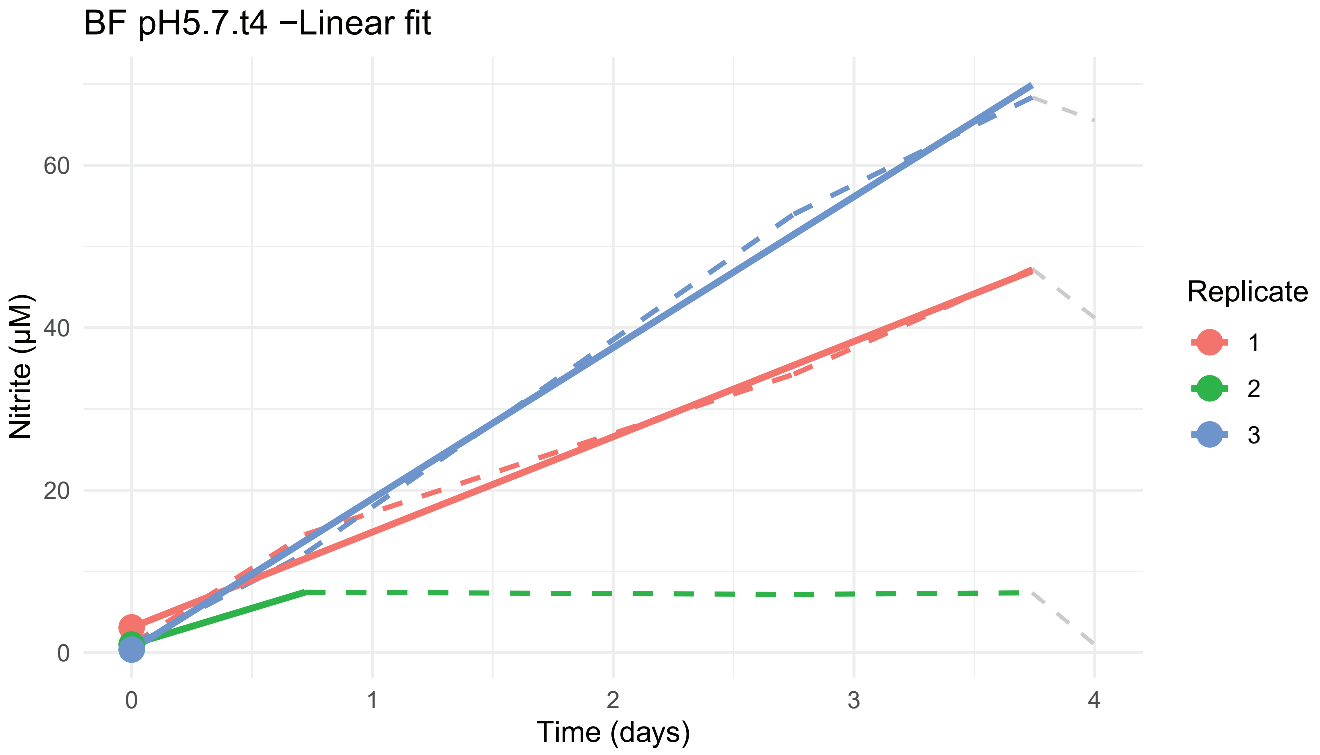
BF pH5.7.t4 −Linear fit
60
40
20
0
Nitrite (µM)
0
1
2
3
4
Time (days)
Replicate
1
2
3

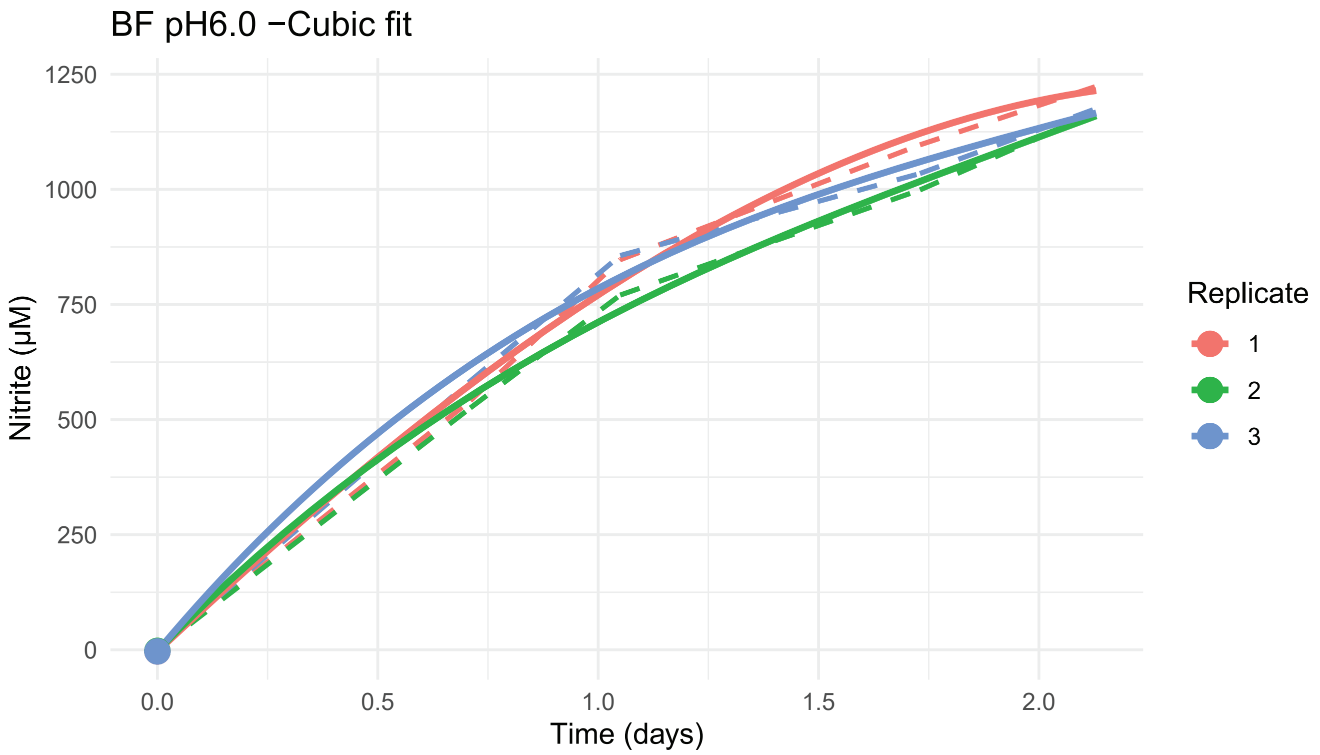
BF pH6.0 −Cubic fit
Nitrite (µM)
Time (days)
Replicate
1
2
3
0
250
500
750
1000
1250
0.0
0.5
1.0
1.5
2.0

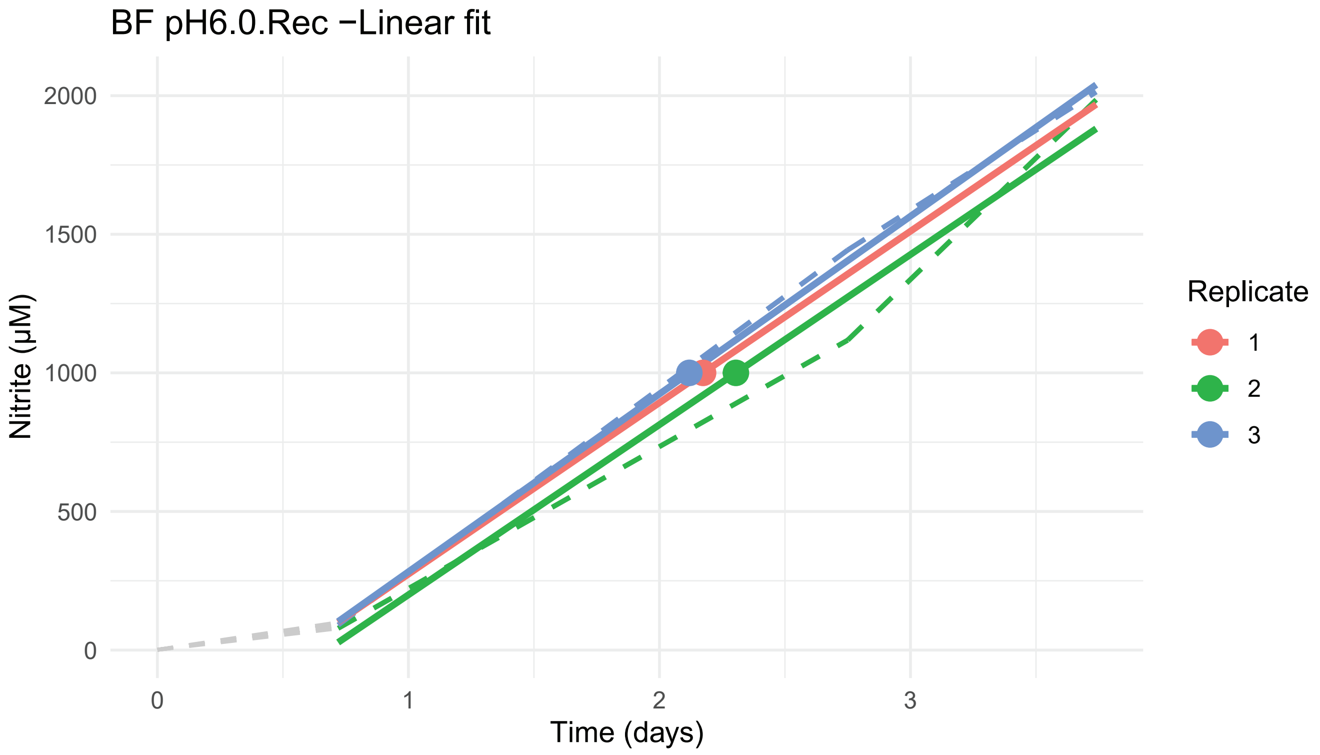
BF pH6.0.Rec −Linear fit
Nitrite (µM)
Time (days)
Replicate
1
2
3
0
500
1000
1500
2000
0
1
2
3

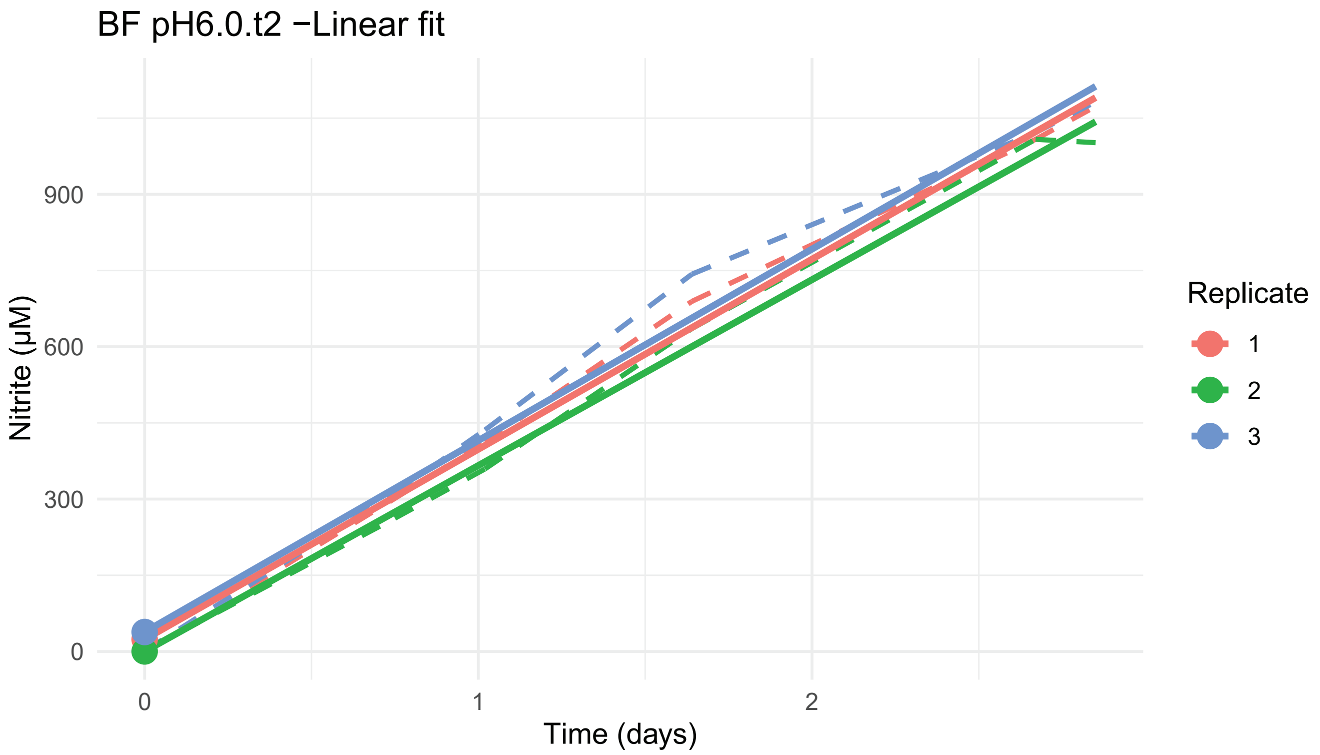
BF pH6.0.t2 −Linear fit
Nitrite (μM)
0
300
600
900
0
1
2
Time (days)
Replicate
1
2
3

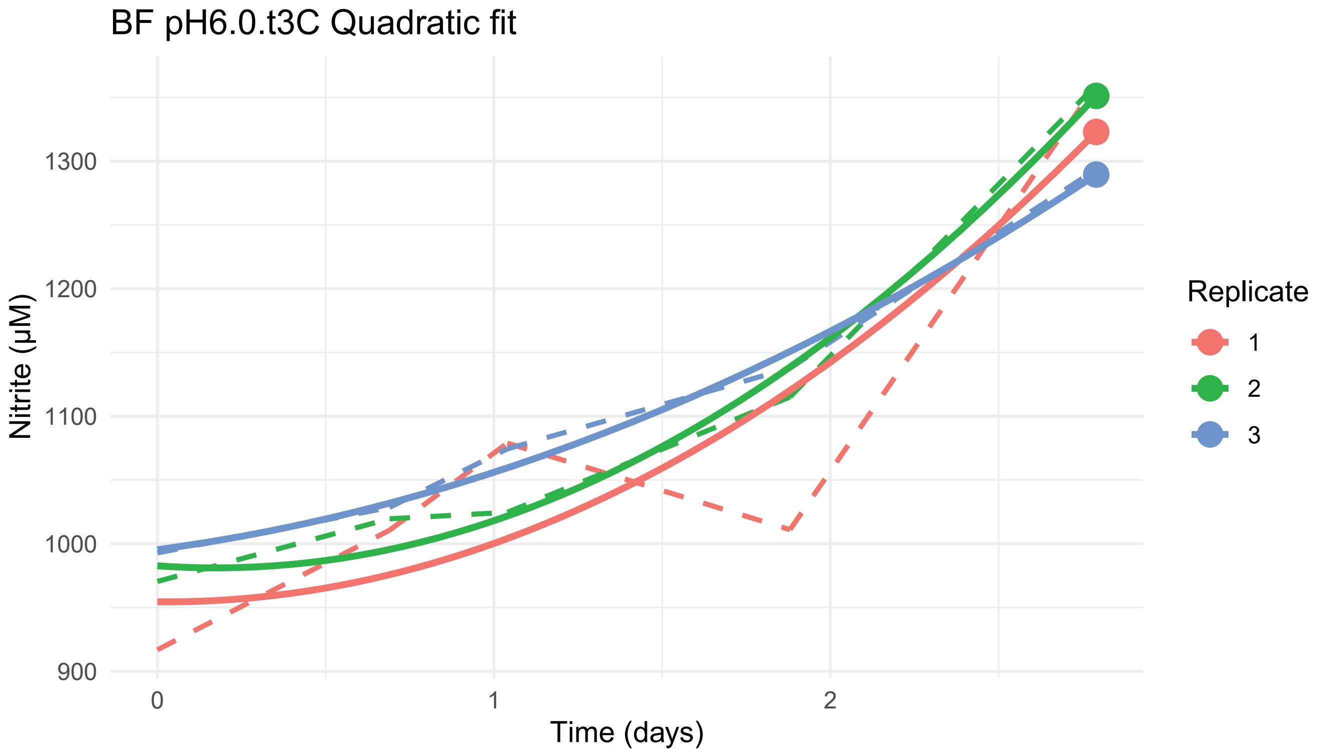
BF pH6.0.t3C Quadratic fit
Nitrite (μM)
900
1000
1100
1200
1300
0
1
2
Time (days)
Replicate
1
2
3

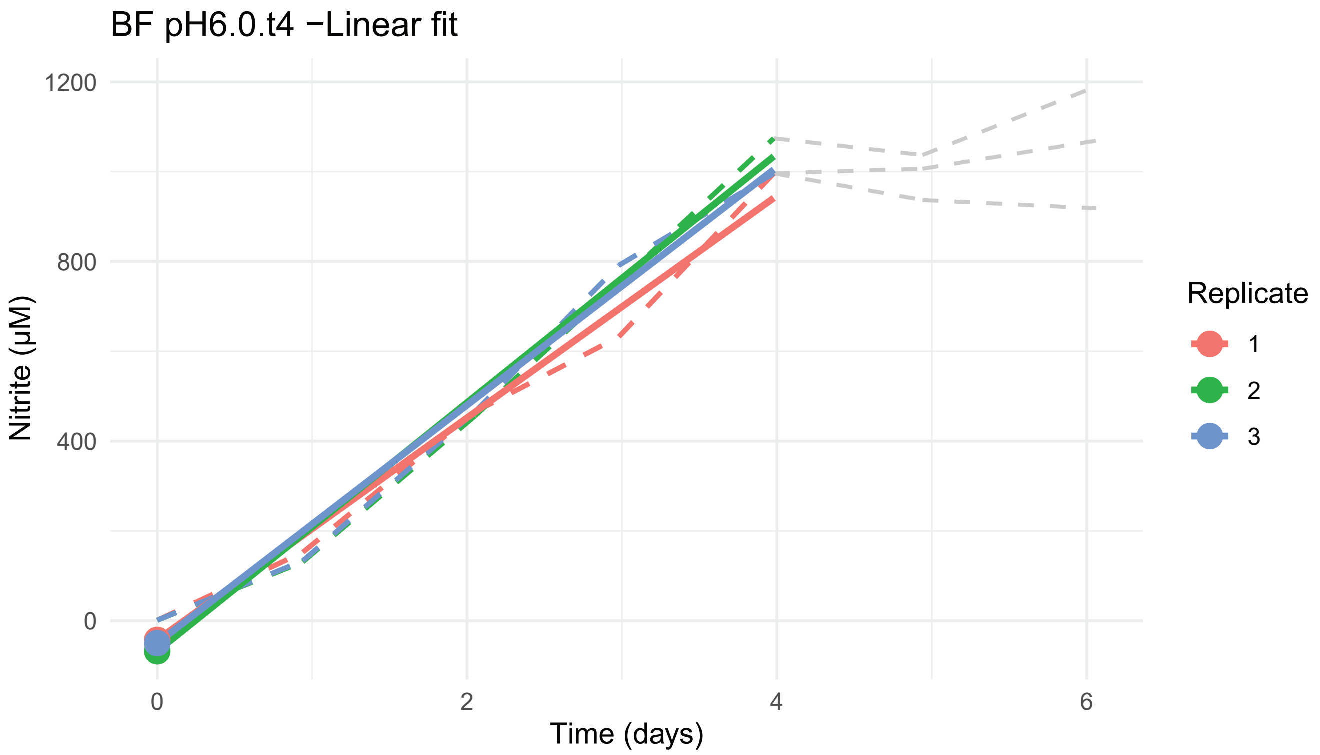
BF pH6.0.t4 −Linear fit
Nitrite (µM)
1200
800
400
0
0
2
4
6
Time (days)
Replicate
1
2
3

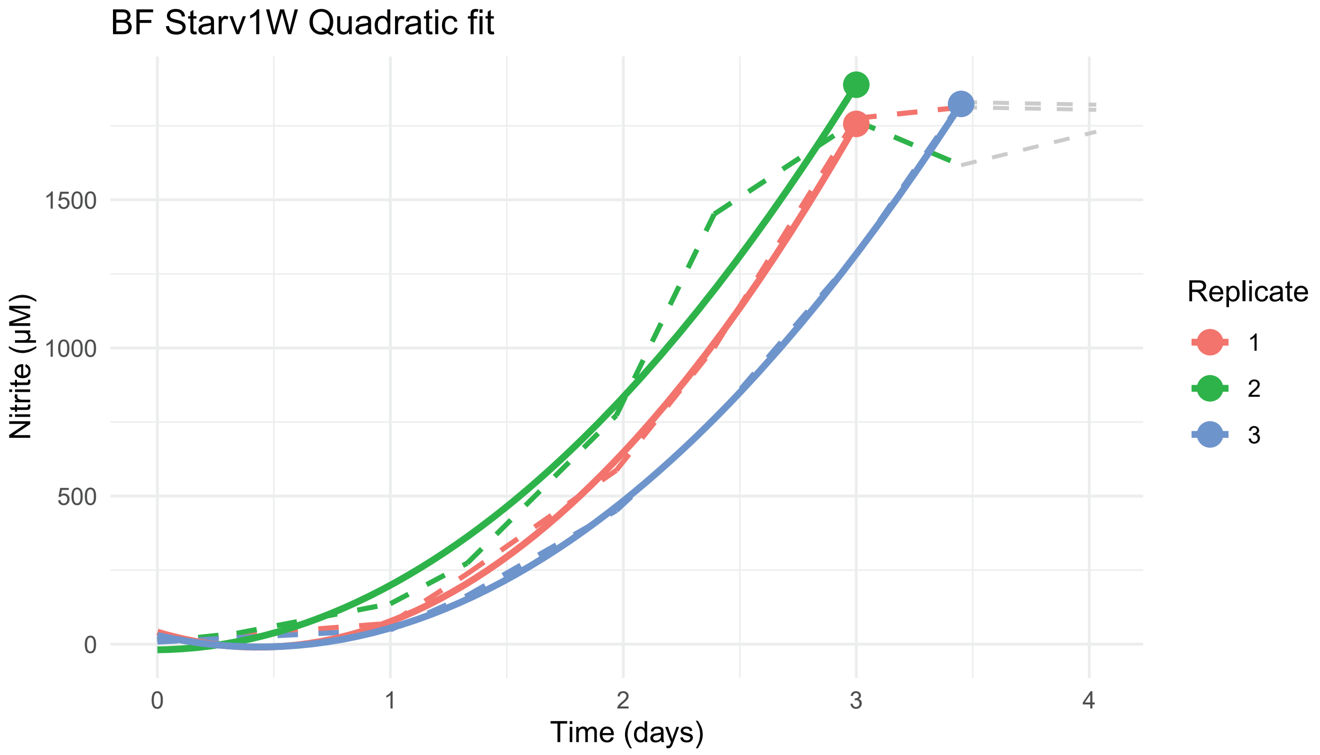
BF Starv1W Quadratic fit
Nitrite (µM)
1500
1000
500
0
0
1
2
3
4
Time (days)
Replicate
1
2
3

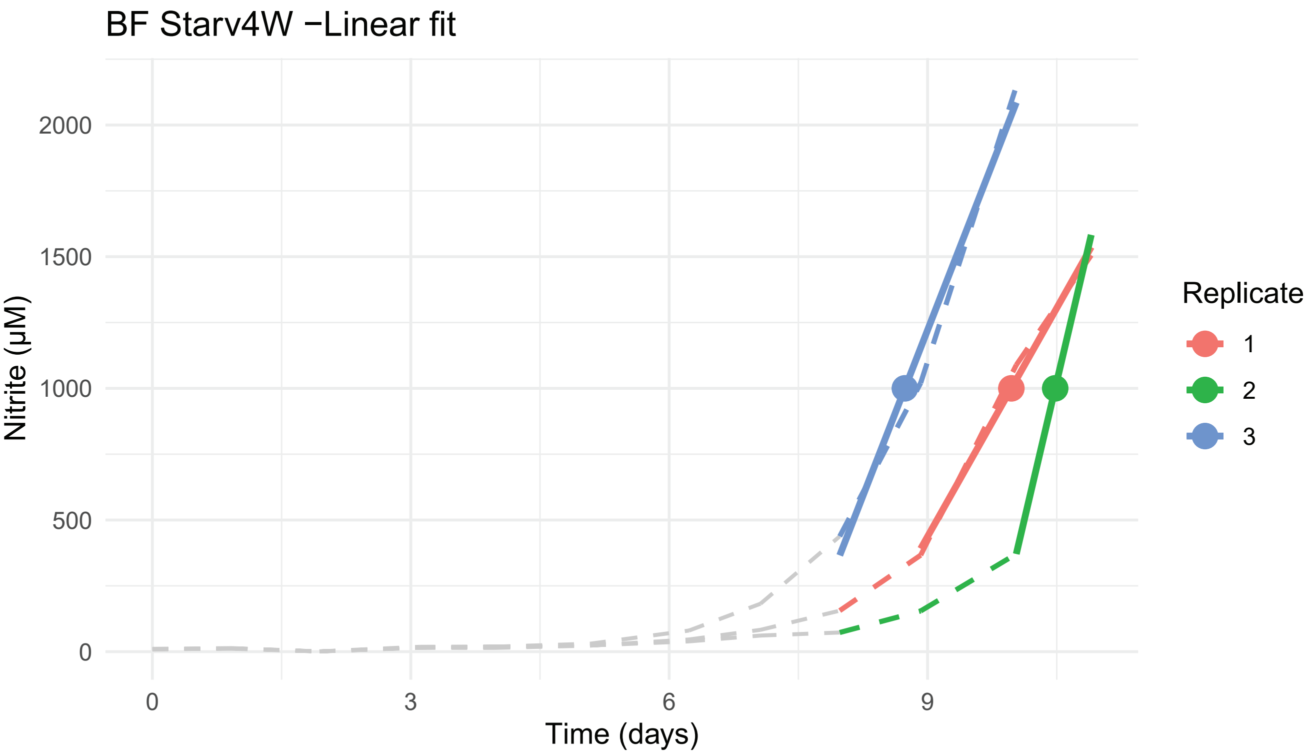
BF Starv4W −Linear fit
2000
1500
1000
500
0
Nitrite (µM)
0
3
6
9
Time (days)
Replicate
1
2
3

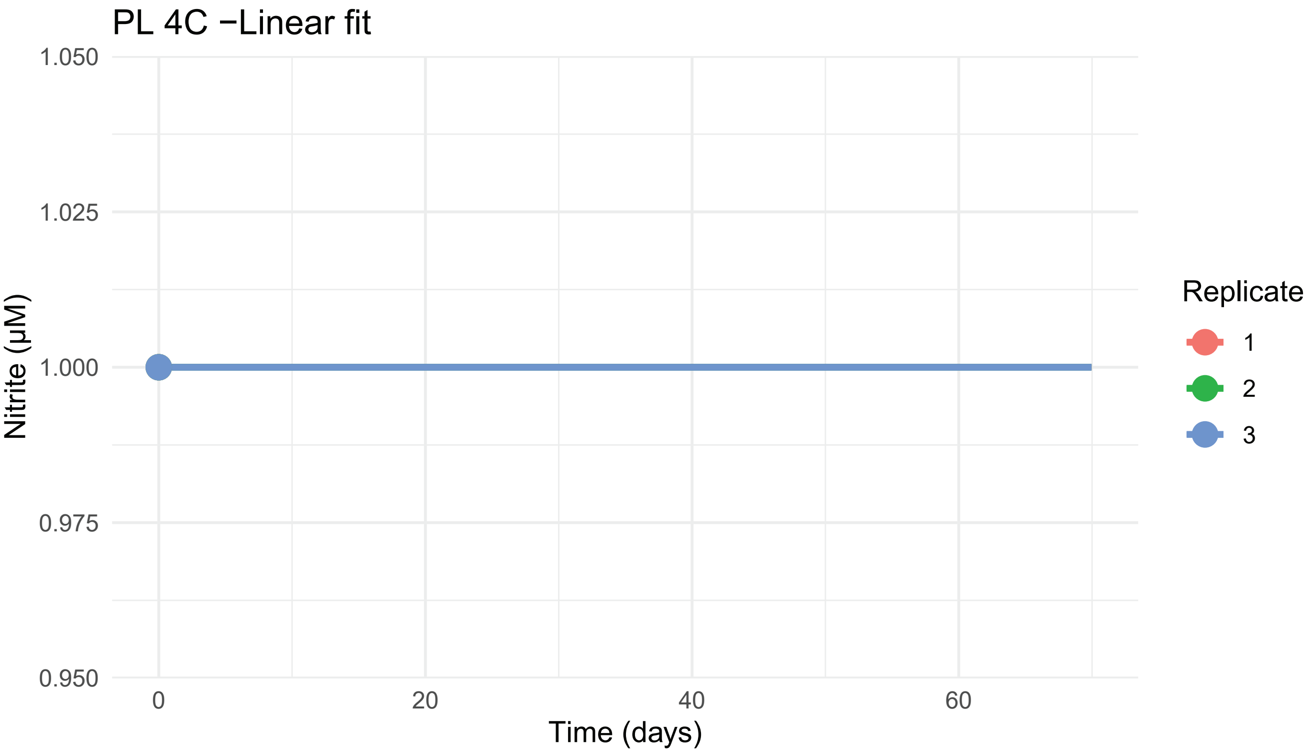
PL 4C −Linear fit
1.050
1.025
1.000
0.975
0.950
Nitrite (µM)
0
20
40
60
Time (days)
Replicate
1
2
3

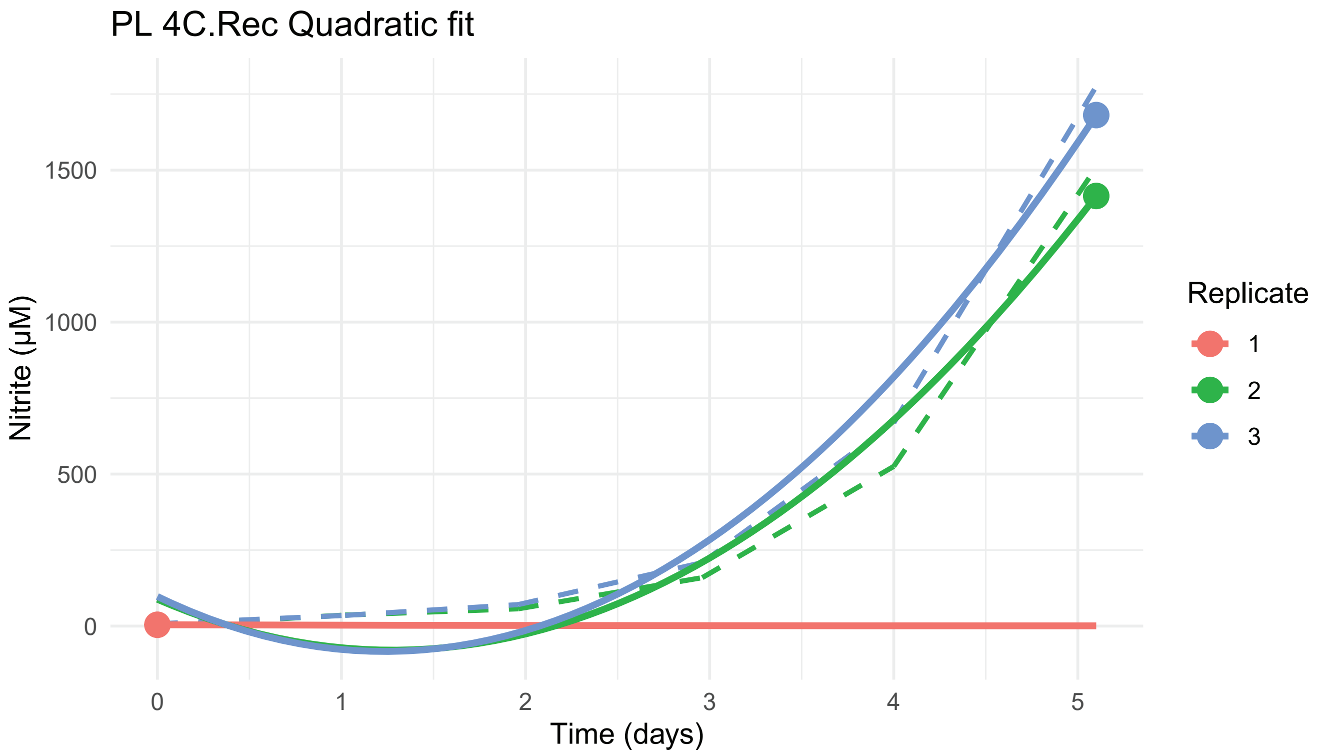

PL 4C.Rec Quadratic fit
Nitrite (µM)
1500
1000
500
0
0
1
2
3
4
5
Time (days)
Replicate
1
2
3


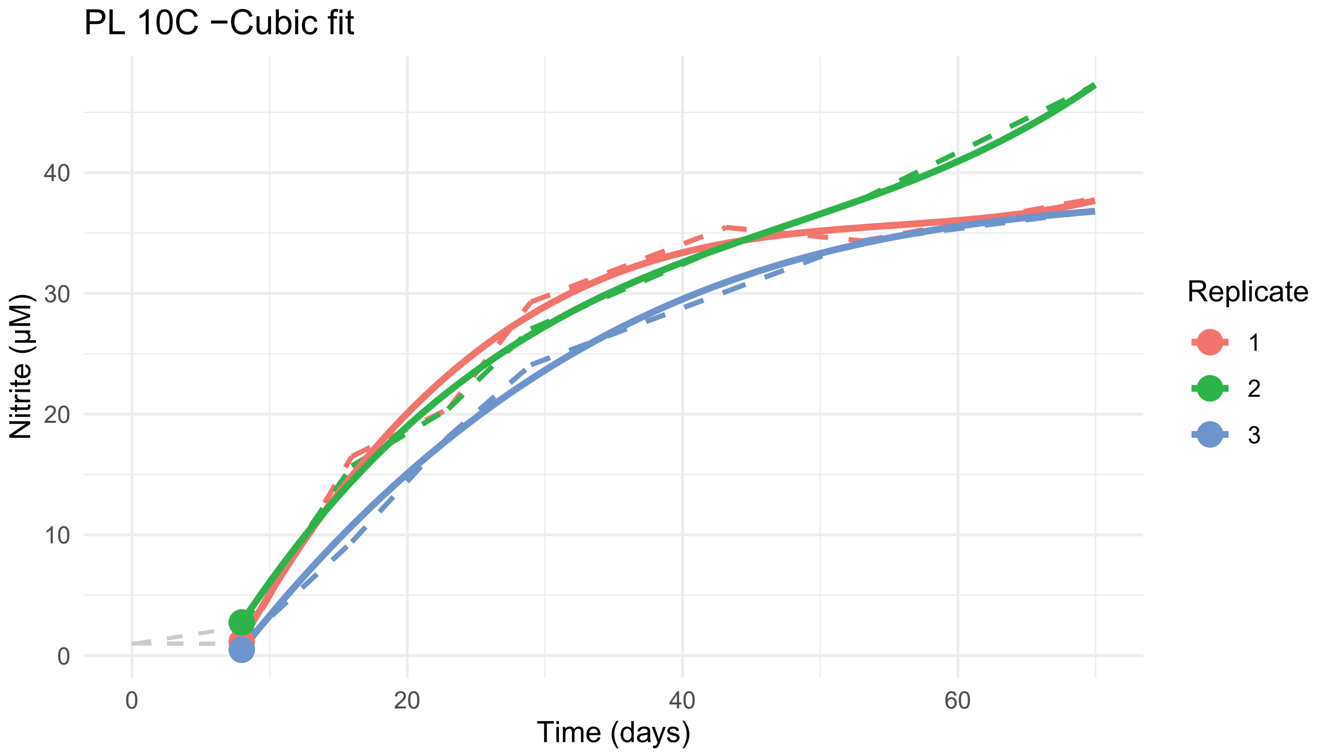

PL 10C −Cubic fit
Nitrite (µM)
40
30
20
10
0
0
20
40
60
Time (days)
Replicate
1
2
3

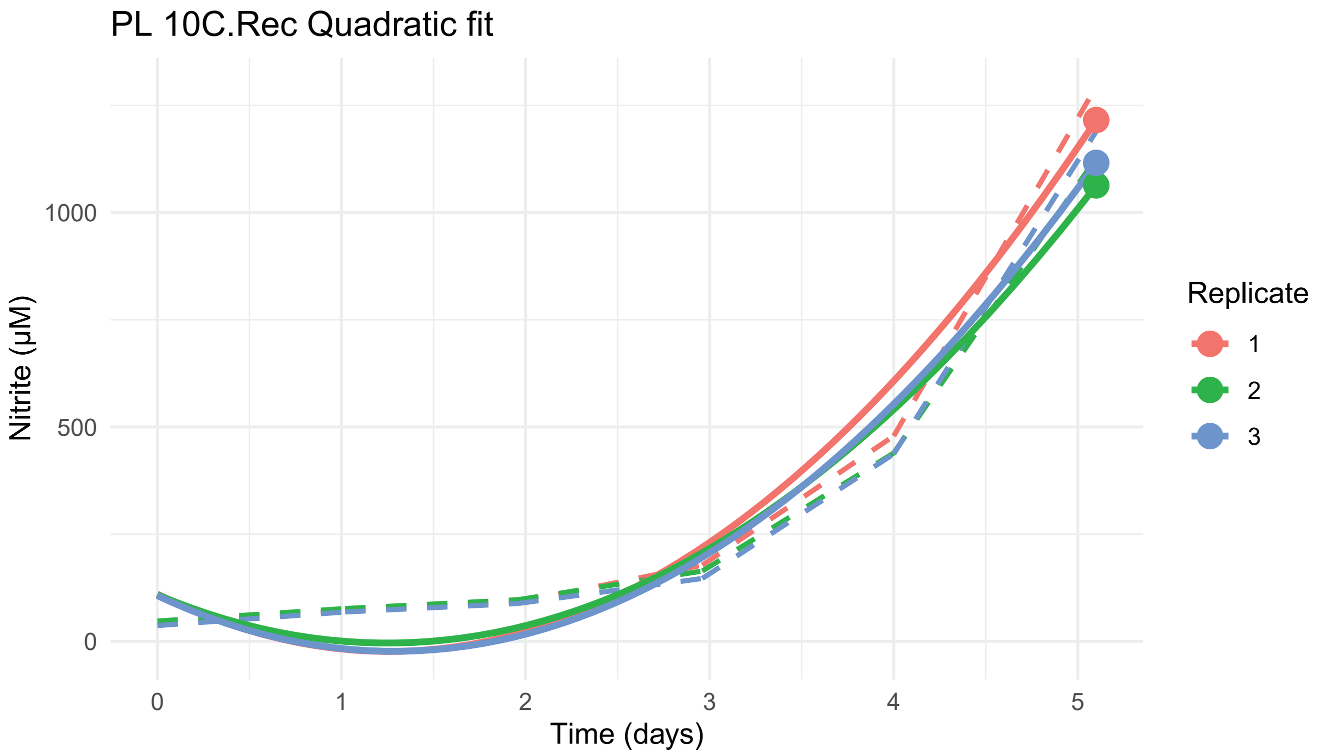

PL 10C.Rec Quadratic fit
Nitrite (μM)
0
500
1000
0
1
2
3
4
5
Time (days)
Replicate
1
2
3


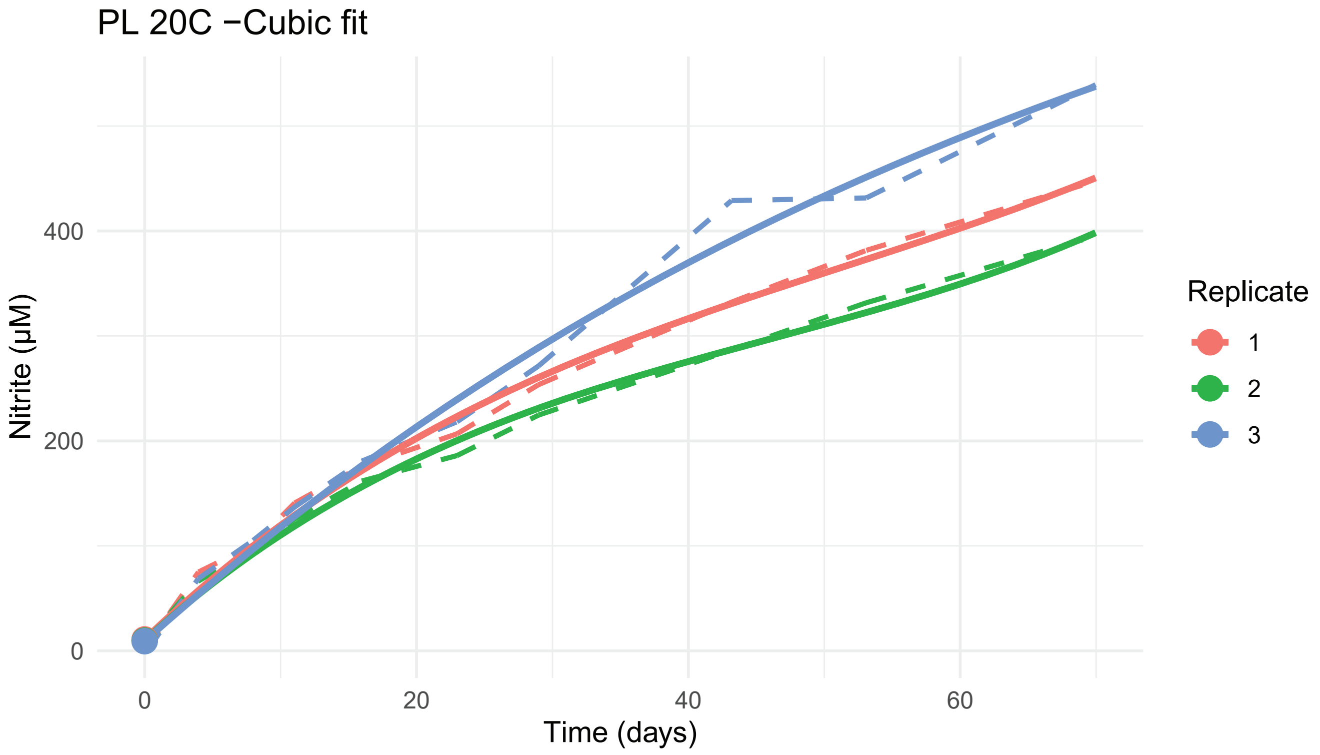

PL 20C −Cubic fit
Nitrite (μM)
0
200
400
0
20
40
60
Time (days)
Replicate
1
2
3

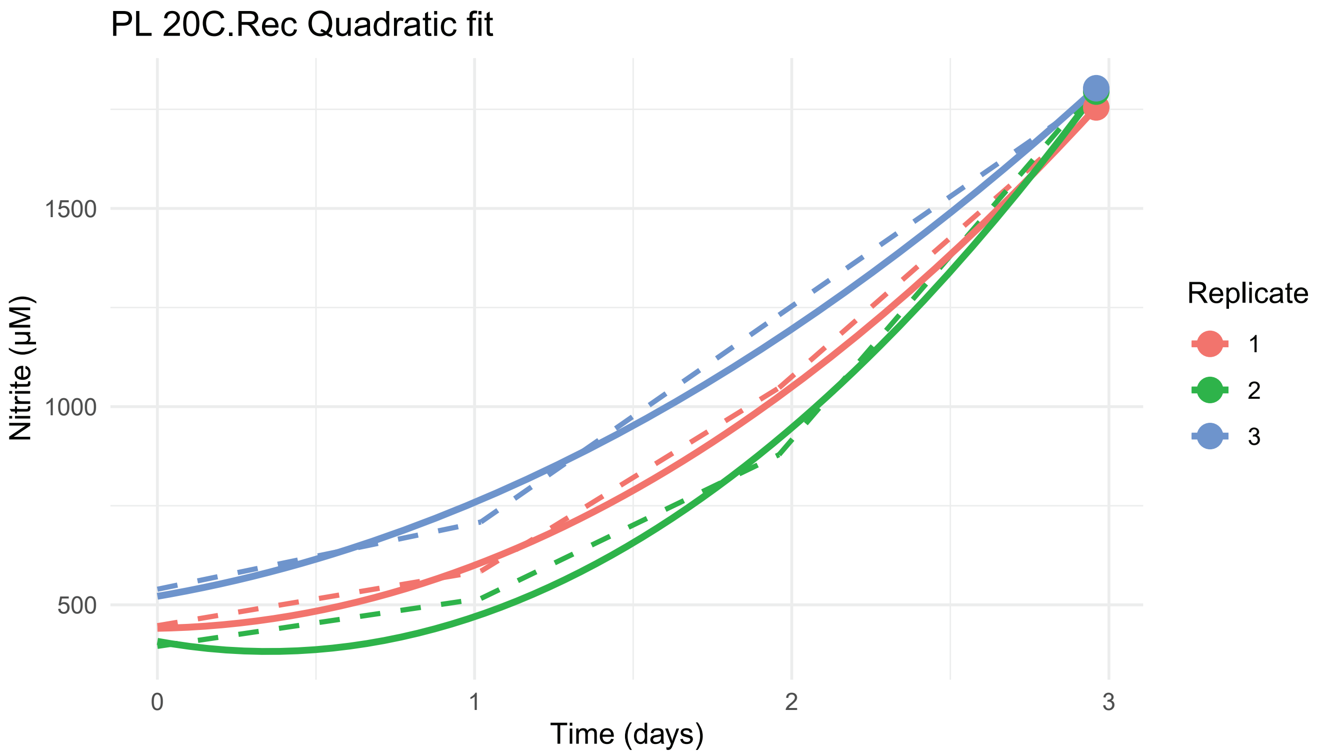
PL 20C.Rec Quadratic fit
Nitrite (µM)
1500
1000
500
0
1
2
3
Time (days)
Replicate
1
2
3

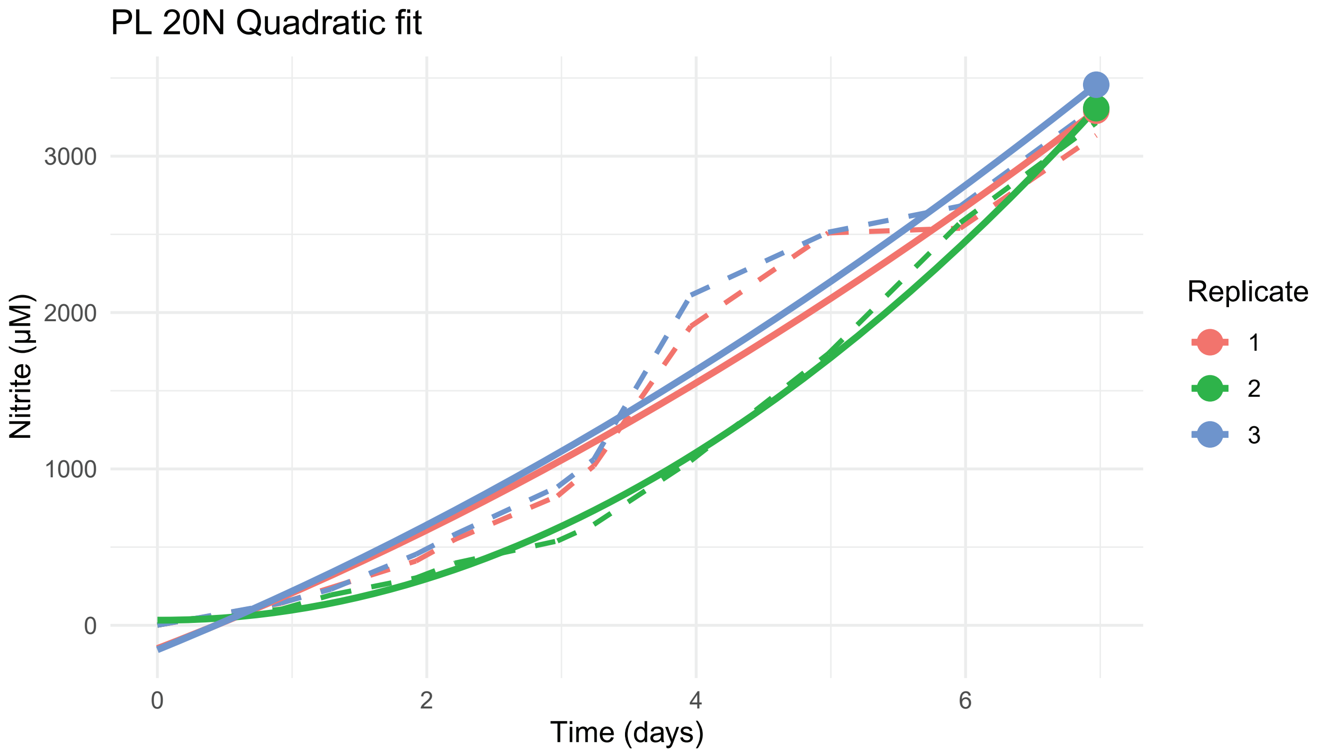
PL 20N Quadratic fit
Nitrite (µM)
3000
2000
1000
0
0
2
4
6
Time (days)
Replicate
1
2
3

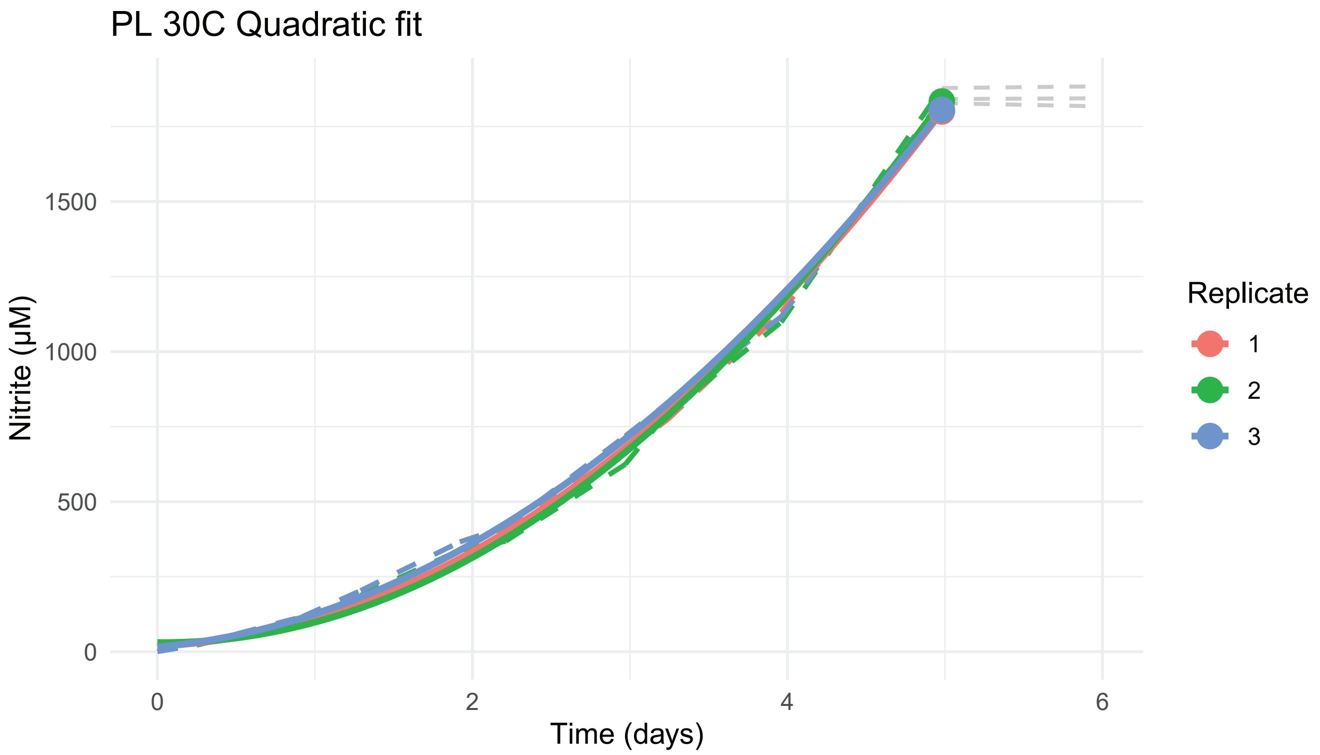
PL 30C Quadratic fit
Nitrite (µM)
0
500
1000
1500
0
2
4
6
Time (days)
Replicate
1
2
3

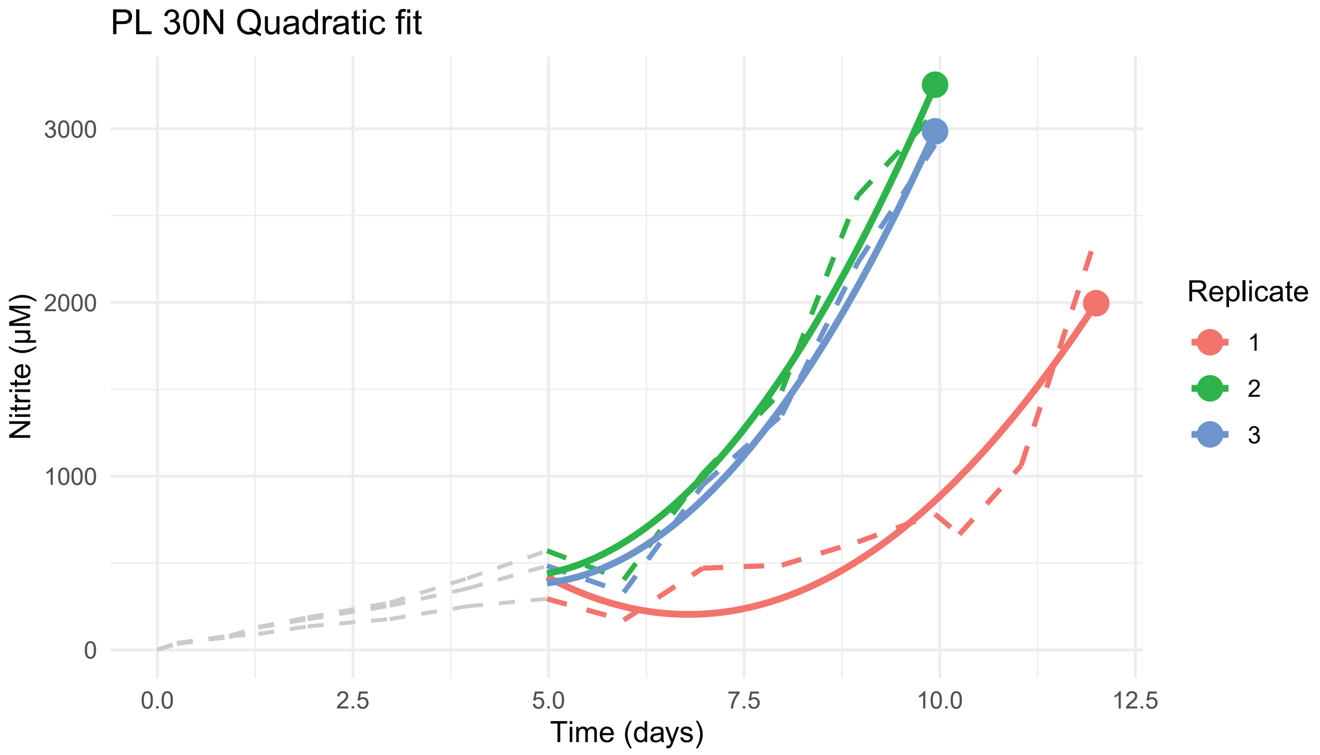
PL 30N Quadratic fit
Nitrite (µM)
0
1000
2000
3000
0.0
2.5
5.0
7.5
10.0
12.5
Time (days)
Replicate
1
2
3

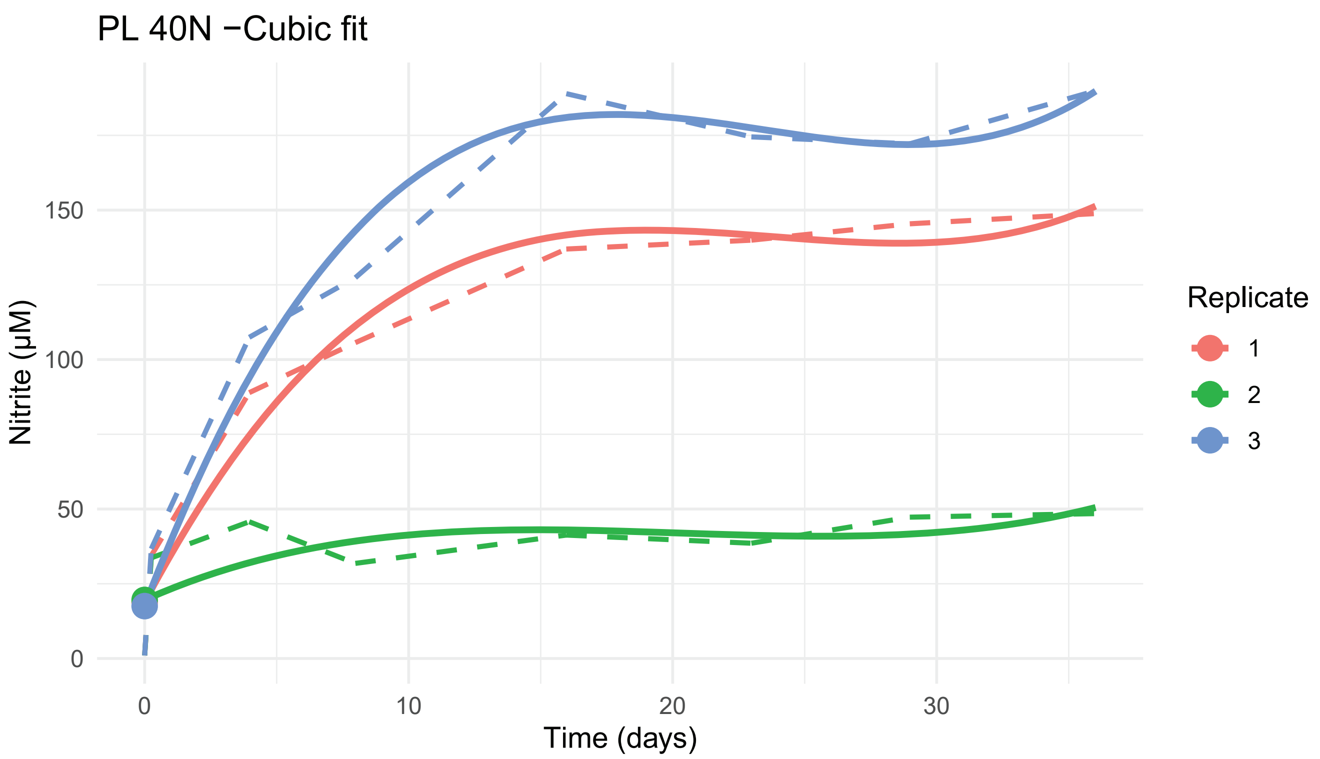
PL 40N −Cubic fit
Nitrite (µM)
Time (days)
Replicate
1
2
3

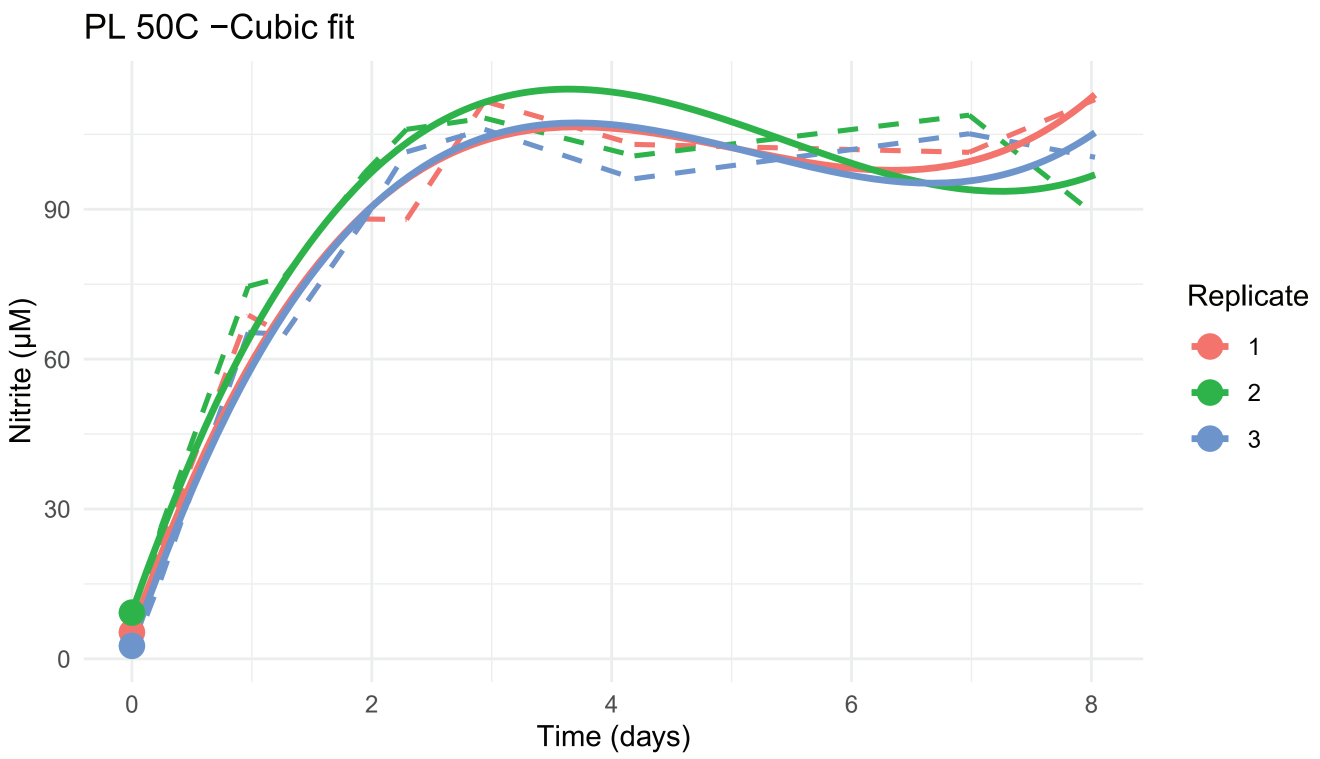
PL 50C −Cubic fit
Nitrite (µM)
Time (days)
Replicate
1
2
3

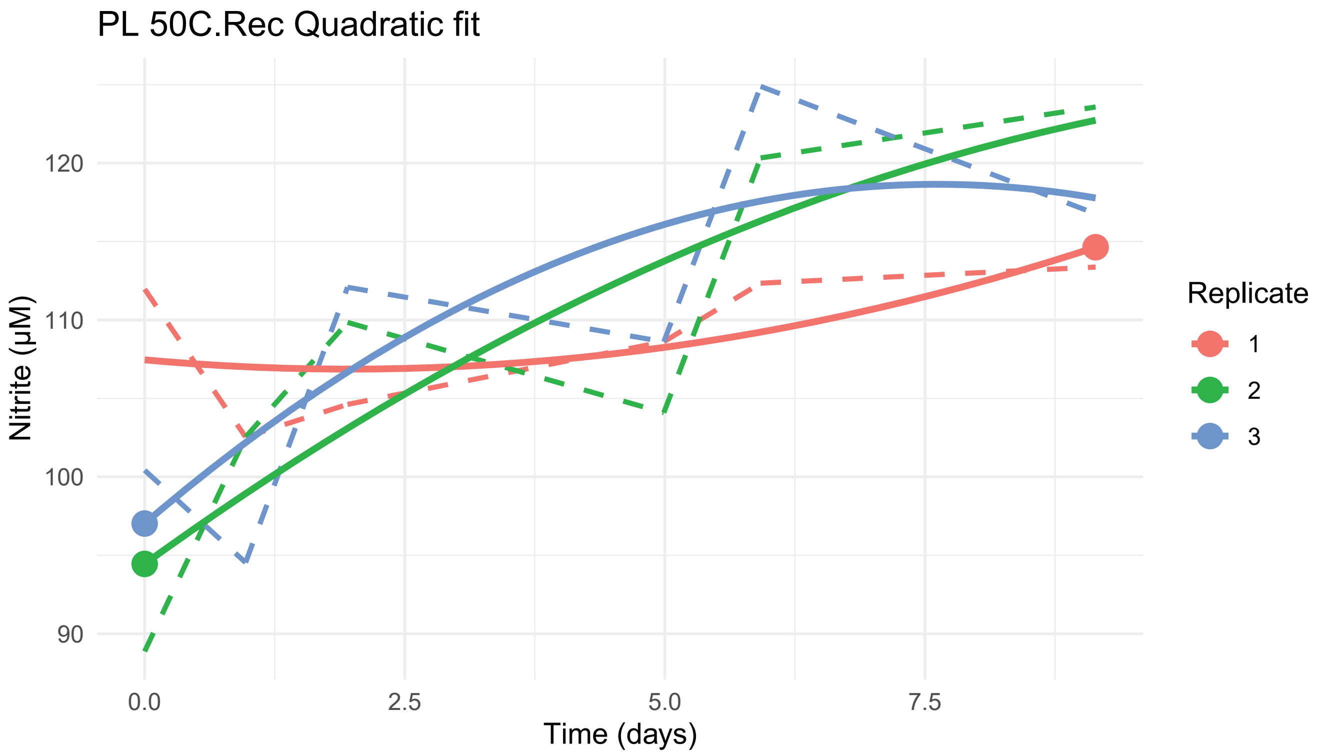
PL 50C.Rec Quadratic fit
Nitrite (µM)
Time (days)
90
100
110
120
0.0
2.5
5.0
7.5
Replicate
1
2
3

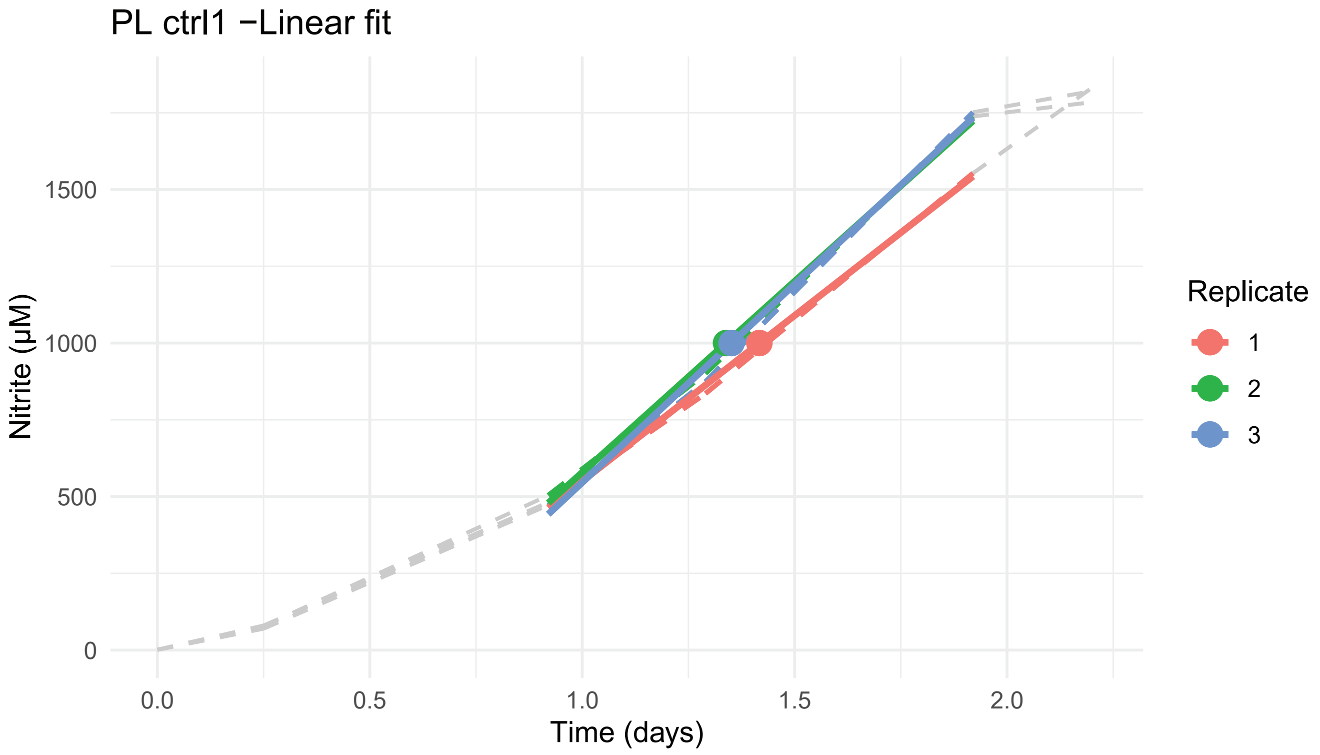
PL ctrl1 −Linear fit
Nitrite (µM)
Time (days)
0
500
1000
1500
0.0
0.5
1.0
1.5
2.0
Replicate
1
2
3

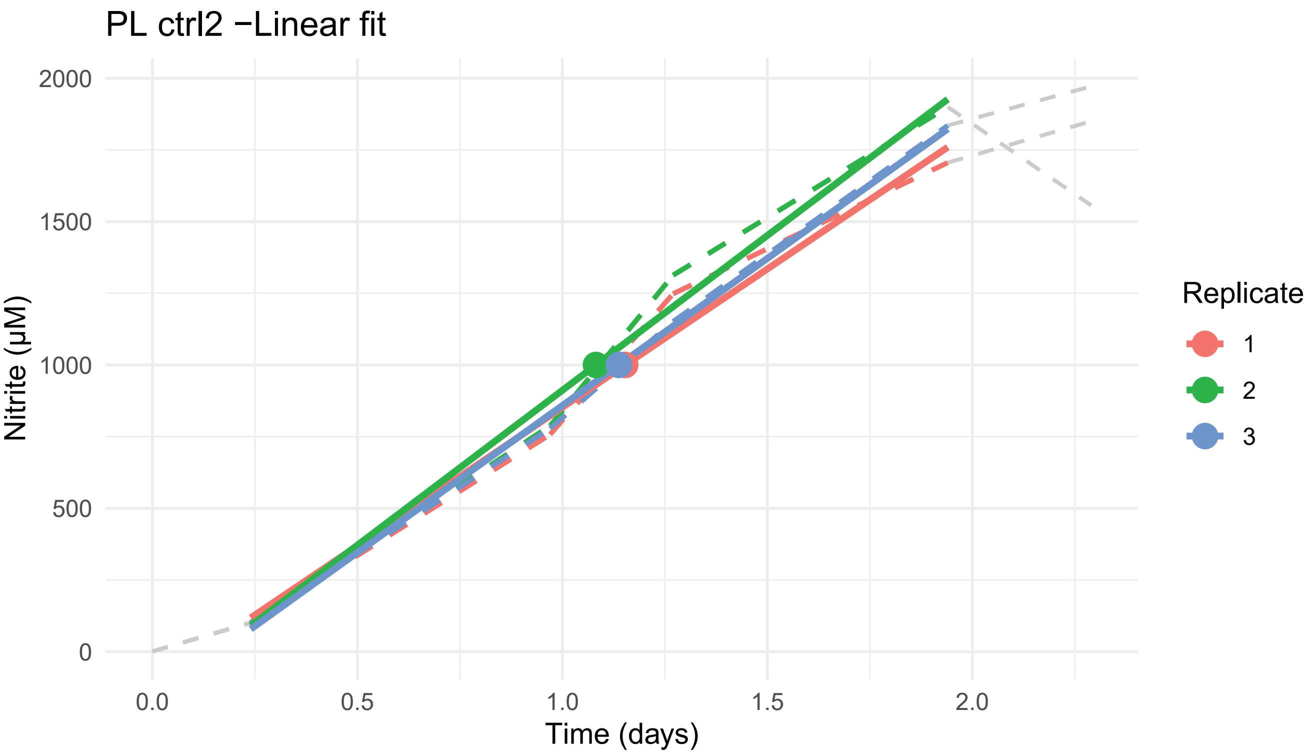
PL ctrl2 −Linear fit
Nitrite (µM)
2000
1500
1000
500
0
0.0
0.5
1.0
1.5
2.0
Time (days)
Replicate
1
2
3

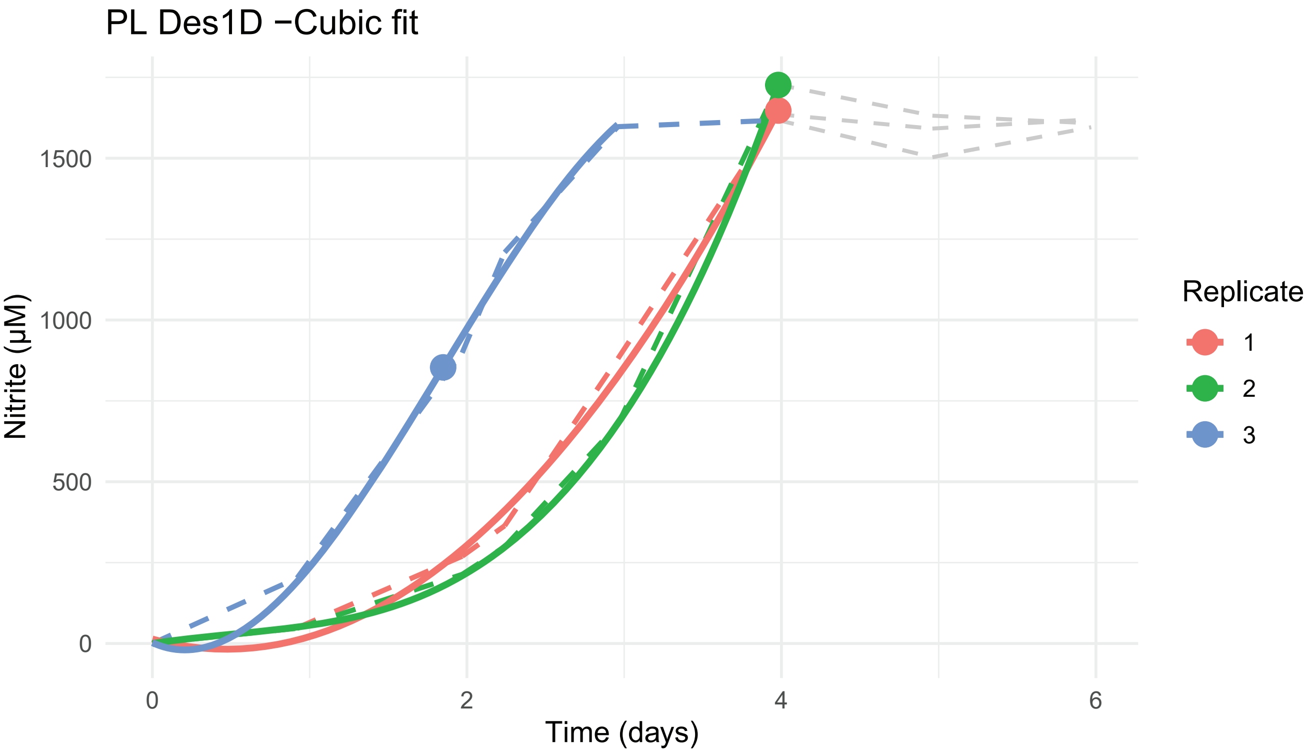
PL Des1D −Cubic fit
Nitrite (µM)
1500
1000
500
0
0
2
4
6
Time (days)
Replicate
1
2
3

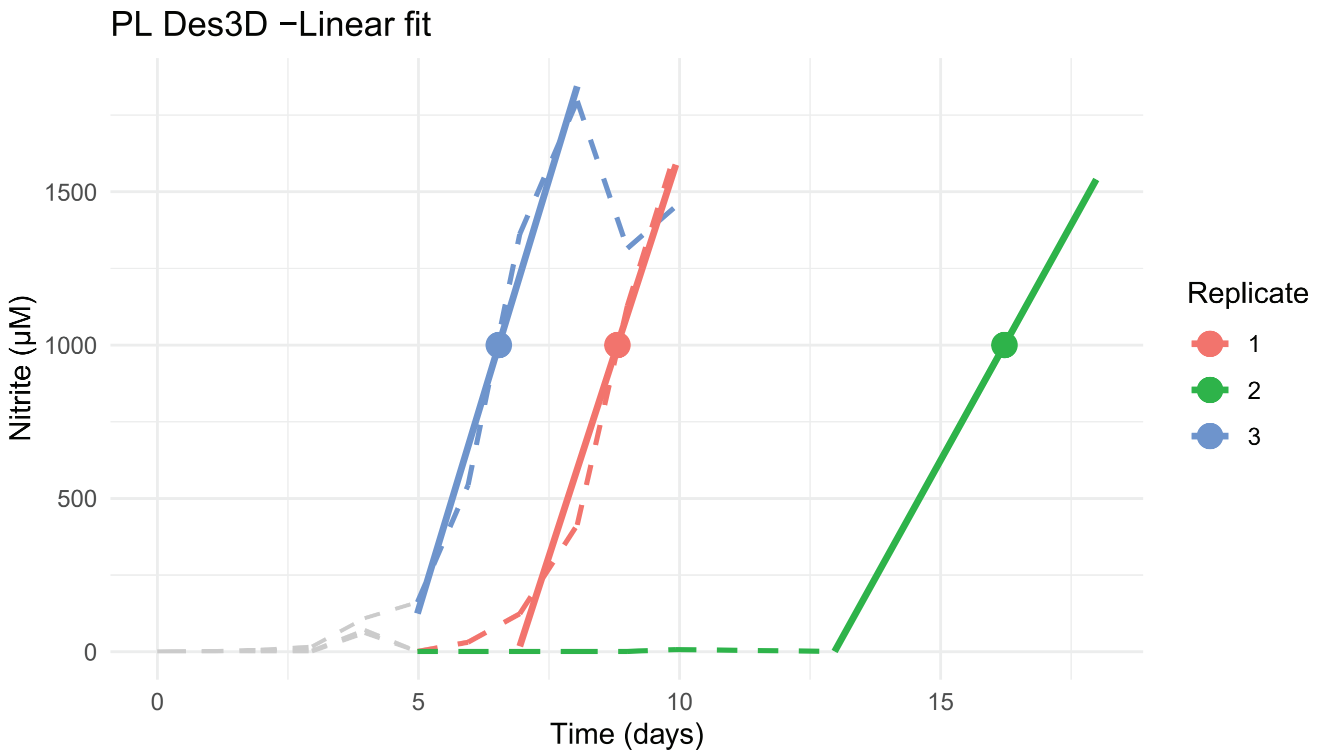
PL Des3D −Linear fit
Nitrite (µM)
0
500
1000
1500
0
5
10
15
Time (days)
Replicate
1
2
3

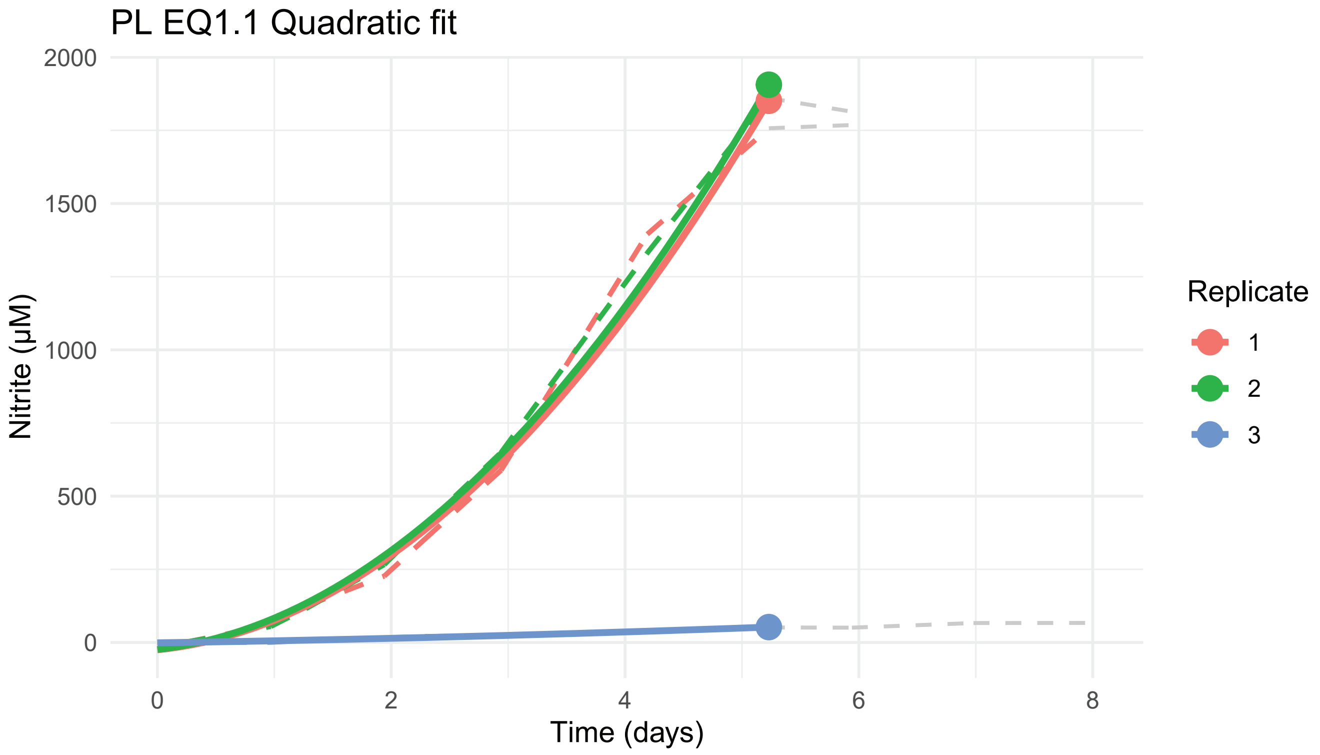
PL EQ1.1 Quadratic fit
Nitrite (µM)
0
500
1000
1500
2000
0
2
4
6
8
Time (days)
Replicate
1
2
3

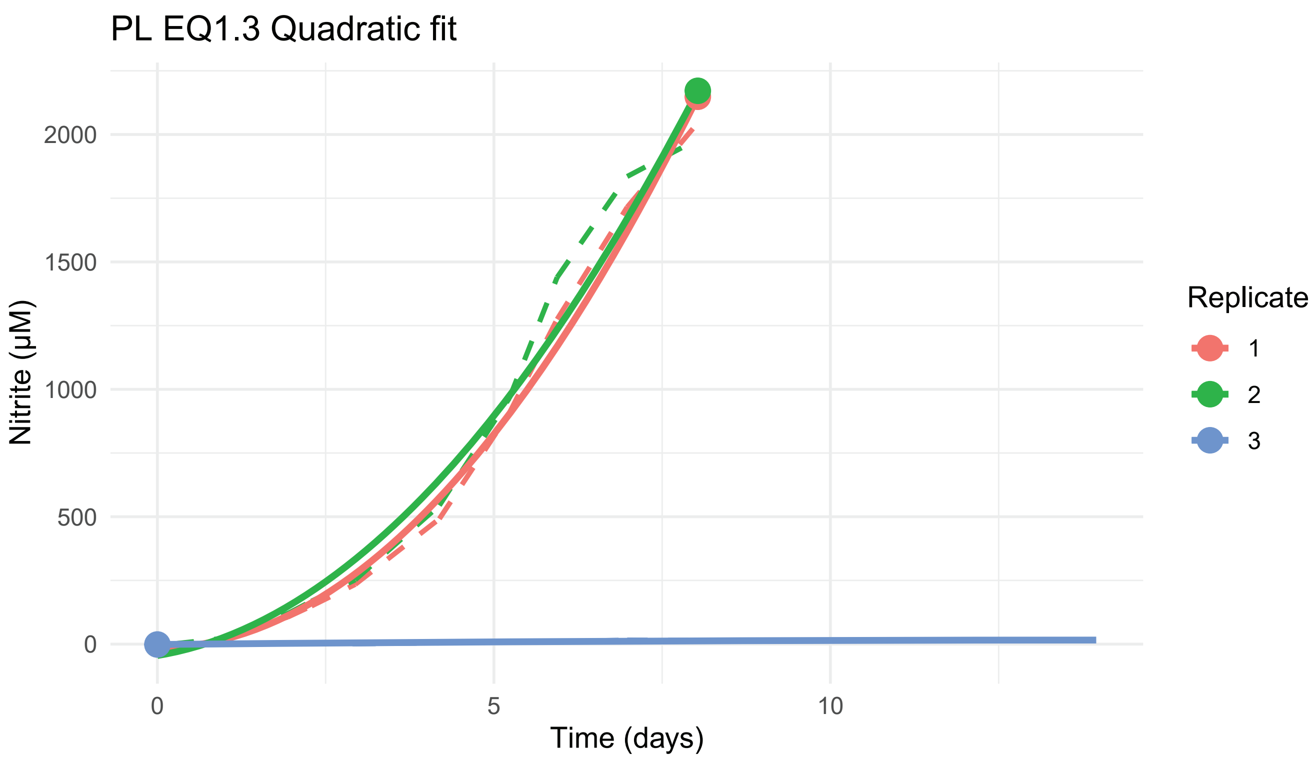
PL EQ1.3 Quadratic fit
Nitrite (µM)
2000
1500
1000
500
0
0
5
10
Time (days)
Replicate
1
2
3

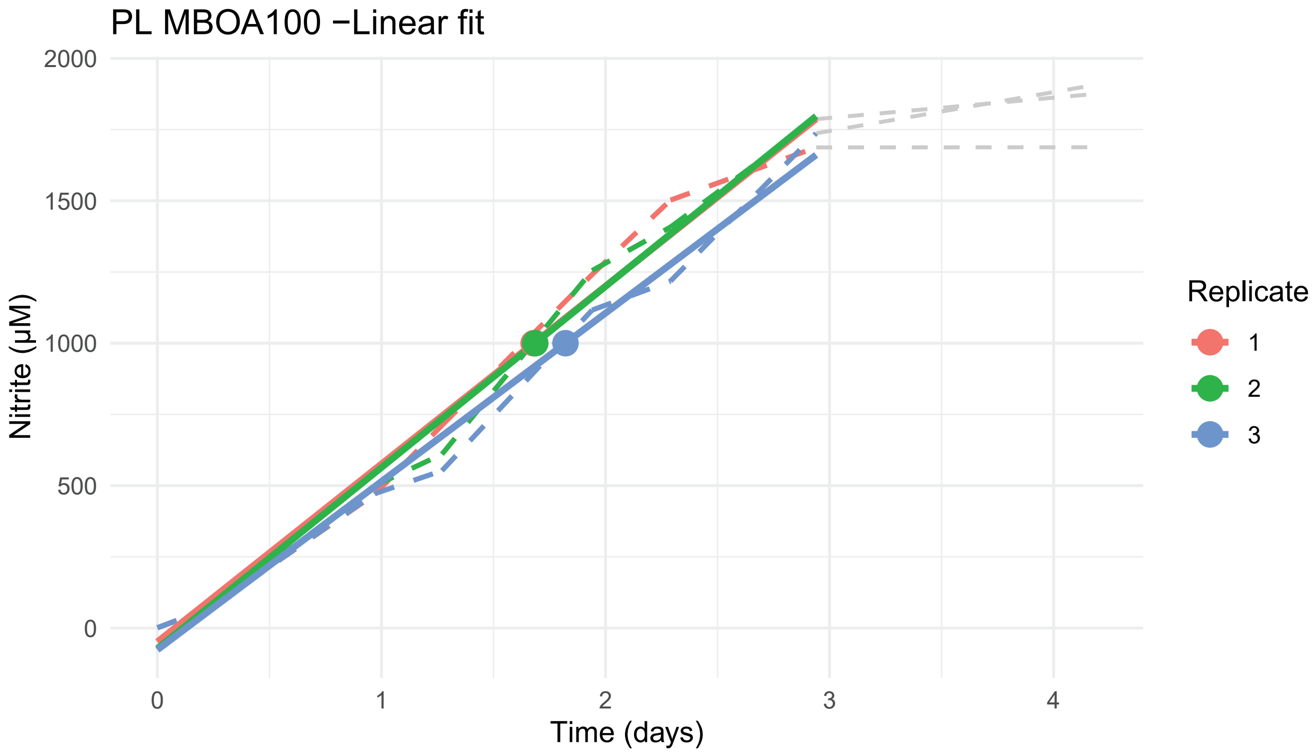
PL MBOA100 −Linear fit
Nitrite (µM)
2000
1500
1000
500
0
0
1
2
3
4
Time (days)
Replicate
1
2
3

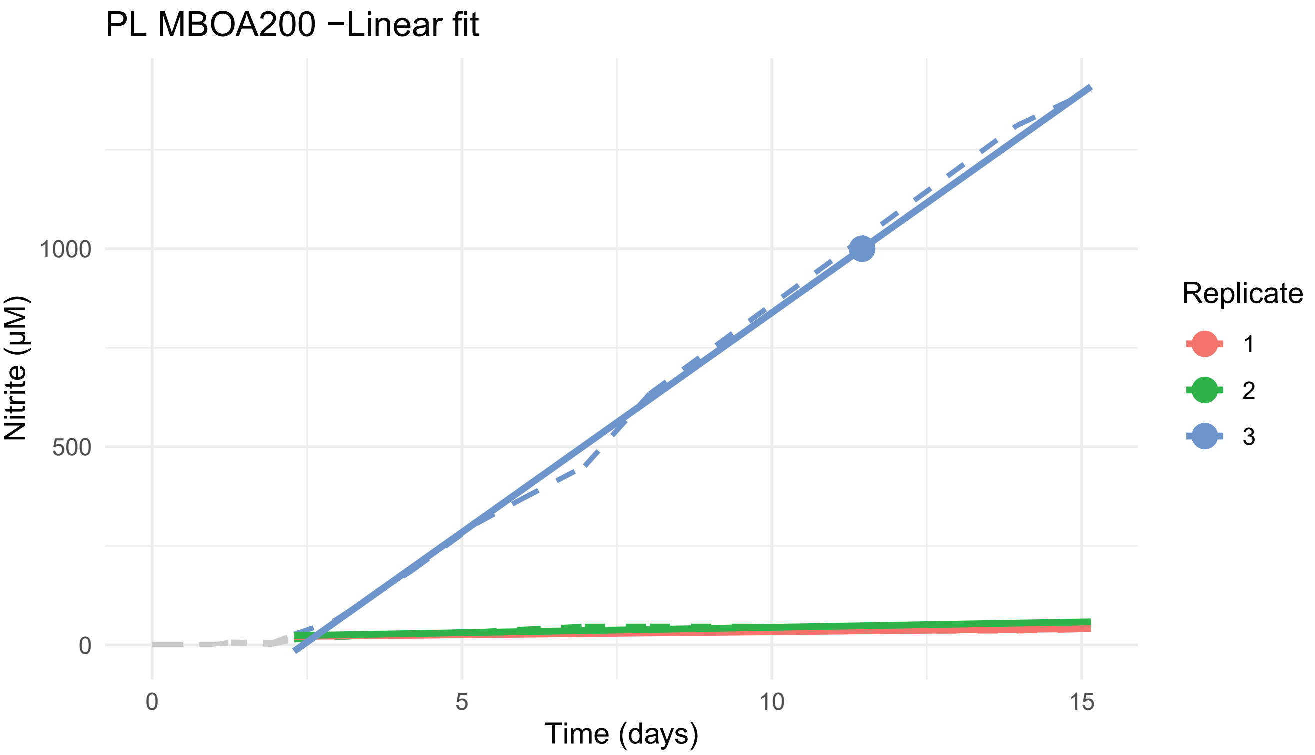
PL MBOA200 −Linear fit
Nitrite (μM)
0
500
1000
Time (days)
0
5
10
15
Replicate
1
2
3

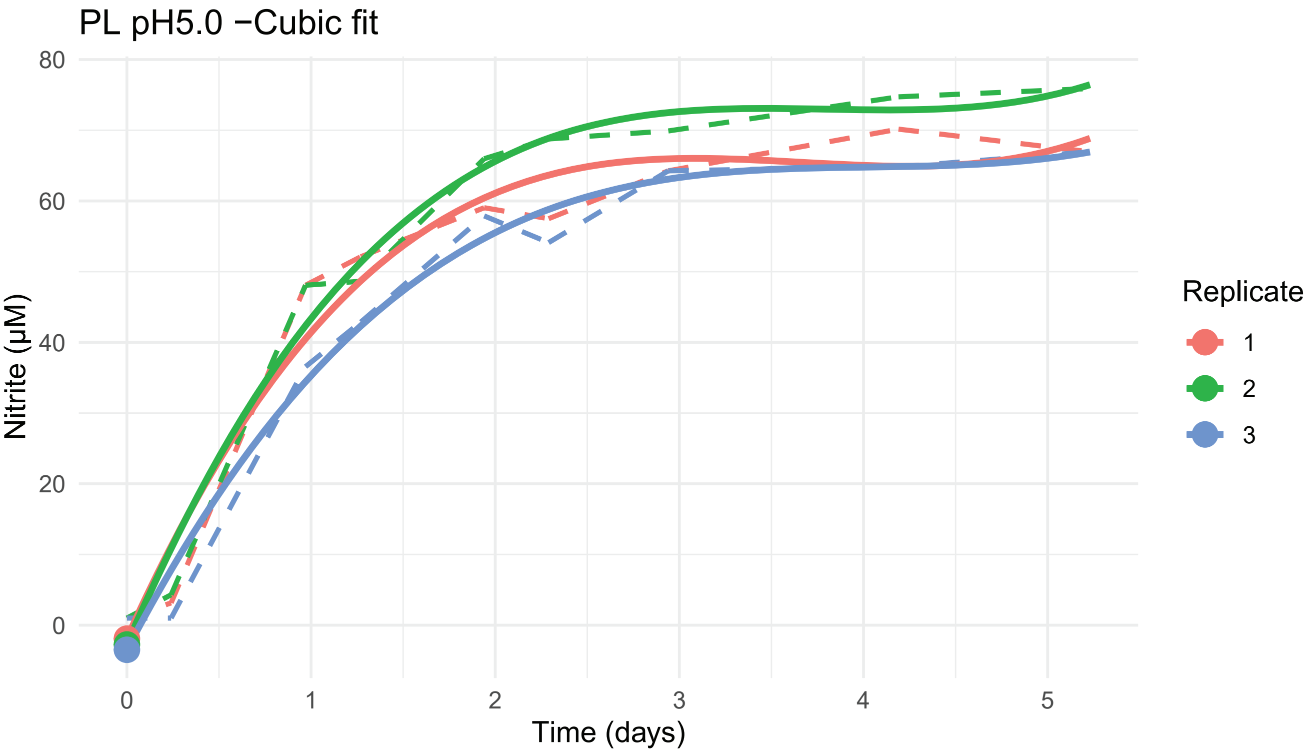
PL pH5.0 −Cubic fit
Nitrite (μM)
0
20
40
60
80
Time (days)
0
1
2
3
4
5
Replicate
1
2
3

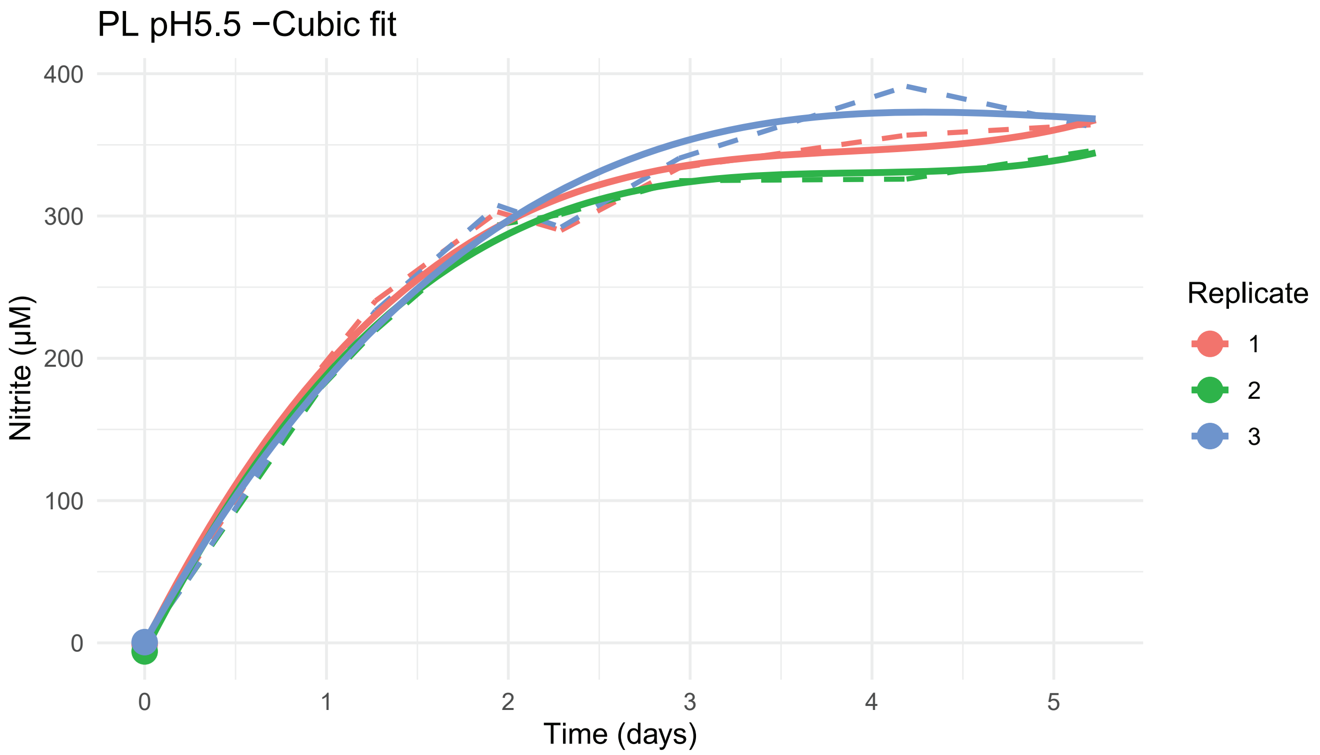
PL pH5.5 −Cubic fit
Nitrite (µM)
Time (days)
Replicate
1
2
3
0
100
200
300
400
0
1
2
3
4
5

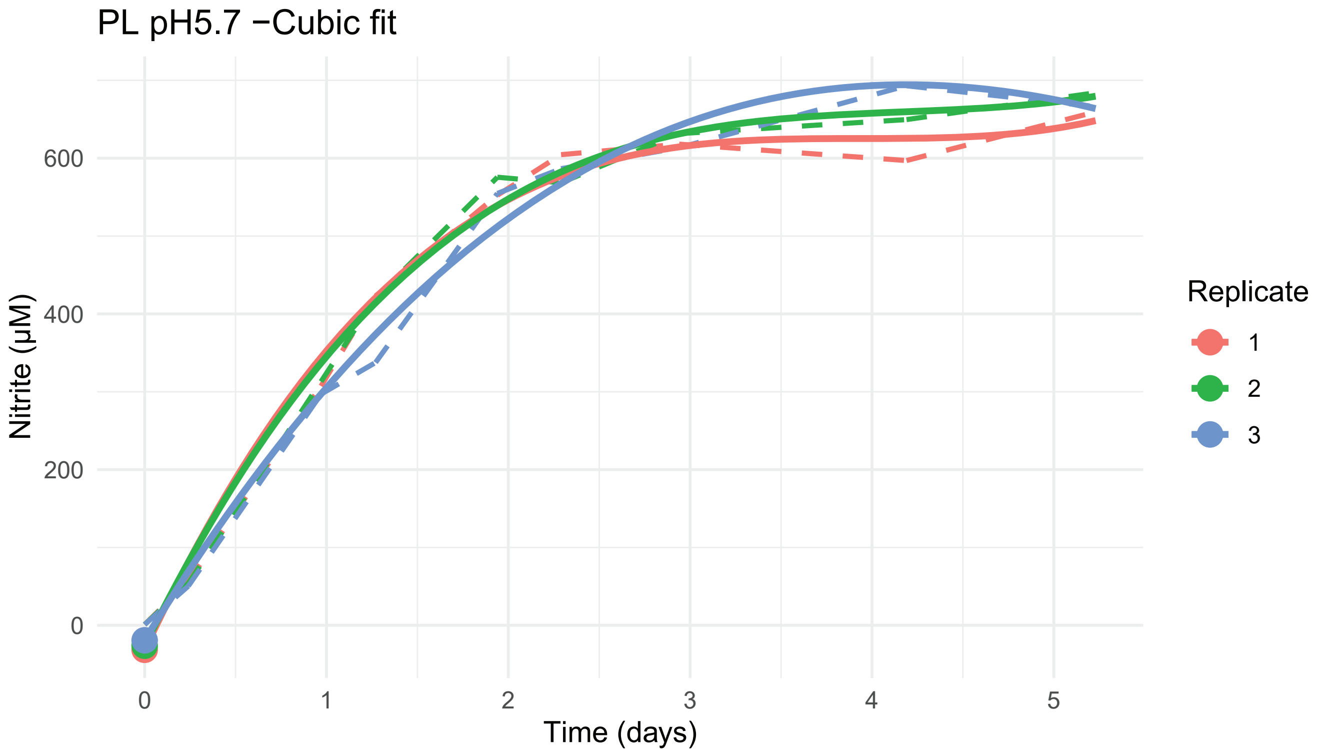
PL pH5.7 −Cubic fit
Nitrite (µM)
Time (days)
Replicate
1
2
3
0
200
400
600
0
1
2
3
4
5

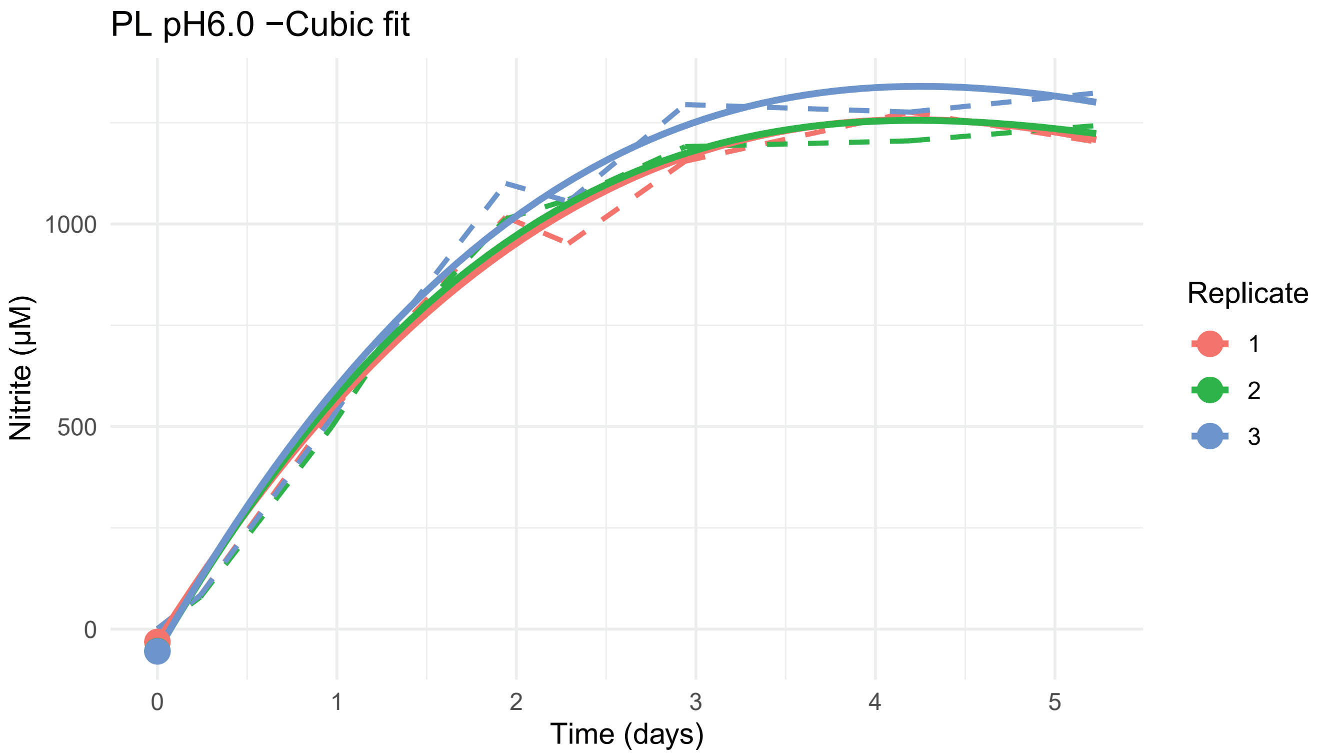
PL pH6.0 −Cubic fit
Nitrite (µM)
Time (days)
0
500
1000
0
1
2
3
4
5
Replicate
1
2
3

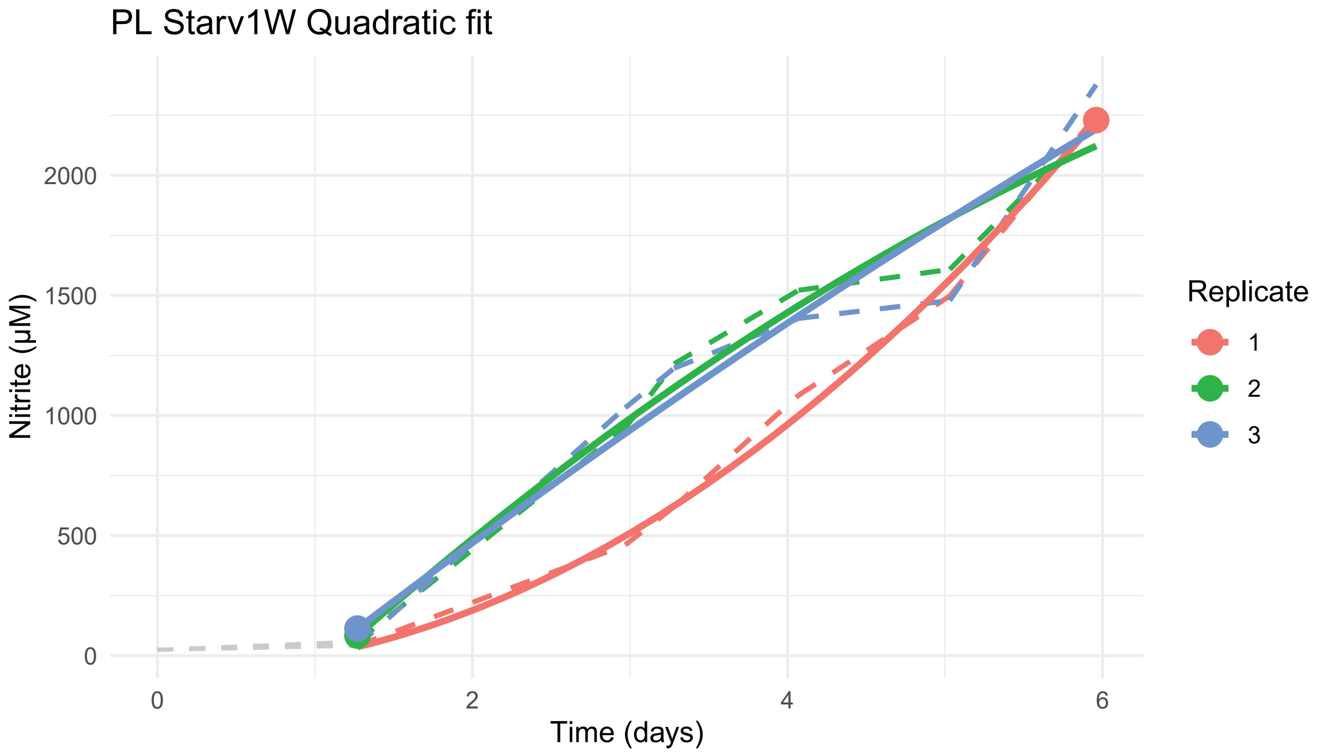
PL Starv1W Quadratic fit
Nitrite (µM)
Time (days)
0
500
1000
1500
2000
0
2
4
6
Replicate
1
2
3

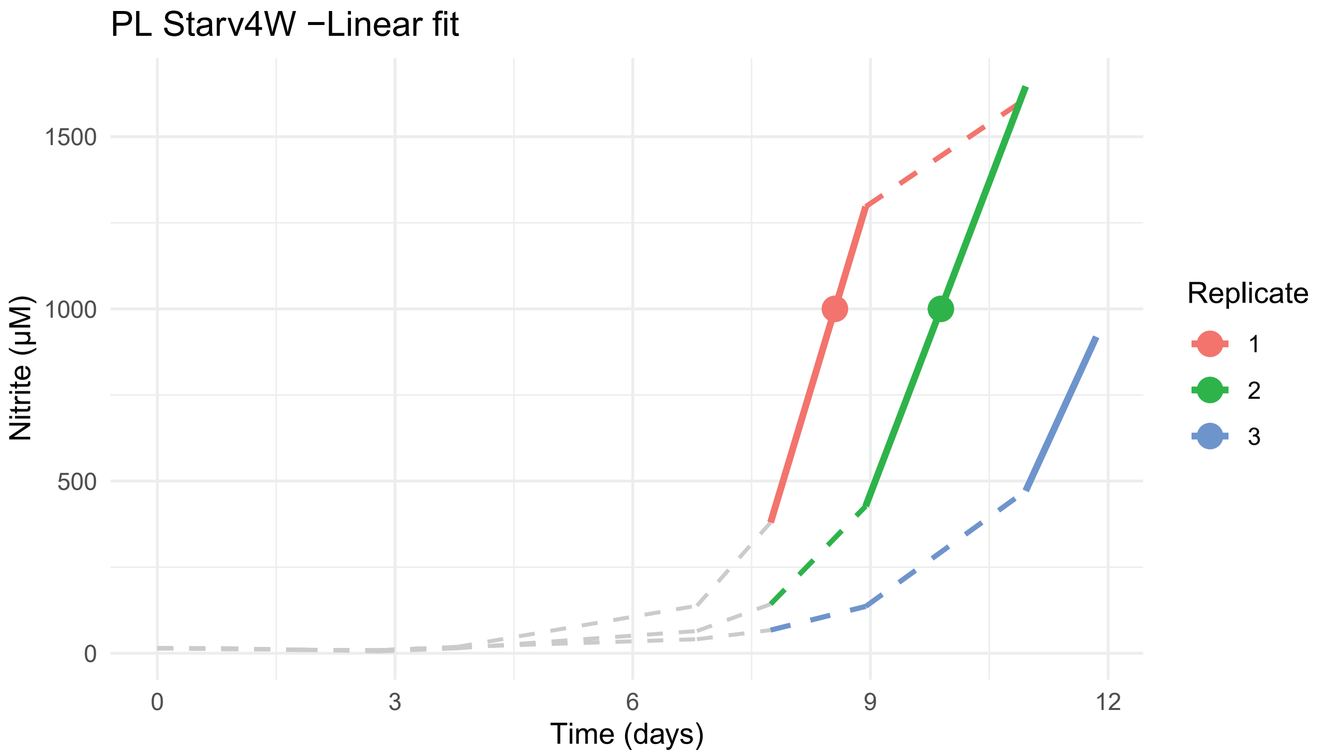
PL Starv4W −Linear fit
Nitrite (µM)
1500
1000
500
0
0
3
6
9
12
Time (days)
Replicate
1
2
3
