## Supplementary Material S3 for "Ecophysiological niche expansion driven by biofilm lifestyle in an archaeal soil nitrifier"

**Models of Compared Growth Curves**

Each graph represents one condition. Condition and the chosen model (linear, quadratic, or cubic) are indicated in the figure titles. Dotted black lines represent growth curves. Solid colored lines represent fitted models. Models only cover regions of the growth curve for which points were used. Large dark green points represent the point in the model of the maximum rate ($n_{max}$). If needed, dashed red line represents the nitrite level for which ($n_{max}$) was estimated.

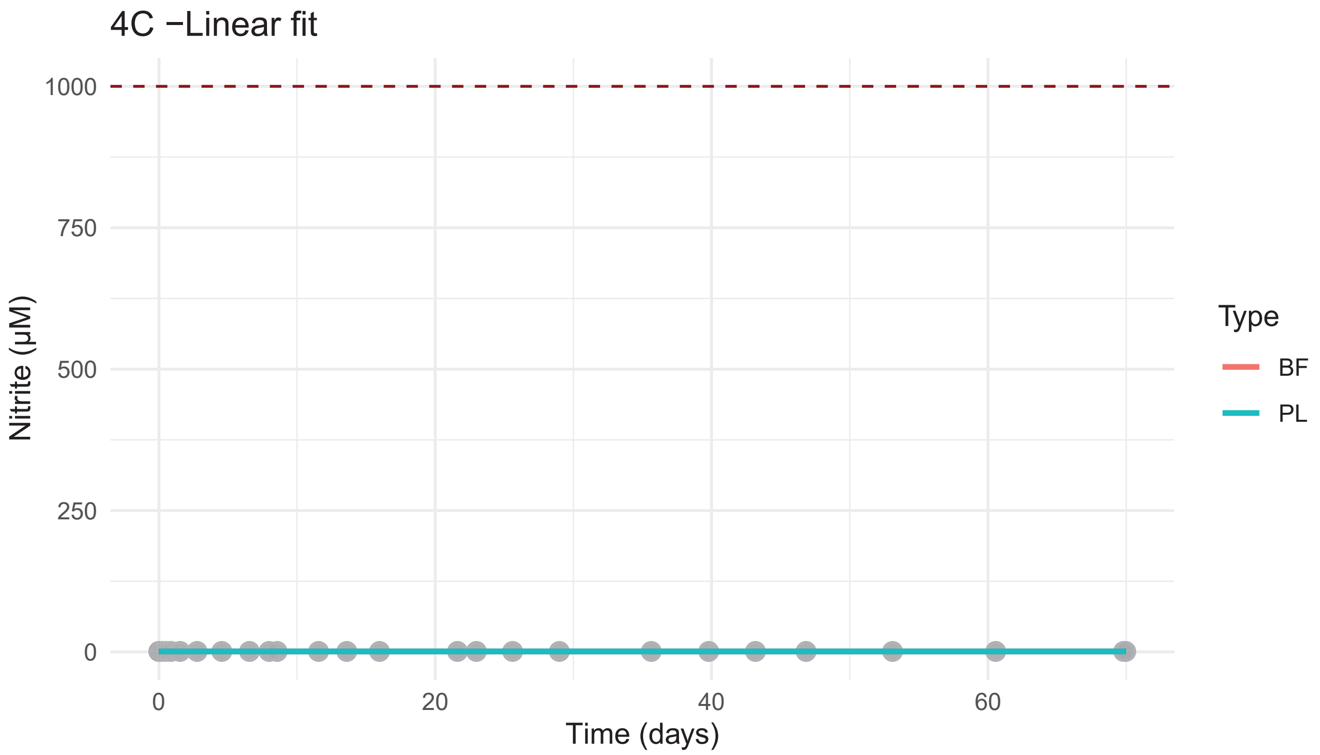

4C −Linear fit
1000
750
500
250
0
Nitrite (μM)
0
20
40
60
Time (days)
Type
BF
PL


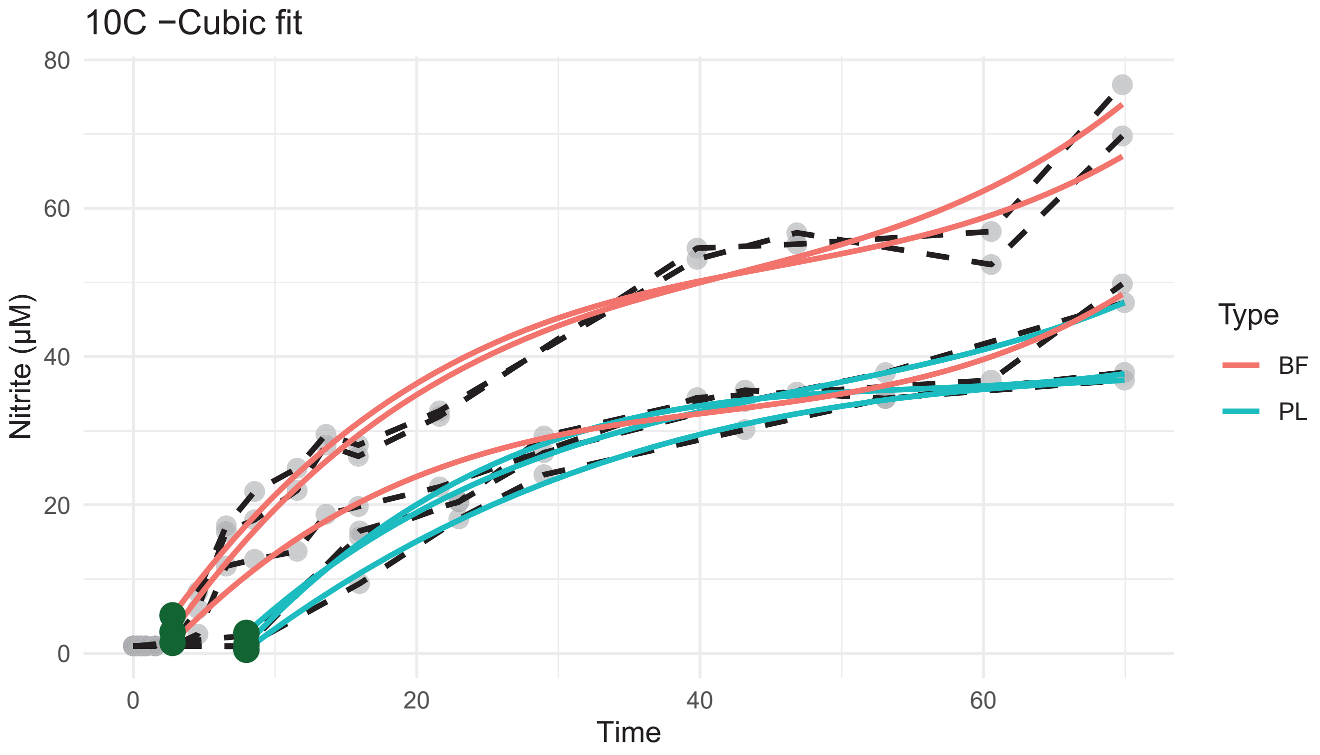

10C −Cubic fit
80
60
40
20
0
Nitrite (μM)
0
20
40
60
Time
Type
BF
PL

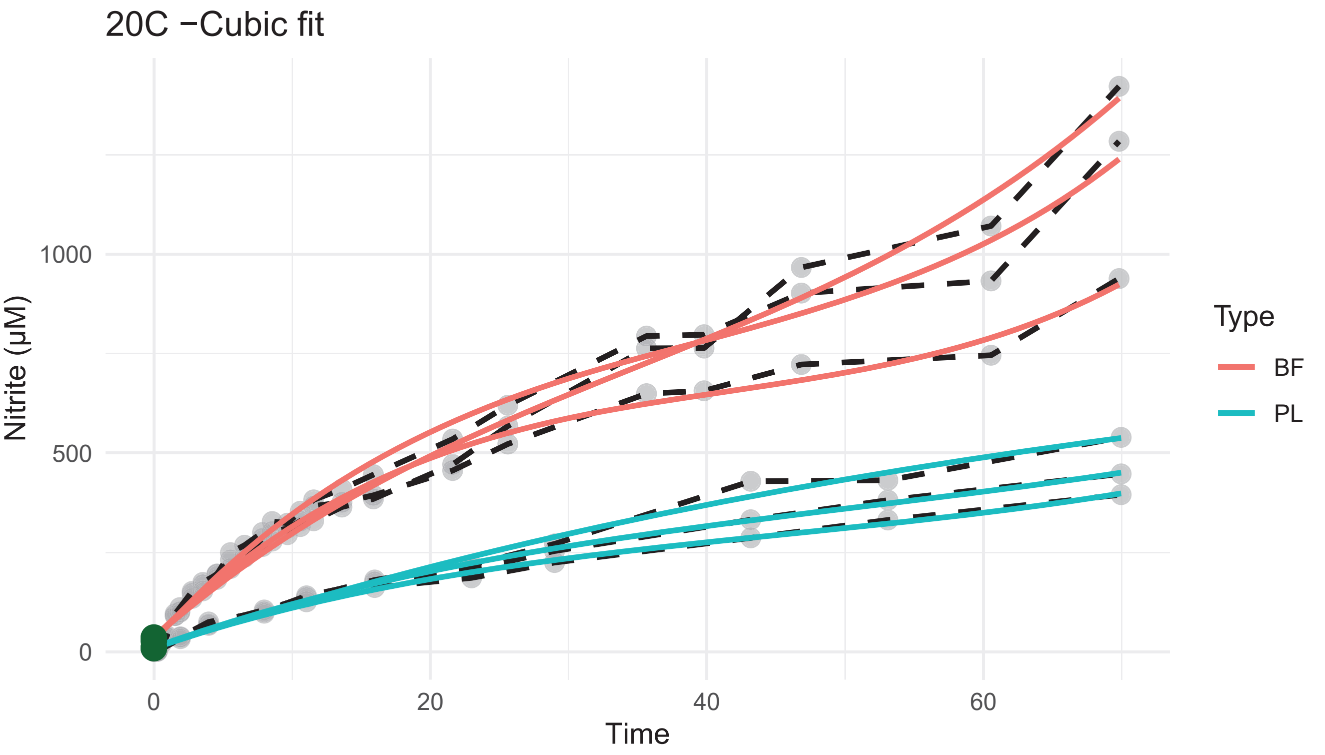
20C −Cubic fit
Nitrite (µM)
Time
Type
BF
PL
0
500
1000
0
20
40
60

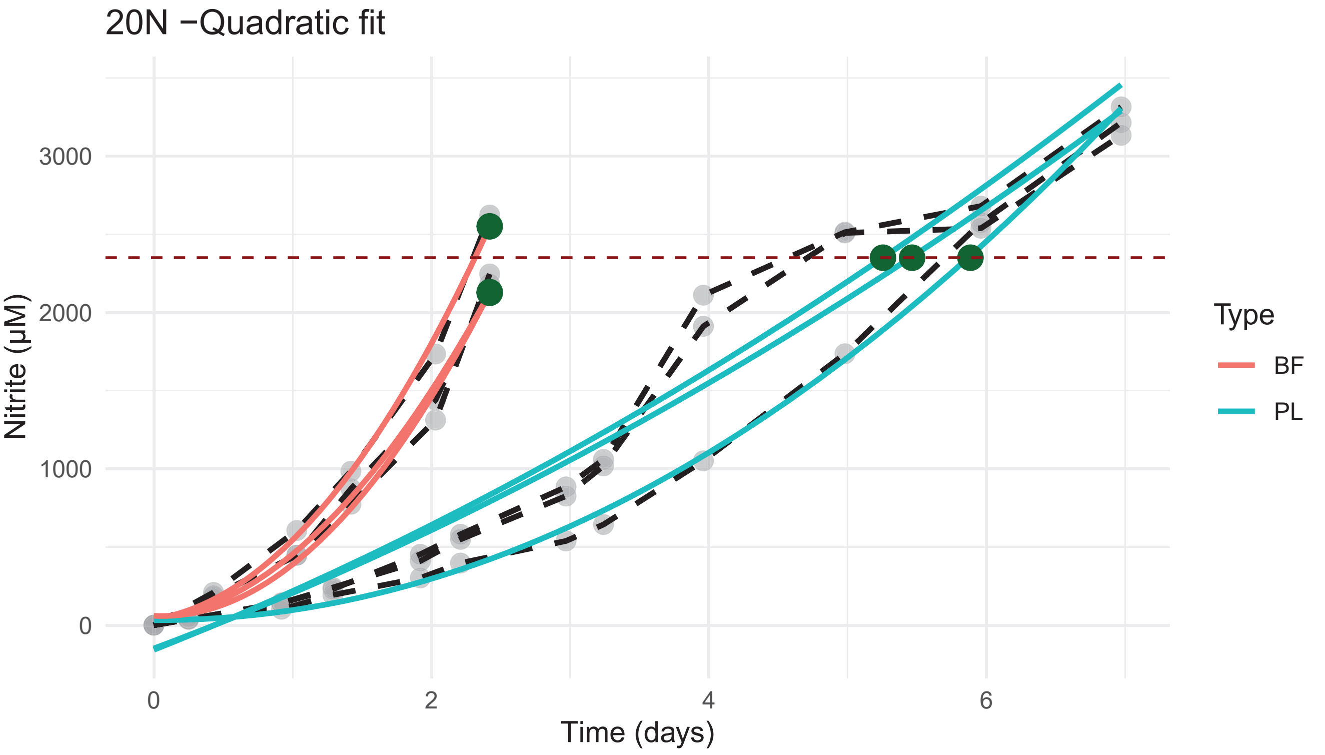
20N −Quadratic fit
Nitrite (µM)
Time (days)
Type
BF
PL
0
1000
2000
3000
0
2
4
6

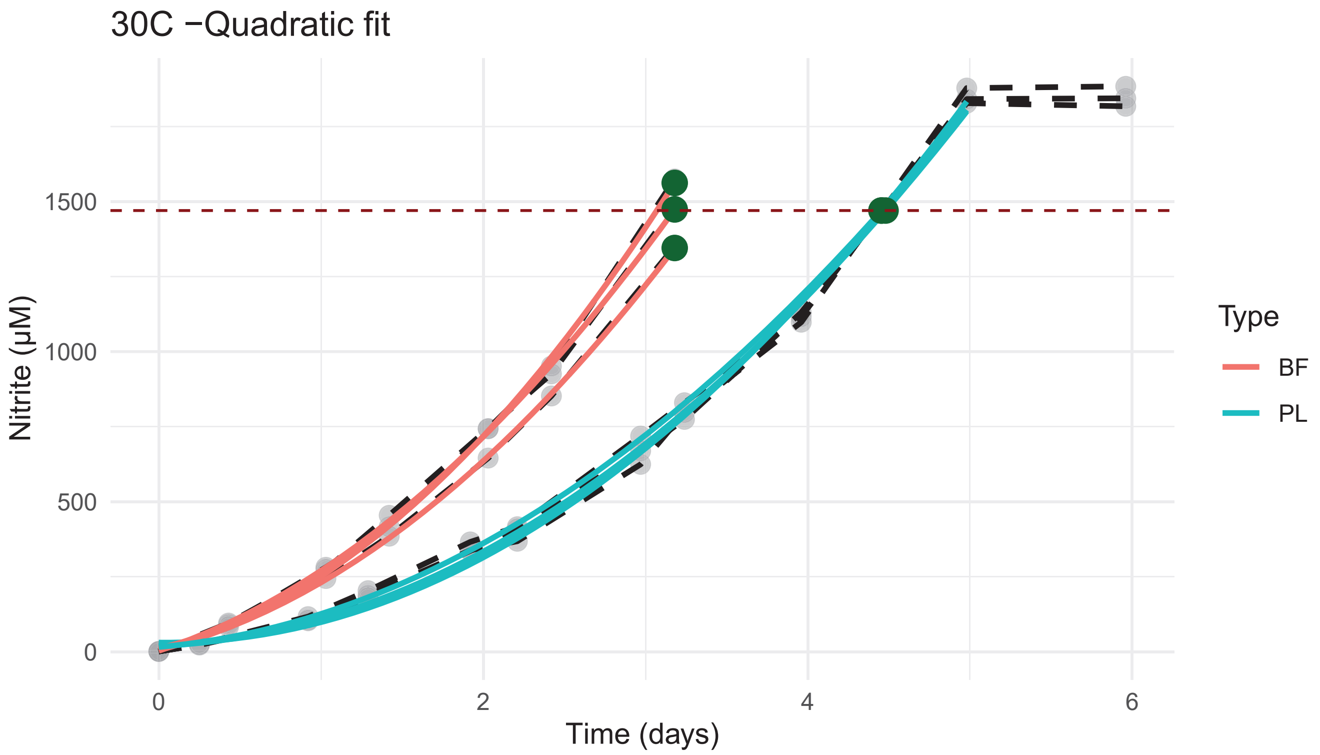
30C −Quadratic fit
Nitrite (μM)
1500
1000
500
0
0
2
4
6
Time (days)
Type
BF
PL

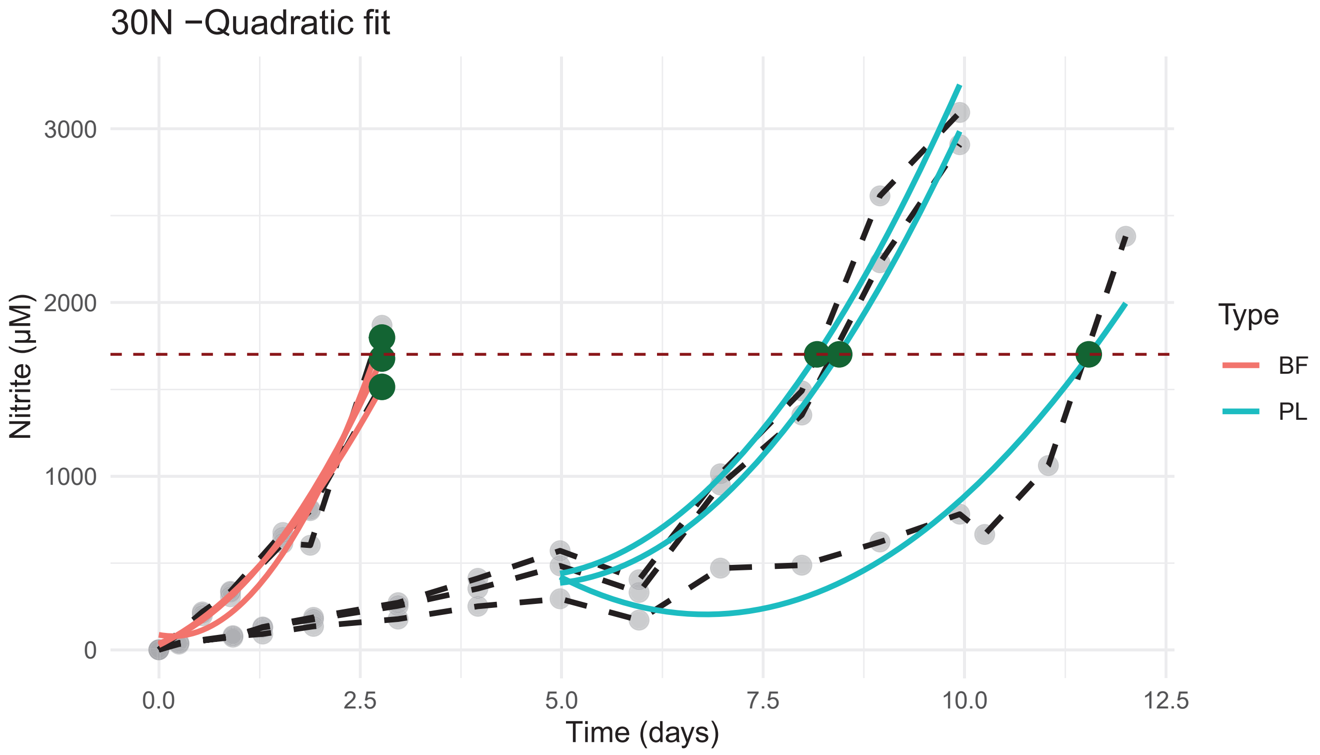
30N −Quadratic fit
Nitrite (μM)
3000
2000
1000
0
0.0
2.5
5.0
7.5
10.0
12.5
Time (days)
Type
BF
PL

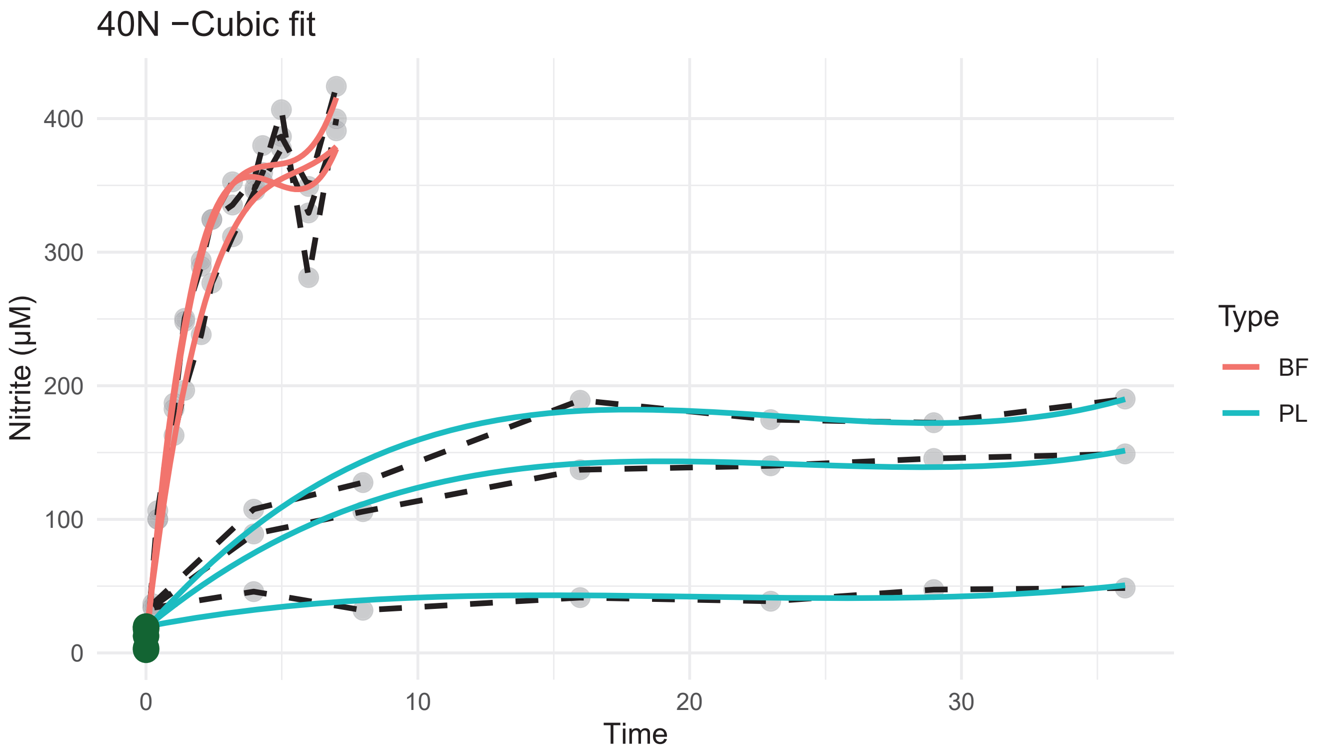
40N −Cubic fit
Nitrite (μM)
0
100
200
300
400
0
10
20
30
Time
Type
BF
PL

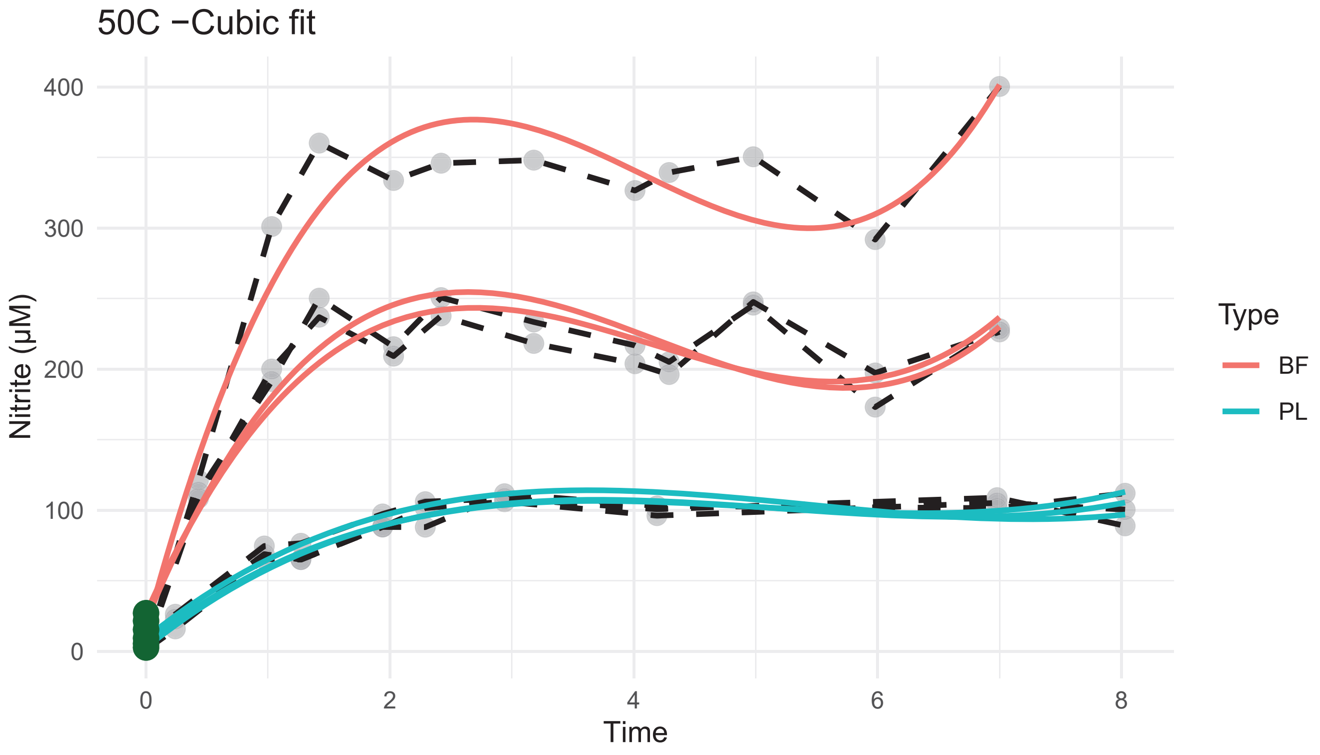
50C −Cubic fit
Nitrite (μM)
0
100
200
300
400
0
2
4
6
8
Time
Type
BF
PL

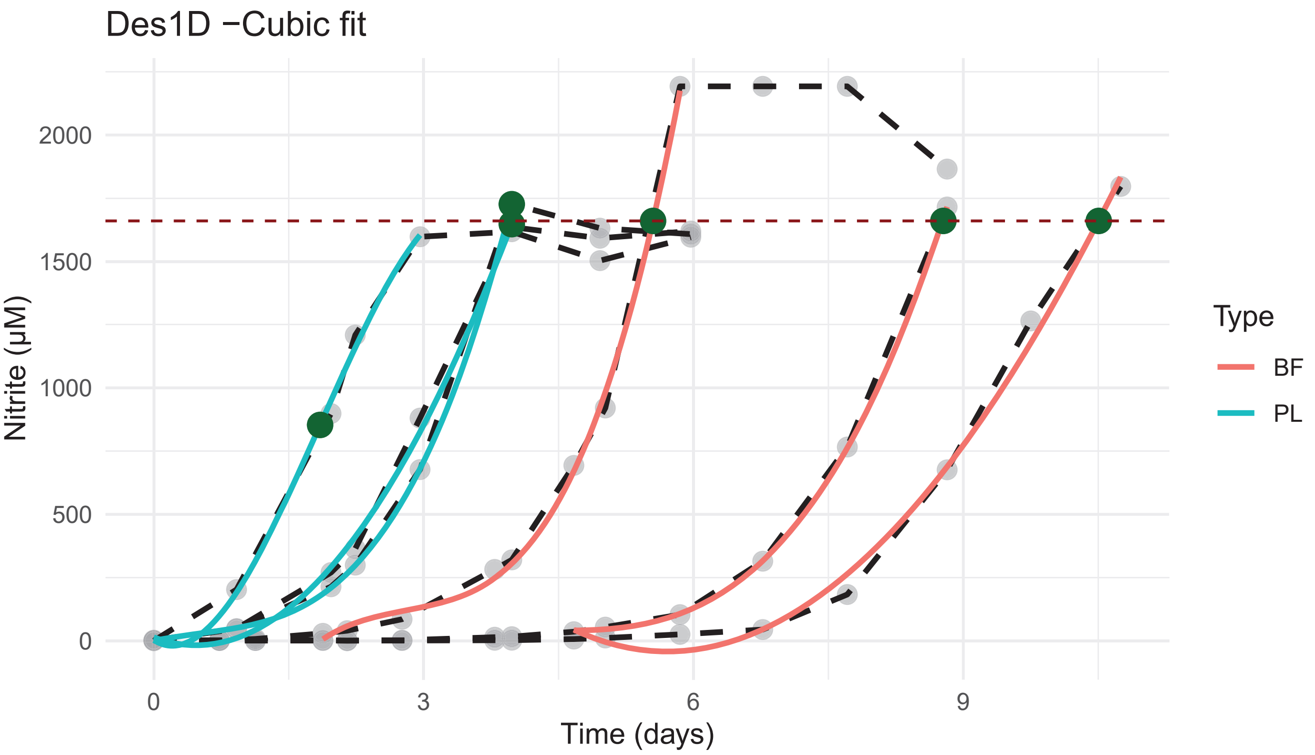
Des1D −Cubic fit
Nitrite (µM)
2000
1500
1000
500
0
0
3
6
9
Time (days)
Type
BF
PL

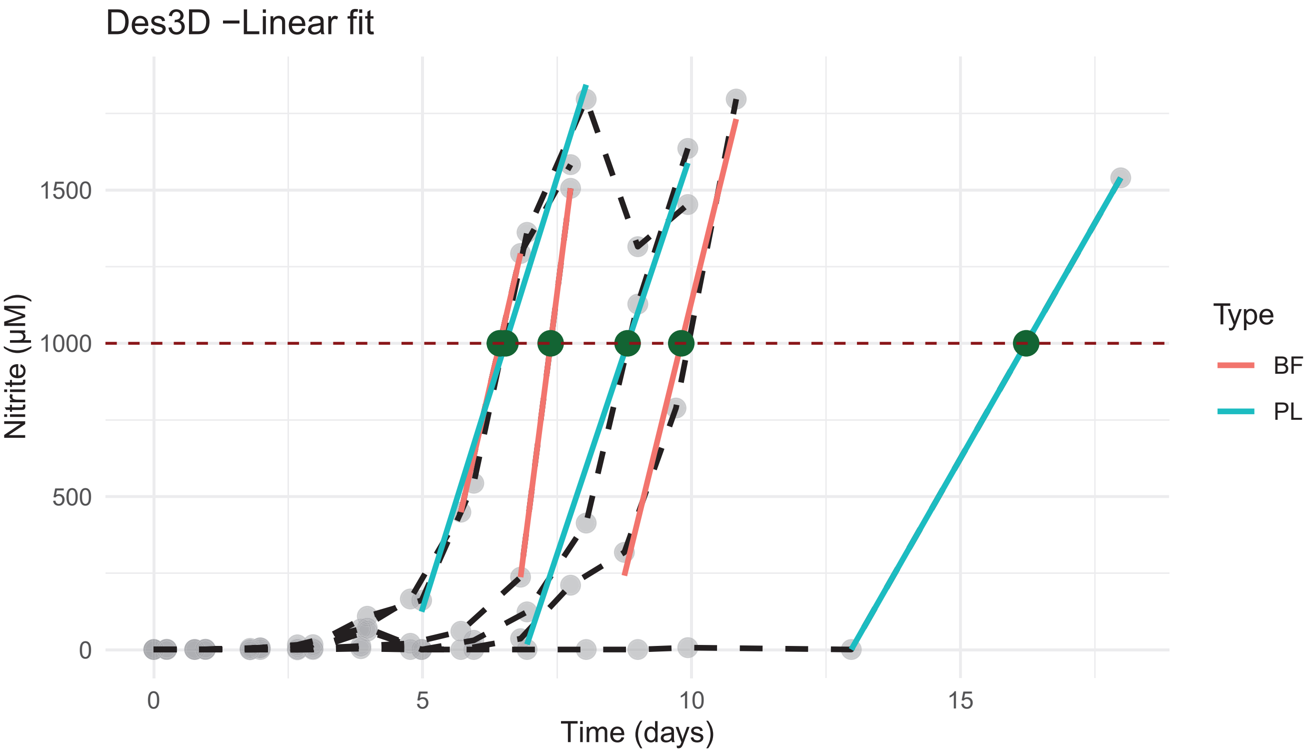
Des3D −Linear fit
Nitrite (µM)
1500
1000
500
0
0
5
10
15
Time (days)
Type
BF
PL

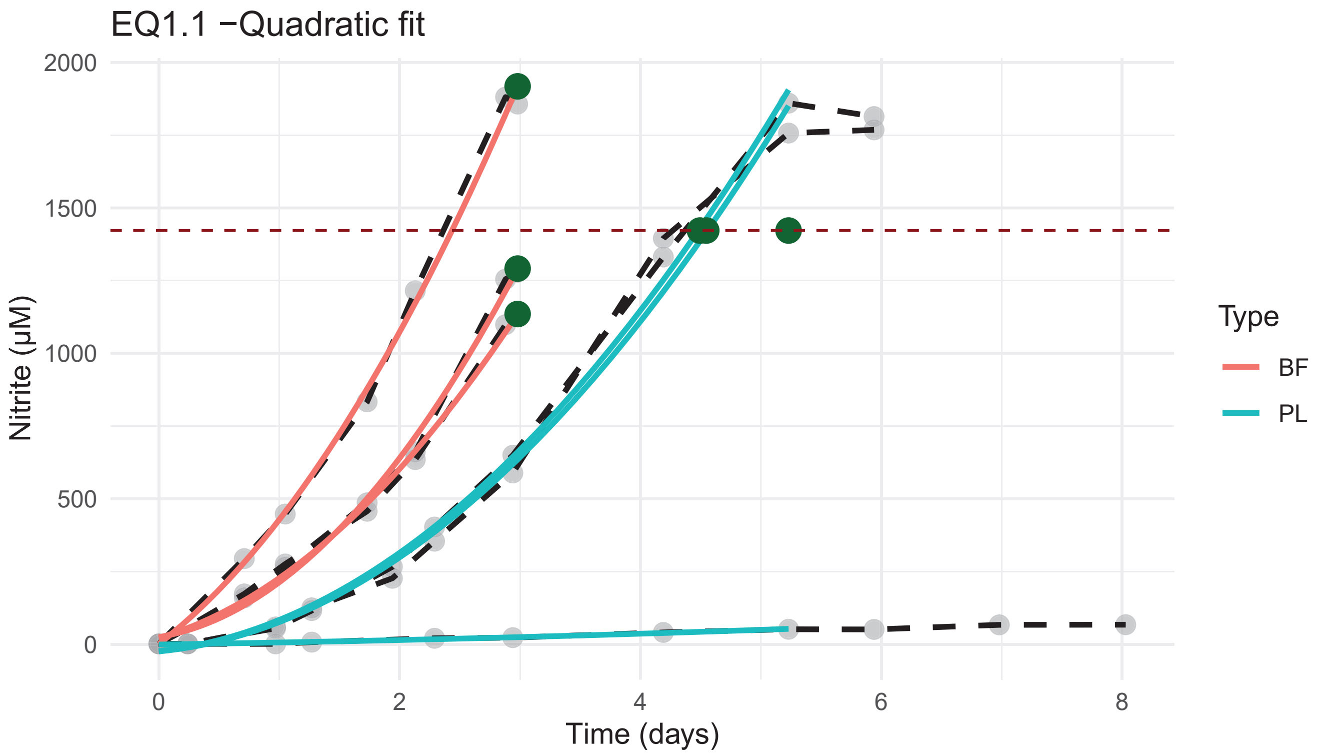
EQ1.1 −Quadratic fit
2000
1500
1000
500
0
Nitrite (µM)
0
2
4
6
8
Time (days)
Type
BF
PL

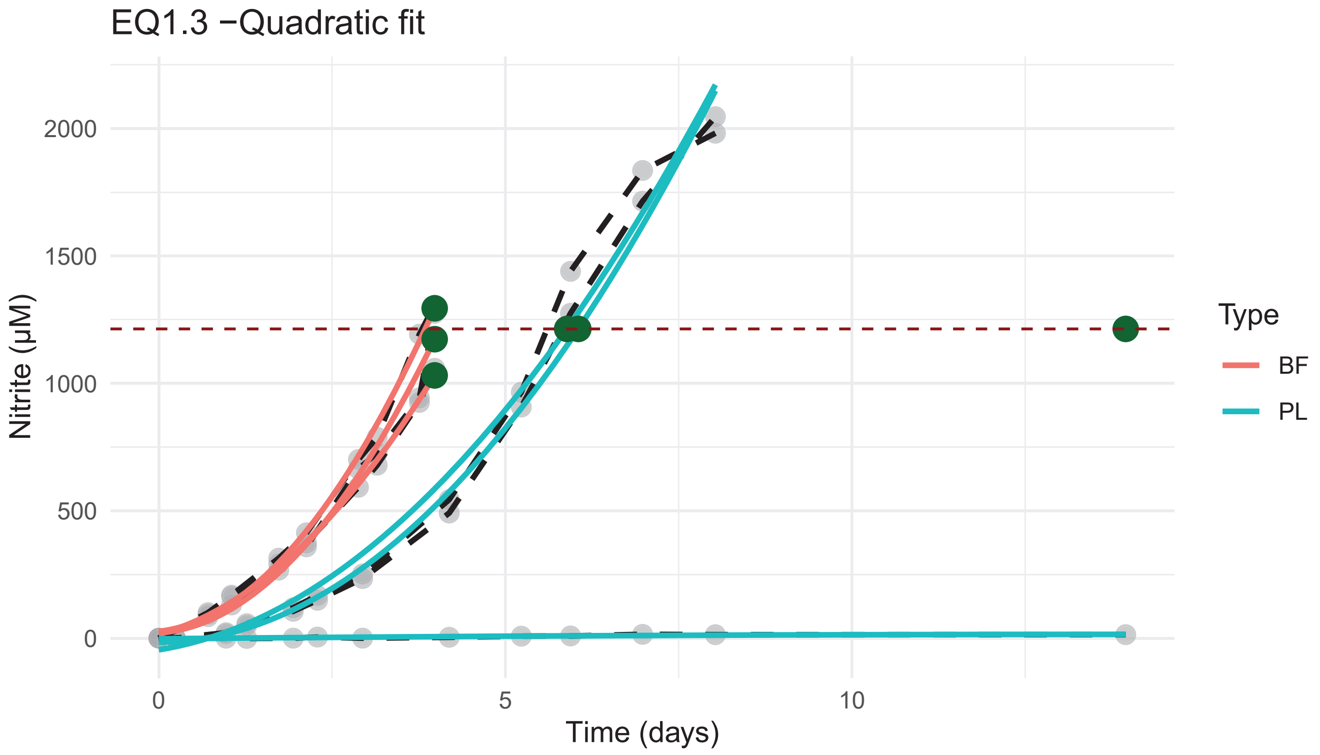
EQ1.3 −Quadratic fit
2000
1500
1000
500
0
Nitrite (µM)
0
5
10
Time (days)
Type
BF
PL

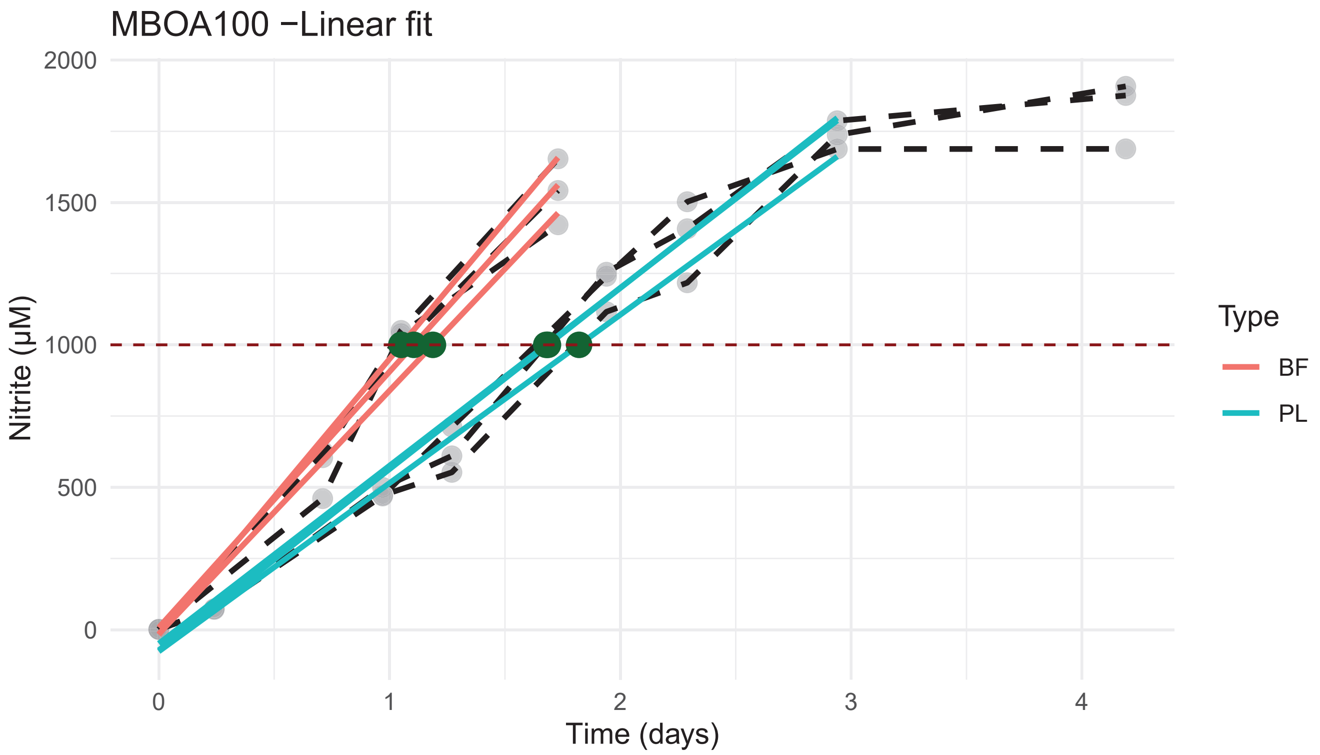
MBOA100 −Linear fit
Nitrite (µM)
2000
1500
1000
500
0
0
1
2
3
4
Time (days)
Type
BF
PL

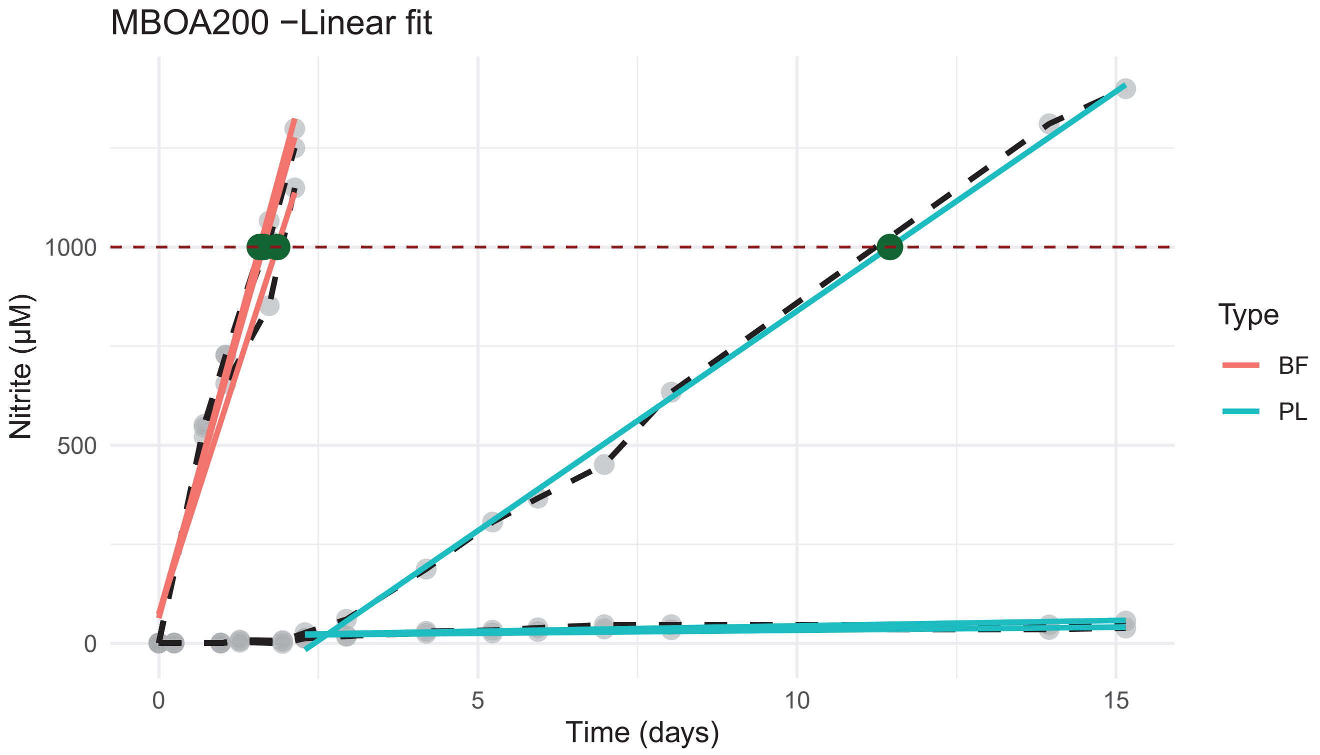
MBOA200 −Linear fit
Nitrite (µM)
1000
500
0
0
5
10
15
Time (days)
Type
BF
PL

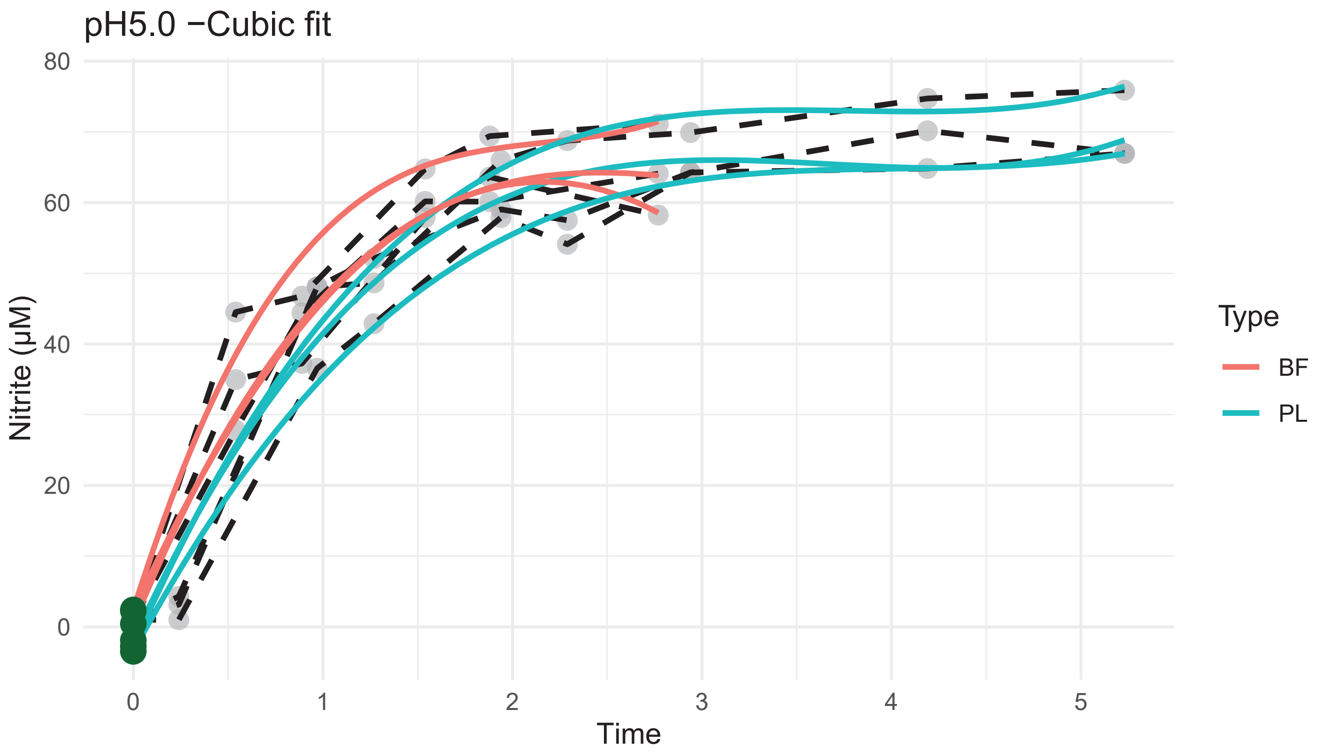
pH5.0 −Cubic fit
80
60
40
20
0
Nitrite (µM)
0
1
2
3
4
5
Time
Type
BF
PL

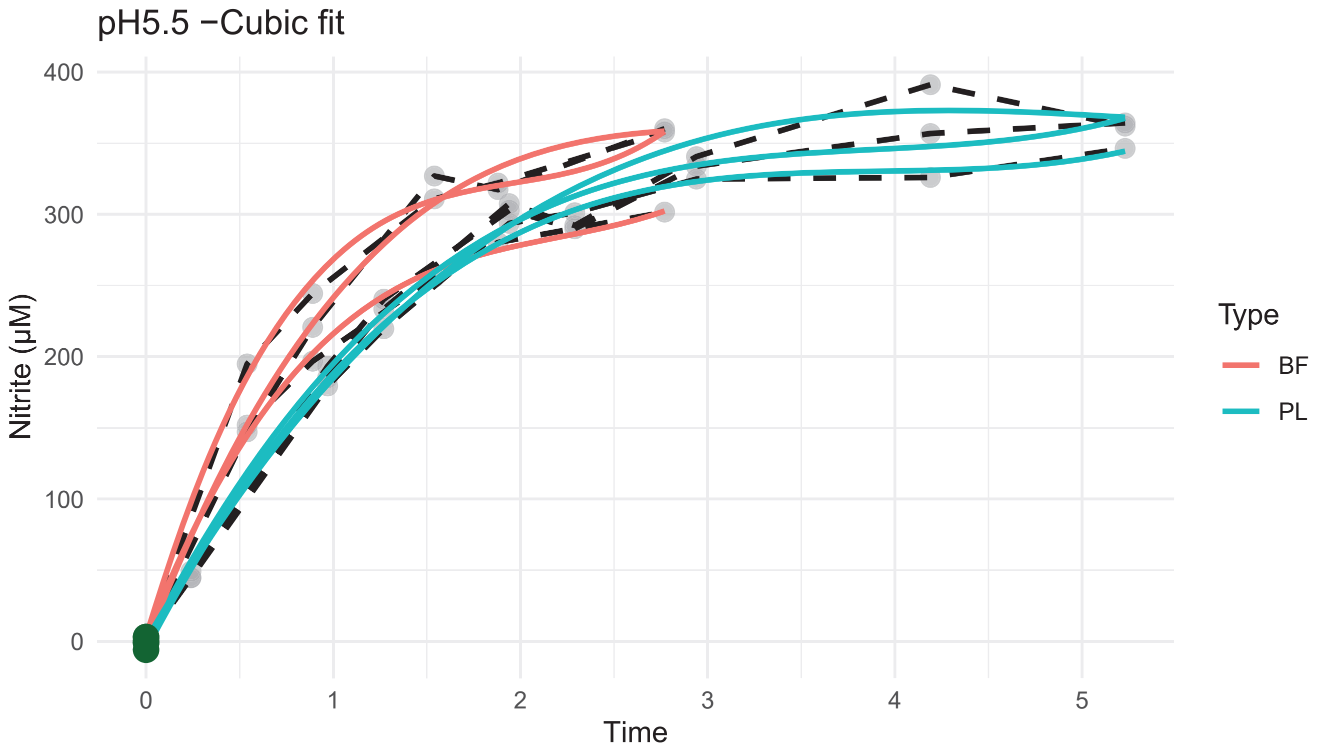
pH5.5 −Cubic fit
400
300
200
100
0
Nitrite (µM)
0
1
2
3
4
5
Time
Type
BF
PL

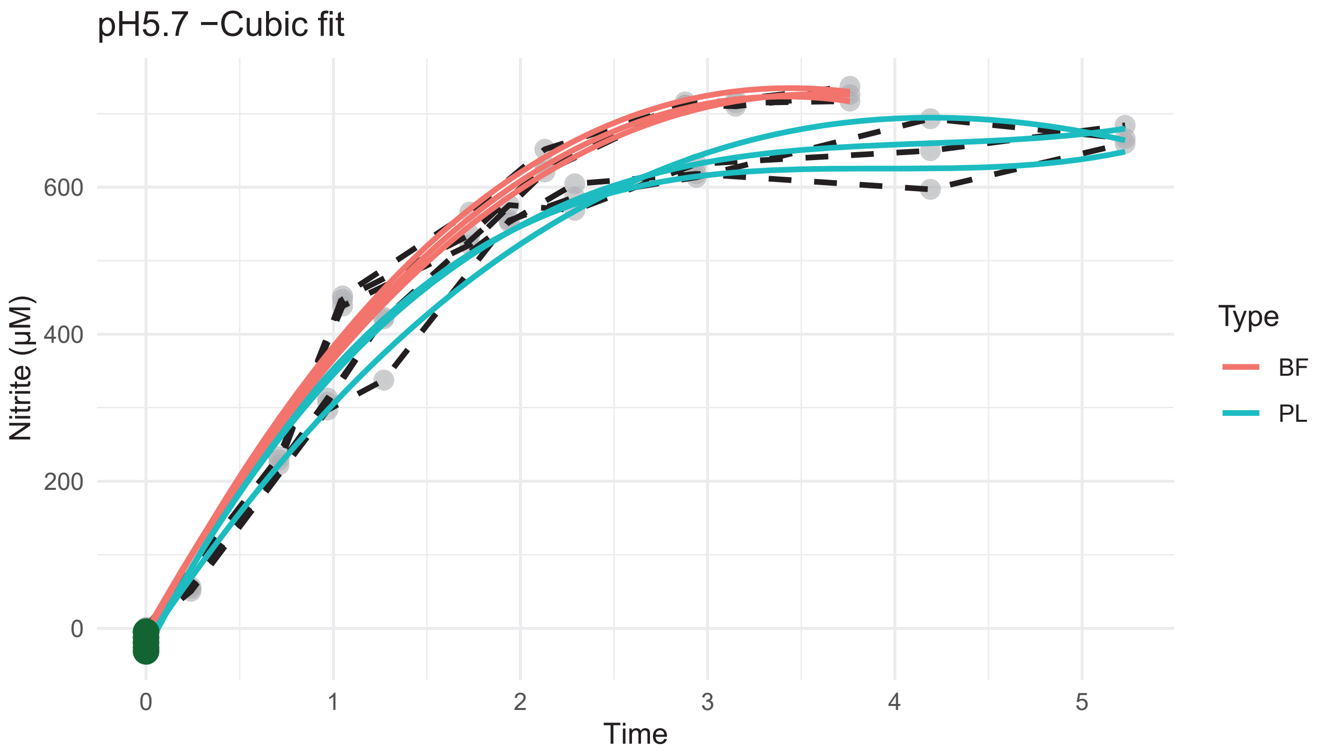
pH5.7 −Cubic fit
Nitrite (µM)
Time
0
200
400
600
0
1
2
3
4
5
Type
BF
PL

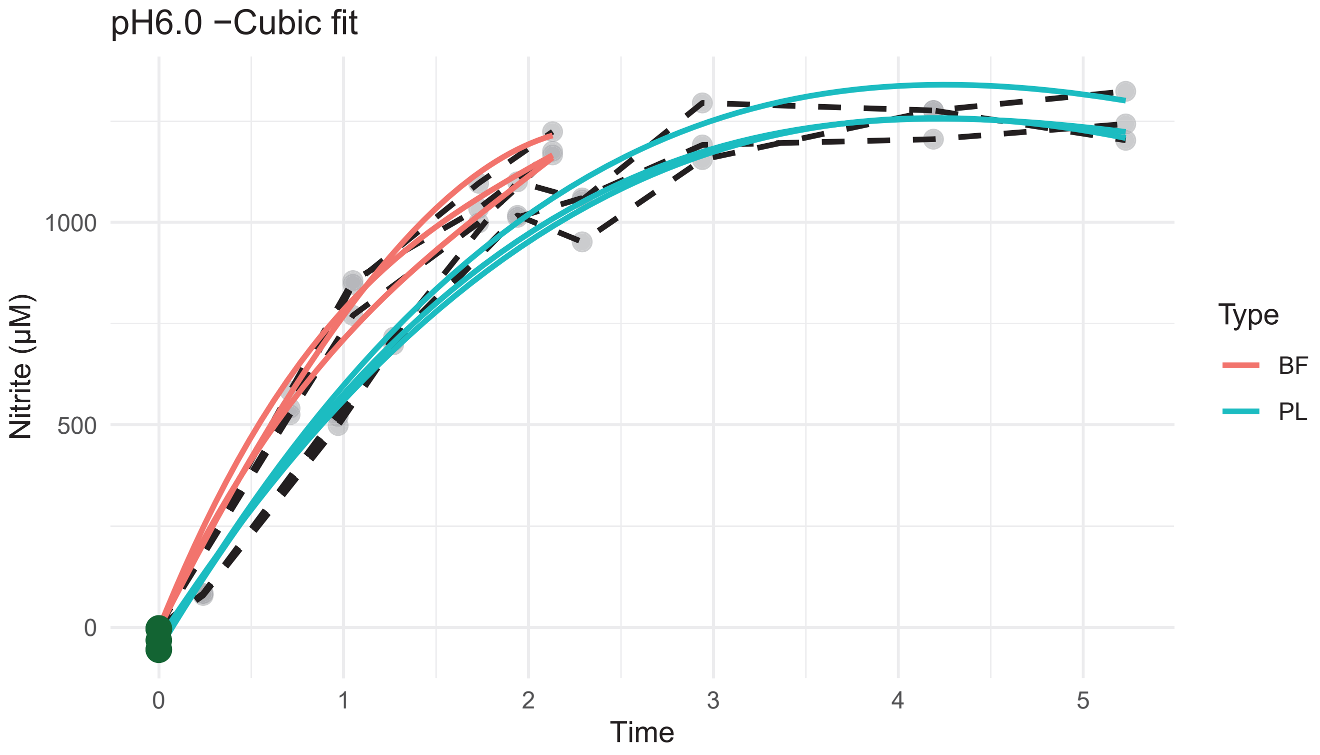
pH6.0 −Cubic fit
Nitrite (µM)
Time
0
500
1000
0
1
2
3
4
5
Type
BF
PL

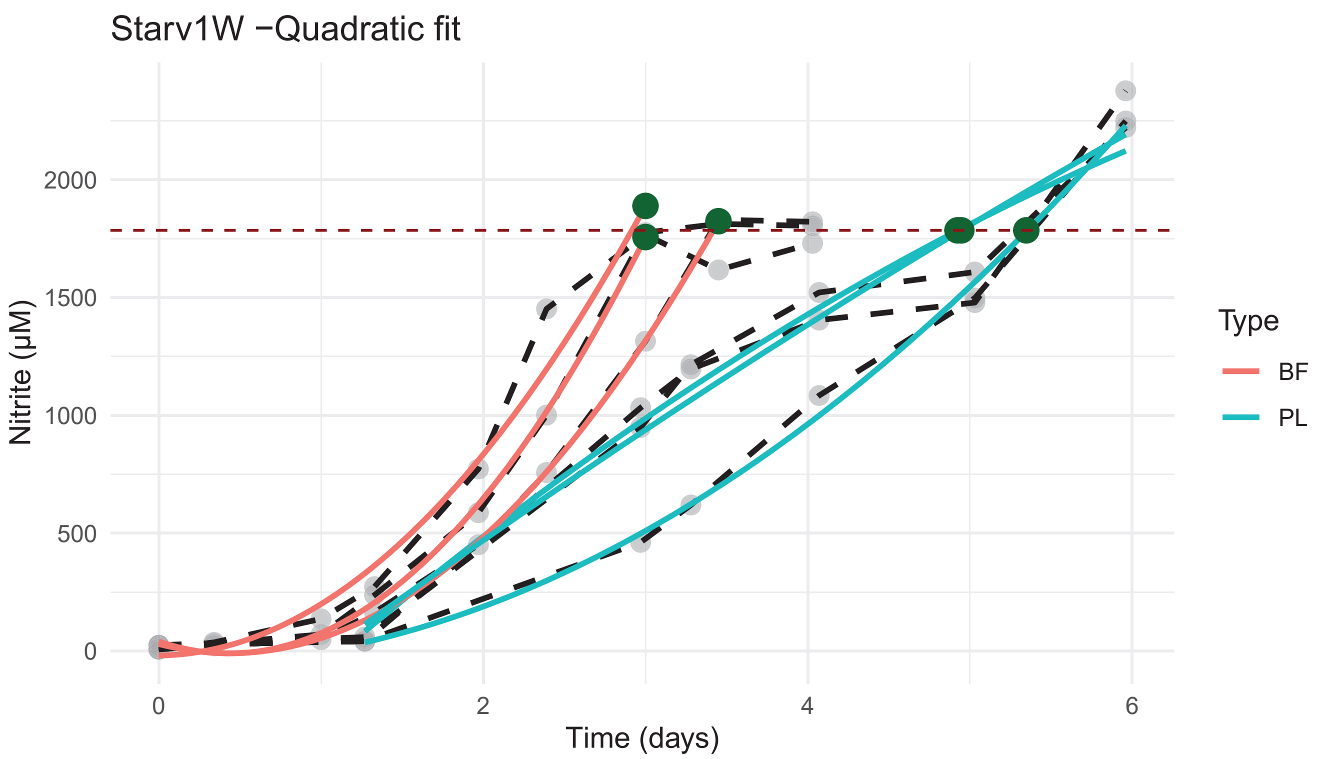

Starv1W −Quadratic fit
Nitrite (µM)
2000
1500
1000
500
0
0
2
4
6
Time (days)
Type
BF
PL


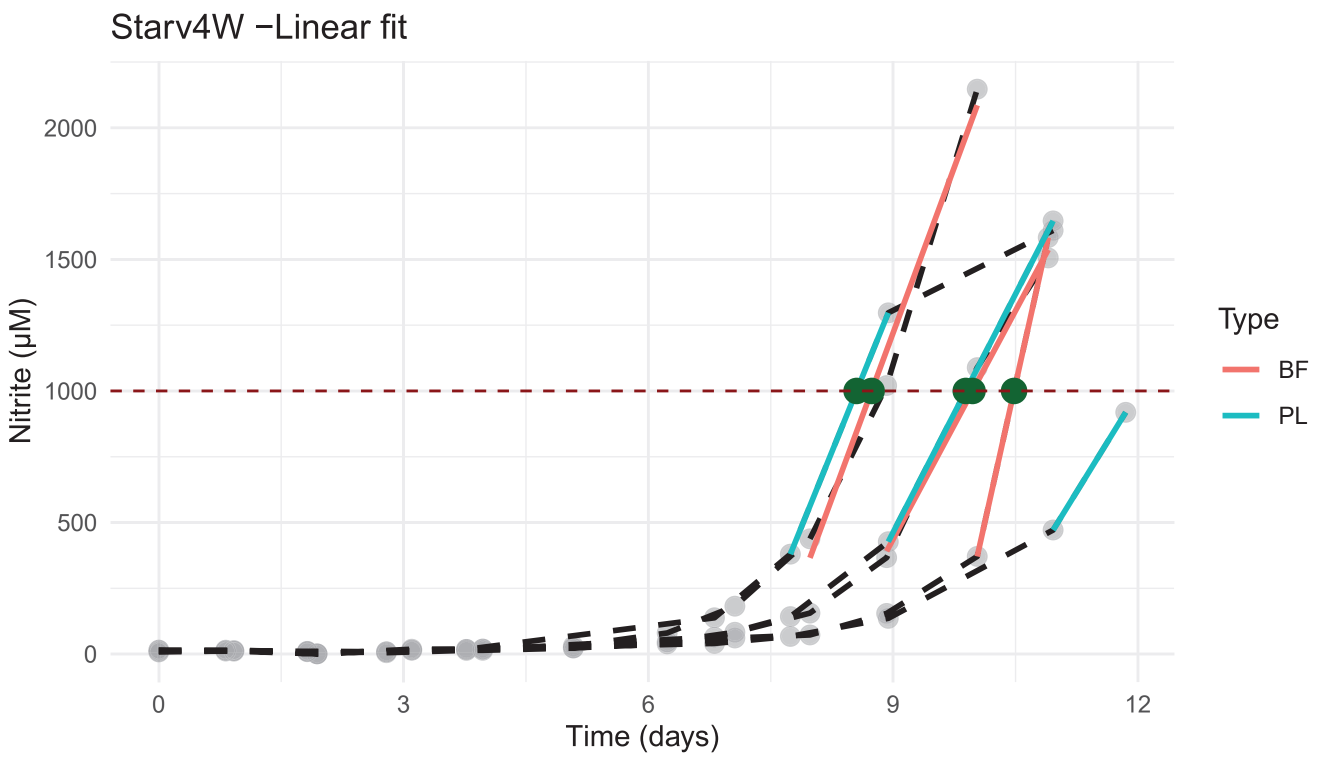

Starv4W −Linear fit
Nitrite (µM)
2000
1500
1000
500
0
0
3
6
9
12
Time (days)
Type
BF
PL
